# Genome wide meta-analysis identifies genomic relationships, novel loci, and pleiotropic mechanisms across eight psychiatric disorders

**DOI:** 10.1101/528117

**Authors:** Cross-Disorder Group of the Psychiatric Genomics Consortium, Jordan W. Smoller

## Abstract

Genetic influences on psychiatric disorders transcend diagnostic boundaries, suggesting substantial pleiotropy of contributing loci. However, the nature and mechanisms of these pleiotropic effects remain unclear. We performed a meta-analysis of 232,964 cases and 494,162 controls from genome-wide studies of anorexia nervosa, attention-deficit/hyperactivity disorder, autism spectrum disorder, bipolar disorder, major depression, obsessive-compulsive disorder, schizophrenia, and Tourette syndrome. Genetic correlation analyses revealed a meaningful structure within the eight disorders identifying three groups of inter-related disorders. We detected 109 loci associated with at least two psychiatric disorders, including 23 loci with pleiotropic effects on four or more disorders and 11 loci with antagonistic effects on multiple disorders. The pleiotropic loci are located within genes that show heightened expression in the brain throughout the lifespan, beginning in the second trimester prenatally, and play prominent roles in a suite of neurodevelopmental processes. These findings have important implications for psychiatric nosology, drug development, and risk prediction.

## INTRODUCTION

Psychiatric disorders affect more than 25% of the population in any given year and are a leading cause of worldwide disability (Global Burden of Disease Injury Incidence Prevalence Collaborators, 2017; Kessler and Wang, 2008). The substantial influence of genetic variation on risk for a broad range of psychiatric disorders has been established by both twin and, more recently, large-scale genomic studies (Smoller et al., 2018). Psychiatric disorders are highly polygenic, with a large proportion of heritability contributed by common variation. Many risk loci have emerged from genome-wide association studies (GWAS) of, among others, schizophrenia (SCZ), bipolar disorder (BIP), major depression (MD), and attention-deficit/hyperactivity disorder (ADHD) from the Psychiatric Genomics Consortium (PGC) and other efforts (Sullivan et al., 2018). These studies have revealed a surprising degree of genetic overlap among psychiatric disorders (Brainstorm Consortium, 2018; Cross-Disorder Group of the Psychiatric Genomics Consoritum, 2013). Elucidating the extent and biological significance of cross-disorder genetic influences has implications for psychiatric nosology, drug development, and risk prediction. In addition, characterizing the functional genomics of cross-phenotype genetic effects may reveal fundamental properties of pleiotropic loci that differentiate them from disorder-specific loci, and help identify targets for diagnostics and therapeutics.

In 2013, analyses by the PGC’s Cross-Disorder Group identified loci with pleiotropic effects across five disorders: autism spectrum disorder (ASD), ADHD, SCZ, BIP, and MD in a sample comprising 33,332 cases and 27,888 controls (Cross-Disorder Group of the Psychiatric Genomics Consortium, 2013). In the current study, we examined pleiotropic effects in a greatly expanded dataset, encompassing 232,964 cases and 494,162 controls, that included three additional psychiatric disorders: Tourette syndrome (TS), obsessive-compulsive disorder (OCD), and anorexia nervosa (AN). We address four major questions regarding the shared genetic basis of these eight disorders: 1) Can we identify a shared etiologic structure within the broad range of these clinically distinct psychiatric disorders? 2) Can we detect additional loci associated with risk for multiple disorders (pleiotropic loci)? 3) Do some of these risk loci have opposite allelic effects across disorders? and 4) Can we identify functional features of the pleiotropic loci that could account for their broad effects on psychopathology?

## RESULTS

We analyzed genome-wide single nucleotide polymorphism (SNP) data for eight neuropsychiatric disorders using a combined sample of 232,964 cases and 494,162 controls (**Table 1; Supplementary Table 1**). The eight disorders included AN (Duncan et al., 2017), ASD (Grove et al., 2017), ADHD (Demontis et al., 2019), BIP (Stahl et al., 2018), MD (Wray et al., 2018), OCD (International Obsessive Compulsive Disorder Foundation Genetics Collaborative (IOCDF-GC) and OCD Collaborative Genetics Association Studies (OCGAS), 2018), TS (Yu et al., In press.), and SCZ (Schizophrenia Working Group of the Psychiatric Genomics, 2014). All study participants were of self-identified European ancestry, which was supported by principal component analysis of genome-wide data.

**Table 1.** Summary of eight neuropsychiatric disorder datasets

| Disorder | #Cases | #Controls | Total Samples | # of GWAS Loci | Population Prevalence (k) | Liability-based SNP heritability (SE) | References |
| --- | --- | --- | --- | --- | --- | --- | --- |
| **ADHD** | 19,099 | 34,194 | 53,293 | 9 | 0.05 | 0.222 (0.014) | Demontis et al. 2019 |
| **AN** | 3,495 | 10,983 | 14,478 | 0 | 0.01 | 0.195 (0.029) | Duncan et al. 2017 |
| **ASD** | 18,381 | 27,969 | 46,350 | 5 | 0.01 | 0.113 (0.010) | Grove et al. 2017 |
| **BIP** | 20,352 | 31,358 | 51,710 | 17 | 0.01 | 0.182 (0.011) | Stahl et al. 2018 |
| **MD** | 130,664 | 330,470 | 461,134 | 44 | 0.15 | 0.085 (0.004) | Wray et al. 2018 |
| **OCD** | 2,688 | 7,037 | 9,725 | 0 | 0.025 | 0.280 (0.041) | IOCDF-GC and OCGAS 2018 |
| **SCZ** | 33,640 | 43,456 | 77,096 | 108 | 0.01 | 0.222 (0.012) | Schizophrenia Working Group of PGC. 2014 |
| **TS** | 4,645 | 8,695 | 13,340 | 0 | 0.008 | 0.200 (0.026) | Yu et al. 2018 |
| **Total** | **232,964** | **494,162** | **727,126** |  |  |  |  |

The number of cases and controls used in the meta-analysis of the present study. The numbers may differ from those reported in the original publications because our study included only European ancestry subjects to avoid potential confounding due to ancestral heterogeneity across distinct disorder studies. SNP heritability was estimated from the GWAS summary statistics using LD score regression.

### Genetic correlations among eight neuropsychiatric disorders indicate three genetic factors

After standardized and uniform quality control, additive logistic regression analyses were performed on individual disorders (Methods). A total of 6,786,994 SNPs were common across all datasets and were retained for further study. Using the summary statistics of these SNPs, we first estimated pairwise genetic correlations among the eight disorders using linkage disequilibrium (LD) score regression analyses (Bulik-Sullivan et al., 2015a) (Methods; **Fig. 1a; Supplementary Table 2**). The results were broadly concordant with previous estimates (Brainstorm Consortium, 2018; Cross-Disorder Group of the Psychiatric Genomics Consoritum, 2013). The genetic correlation was highest between SCZ and BIP (r_g_ = 0.70 ±0.02), followed by OCD and AN (r_g_ =0.50 ±0.12). Interestingly, based on genome-wide genetic correlations, MD was closely correlated with ASD (r_g_=0.45 ±0.04) and ADHD (r_g_=0.44 ±0.03), two childhood-onset disorders. Despite variation in magnitude, significant genetic correlations were apparent for most pairs of disorders, suggesting a complex, higher-order genetic structure underlying psychopathology (**Fig. 1b**).

**Figure 1.**
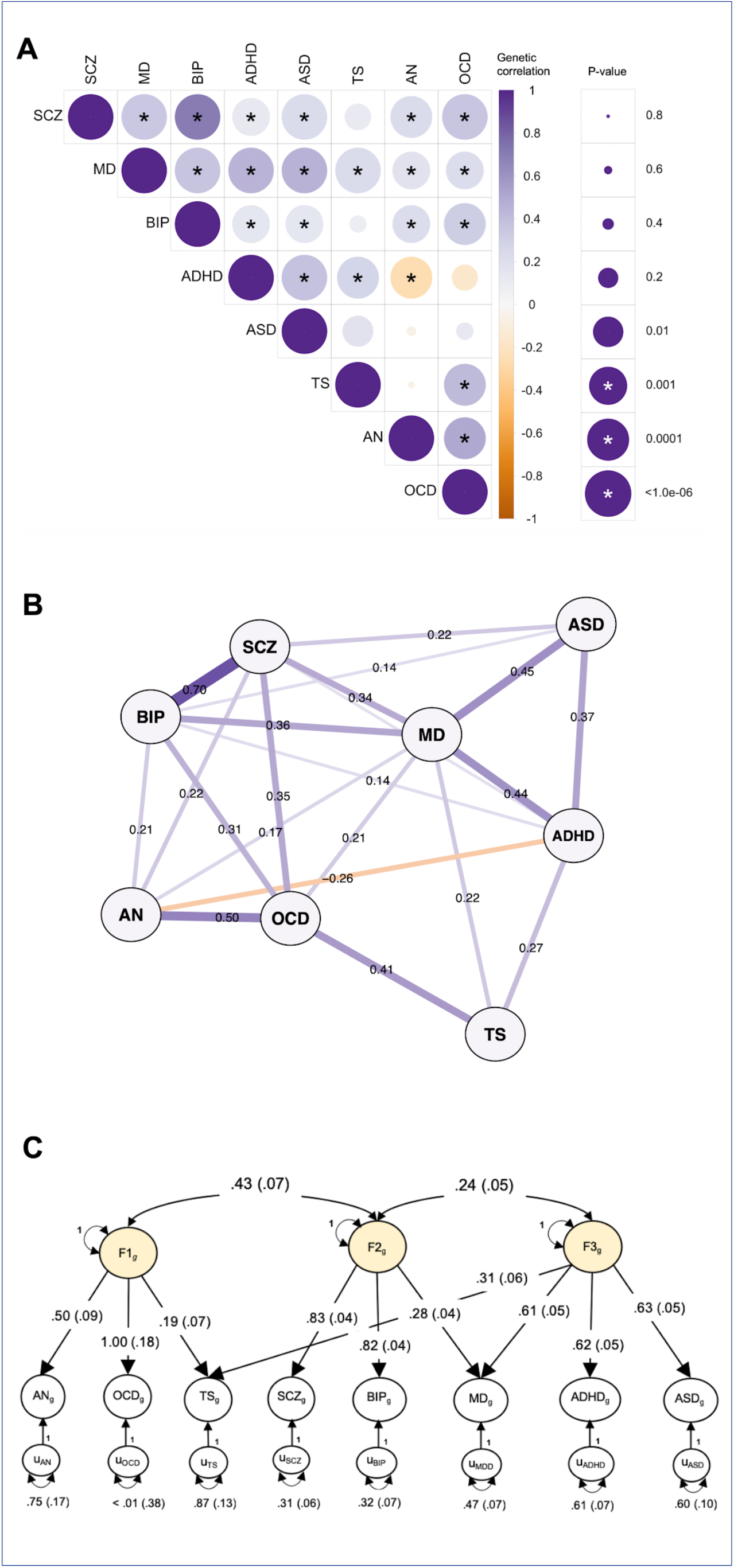
Genetic relationships between eight psychiatric disorders. A) SNP-based genetic correlations (r_g_) were estimated between eight neuropsychiatric disorders using LDSC. The size of the circles scales with the significance of the p-values. The darker the color, the larger the magnitude of r_g_. Star sign (*) indicates statistical significance after Bonferroni correction. (B) SNP-based genetic correlations between eight disorders were depicted using an in-directed graph to reveal complex genetic relationships. Only significant genetic correlations after Bonferroni correction in (A) were displayed. Each node represents a disorder, with edges indicating the strength of the pairwise correlations. The width of the edges increases, while the length decreases, with the absolute values of r_g_. (C) Based on results of an exploratory factor analysis of the genetic correlation matrix, a confirmatory factor model with three correlated genetic factors was specified using Genomic SEM and estimated with the weighted least squares algorithm. Two-headed arrows connecting the three factors to one another represent their correlations. Two-headed arrows connecting the genetic components of the individual psychiatric disorders to themselves represent residual genetic variances and correspond to the proportion of heritable variation in liability to each individual psychiatric disorder that is unexplained by the three factors. Standardized parameters are depicted with their standard errors in parentheses. Paths labeled 1 with no standard errors reported are fixed parameters, which are used for scaling.

We modeled the genome-wide joint architecture of the eight neuropsychiatric disorders using an exploratory factor analysis (EFA) (Gorsuch, 1988), followed by genomic structural equation modeling (SEM) (Grotzinger et al., 2018) (Methods). EFA identified three correlated factors, which together explained 51% of the genetic variation in the eight neuropsychiatric disorders (**Supplementary Table 3**). The first factor consisted primarily of disorders characterized by compulsive behaviors, specifically AN, OCD, and, more weakly, TS. The second factor was characterized by mood and psychotic disorders (MD, BIP, and SCZ), and the third factor by three early-onset neurodevelopmental disorders (ASD, ADHD, TS) as well as MD. Similar to our EFA results, hierarchical clustering analyses also identified three sub-groups among the eight disorders (**Supplementary Fig. 1**).

### Cross-disorder meta-analysis identifies 109 pleiotropic loci

The factor structure described above is based on average effects across the genome, but does not address more fine-grained cross-disorder effects at the level of genomic regions or individual loci. To identify genetic loci with shared risk, we performed a meta-analysis of the eight neuropsychiatric disorders using a fixed-effects-based method (Bhattacharjee et al., 2012) that accounts for the differences in sample sizes, existence of subset-specific effects, and overlapping subjects across datasets (Methods). There was no evidence of genomic inflation (λ_1000_ = 1.005; **Fig. 2a**). Using the primary fixed-effects-based meta-analysis, we identified 136 LD-independent regions with genome-wide significant association (*P*_meta_ ≤ 5×10^−8^). Due to the known extensive LD at the major histocompatibility complex (MHC) region (chromosome 6 region at 25-35 Mb), we considered the multiple signals present there as one locus. 101 of the 136 (74.3%) significantly associated regions overlapped with previously reported genome-wide significant regions from at least one individual disorder, while 35 loci (25.7%) represented novel genome-wide significant associations.

**Figure 2.**
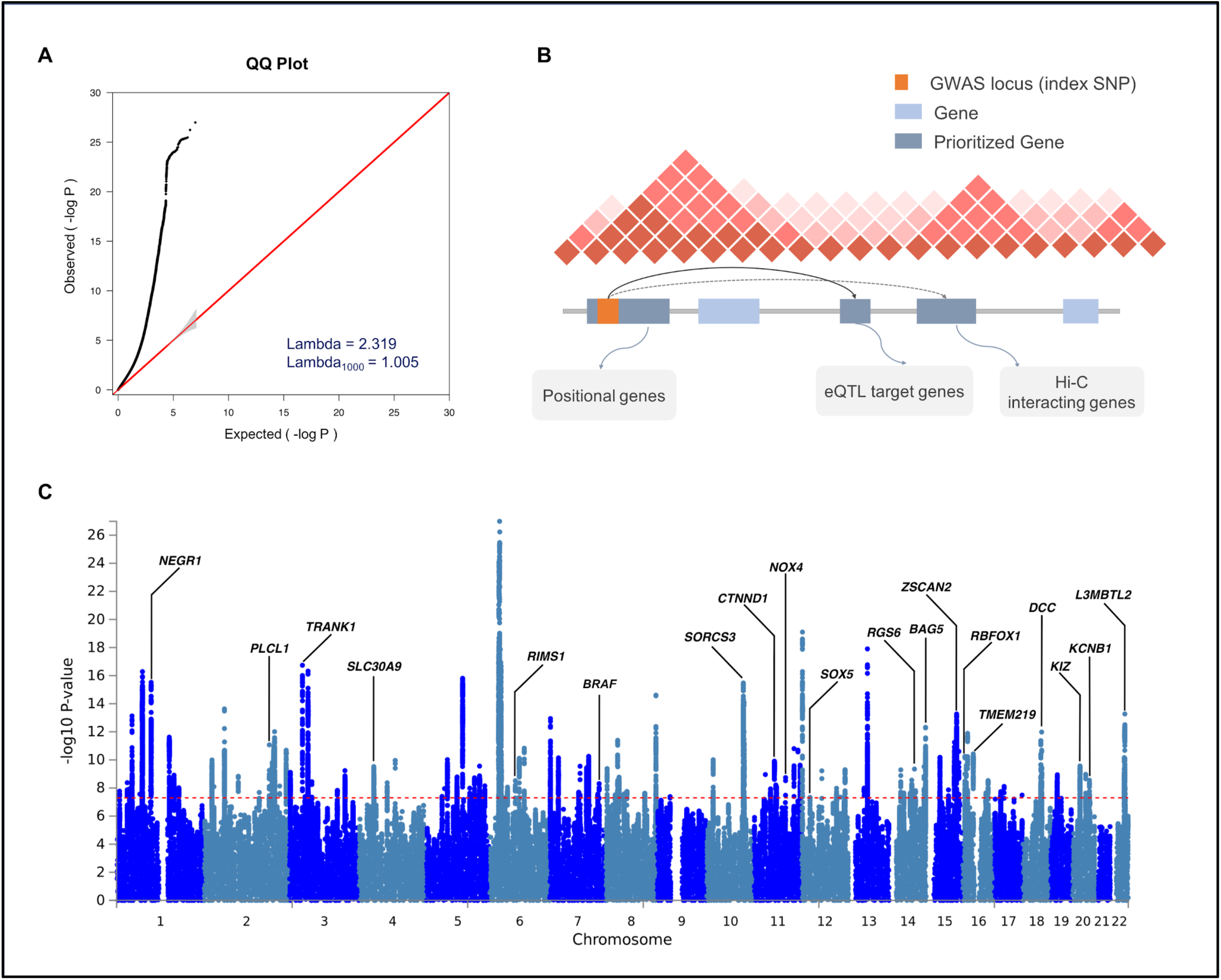
Results of cross-disorder meta-analysis and candidate gene mapping. (A) Quantile-quantile (QQ) plot displaying the observed meta-analysis statistics vs. the expected statistics under the null model of no associations in the -log_10_(p-value) scale. Although a marked departure is notable between the two statistics, the estimated lambda_1000_ and the estimated LD Score regression intercept indicate that the observed inflation is mainly due to polygenic signals rather than major confounding factors including population stratification. (B) Gene prioritization strategies for significantly associated loci. Candidate genes were mapped on each locus if the index SNP and credible SNPs reside within a protein-coding gene, are eQTL markers of the gene in the brain tissue, or interact with promoter regions of the gene based on brain Hi-C data. (C) Manhattan plot displaying the cross-disorder meta-analysis results highlighting candidate genes mapped to top pleiotropic regions.

Within these 136 loci, multi-SNP-based conditional analysis (Yang et al., 2012) identified 10 additional SNPs with independent associations, resulting in a total of 146 independent lead SNPs (**Supplementary Table 4)**. To provide a quantitative estimate of the best fit configuration of cross-disorder genotype-phenotype relationships, we estimated the posterior probability of association (referred to as the *m-value)* with each disorder using a Bayesian statistical framework (Han and Eskin, 2012) (Methods; **Supplementary Table 5**). As recommended, an *m*-value threshold of 0.9 was used to predict with high confidence that a particular SNP was associated with a given disorder. Also, *m*-values of < 0.1 were taken as strong evidence against association. Plots of the SNP *p*-value vs. *m*-value for all 146 lead SNPs are shown in **Supplementary Fig. 2**. Nearly 75% (N = 109/146) of the genome-wide significant SNPs were pleiotropic (i.e., associated with more than one disorder). As expected, configurations of disease association reflected the differences in the statistical power and genetic correlations between the samples (**Supplementary Fig. 3**). Of the 109 pleiotropic loci, 83% and 72% involved SCZ and BIP, respectively. MD, which had the largest case-control sample, was associated with 48% of the pleiotropic loci (N=52/109). Despite the relatively small sample size, ASD was implicated in 36% of the pleiotropic loci. Most of the ASD associations co-occurred with SCZ and BIP. The other disorders, ADHD, TS, OCD, and AN featured associations in 16%, 14%, 11%, and 7% of the pleiotropic loci, respectively. Of the single-disorder-specific loci, 81% and 16% were associated with SCZ and MD, respectively.

**Figure 3.**
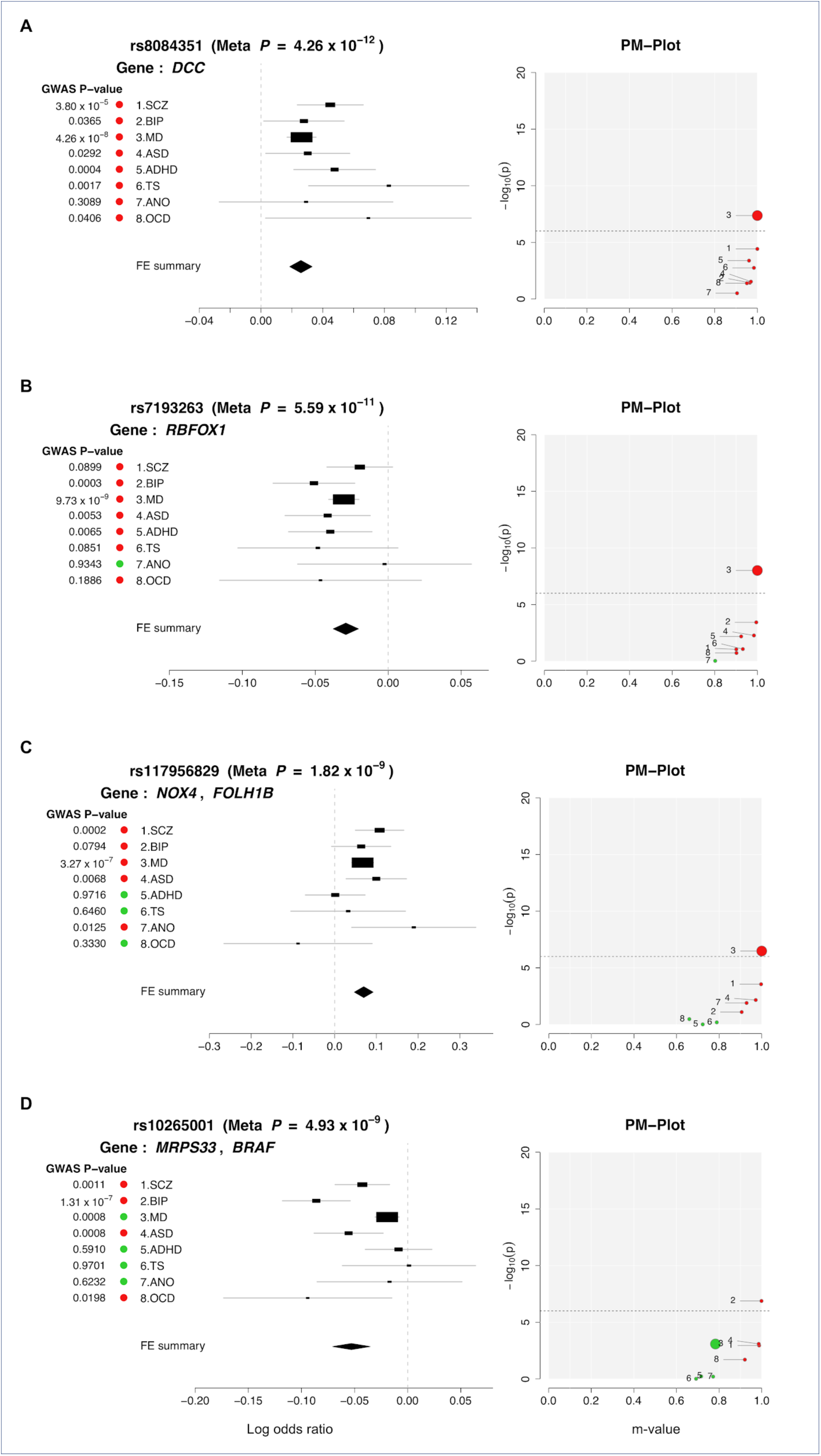
Profile of disorder associations for illustrative pleiotropic loci: (A) rs8084351 on 18q21.2; (B) rs7193263 on 16p13.3; (C) rs117956829 on 11q14.3; and (D) rs10265001 on 7q34. For each locus, disorder-specific effects of the index SNP are shown using ForestPMPlot. The first panel is the forest plot, displaying disorder-specific association p-value, log odds ratios (ORs), and standard errors of the SNP. The meta-analysis p-value and the corresponding summary statistic are displayed on the top and the bottom of the forest plot, respectively. The second panel is the PM-plot in which X-axis represents the m-value, the posterior probability that the effect eixsts in each disorder, and the Y-axis represents the disorder-specific association p-value as -log_10_(p-value). Disorders are depicted as a dot whose size represents the sample size of individual GWAS. Disorders with estimated m-values of at least 0.9 are colored in red, while those with m-values less than 0.9 are marked in green.

**Table 2** summarizes 23 pleiotropic loci associated with at least four of the disorders. Among these loci, heterogeneity of effect sizes was minimal (*p*-value of Q > 0.1). Eleven of the 23 regions map to the intron of a protein-coding gene, and seven additional lead SNPs had at least one protein-coding gene within 100 kb. We used an array of functional genomics resources, including brain eQTL and Hi-C data (Wang et al., 2018; Won et al., 2016) to prioritize potential candidate genes to the identified regions (Methods; **Fig. 2b**). The Manhattan plot in **Fig. 2c** highlights the prioritized candidate genes.

**Table 2.** Summary of 23 loci with the broadest cross-disorder association

| SNP | CHR | BP | Candidate Gene (evidence) | ADHD | ANO | ASD | BIP | MD | OCD | SCZ | TS | m |
| --- | --- | --- | --- | --- | --- | --- | --- | --- | --- | --- | --- | --- |
| rs8084351 | 18 | 50726559 | DCC(g,q) | 0.961 | 0.905 | 0.97 | 0.965 | 1 | 0.951 | 1 | 0.984 | 8 |
| rs7193263 | 16 | 6315880 | RBFOX1(g) | 0.924 | 0.802 | 0.984 | 0.995 | 1 | 0.902 | 0.901 | 0.932 | 7 |
| rs12658451 | 5 | 103904037 | - | 0.963 | 0.165 | 0.999 | 0.972 | 1 | 0.574 | 1 | 0.963 | 6 |
| rs34215985 | 4 | 42047778 | SLC30A9(g;q) | 0.908 | 0.926 | 0.992 | 0.843 | 1 | 0.88 | 0.929 | 0.913 | 6 |
| rs61867293 | 10 | 106563924 | SORCS3(g) | 0.987 | 0.954 | 0.992 | 0.985 | 1 | 0.854 | 1 | 0.886 | 6 |
| rs9360557 | 6 | 73132745 | KCNQ5(ha;hf)<br>KCNQ5-IT1(hf) | 0.905 | 0.938 | 0.976 | 0.984 | 0.993 | 0.897 | 1 | 0.892 | 6 |
| rs10149470 | 14 | 104017953 | - | 0.844 | 0.833 | 0.998 | 0.979 | 1 | 0.868 | 0.997 | 0.97 | 5 |
| rs11570190 | 11 | 57560452 | CTNND1(g)<br>OR5AK2(q) | 0.927 | 0.79 | 0.97 | 0.58 | 1 | 0.916 | 1 | 0.832 | 5 |
| rs117956829 | 11 | 89339666 | TRIM64B(ha)<br>GRM5(hf)<br>TRIM77(hf)<br>TYR(hf) | 0.723 | 0.929 | 0.972 | 0.906 | 1 | 0.66 | 0.997 | 0.789 | 5 |
| rs1484144 | 4 | 80217597 | - | 0.97 | 0.884 | 0.973 | 0.98 | 1 | 0.84 | 0.998 | 0.85 | 5 |
| rs6969410 | 7 | 110069015 | - | 0.836 | 0.827 | 0.987 | 0.93 | 0.999 | 0.917 | 1 | 0.729 | 5 |
| rs7531118 | 1 | 72837239 | - | 0.74 | 0.949 | 0.963 | 0.785 | 1 | 0.858 | 0.973 | 0.921 | 5 |
| rs9787523 | 10 | 106460460 | SORCS3(g) | 0.944 | 0.855 | 0.972 | 0.877 | 1 | 0.853 | 0.999 | 0.963 | 5 |
| rs10265001 | 7 | 140665521 | - | 0.716 | 0.772 | 0.986 | 0.999 | 0.783 | 0.921 | 0.988 | 0.692 | 4 |
| rs11688767 | 2 | 57988194 | BCL11A(h) | 0.845 | 0.899 | 0.929 | 0.983 | 1 | 0.849 | 1 | 0.698 | 4 |
| rs12129573 | 1 | 73768366 | - | 0.929 | 0.835 | 0.894 | 0.948 | 1 | 0.85 | 1 | 0.539 | 4 |
| rs1518367 | 2 | 198807015 | PLCL1(g)<br>SF3B1(q) | 0.897 | 0.783 | 0.913 | 0.991 | 1 | 0.674 | 1 | 0.865 | 4 |
| rs2332700 | 14 | 72417326 | RGS6(g) | 0.755 | 0.884 | 0.951 | 0.948 | 0.999 | 0.885 | 1 | 0.817 | 4 |
| rs5758265 | 22 | 41617897 | L3MBTL2(g)<br>CHADL(g) | 0.735 | 0.885 | 0.89 | 0.885 | 1 | 0.913 | 1 | 0.978 | 4 |
| rs6125656 | 20 | 48090779 | KCNB1(g) | 0.768 | 0.885 | 0.986 | 0.995 | 0.985 | 0.731 | 0.999 | 0.707 | 4 |
| rs7405404 | 16 | 13749859 | - | 0.763 | 0.765 | 0.99 | 0.939 | 1 | 0.726 | 1 | 0.562 | 4 |
| rs78337797 | 12 | 23987925 | SOX5(g) | 0.849 | 0.797 | 0.97 | 0.954 | 1 | 0.831 | 0.996 | 0.885 | 4 |
| rs79879286 | 7 | 24826589 | - | 0.865 | 0.854 | 0.966 | 0.999 | 1 | 0.734 | 0.999 | 0.798 | 4 |

SNP ID, location, prioritized candidate gene, disorder-specific m-values for 23 most pleiotropic loci.
The number of disorders with high confidence association (m-values $\geq 0.9$ ) is shown in the last column. Evidence for candidate gene mapping include: g (genic SNP); q (brain cis-eQTLs); h (hi-C interacting gene based on FUMA); hf (hi-C-based interaction between associated SNP and target gene in the fetal brain from Won et al. 2016); ha (hi-C-based interaction in the adult brain from Wang et al. 2018); and tss (transcription start sites).

Of the 109 risk loci with shared effects, the 18q21.2 region surrounding SNP rs8084351 at the netrin 1 receptor gene *DCC* featured the most pleiotropic association (*P*_meta_ =4.26 x 10^−12^; **Fig. 3a**). This region showed association with all eight psychiatric disorders, and has been previously associated with both MD and neuroticism (Turley et al., 2018; Wray et al., 2018). The product of *DCC* plays a key role in guiding axonal growth during neurodevelopment and serves as a master regulator of midline crossing and white matter projections (Bendriem and Ross, 2017). Gene expression data indicate that *DCC* expression peaks during early prenatal development (**Supplementary Fig. 4a**).

The second most pleiotropic locus in our analysis was identified in an intron of *RBFOX1* (RNA Binding Fox-1 Homolog 1) on 16p13.3 (lead SNP rs7193263; *P*_meta_ = 5.59 x 10^−11^). The lead SNP showed association with all of the disorders except AN (**Fig. 3b**). *RBFOX1* (also called *A2BP1*) encodes a splicing regulator mainly expressed in neurons and known to target several genes important to neuronal development, including *NMDA* receptor 1 and voltage-gated calcium channels (Hamada et al., 2015). Knock-down and silencing of *RBFOX1* during mouse corticogenesis impairs neuronal migration and synapse formation (Hamada et al., 2015; Hamada et al., 2016), implying its pivotal role in early cortical maturation. In contrast to *DCC*, however, developmental gene-expression of *RBFOX1* showed gradually increasing gene expression throughout the prenatal period (**Supplementary Fig. 4b**). Animal models and association studies have implicated *RBFOX1* in aggressive behaviors, a trait observed in several of the disorders in our analysis (Fernandez-Castillo et al., 2017).

Of the 109 pleiotropic loci, 76 were identified in the GWAS of individual disorders, while the remaining 33 are novel. The most pleiotropic among these novel loci was a region downstream of *NOX4* (NADPH Oxidase 4) that was associated with SCZ, BIP, MD, ASD, and AN (rs117956829; *P*_meta_ = 1.82 x 10^−9^; **Fig. 3c**). Brain Hi-C data (Wang et al., 2018; Won et al., 2016) detected a direct interaction of the cross-disorder association region with *NOX4* in both adult and fetal brain (interaction *p*=3.2×10^−16^ and 9.324×10^−6^, respectively). As a member of the family of *NOX* genes that encode subunits of *NADPH* oxidase, *NOX4* is a major source of superoxide production in human brain and a promoter of neural stem cell growth (Kuroda et al., 2014; Topchiy et al., 2013).

**Figure 3d** illustrates another novel psychiatric risk locus associated with SCZ, BIP, ASD, and OCD (*P*_meta_ = 3.58 x 10^−8^). The lead SNP rs10265001 resides between *MRPS33* (Mitochondrial Ribosomal Protein S33) and *BRAF* (B-Raf Proto-Oncogene, Serine/Threonine Kinase) on 7q34. The brain Hi-C data indicated interaction of the associated region with the promoters of two nearby genes: *BRAF*, which contributes to the MAP kinase signal transduction pathway and plays a role in postsynaptic responses of hippocampal neurons (Grantyn and Grantyn, 1973), and *KDM7A* (encoding Lysine Demethylase 7A), which plays a central role in the nervous system and midbrain development (Horton et al., 2010; Qi et al., 2010; Tsukada et al., 2010).

Our prior cross-disorder meta-analysis of five psychiatric disorders(Cross-Disorder Group of the Psychiatric Genomics Consortium, 2013) found no evidence of SNPs with antagonistic effects on two or more disorders. Here, we examined whether any variants with meta-analysis *p* ≤ 1×10^−6^ had opposite directional effects between disorders (Methods). After adjusting for having examined 206 loci across eight disorders (*q* < 0.001), we identified 11 loci with evidence of opposite directional effects on two or more disorders (**Fig. 4; Supplementary Table. 6**). The disorder configuration of opposite directional effects varied for the 11 loci, including three loci with opposite directional effects on SCZ and MD (rs301805, rs1933802, rs3806843), two loci between SCZ and ASD (rs9329221, rs2921036), and one locus (rs75595651) with opposite directional effects on the two mood disorders, BIP and MD. Notably, all of the six loci involving SCZ and BIP exhibited the same directional effect on the two disorders (*P*_*binom*_ < 0.05), in line with their strong genome-wide genetic correlation.

**Figure 4.**
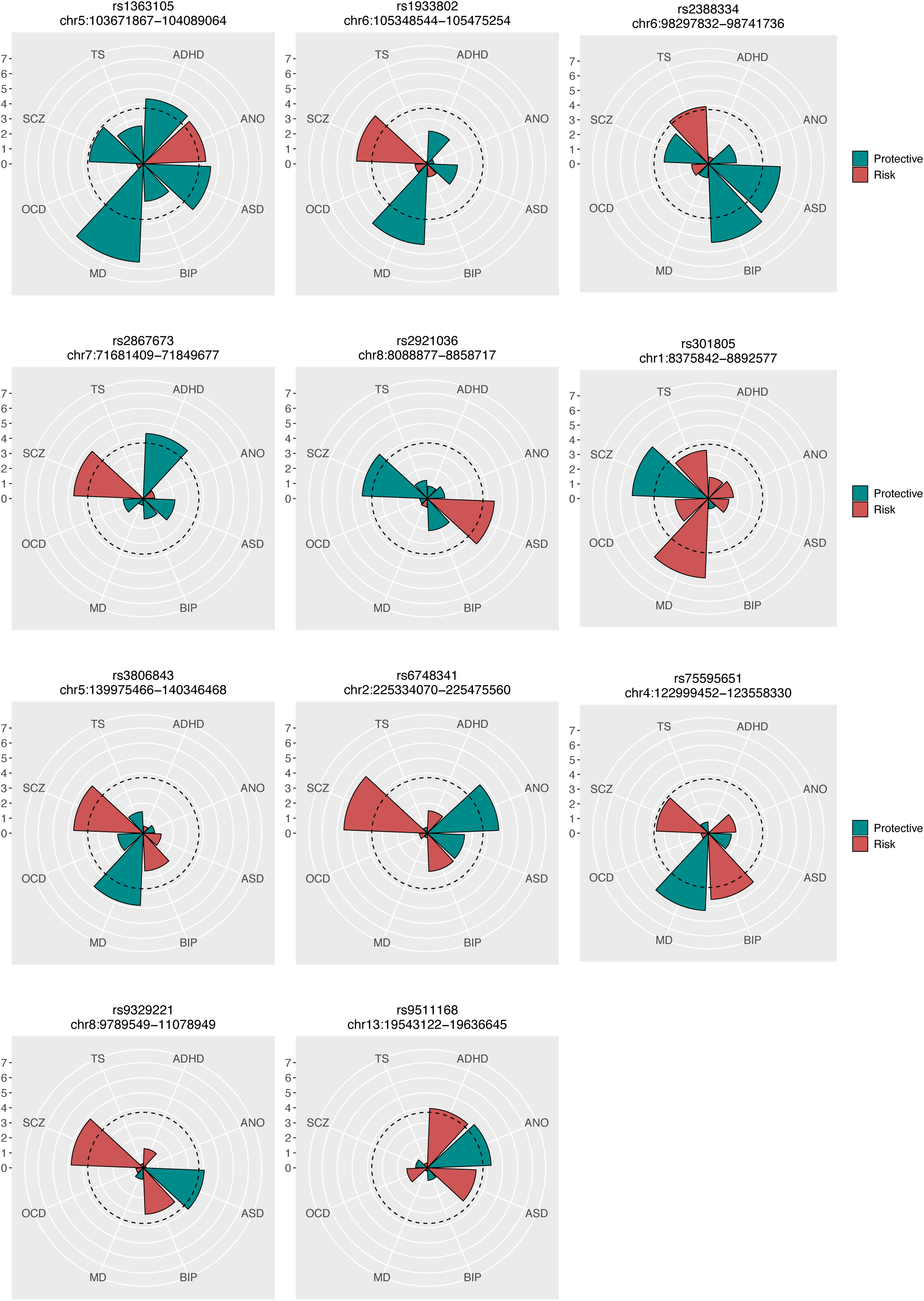
Eleven loci with opposite directional effects. The radius of each wedge corresponds to the absolute values of the Z-scores (log(Odds ratios)/S.E) obtained from association tests of the SNP for eight disorders. The color indicates whether the examined SNP carries risk (red) or protective effects (green) for each disorder. The dotted line around the center indicates statistically significant SNP effects that account for multiple testing of 206 SNPs with the q-value of 0.001.

### Functional characterization of pleiotropic risk loci

We conducted a series of bioinformatic analyses that examined whether loci with shared risk effects on multiple neuropsychiatric disorders had characteristic features that distinguished them from non-pleiotropic risk loci. First, we annotated the functional characteristics of 146 lead SNPs using various public data sources (Methods; **Supplementary Table 7-9**). Overall, they showed significant enrichment of genes expressed in the brain (*beta*=0.123, SE=0.0109, enrichment *p* = 1.22×10^−29^) and pituitary (*beta*=0.0916, SE=0.0136, *p* = 8.74 x 10^−12^), but not in the other Genotype-Tissue Expression (GTEx) tissues. (**Supplementary Table 10; Fig. 5a)**. A separate analysis of 109 pleiotropic risk loci also showed specific enrichment of genes expressed in multiple brain tissues (*p* = 1.55 x 10^−5^; **Supplementary Table 11**), while disorder-specific loci showed nominally enriched brain gene expression in the cortex (*p* =2.14 x 10^−2^; **Supplementary Table 12**).

**Figure 5.**
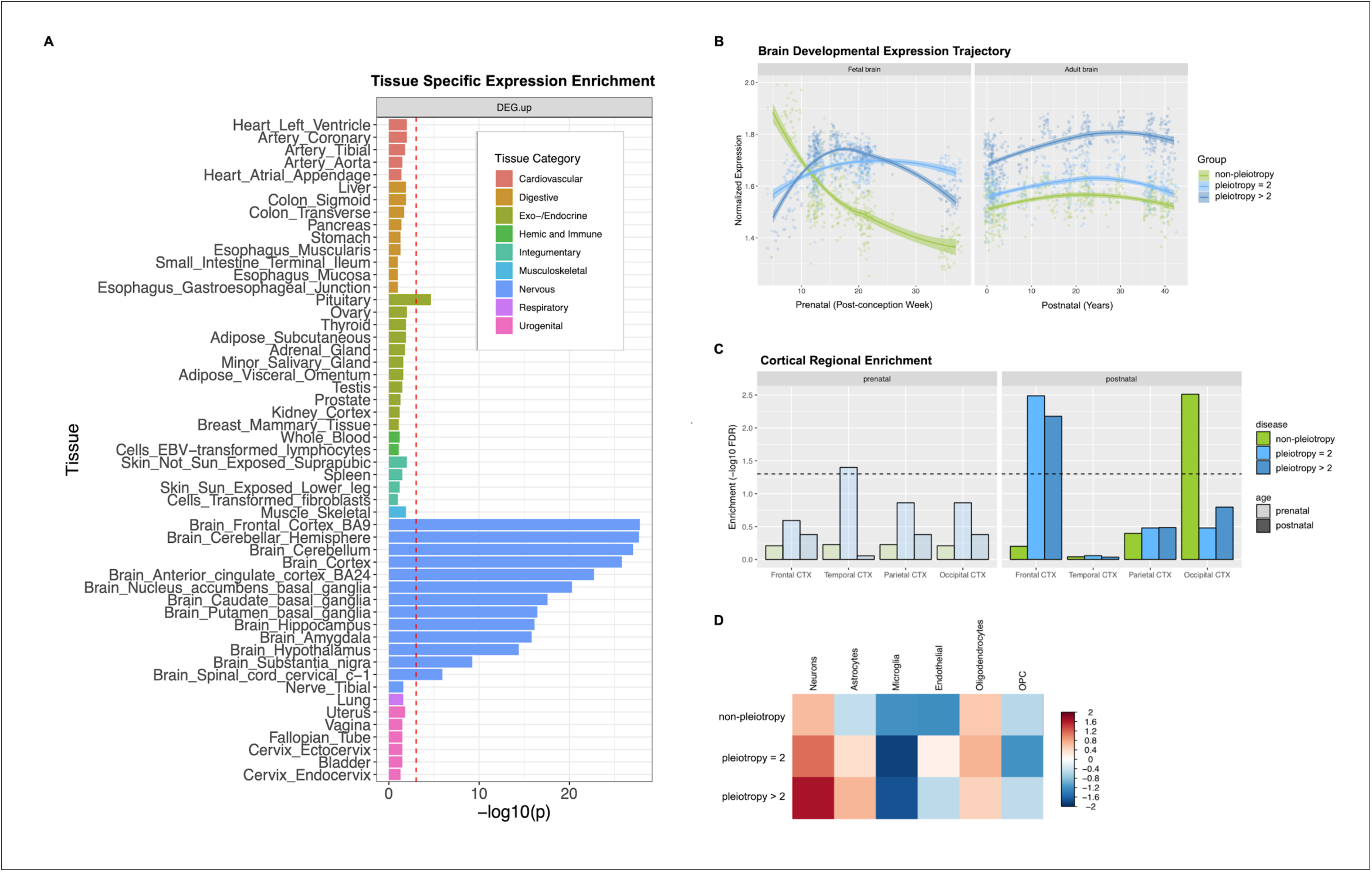
Results of functional genomics data analysis for pleiotropic vs. disorder-specific loci. (A) GTEX tissue-specific enrichment results for 146 risk loci associated with at least one of eight neuropsychiatric disorders. GTEX tissues were classified as 9 distinct categories, of which the brain tissues were colored in blue. The dotted red line indicates a statistically significant p-value after conducting Bonferroni correction for multiple testing. Psychiatric disorder-associated loci show significant enrichment in genes expressed in pituitary and all brain tissues except nerve_tibal. (B) Brain developmental expression trajectory displayed for the three groups of genes based on Kang et al. The 146 genome-wide significant loci from the cross-disorder meta analysis were clustered into three groups based on predicted disorder-specific associations: (1) no-pleiotopy; (2) pleiotropy=2; and (3) pleiotropy>2. The “no-pleiotropy” group included 37 loci that showed a single-disorder-specific association, while the “pleiotropy=2” and “pleiotropy>2” groups included 60 and 49 loci that were associated with two and more than two disorders, respectively. (C) In the adult cortex, genes mapped to pleiotropic loci were enriched for frontal cortex specific genes, while genes mapped to non-pleiotropic loci are enriched for occipical cortex specific genes. (D) Genes mapped to 146 risk loci show higher expression values in neurons and oligodendrocytes, with much higher neuronal specificity for pleiotropic loci.

Gene-set enrichment analyses using Gene Ontology data suggested involvement of pleiotropic risk loci in neurodevelopmental processes (**Supplementary Table 13,14**). The 109 pleiotropic risk loci were enriched for genes involved in neurogenesis (gene-set enrichment *p* = 9.67 x 10^−6^), regulation of nervous system development (*p* = 3.41 x 10^−5^), and neuron differentiation (*p* = 3.30 x 10^−5^), while enrichment of these gene-sets was not seen for disorder-specific risk loci (adjusted enrichment *p* > 0.05). Pleiotropic risk loci also showed enrichment of genes involved in specific neurotransmitter-related pathways – glutamate receptor signaling (*p* = 2.45 x 10^−6^) and voltage-gated calcium channel complex (*p* = 5.72 x 10^−4^) – while non-pleiotropic risk loci, which were predominantly SCZ-associated, were over-represented among acetylcholine receptor genes (*p* = 7.25 x 10^−8^). Analysis of cortical gene expression data also suggested enrichment of pleiotropic risk genes in cortical glutamatergic neurons through layers 2-6 (**Supplementary Table 15**), further supporting the shared role of glutamate receptor signaling in the pathogenesis of diverse neuropsychiatric disorders.

In contrast to the differences in neuronal development and neuronal signaling pathways, pleiotropic and non-pleiotropic risk loci shared several characteristics related to genomic function. For instance, gene-set enrichment analyses indicated that both pleiotropic and non-pleiotropic risk loci were enriched for genes involved in the regulation of synaptic plasticity, neurotransmission, and synaptic cellular components. More than 41% of the genes associated with our genome-wide significant loci, both pleiotropic and non-pleiotropic, were intolerant of loss of function mutations (pLI score ≥ 0.9); this is highly unlikely to occur by chance (Fisher’s exact *p*=4.90×10^−8^). This finding was consistent when examining pleiotropic (*p*=2.85×10^−11^) and non-pleiotropic risk loci (*p*=1.56×10^−3^) separately.

Next, we compared spatio-temporal gene-expression patterns for the 109 pleiotropic risk loci and the 37 disorder-specific loci using post-mortem brain data. On average, disorder-specific and pleiotropic risk loci showed a similar level of gene expression in both prenatal and postnatal development after multiple testing correction (t-test *p* > 0.025 x10^−2^; **Supplementary Fig. 5**). During prenatal development, non-pleiotropic loci (mainly SCZ-associated) showed peak expression in the first trimester, after which expression rapidly decreased, while pleiotropic genes associated with only 2 disorders (“*pleiotropy=2*”; 60 loci) and those associated with more than 2 (“*pleiotropy>2*”, 49 loci) showed peak expression around the second trimester (**Fig. 5b**). After birth, all three groups showed gradually increasing gene expression until adulthood. Expression levels were associated with the degree of pleiotropy, with the *pleiotropy>2* group showing higher gene expression than either the *pleiotropy=2* group (t-test p < 2.10×10^−4^) or non-pleiotropic risk loci (t-test p < 2.2×10^−16^).

Enrichment analyses using the genes preferentially expressed in specific cortical regions suggested that pleiotropic loci were over-represented among genes expressed in the frontal cortex, while non-pleiotropic loci were enriched in the occipital cortex (FDR *q*<0.05; **Fig. 5c**). Cell-type-specific analysis indicated that genes implicated in pleiotropic loci were mainly expressed in neurons (FDR *q*<0.05) but not in glial cell types. Further, enrichment of pleiotropic loci in neuronal cells was also associated with the degree of pleiotropy, as highlighted in **Fig. 5d**.

Previous studies of model organisms using gene knock-out experiments suggested that pleiotropic risk loci may undergo stronger selection than non-pleiotropic loci (Hill and Zhang, 2012). However, we found no evidence that pleiotropic risk variants are under stronger evolutionary constraints (**Supplementary Table 16**). Various comparative genomics resources, including PhyloP (Pollard et al., 2010), PhastCons (Siepel et al., 2005), and GERP++ (Davydov et al., 2010), showed our top loci to have similar properties regardless of the extent of pleiotropy. Neither did we find differences between disorder-specific lead SNPs and pleiotropic SNPs with respect to their minor allele frequencies, average heterozygosity, or predicted allele ages (Kiezun et al., 2013). Pleiotropic and non-pleiotropic SNPs also did not differ in terms of the distance to nearest genes, distance to splicing sites, chromosome compositions, and predicted functional consequences of non-coding regulatory elements.

### Relationship between cross-disorder genetic risk and other brain-related traits and diseases

To explore the genetic relationship of cross-disorder genetic risk with other traits, we treated this 8-disorder GWAS meta-analysis as a single “cross-disorder phenotype.” We applied LDSC to estimate SNP heritability (*h*^*2*^_SNP_) and genetic correlations with other phenotypes, using block jackknife-based standard errors to estimate statistical significance. The estimated *h*^*2*^_SNP_ of the cross-disorder phenotype was 0.146 (SE 0.0058; observed scale). Using data for 28 brain-related traits selected from LDHub (Zheng et al., 2017), we found significant genetic correlations of the cross-disorder phenotype with seven traits (at a Bonferroni-corrected p-value threshold 0.002): never/ever smoking status, years of education, neuroticism, subjective well-being, and three sleep-related phenotypes (chronotype, insomnia, and excessive daytime sleepiness) (**Supplementary Table 17**).

GWAS catalog data for the 109 pleiotropic risk loci showed enrichment of implicated genes in a range of brain-related traits (**Supplementary Table 18**). As expected, the associated traits included previous studies of neuropsychiatric disorders including SCZ, BIP, and ASD. In addition, the pleiotropic risk loci were enriched among genes previously associated with neuroticism (corrected enrichment *p*= 5.28×10^−6^; *GRIK3, CTNND1, DRD2, RGS6, RBFOX1, ZNF804A, L3MBTL2, CHADL, RANGAP1, RSRC1,GRM3*), cognitive ability (corrected *p*= 7.15×10^−5^; *PTPRF, NEGR1, ELOVL3, SORCS3, DCC, CACNA1I),* and night sleep phenotypes *(corrected p*= 1.86×10^−2^; *PBX1, NPAS3, RGS6, GRIN2A, MYO18A, TIAF1, CNTN4, PPP2R2B, TENM2, CSMD1*). We also found significant enrichment of pleiotropic risk genes in multiple measures of body mass index (BMI), supporting previous studies suggesting a shared etiologic basis between a range of neuropsychiatric disorders and obesity (Hartwig et al., 2016; Lopresti and Drummond, 2013; Milaneschi et al., 2018)

## DISCUSSION

In the largest cross-disorder GWAS meta-analysis of neuropsychiatric disorders to date, comprising more than 725,000 cases and controls across eight disorders, we identified 146 LD-independent lead SNPs associated with at least one disorder, including 35 novel loci. Of these, 109 loci were found to affect two or more disorders, although characterization of this pleiotropy is partly dependent on per-disorder sample size. Our results provide four major insights into the shared genetic basis of psychiatric disorders.

First, modeling of genetic correlations among the eight disorders using two different methods (EFA and hierarchical clustering) identified three groups of disorders based on shared genomics: one comprising disorders characterized by compulsive behaviors (AN, OCD and TS), a second comprising mood and psychotic disorders (MD, BIP and SCZ), and a third comprising two early-onset neurodevelopmental disorders (ASD and ADHD) and one disorder each from the first two factors (TS and MD). The loading of MD on two factors may reflect biological heterogeneity within MD, consistent with recent evidence showing that early-onset depression is associated with genetic risk for ADHD and with neurodevelopmental phenotypes (Rice et al., 2018). Overall, these results indicate a substantial pairwise genetic correlation between multiple disorders along with a higher-level genetic structure that point to broader domains underlying genetic risk to psychopathology. These findings are at odds with the classical, categorical classification of mental illness.

Second, variant-level analyses support the existence of substantial pleiotropy, with nearly 75% of the 146 genome-wide significant SNPs influencing more than one of the eight examined disorders. We also identified a set of 23 loci with particularly extensive pleiotropic profiles, affecting four or more disorders. The most highly pleiotropic locus in our analyses, with evidence of association with all eight disorders, maps within *DCC,* a gene fundamental to the early development of white matter connections in the brain (Bendriem and Ross, 2017). Prior studies showed that *DCC* is a master regulator of axon guidance (through its interactions with netrin-1 and draxin (Liu et al., 2018). Loss of function mutations in *DCC* cause severe neurodevelopmental syndromes involving loss of midline commissural tracts and diffuse disorganization of white matter tracts (Bendriem and Ross, 2017; Jamuar et al., 2017; Marsh et al., 2017). A highly pleiotropic effect of variation in *DCC* on diverse psychiatric disorders with childhood and adolescent onset would be consistent with its role in both early organization of neuronal circuits and the maturation of mesolimbic dopaminergic connections to the prefrontal cortex during adolescence (Hoops and Flores, 2017; Reynolds et al., 2018; Vosberg et al., 2018).

We also identified a set of loci that have opposite effects on risk of psychiatric disorders. Notably, these included loci with opposing effects on pairs of disorders that are genetically correlated and have common clinical features. For example, a SNP within *MRSA* was associated with opposing effects on two neurodevelopmental disorders (ASD and SCZ), and a variant within *KIAA1109* had opposite directional effects on major mood disorders (BIP and MD) (**Supplementary Table 6**). These results underscore the complexity of genetic relationships among related disorders and suggest that overall genetic correlations may obscure antagonistic biological mechanisms that operate at the level of component loci and pathways as seen in immune-mediated diseases (Baurecht et al., 2015; Lettre and Rioux, 2008; Schmitt et al., 2016). This heterogeneity of effects between genetically correlated disorders is also consistent with a recent analysis that revealed loci contributing to biological differences between BIP and SCZ and found polygenic risk score associations with specific symptom dimensions (Bipolar Disorder and Schizophrenia Working Group of the Psychiatric Genomics Consortium, 2018).

Third, we found extensive evidence that neurodevelopmental effects underlie the cross-disorder genetics of mental illness. In addition to *DCC*, a link between pleiotropy and genetic effects on neurodevelopment was also seen for other top loci in our analysis, including *RBFOX1, BRAF*, and *KDM7A*, all of which have been shown in prior research to influence aspects of nervous system development. Gene enrichment analyses showed that pleiotropic loci were distinguished from disorder-specific loci by their involvement in neurodevelopmental pathways including neurogenesis, regulation of nervous system development, and neuron differentiation. These results are consistent with those of a smaller recent analysis in the population-based Danish iPSYCH cohort (comprising 46,008 cases and 19,526 controls across six neuropsychiatric disorders) (Schork et al., 2017). In that analysis, consistent with the present findings, functional genomic characterization of cross-disorder loci implicated fetal neurodevelopmental processes, with greater prenatal than postnatal expression. However, the specific loci, cell types, and pathways implicated in the iPSYCH analysis differed from those identified in our study. Of note, however, *SORCS3* emerged as a genome-wide significant cross-disorder locus in both studies.

Fourth, our analyses of spatiotemporal gene expression profiles revealed that pleiotropic loci are enriched among genes expressed in neuronal cell types, particularly in frontal or prefrontal regions. They also demonstrated a distinctive feature of genes related to pleiotropic loci: compared with disorder-specific loci, they are on average expressed at higher levels both prenatally and postnatally (Figure 4). More specifically, single-disorder (mainly SCZ) loci were related to genes that were preferentially expressed in the first fetal trimester followed by a decline over the prenatal period and then relatively stable levels postnatally. In contrast, expression of genes related to pleiotropic loci peaked in the second trimester and remained overexpressed throughout the lifespan. When dividing the pleiotropic loci into bins of those associated with two disorders (mainly SCZ and BIP) vs. three or more disorders, we observed a consistent gradient of greater expression associated with broader pleiotropy.

Taken together, our results suggest that pleiotropic loci appear to be distinguished by both their differential importance in neurodevelopmental processes and their heightened brain expression after the first trimester. Apart from this, however, pleiotropic loci were similar to non-pleiotropic loci across a range of other functional features, including intolerance to loss-of-function mutations, evidence of selection, minor allele frequencies, and genomic position relative to functional elements.

Overall, our results identify a range of pleiotropic effects among loci associated with psychiatric disorders. Consistent with prior research (Brainstorm Consortium, 2018; Cross-Disorder Group of the Psychiatric Genomics Consoritum, 2013), we found substantial pairwise genetic correlations across child-and adult-onset disorders and extended these findings by demonstrating clusters of genetically-related disorders. These results augment a substantial body of research demonstrating that genetic influences on psychopathology do not map cleanly onto the clinical nosology instantiated in the DSM or ICD (Smoller et al., 2018). Using a range of bioinformatic and functional genomic analyses, we find that loci with pleiotropic effects are distinguished by their involvement in early neurodevelopment and increased expression beginning in the second trimester of fetal development and persisting throughout adulthood. Taken together, the analyses presented here suggest that genetic influences on psychiatric disorders comprise at least two general classes of loci. The first comprises a set of genes that confer relatively broad liability to psychiatric disorders by acting on early neurodevelopment and the establishment of brain circuitry. These pleiotropic genes begin to come online by the second trimester of fetal development and exhibit differentially high expression thereafter. Such loci may underlie a latent general psychopathology factor (the “p” factor) (Caspi et al., 2014) that has been identified in developmental studies of mental disorders, comprising transdiagnostic symptom clusters (internalizing, externalizing, and psychotic) (Caspi et al., 2014). The expression and differentiation of this generalized genetic risk into discrete psychiatric syndromes (e.g., ASD, BIP, AN) may then involve direct and/or interactive effects of additional sets of loci and environmental factors, possibly mediated by epigenetic effects, that shape phenotypic expression via effects on brain structure/function and behavior. Further research will be needed to clarify the nature of such effects.

Our results should be interpreted in light of several limitations. First, while our dataset is the largest genome-wide cross-disorder analysis to date, data available for individual disorders varied substantially—from a minimum of 9,725 cases and controls for OCD to 461,134 cases and controls for MD. This imbalance of sample size may have limited our power to detect pleiotropic effects on underrepresented disorders. Second, it is possible that comorbidity among disorders contributed to apparent pleiotropy; however we found that less than 2% of cases overlapped between disorder datasets (excluding 23andMe data) and we adjusted for overlap in meta-analysis. Third, the method we applied to detect cross-phenotype association, which combines an all-subsets fixed-effects GWAS meta-analysis with a Bayesian method for evaluating the best-fit configuration of genotype-phenotype associations, is one of several approaches (Solovieff et al., 2013). However, we have previously shown that this method outperforms a range of alternatives for detecting pleiotropy under various settings (Zhu et al., 2018). Fourth, our designation of loci as pleiotropic vs. non-pleiotropic loci refers only to their observed effects on the eight target brain disorders. Thus, some of the “non-pleiotropic” loci may have additional effects on psychiatric phenotypes that were not included in our meta-analysis and/or on non-psychiatric phenotypes. Fifth, our functional genomic analyses were constrained by the limitations of existing resources (e.g. spatiotemporal gene expression data resources). Our work underscores the need for more comprehensive functional data including single cell transcriptomic and epigenomic profiles across development and brain tissues. Lastly, we included only individuals of European ancestry to avoid potential confounding due to ancestral heterogeneity across distinct disorder studies. Similar efforts are needed to examine these questions in other populations.

In sum, in a large-scale cross-disorder genome-wide meta-analysis, we identified three genetic factors underlying the genetic basis of eight psychiatric disorders. We also identified 109 genomic loci with pleiotropic effects, of which 33 have not previously been associated with any of the individual disorders. In addition, we identified 11 loci with opposing directional effects on two or more psychiatric disorders. These results highlight disparities between our clinically-defined classification of psychiatric disorders and underlying biology. Future research is warranted to determine whether more genetically-defined influences on cross-diagnostic traits or subtypes of dissect may inform a biologically-informed reconceptualization of psychiatric nosology. Finally, we found that genes associated with multiple psychiatric disorders are disproportionately associated with biological pathways related to neurodevelopment and exhibit distinctive gene expression patterns, with enhanced expression beginning in the second prenatal trimester and persistently elevated expression relative to less pleiotropic genes. Therapeutic modulation of pleiotropic gene products could have broad-spectrum effects on psychopathology.

## AUTHOR CONTRIBUTIONS

*Writing Group*: P.H.L, Y.A.F., V.A., S.V.F., B.M.N., K.S.M., L.T., S.R., N.W., M.N., G.B., J.W., J.S., H.E., J.G., O.A., M.B.S., C.A.M., A.F., S.S., M.M., A.Z., A.B., K.S.K., J.W.S. (Chair). *Analysis Group*: P.H.L. (Lead), V.A., H.W., Y.A.F, J.R., Z.Z., E.T-D., M.G.N., A.D.G, D.P., M, M-J, W. Disorder-specific data collection, analysis, and identification of duplicate subjects were conducted by D.Y., S.R., E.S., R.A., R.W., D.M., M.M., A.B., 23andme, and L.E.D. *Editorial Revisions Group*: S.B., J.L., H.K., A.K., E.H.C., G.K., G.C., J.K., C.C.Z., P.J.H., T.B., L.A.R., B.F., J.I.N. The remaining authors contributed to the recruitment, genotyping, or data processing for the contributing components of the study. All other authors saw, had the opportunity to comment on, and approved the final draft.

## ACKNOWLEDGMENTS

Full acknowledgements and a list of funding are in the Supplementary Note. The work of the contributing groups was supported by numerous grants from governmental and charitable bodies as well as philanthropic donation. Specifically, P.H.L. (R00MH101367) and J.W.S. (R01MH106547; R01MH117599; U01HG008685). We thank the research participants and employees of 23andMe, Inc. for their contribution to this study.

## Supplementary Figure Titles and Legends

**Figure S1.**
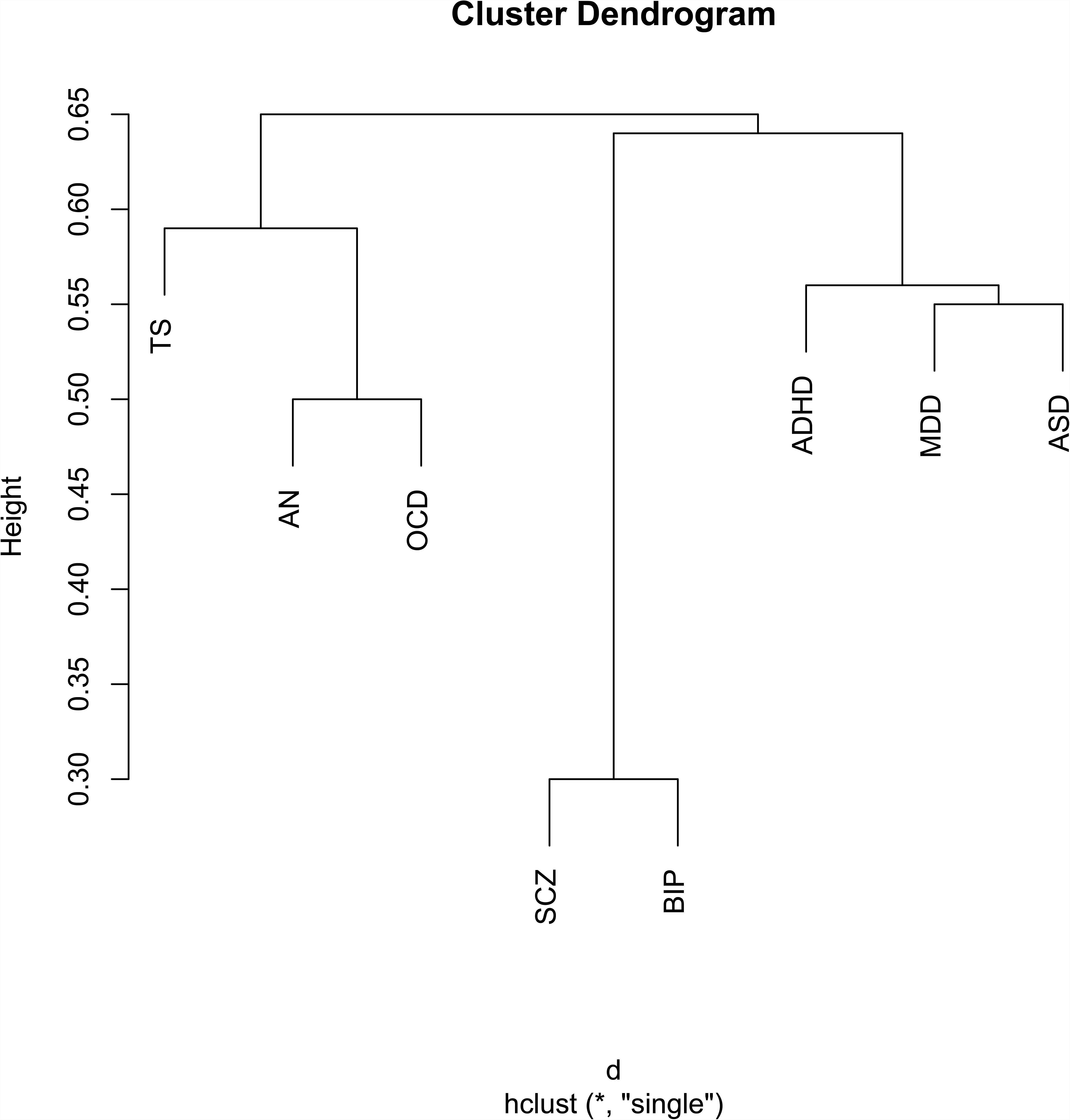
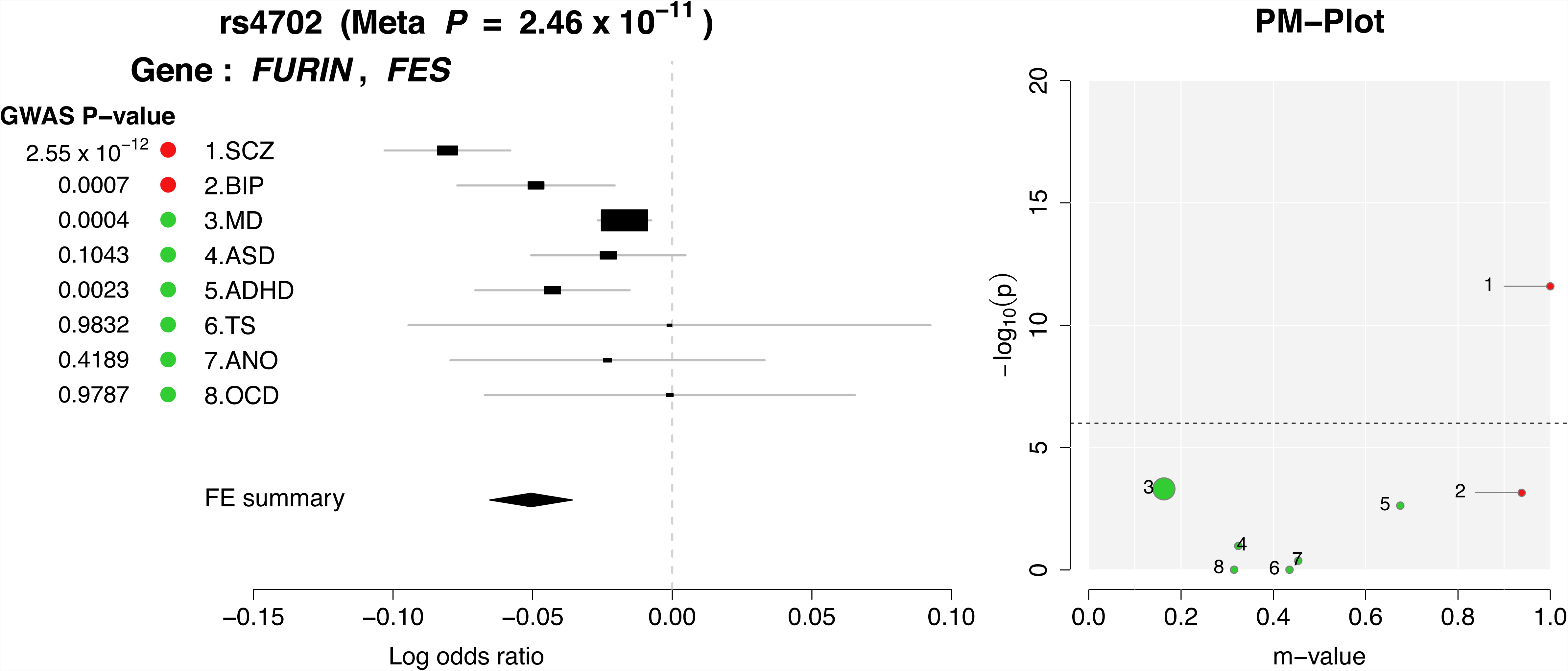
Hierarchical clustering of genetic architecture across eight neuropsychiatric disorders. (Related to Table S3) Hierarchical clustering revealed three sub-groups within the eight disorders. Results were very similar to those obtained using exploratory factor analysis and genomic structural equation modeling.

**Figure S2.**
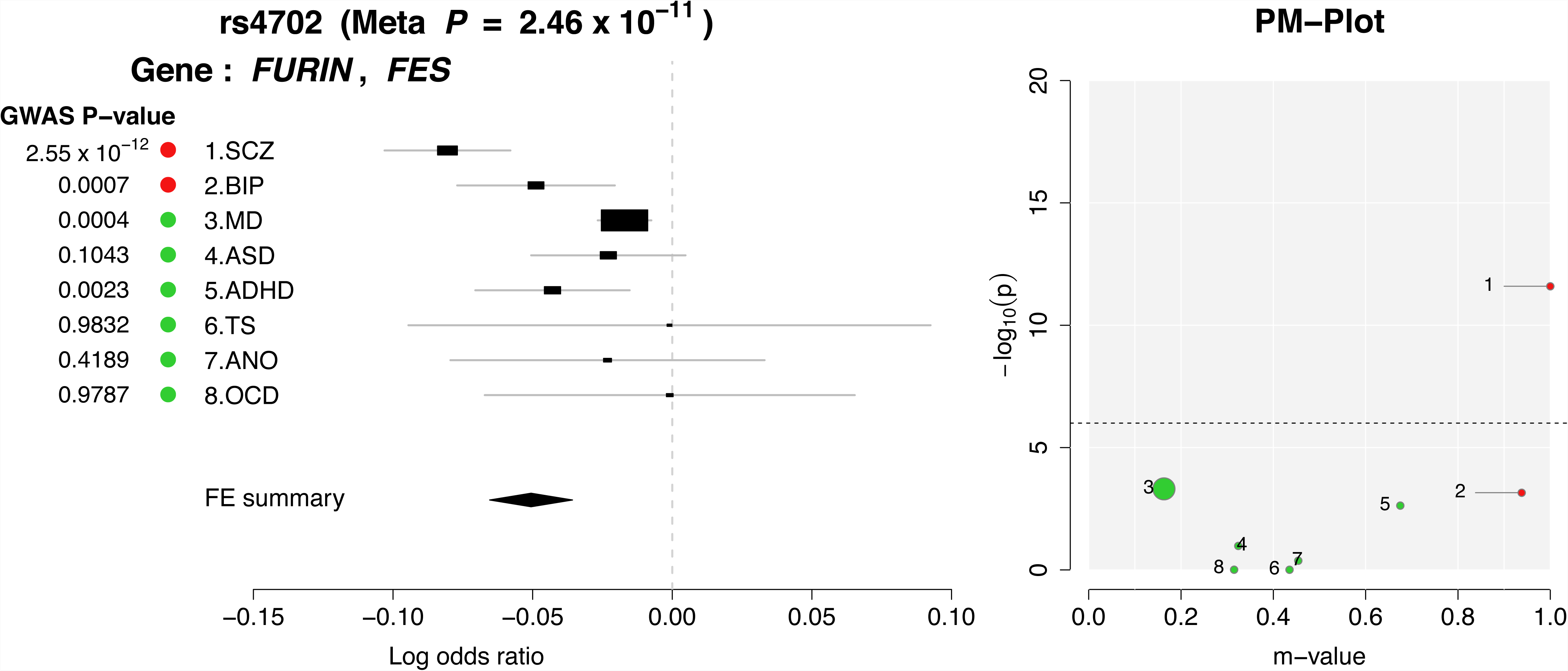
P-M plots for top 146 loci. (Related to Figure 3 and Table S5) M-values were generated for each locus for each disorder and indicate the presence or absence of a disorder-specific effect. M-values were plotted with the negative log odds of the corresponding p-values for each disorder for a given SNP; P-M plots were generated for each of the top 146 loci. Log odds ratios for the effect of the SNP on each disorder and a summary across disorders was also plotted. As expected, most of the top SNPs (109/146) were pleiotropic.

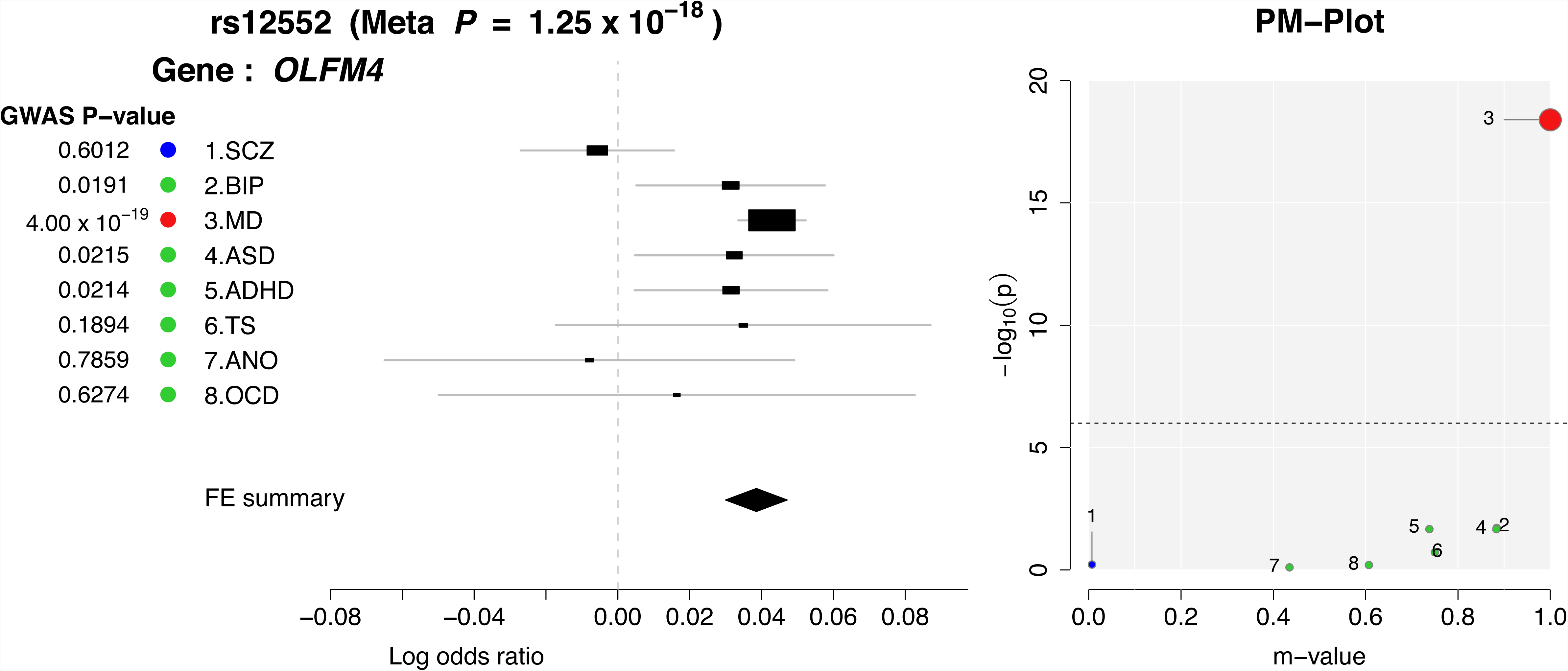

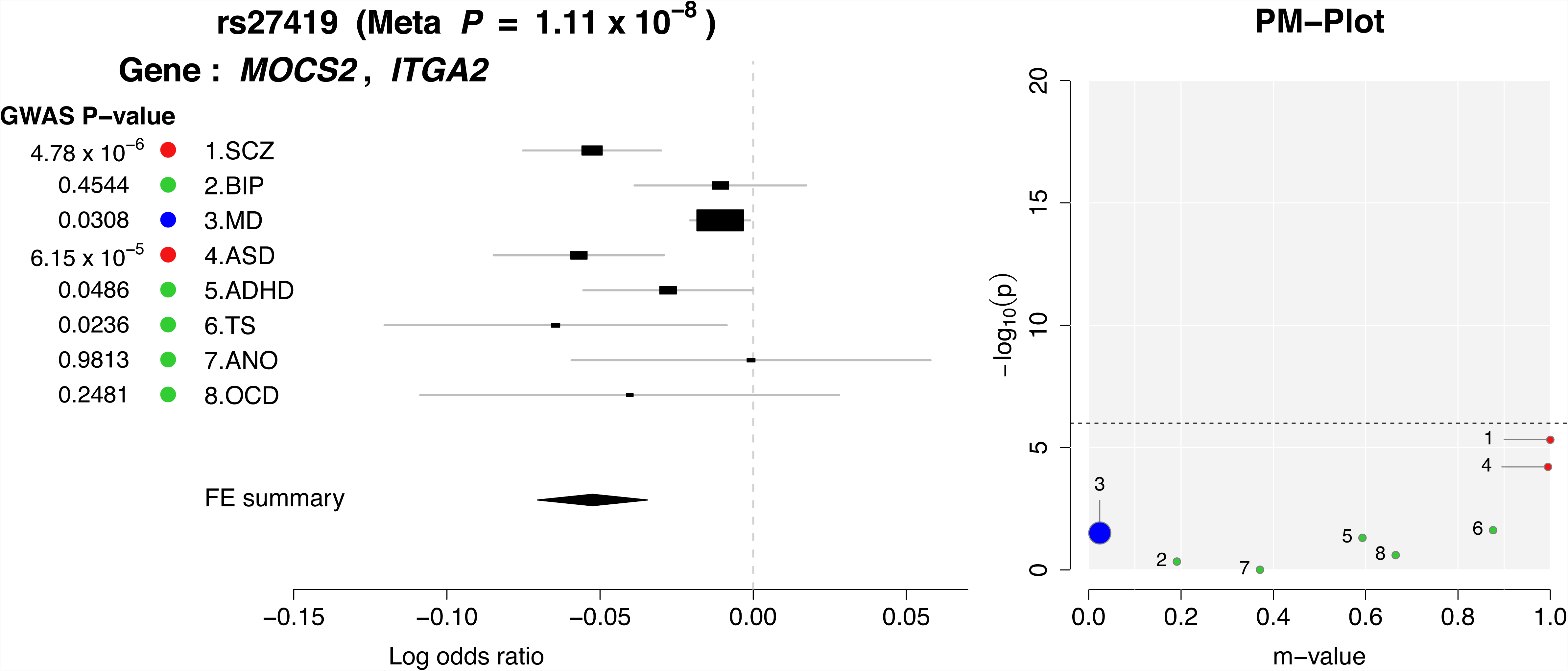

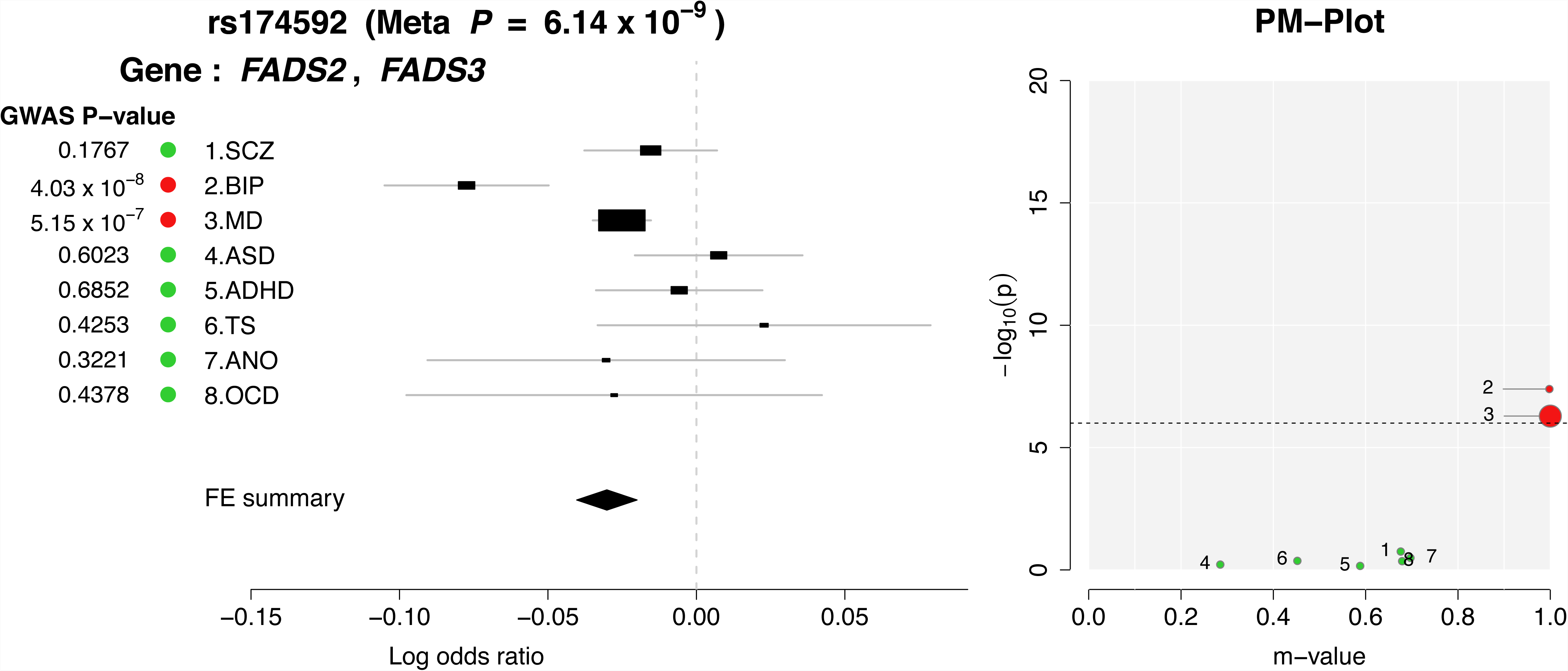

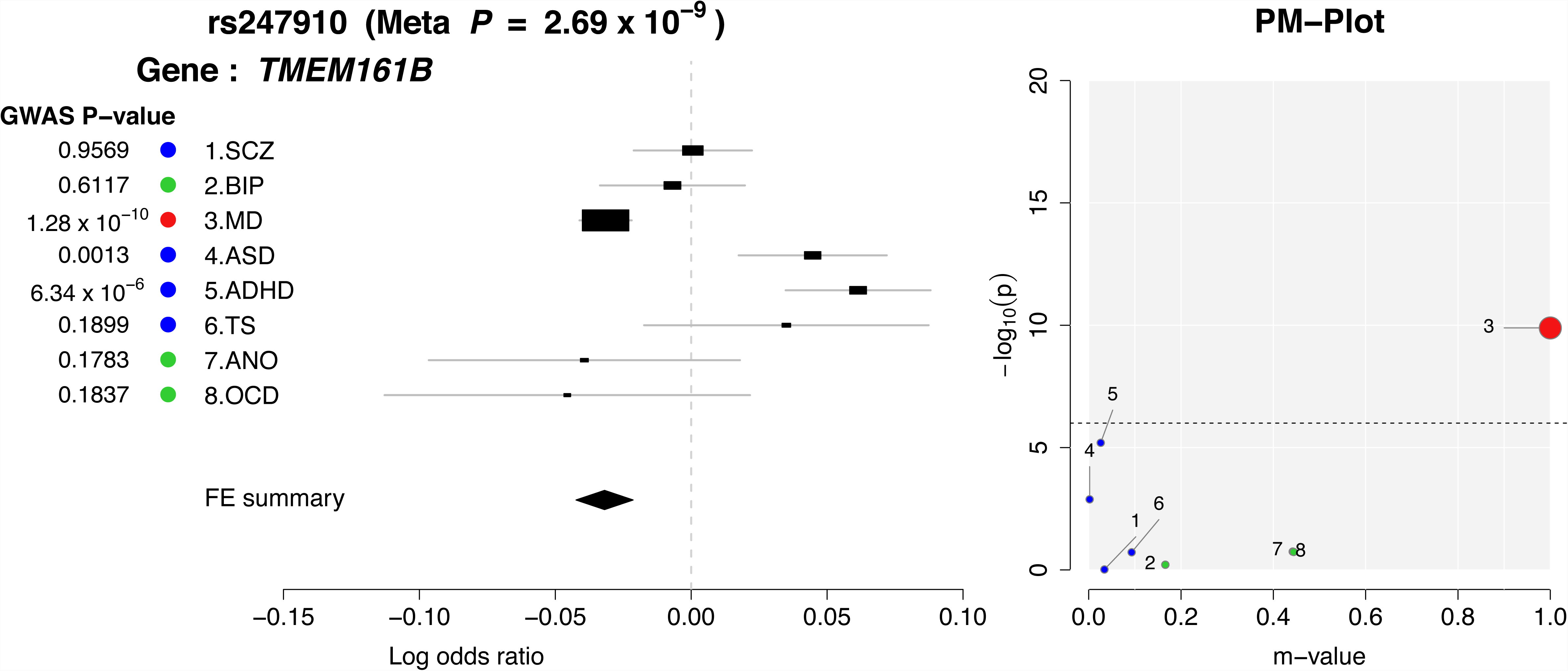

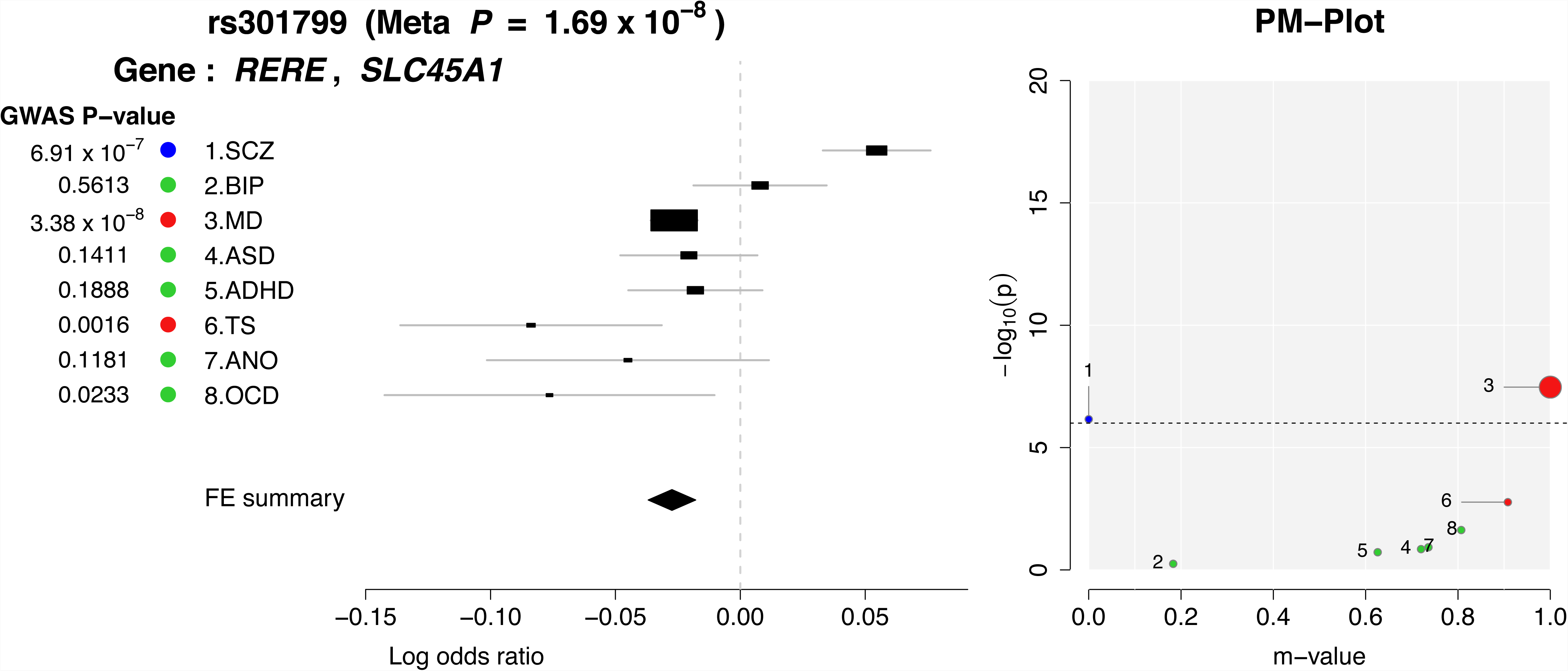

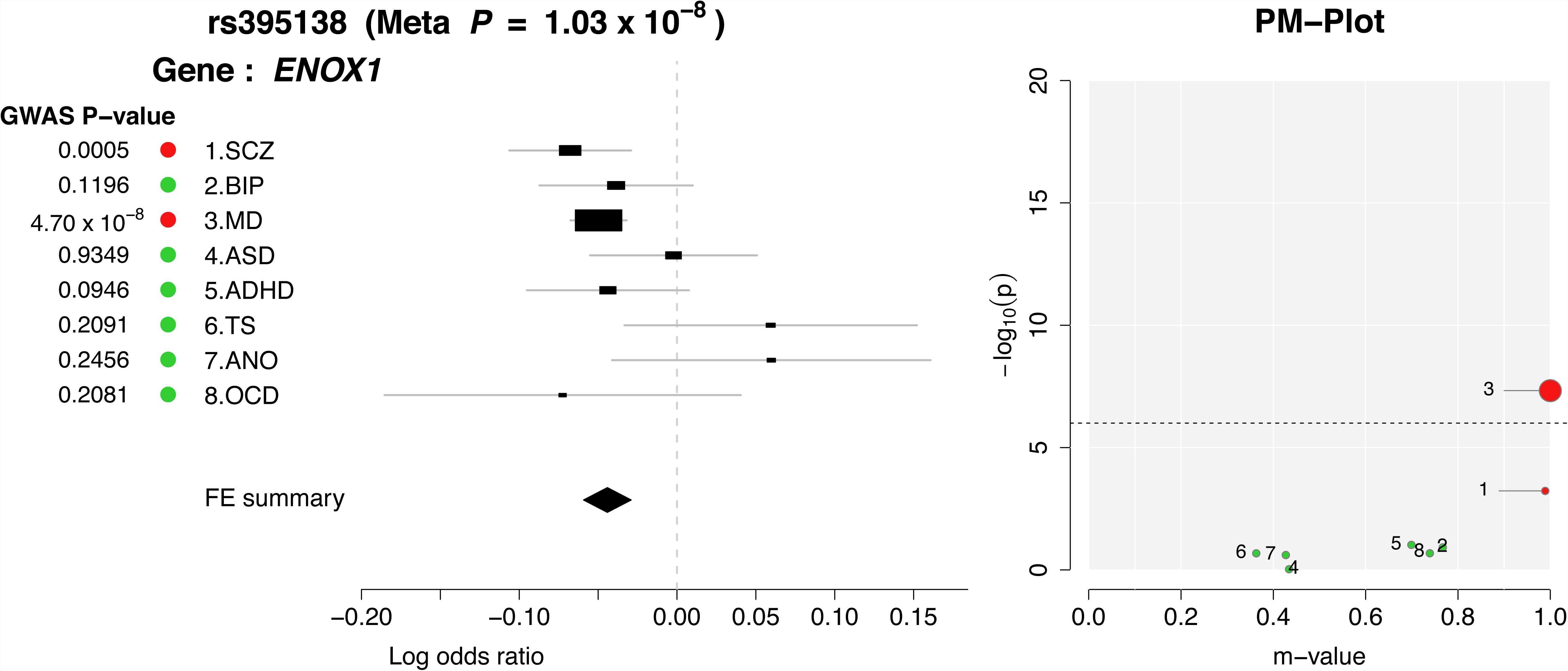

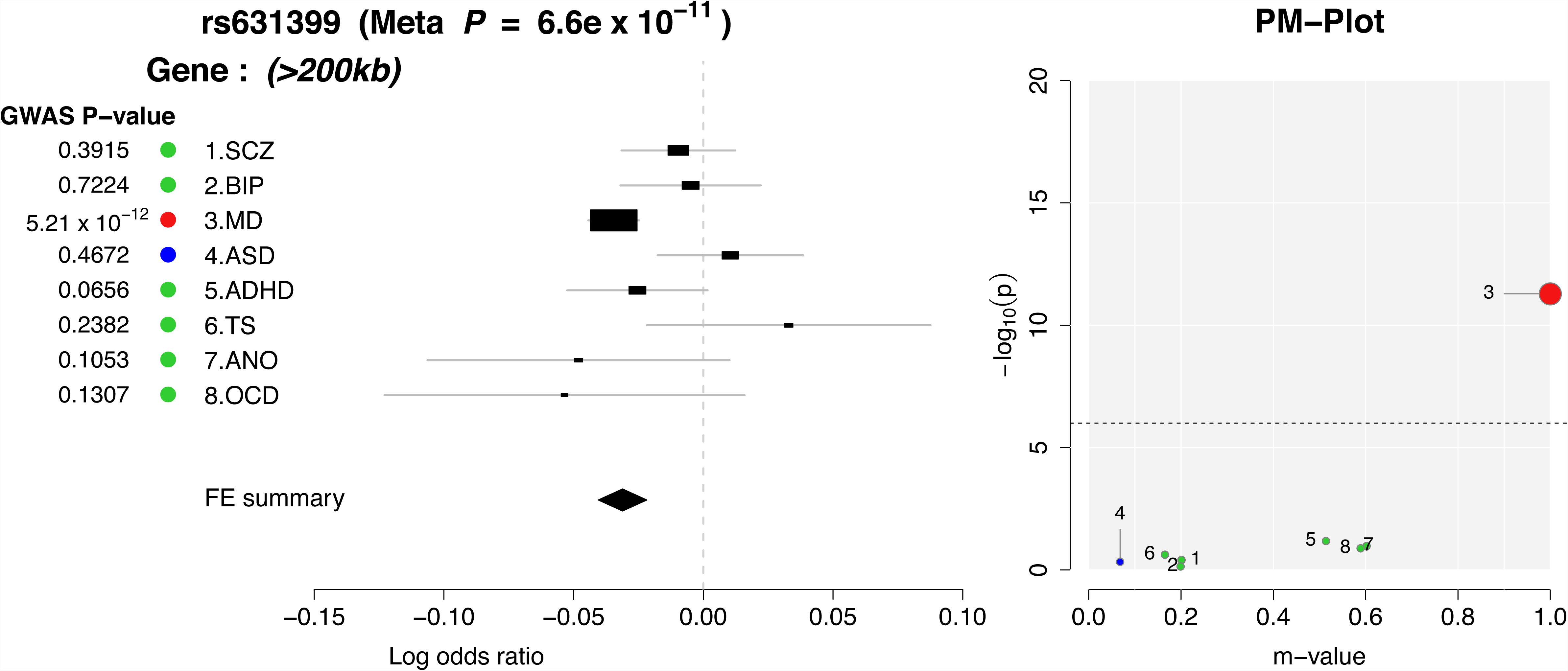

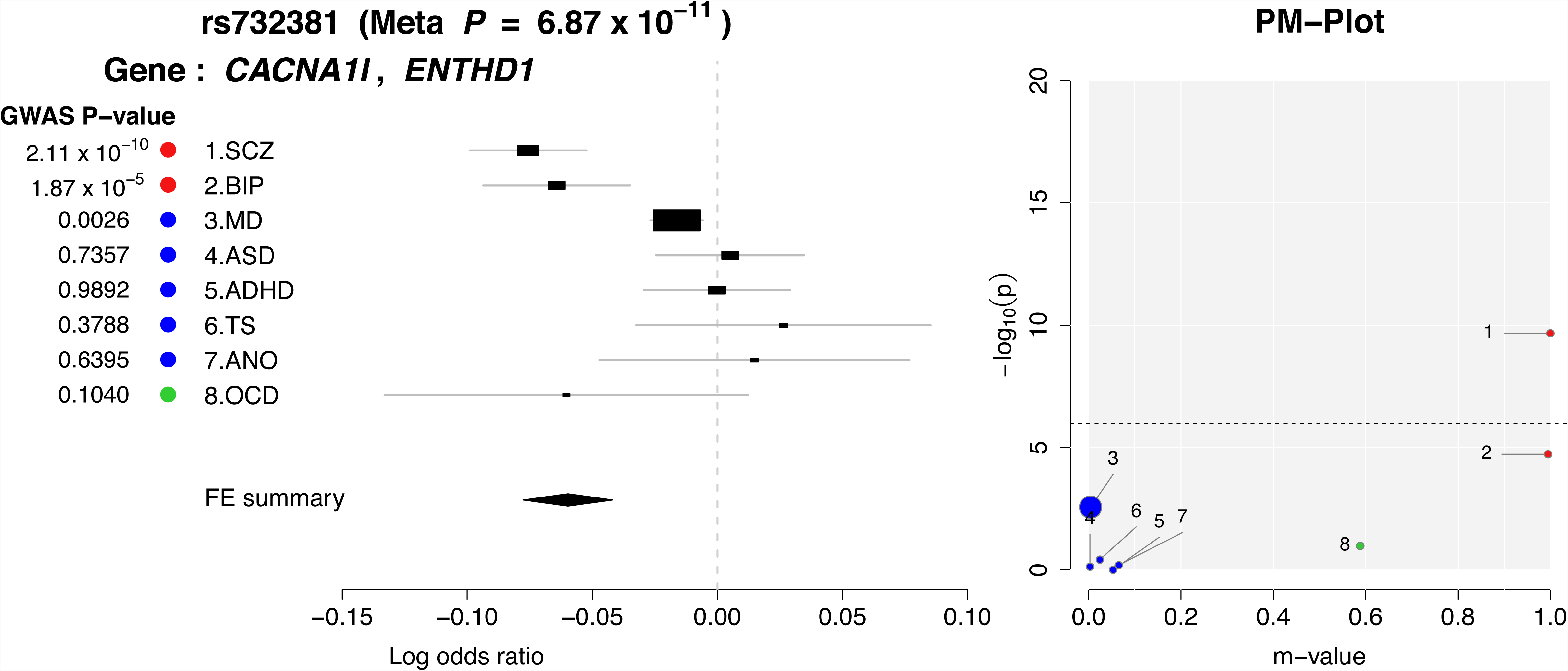

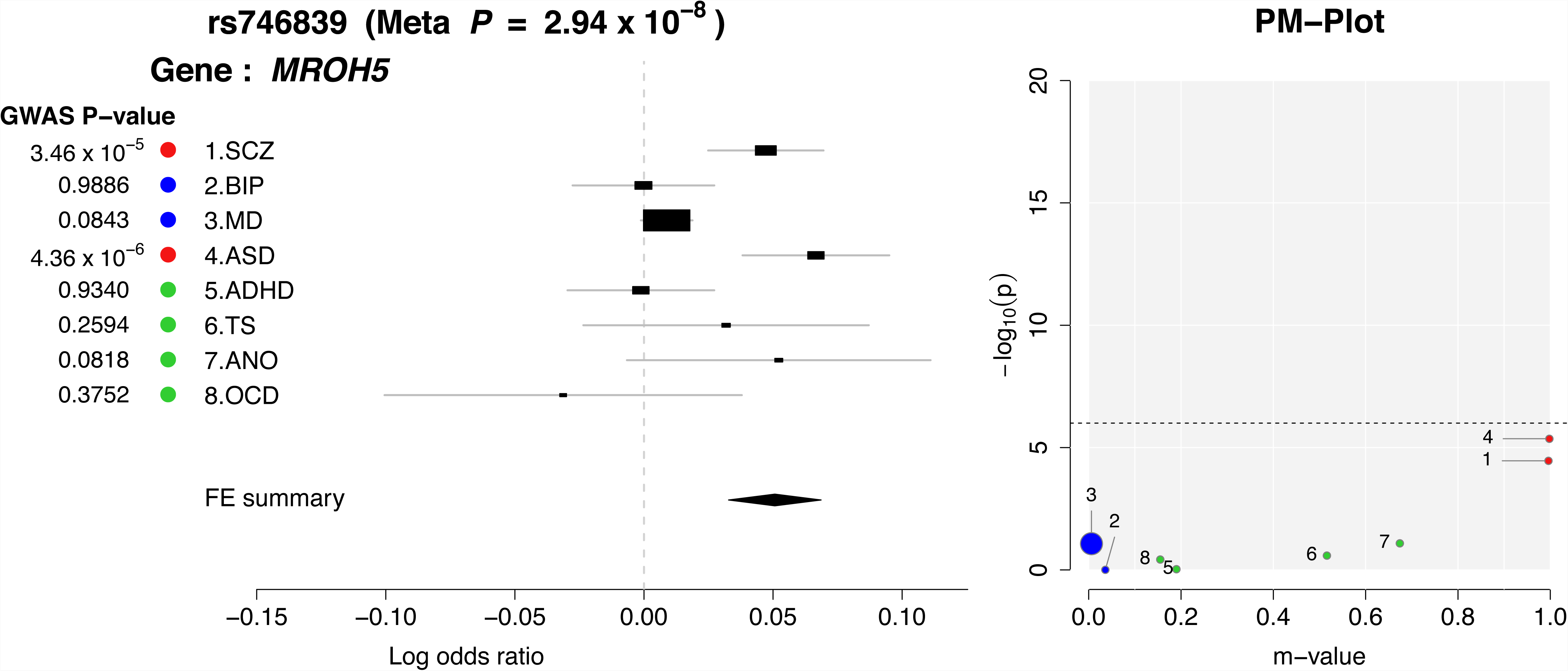

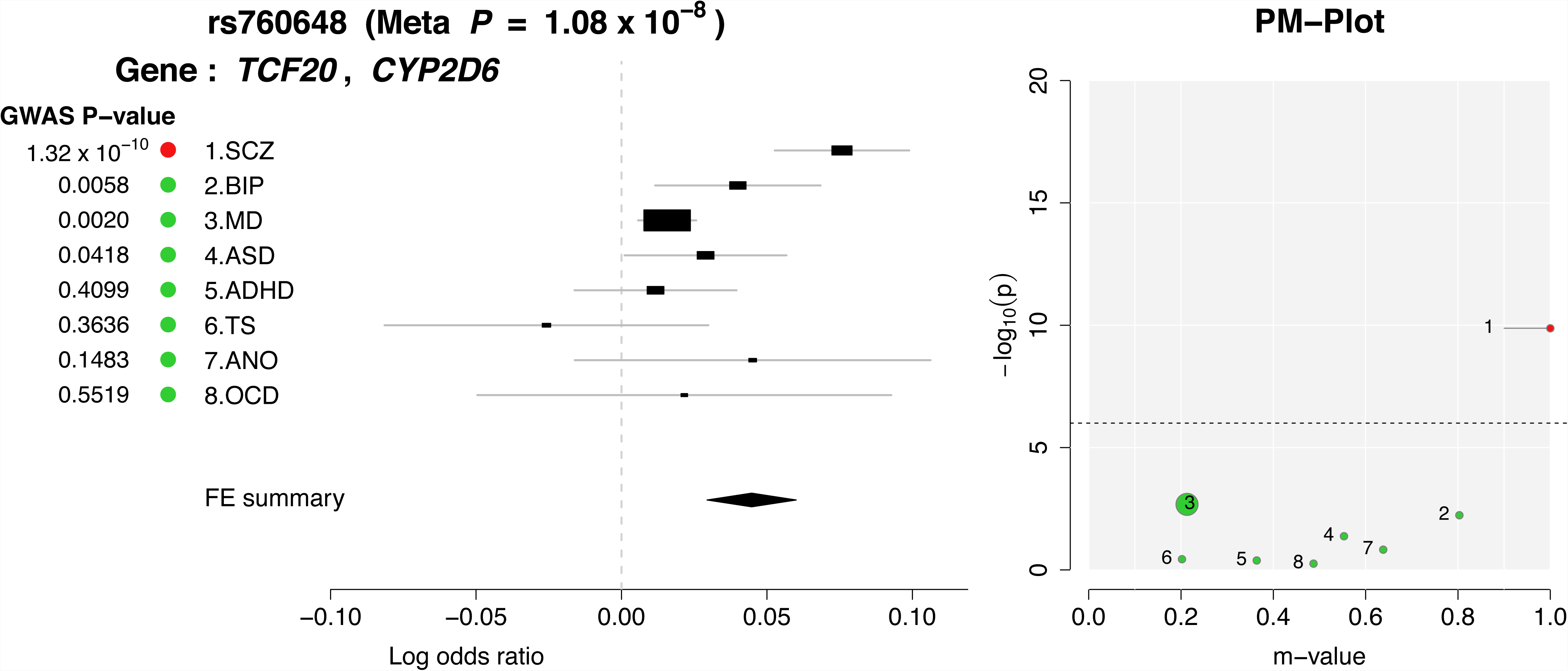

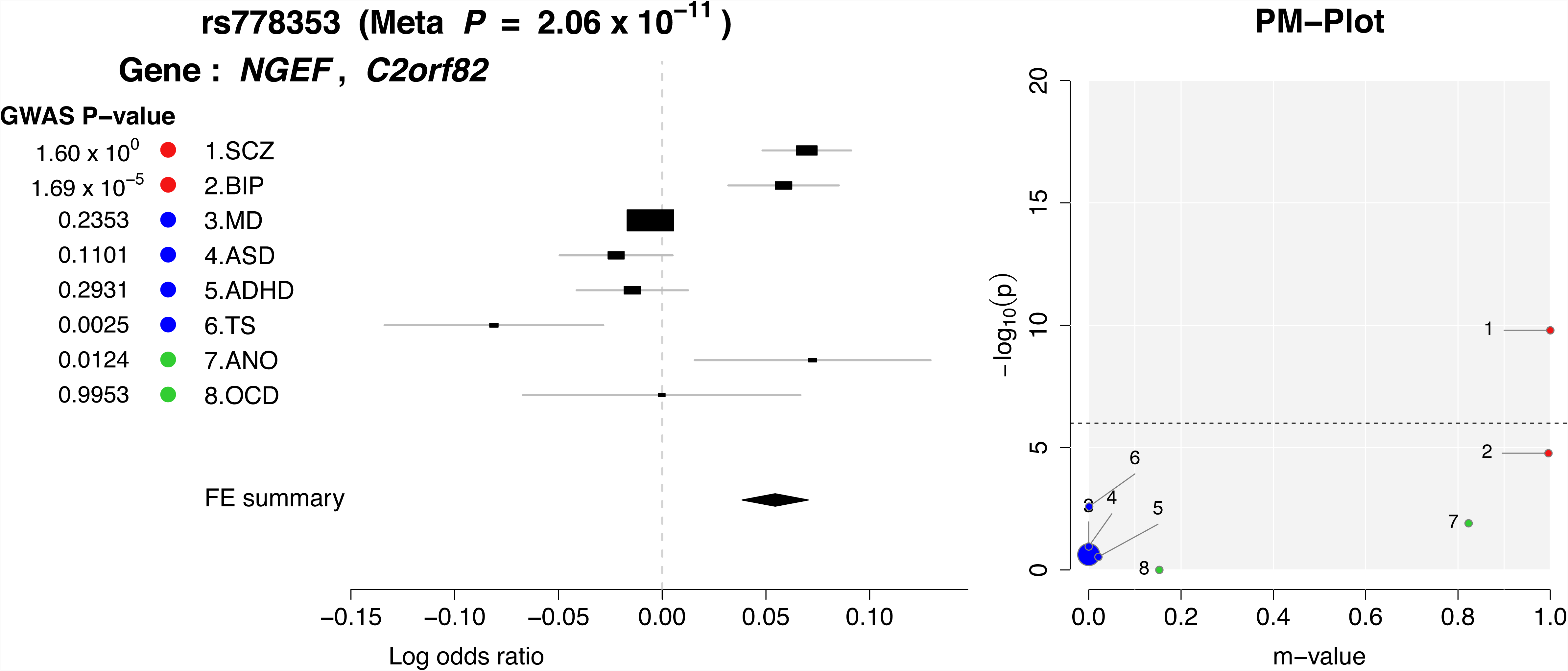

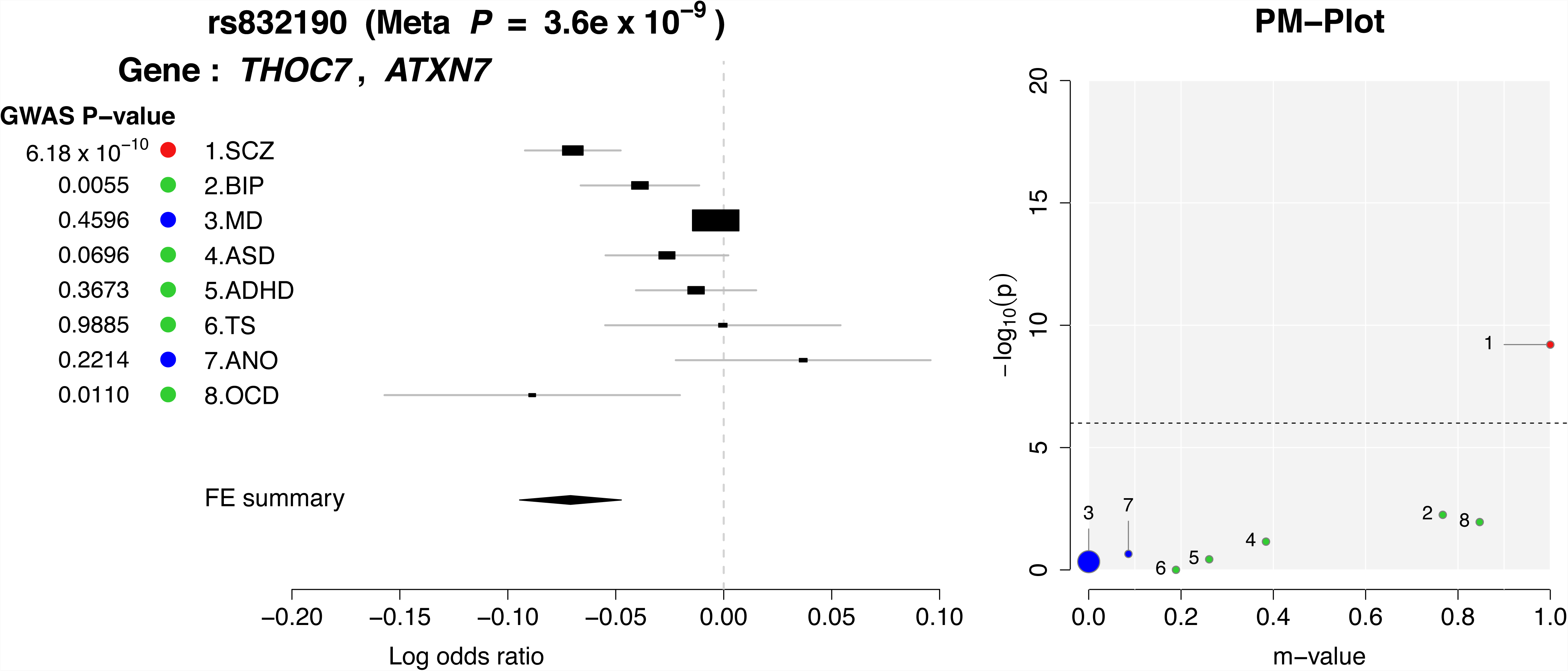

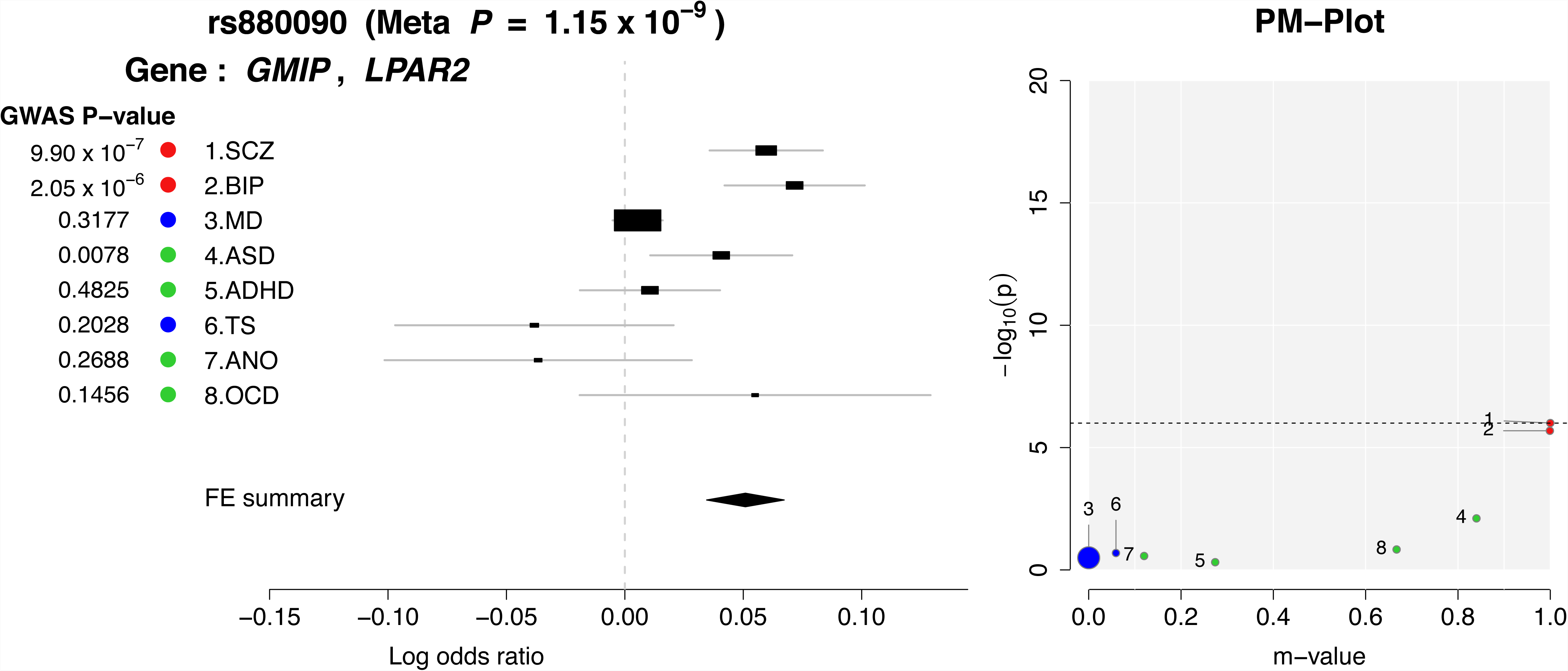

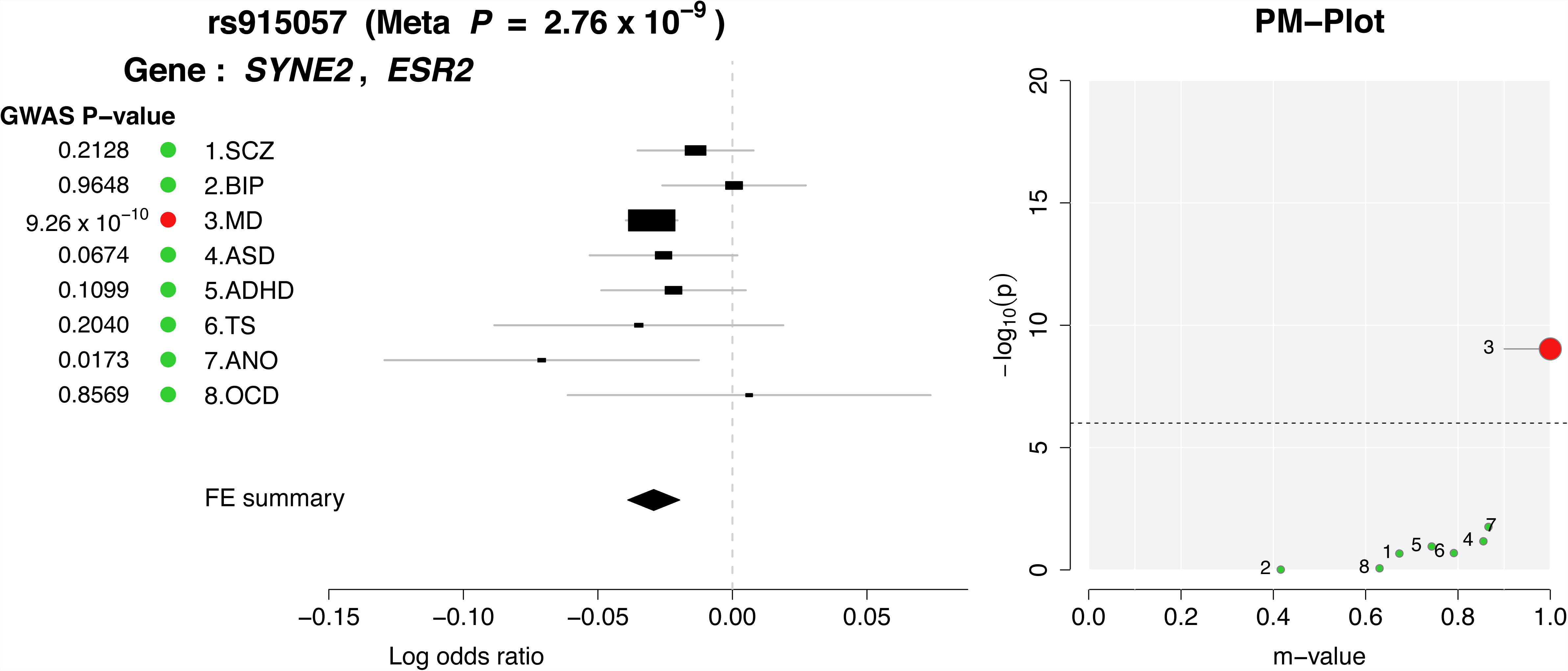

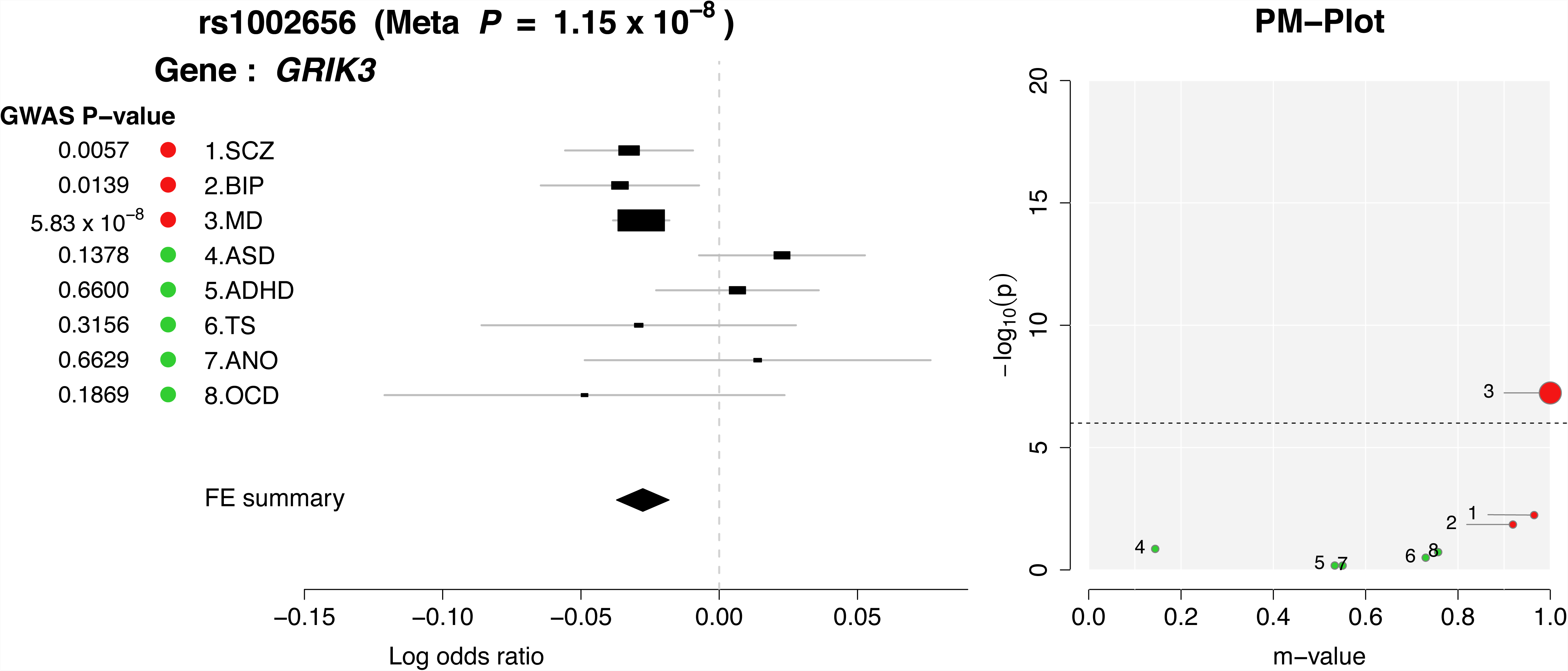

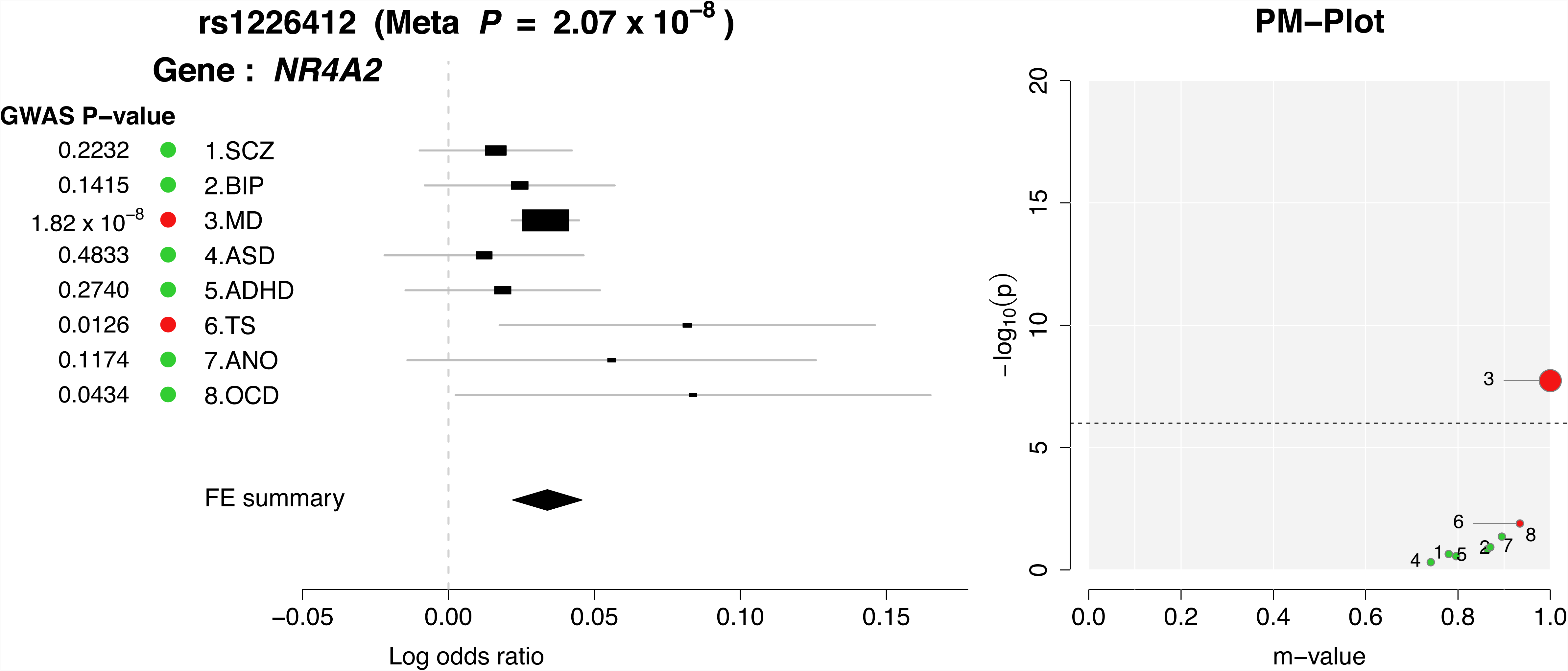

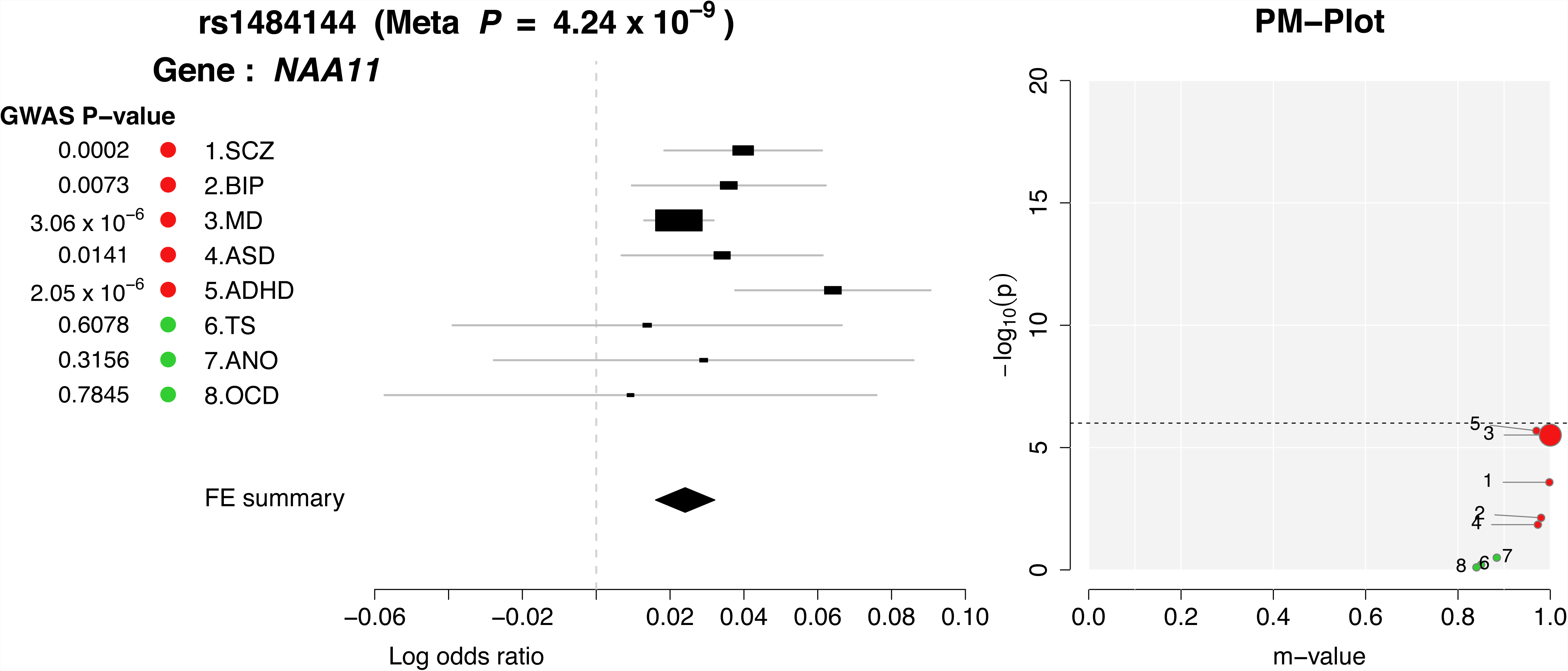

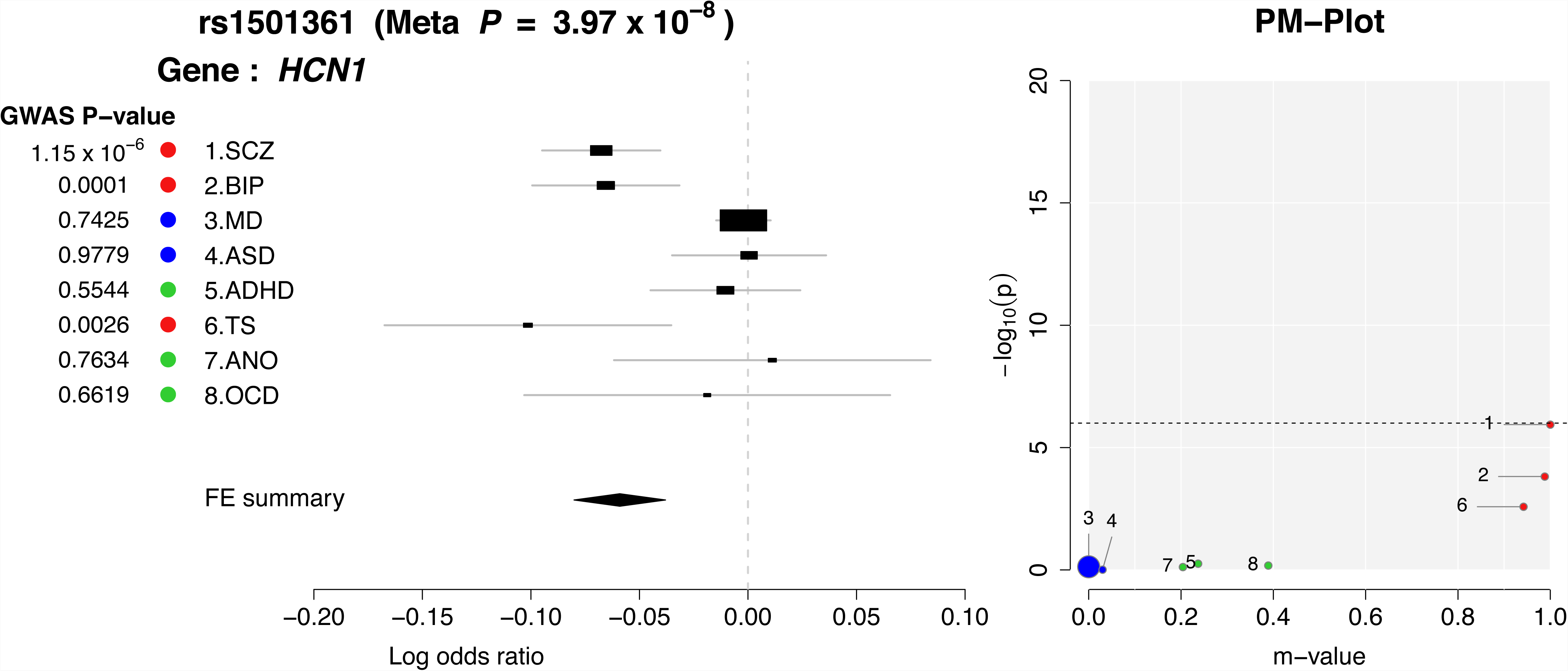

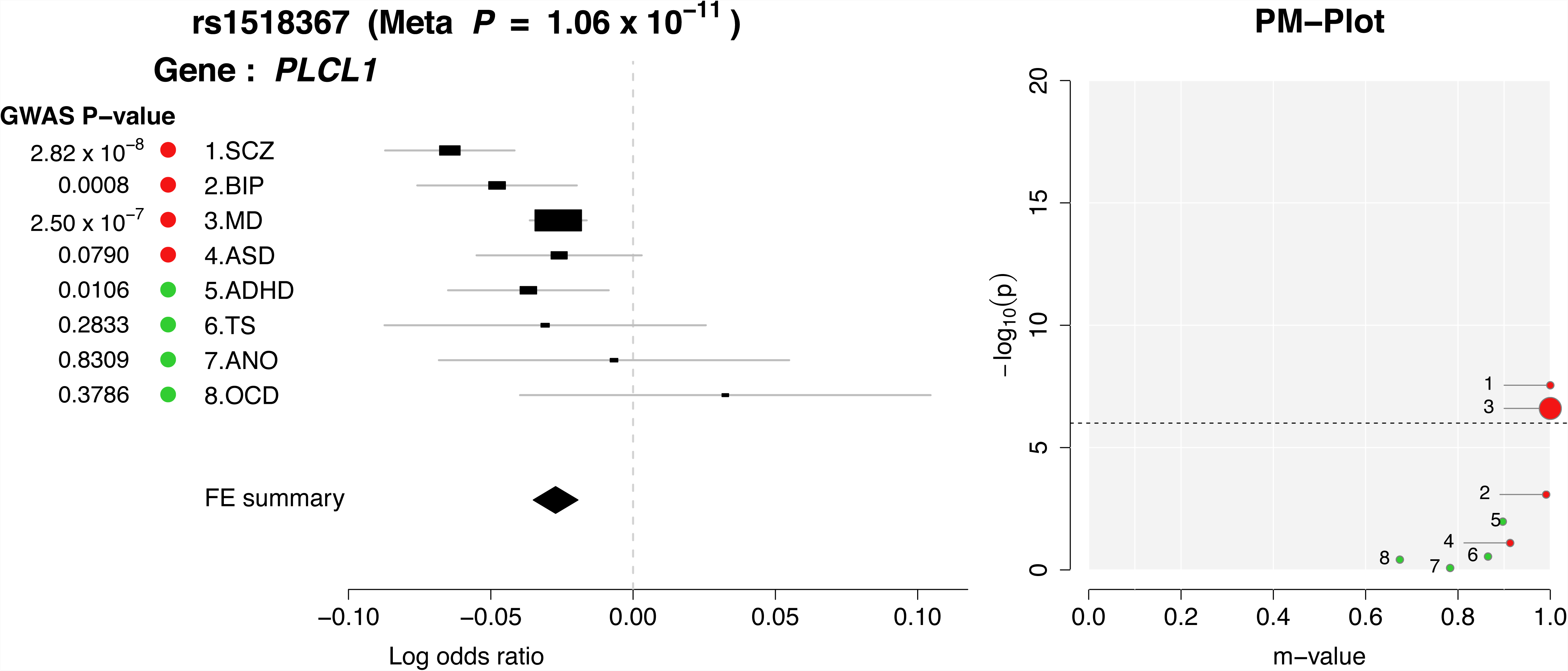

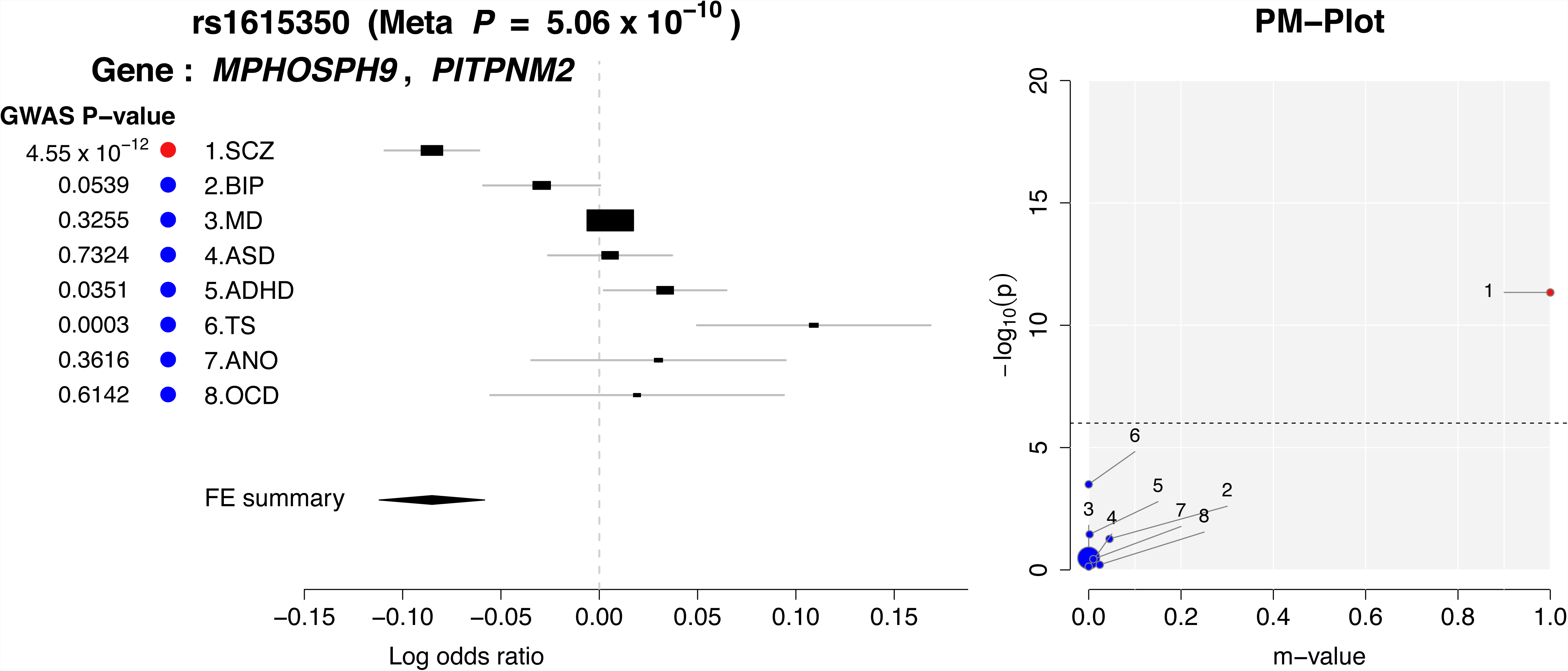

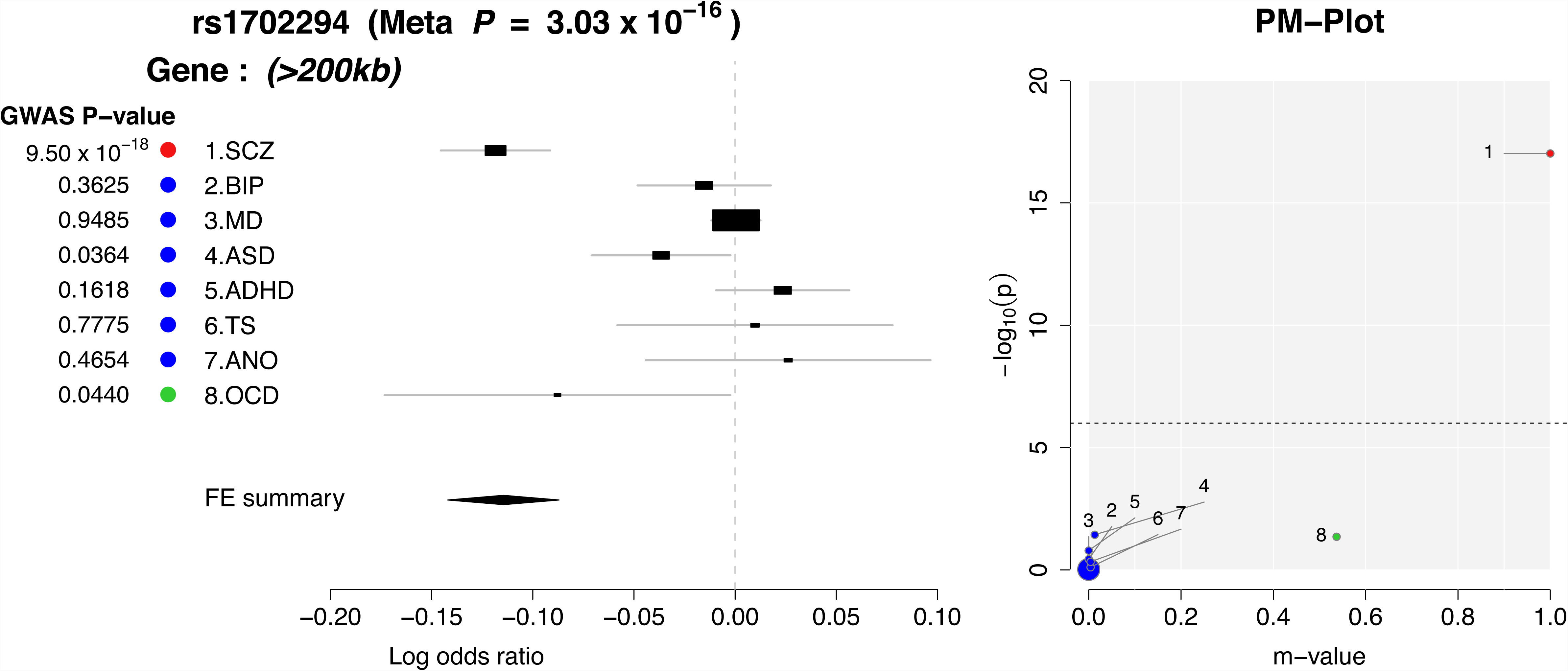

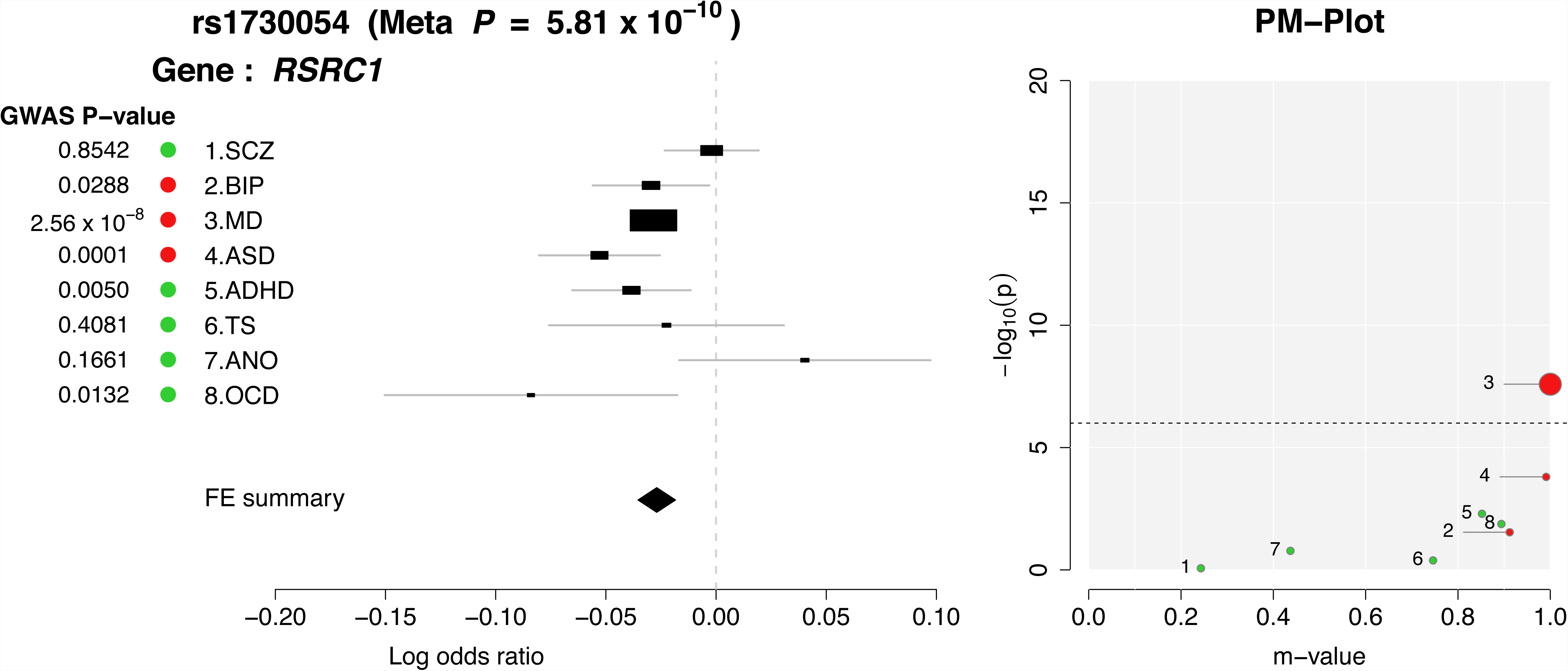

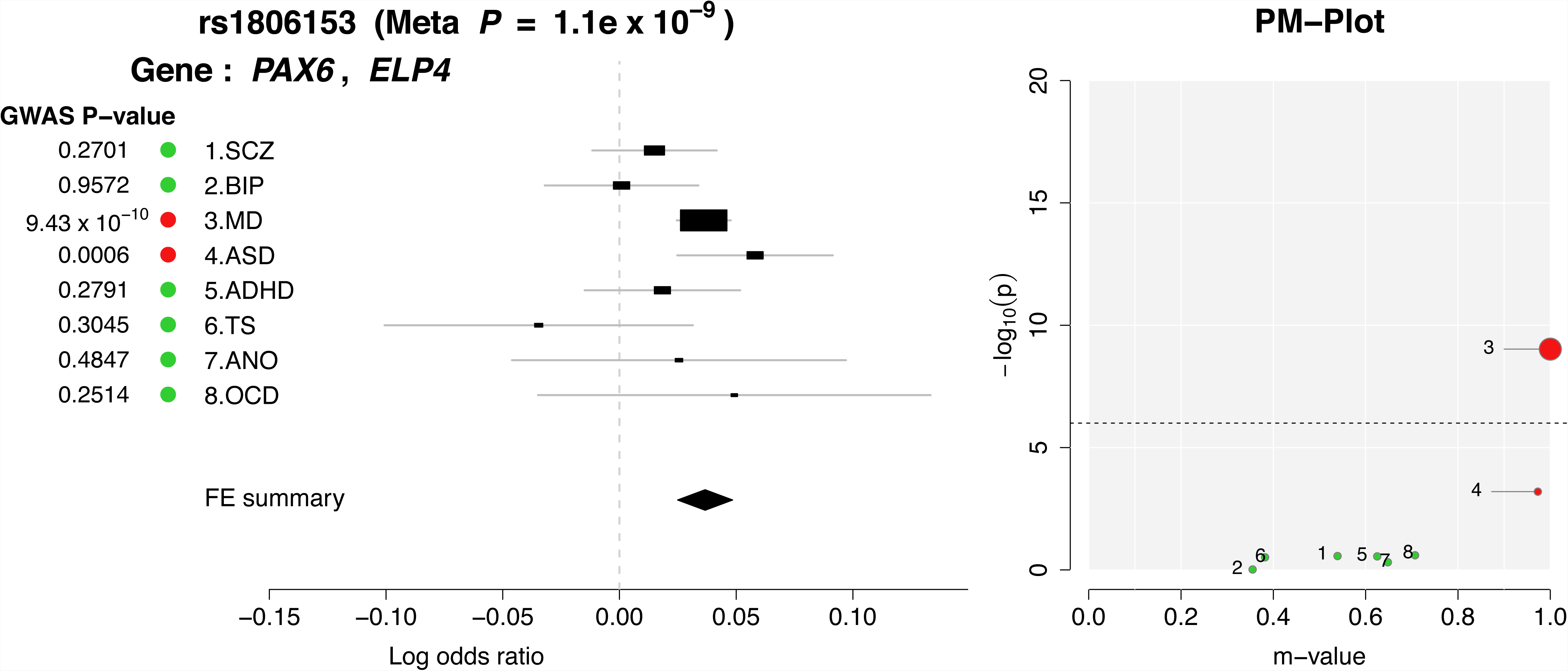

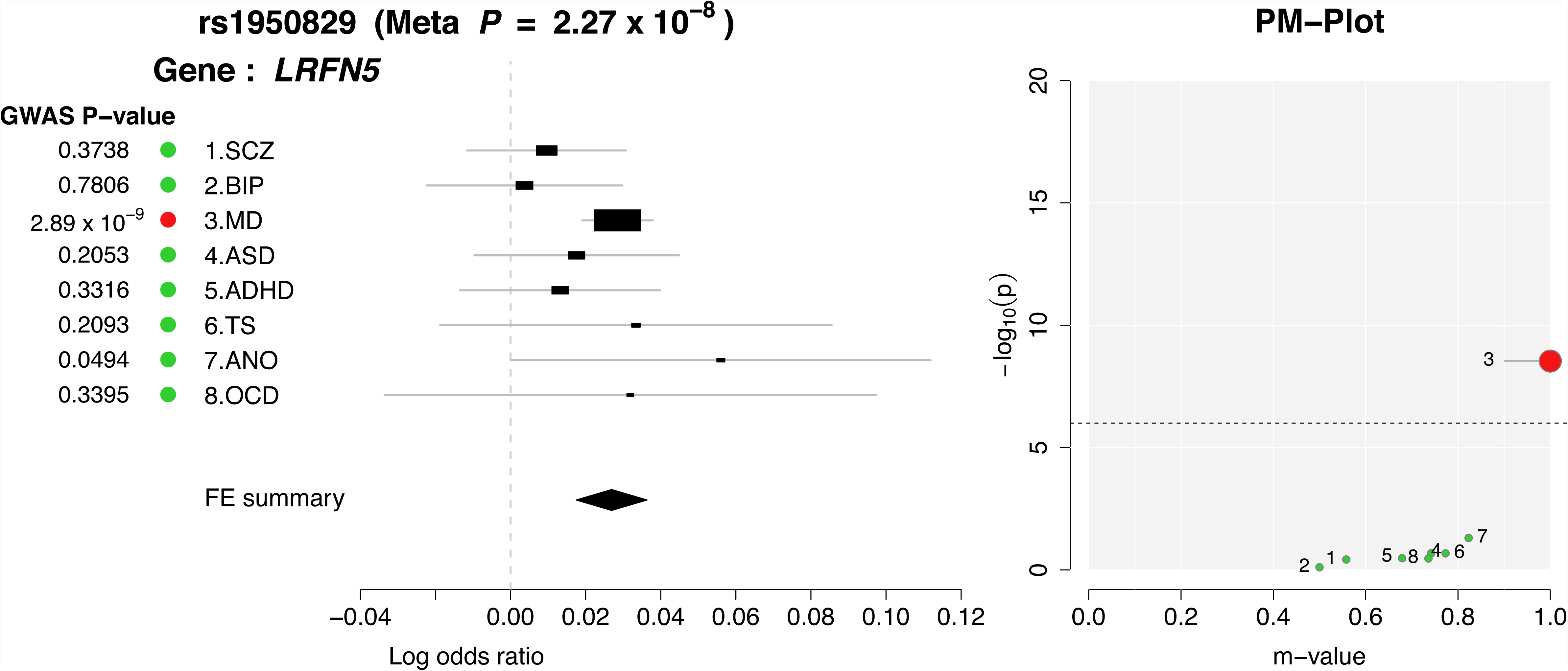

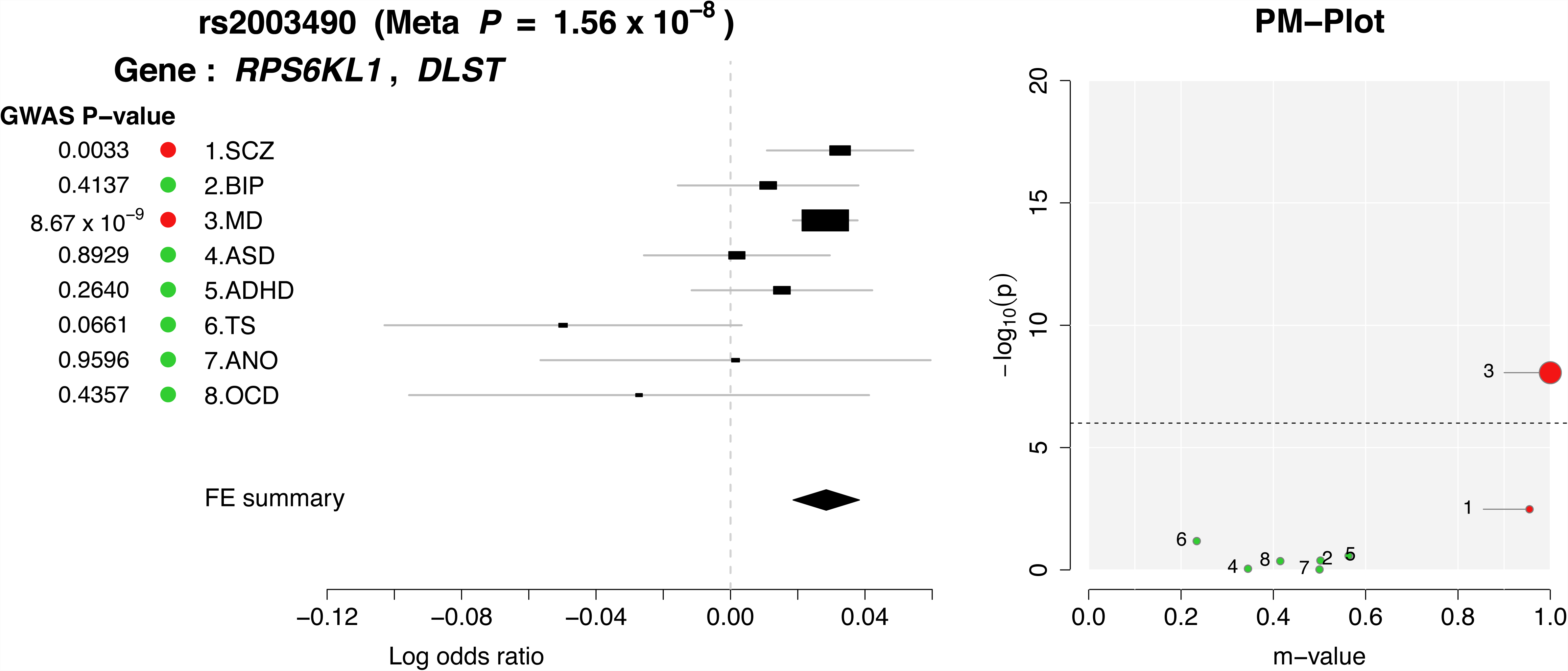

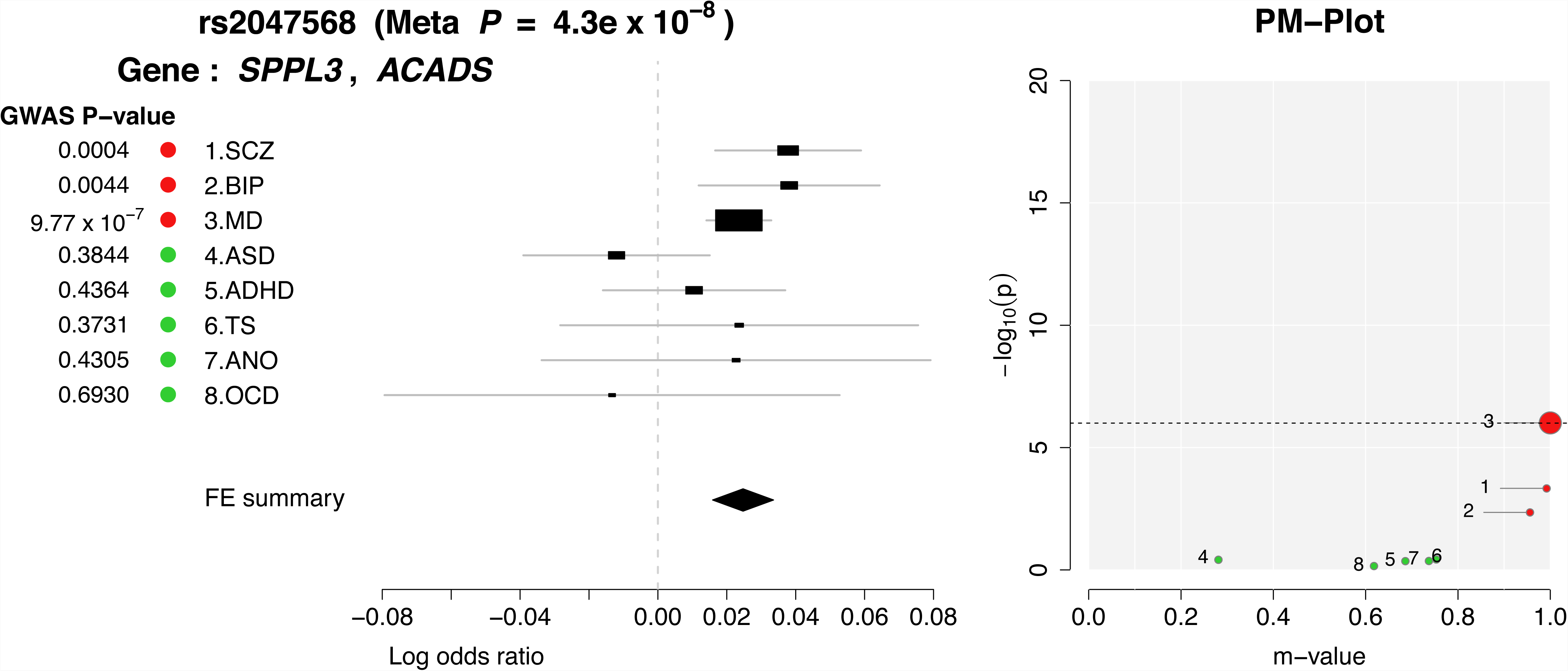

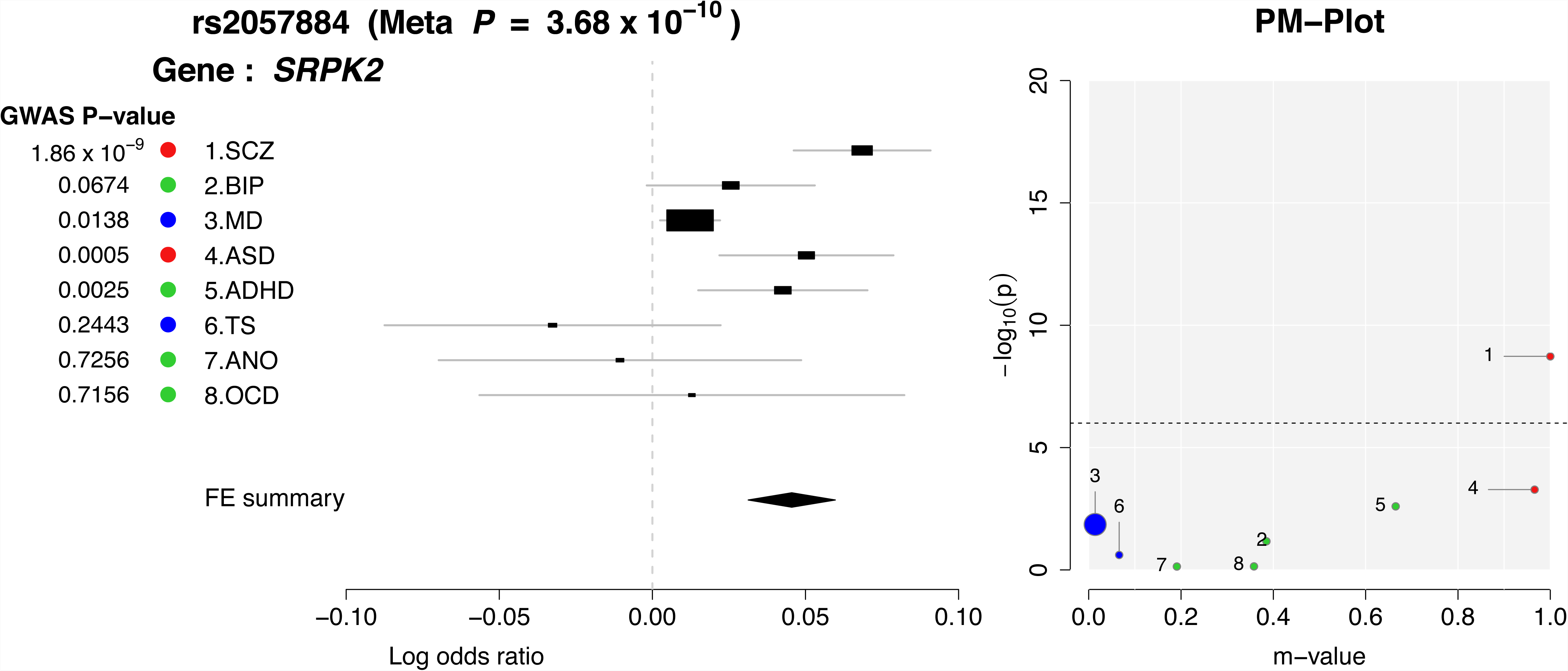

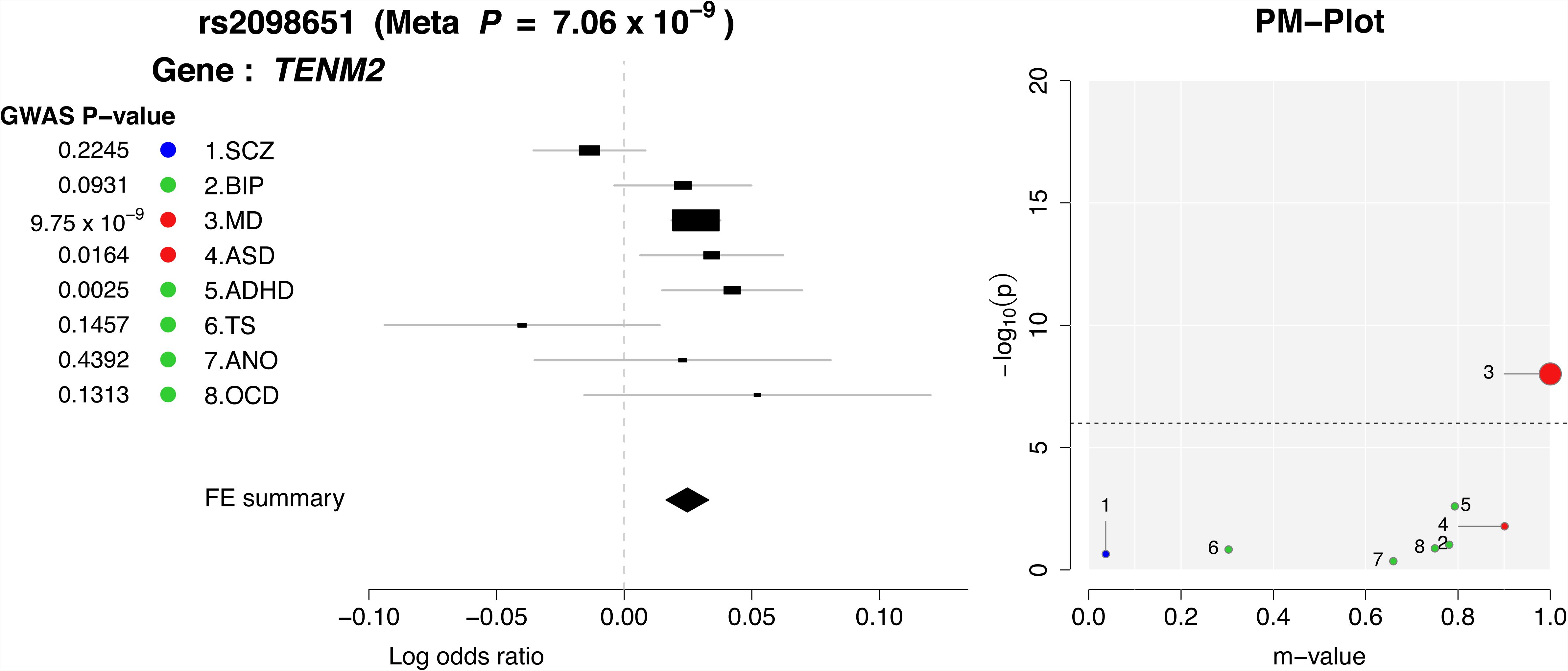

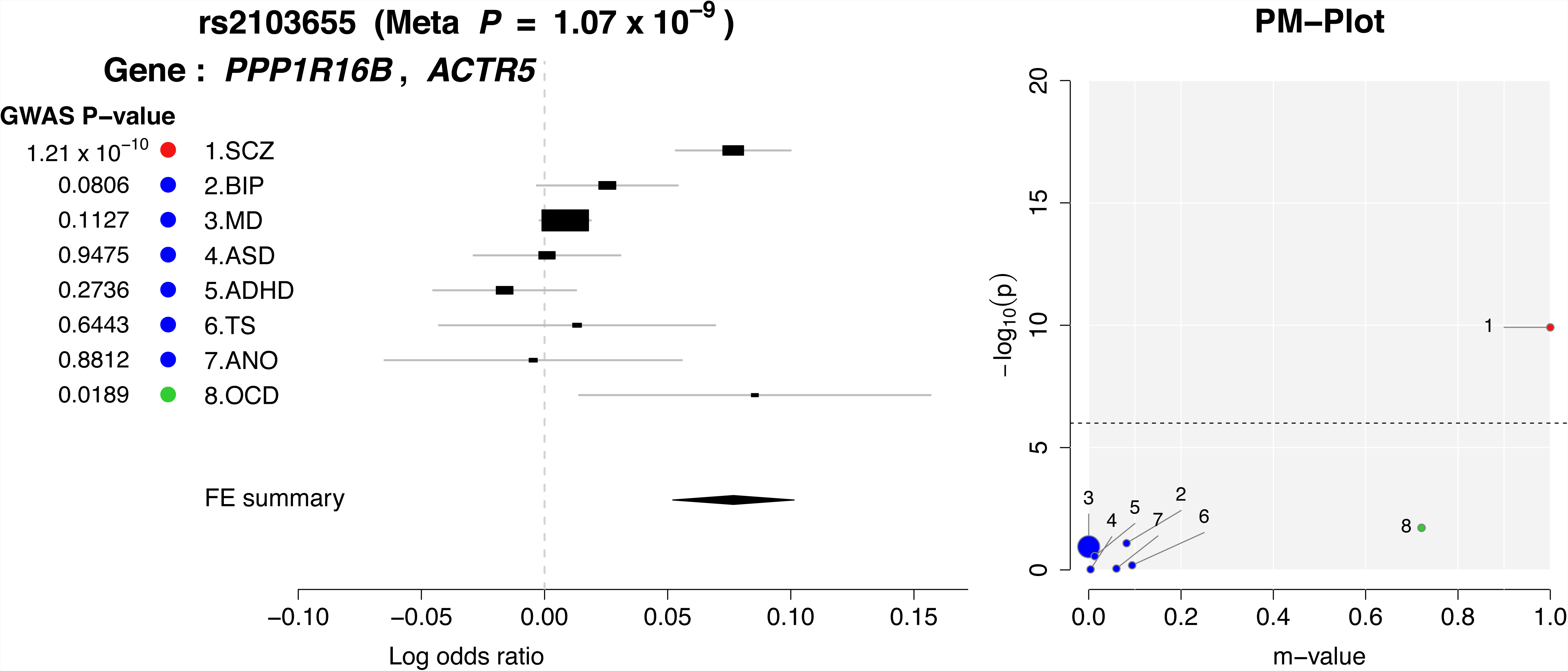

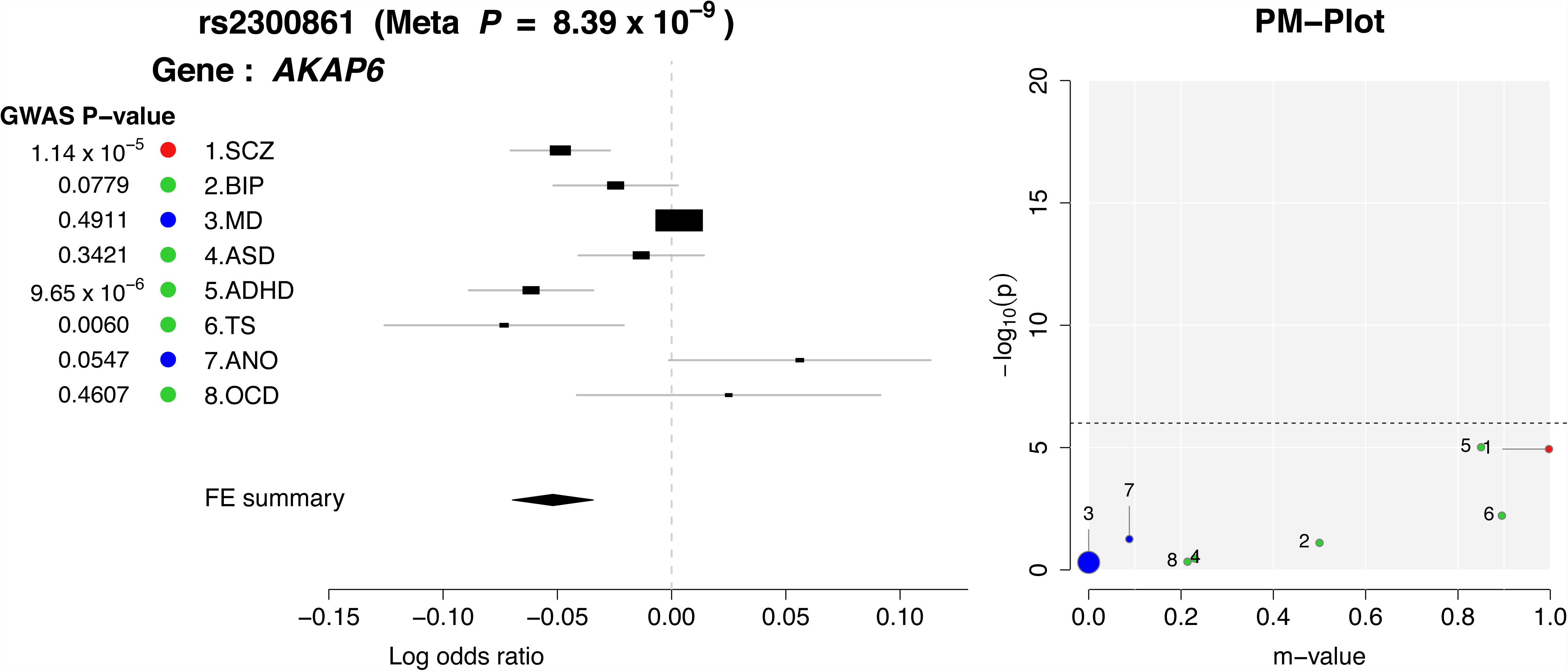

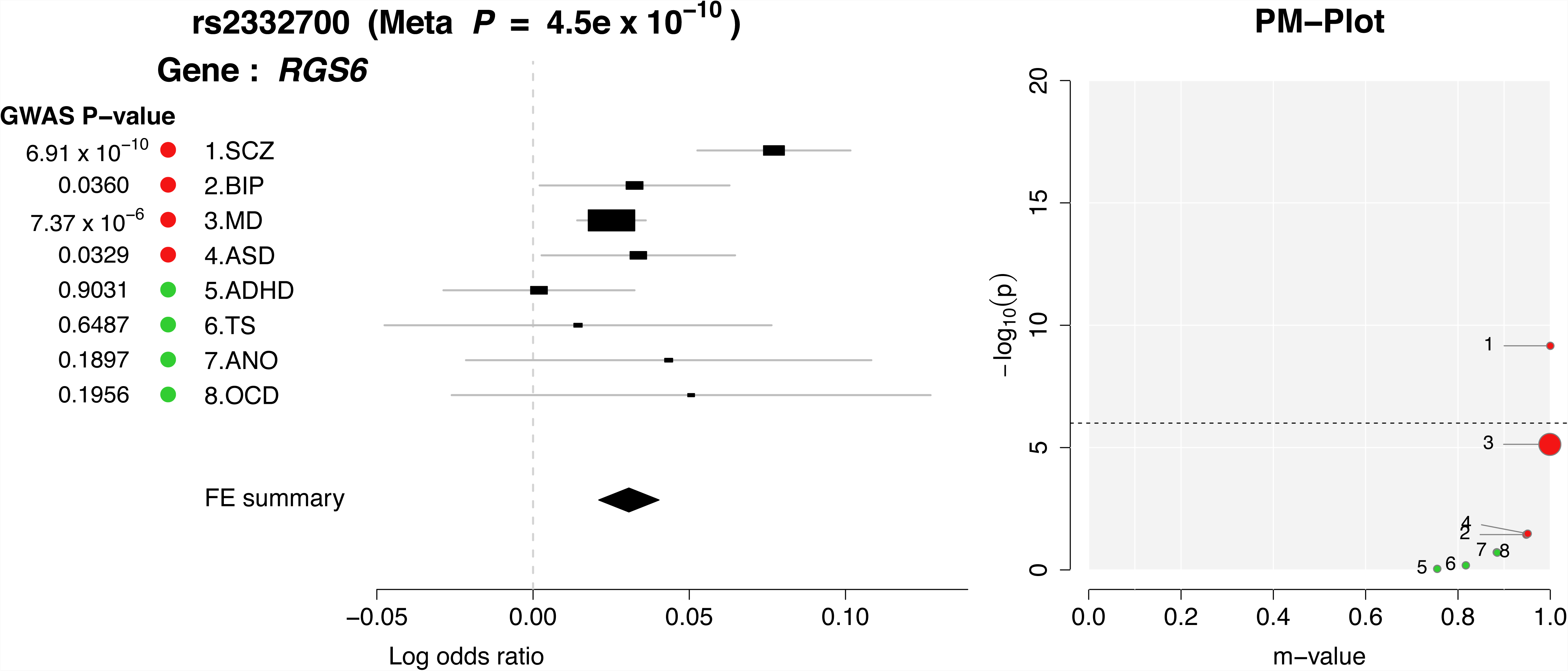

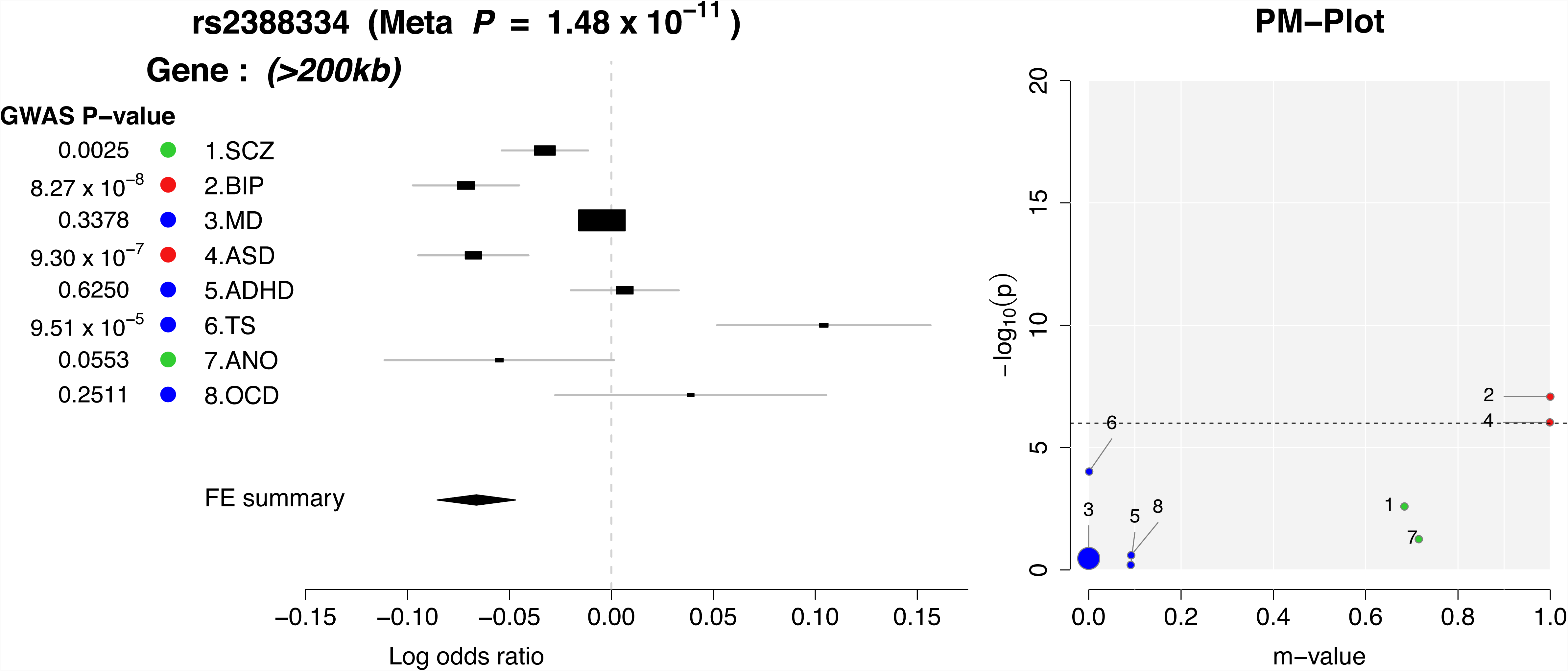

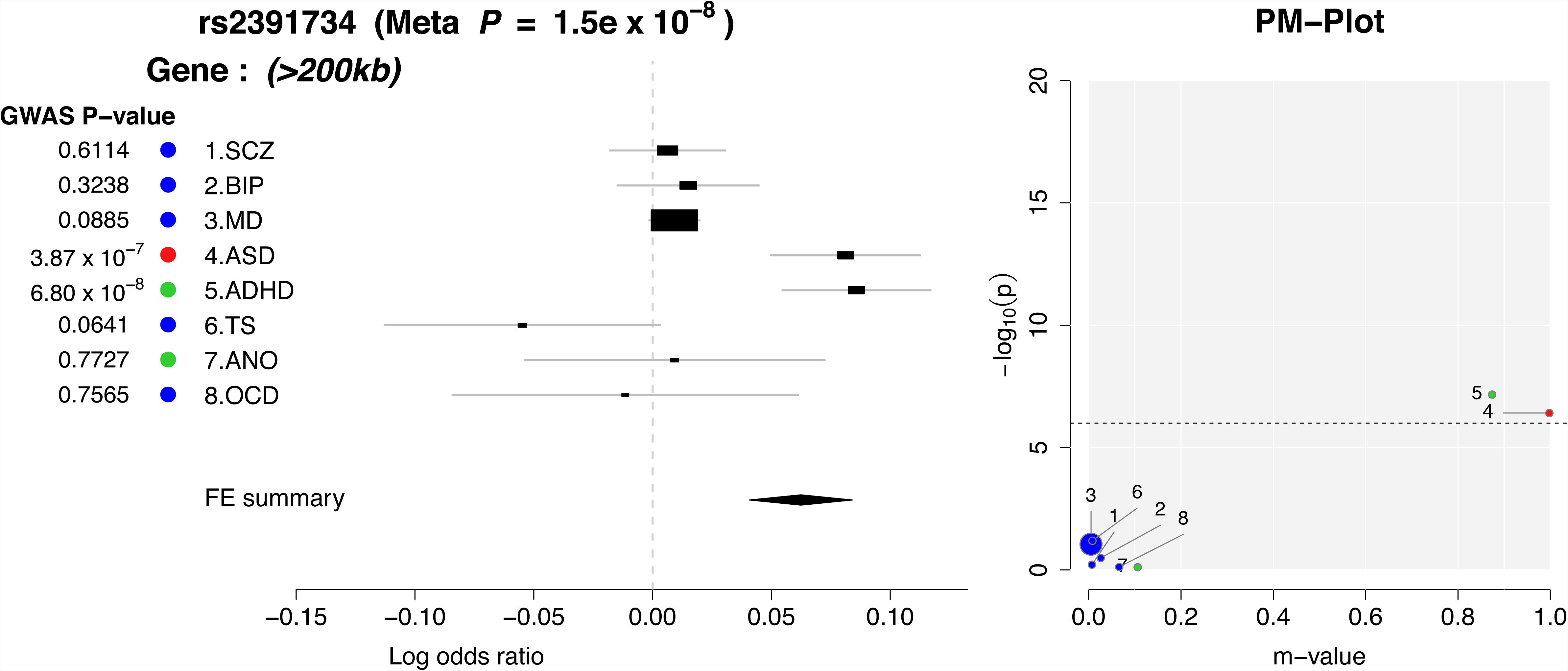

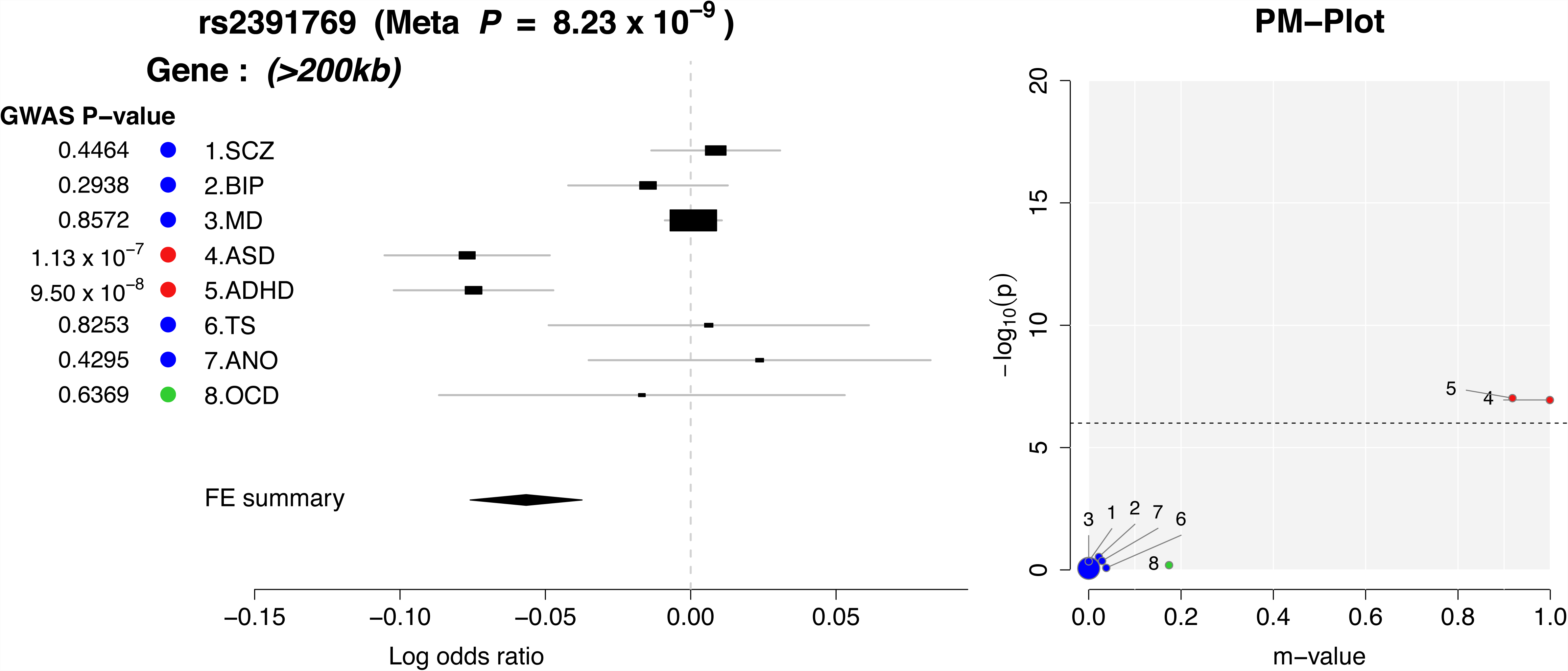

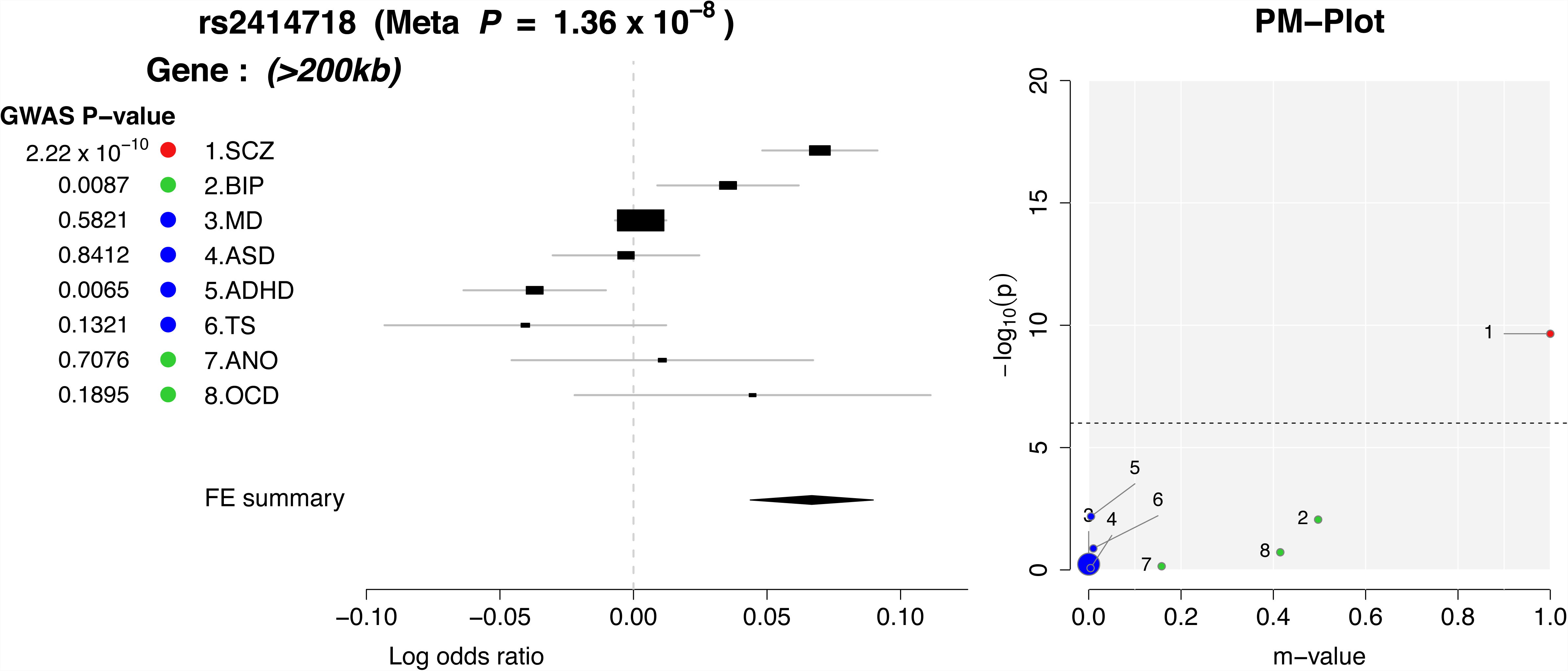

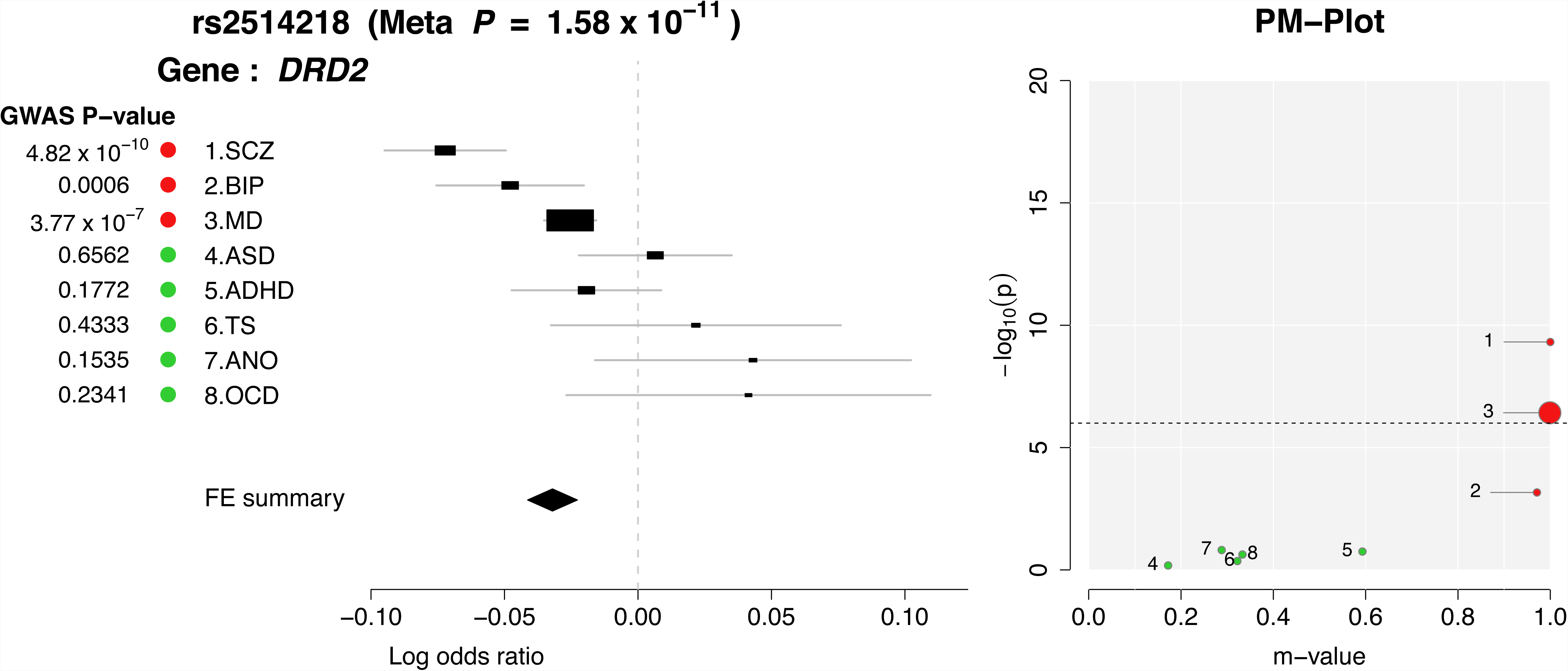

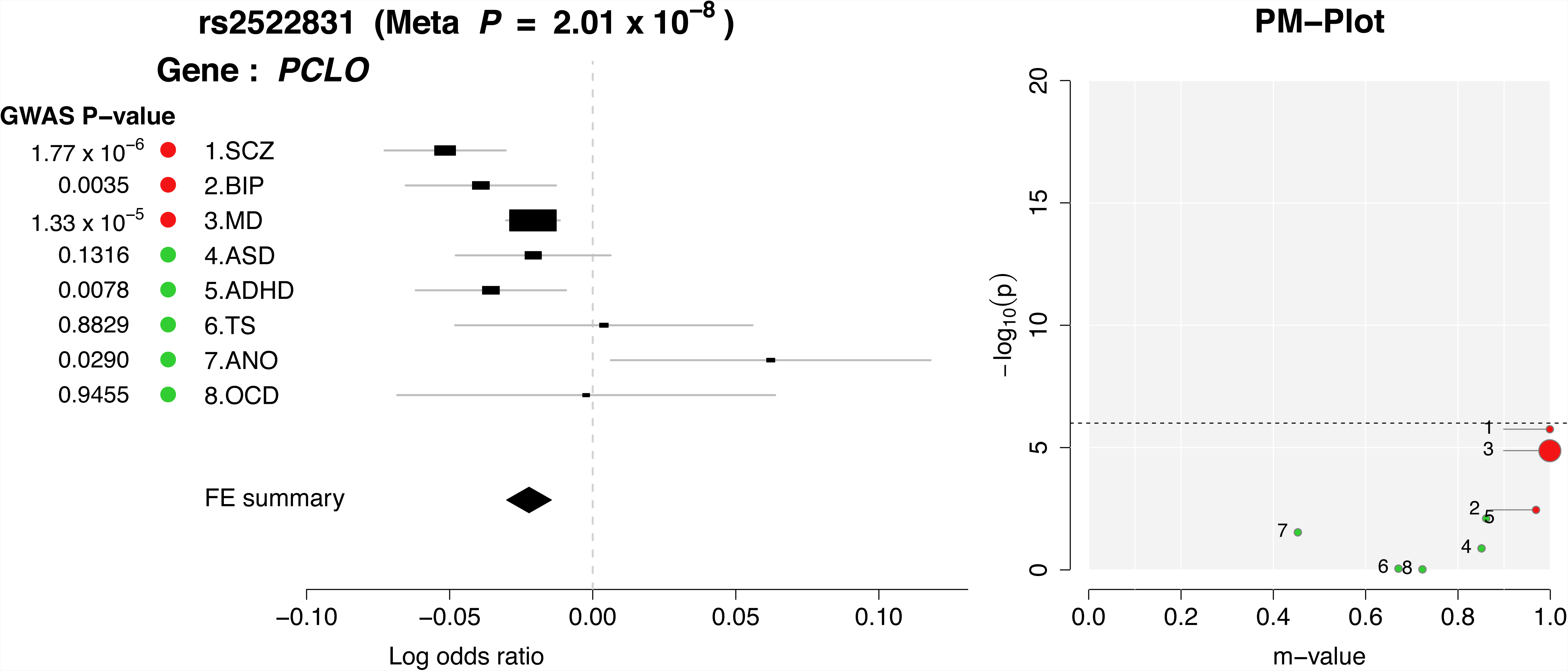

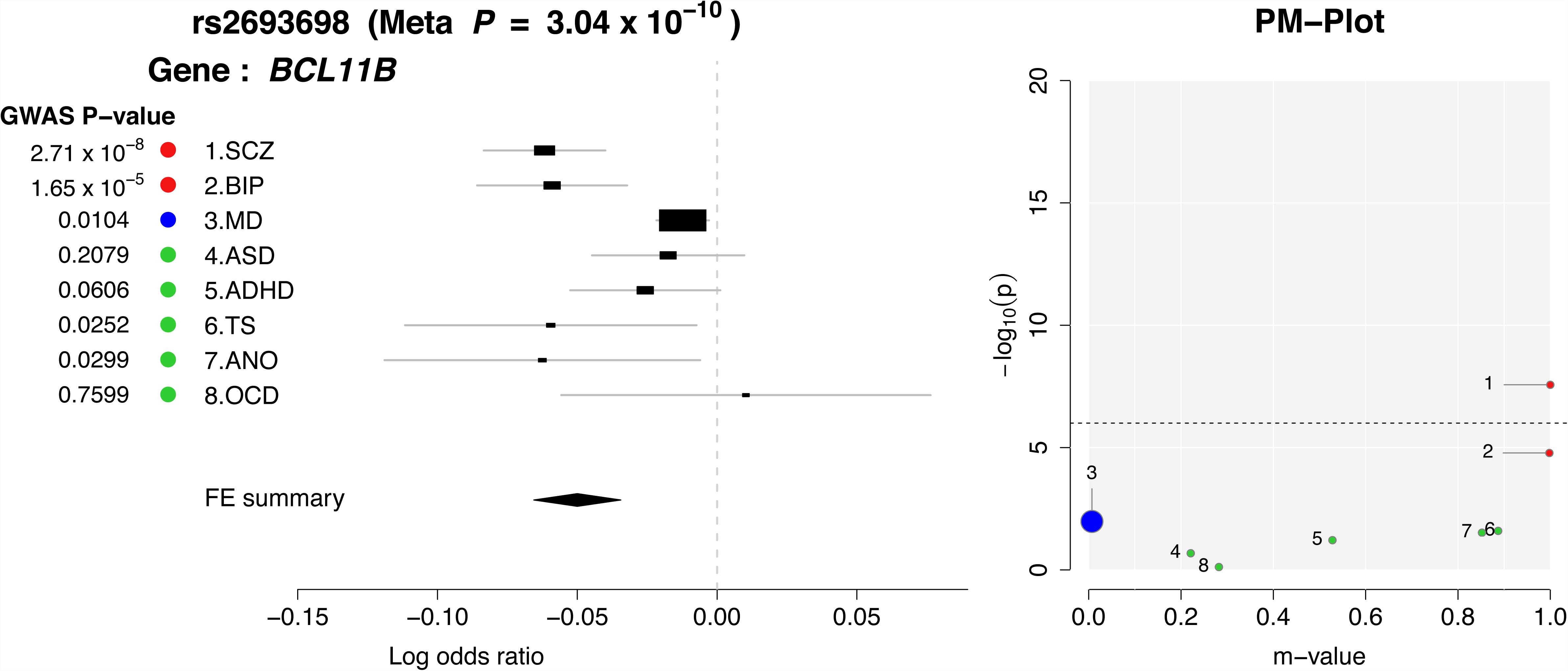

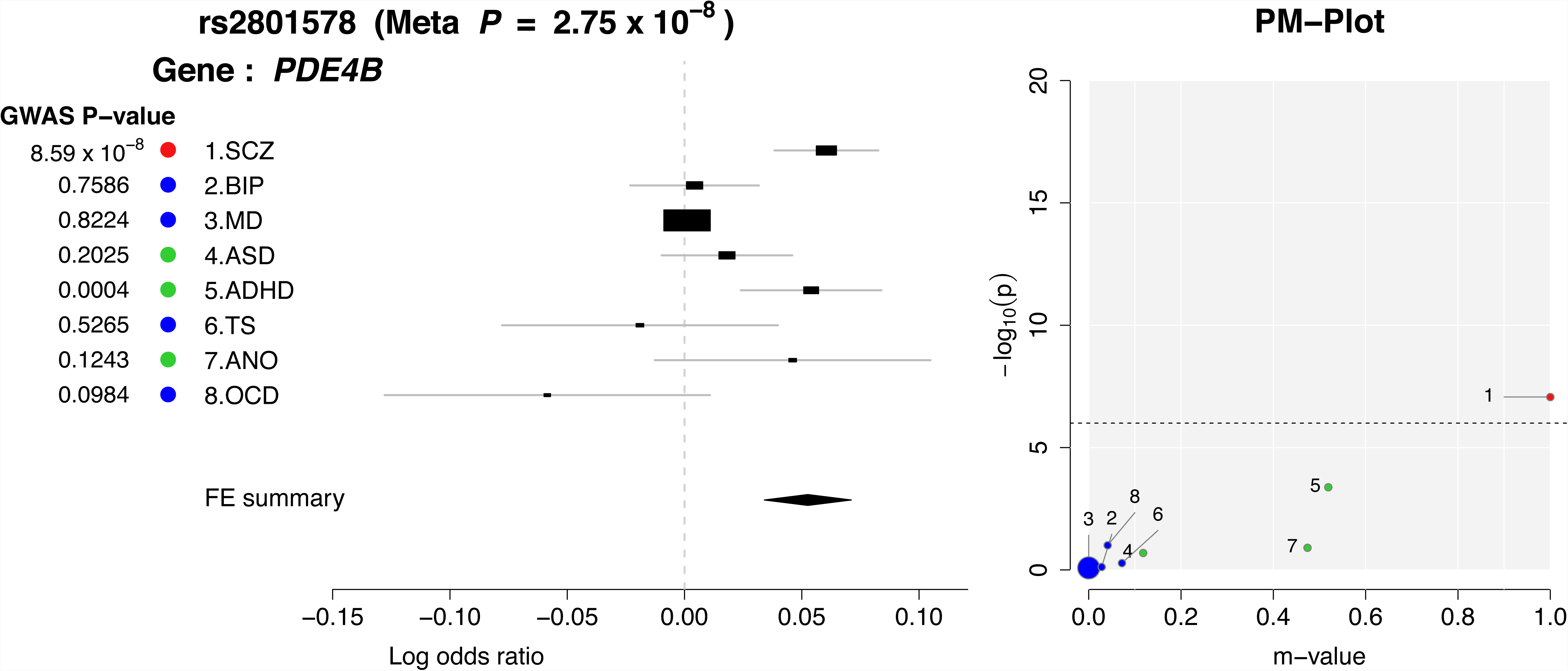

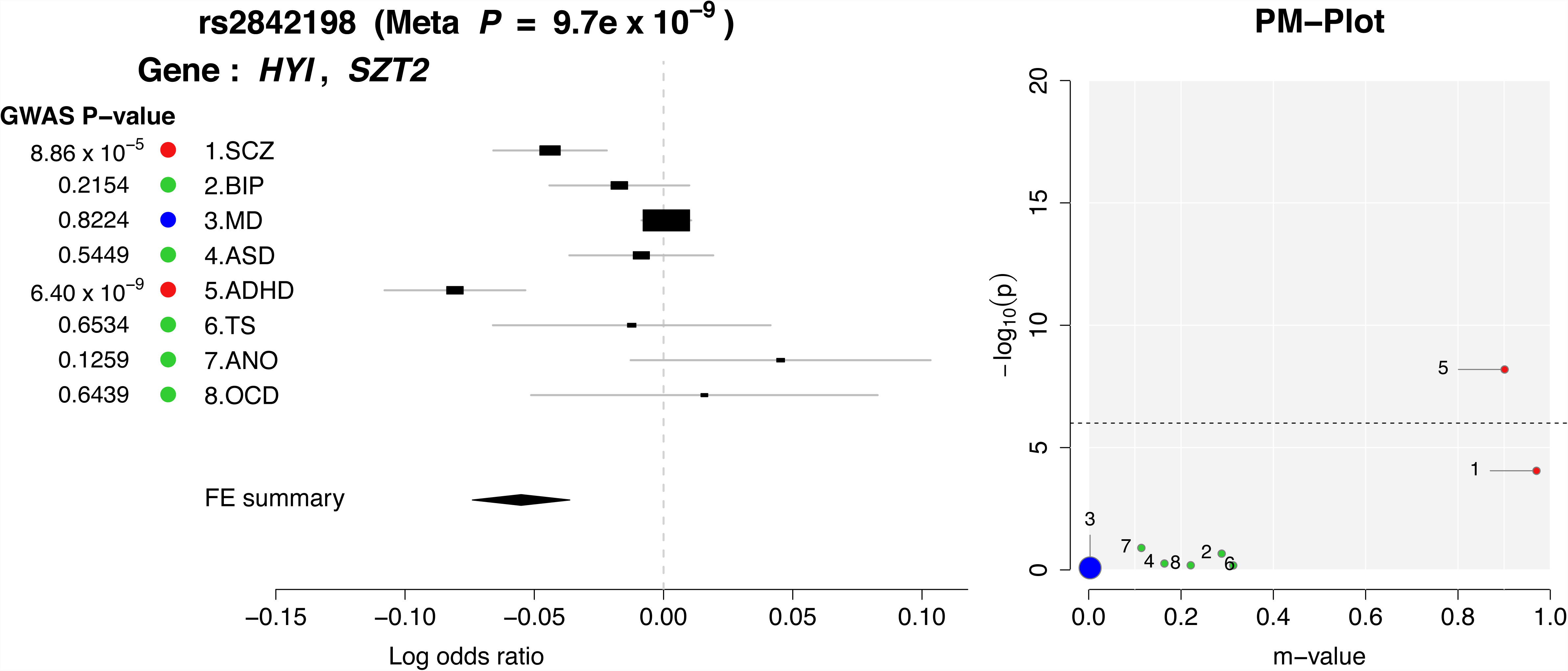

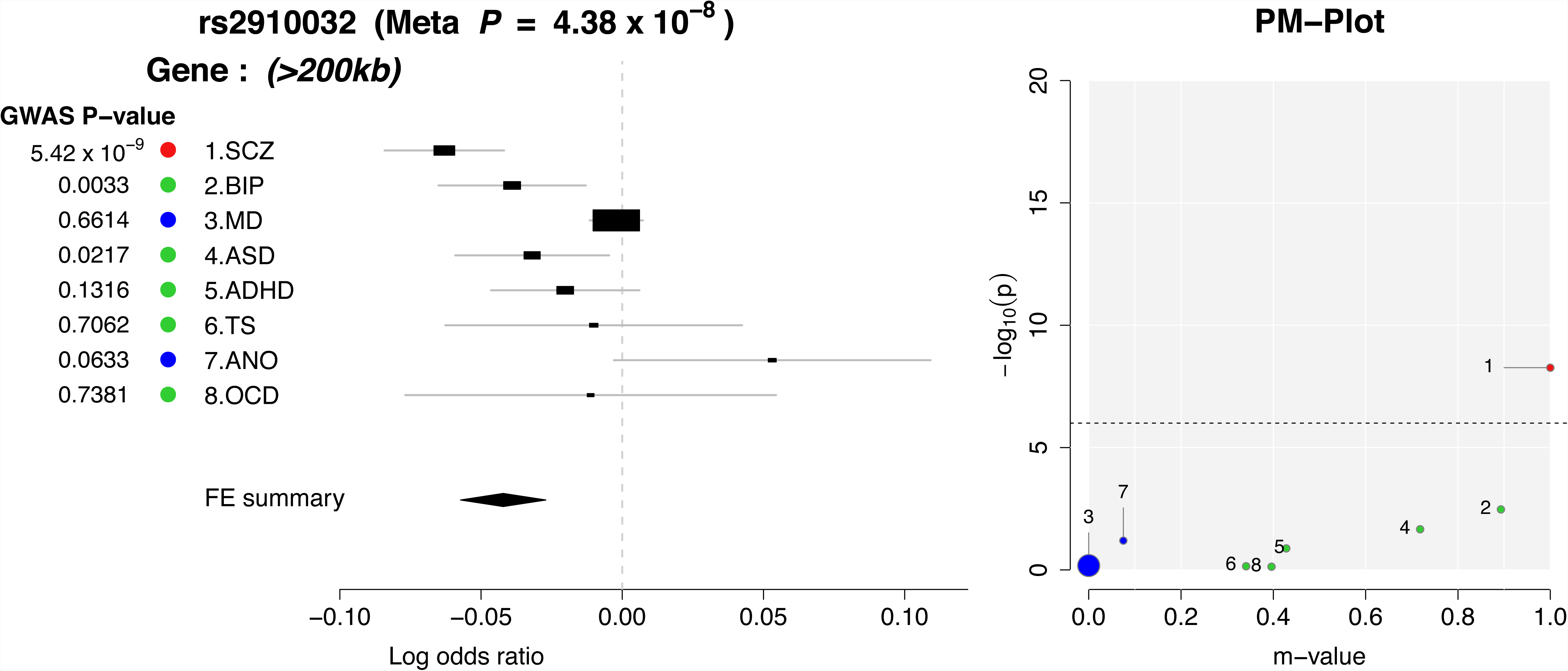

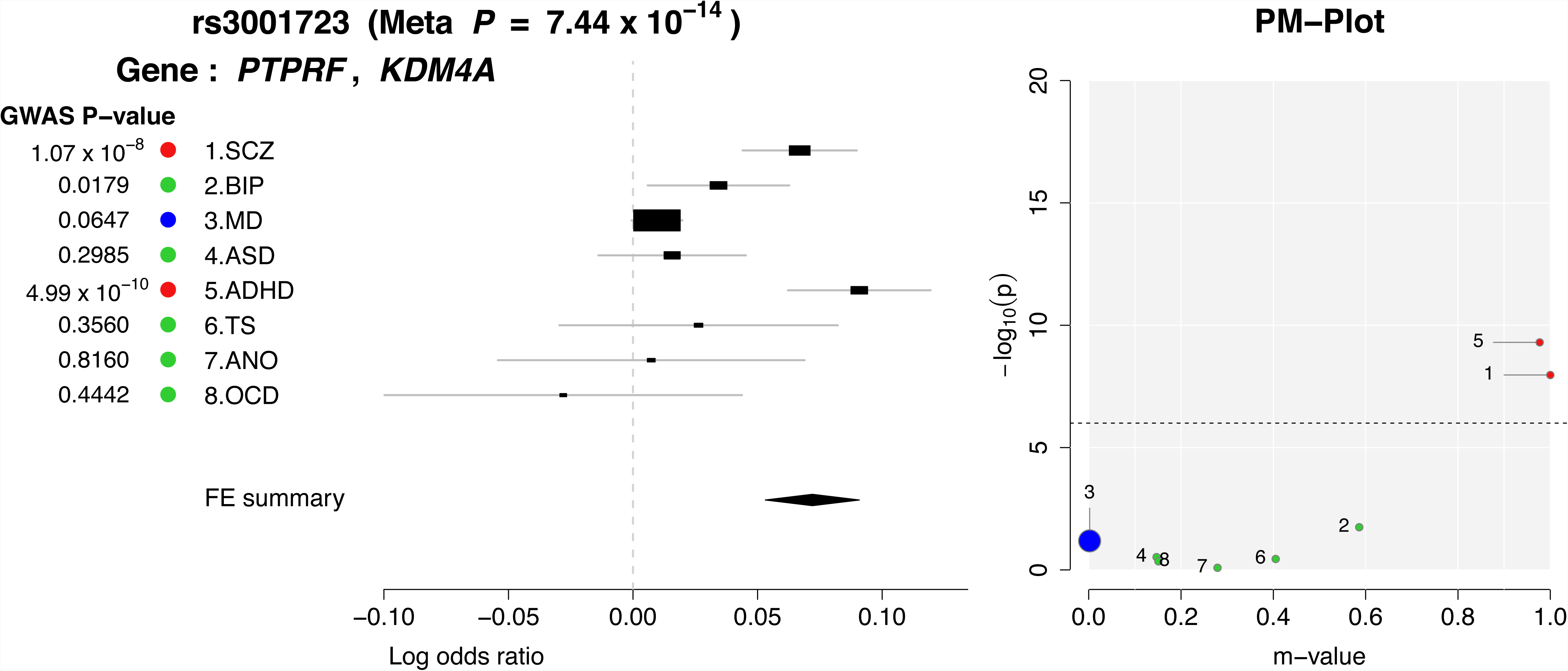

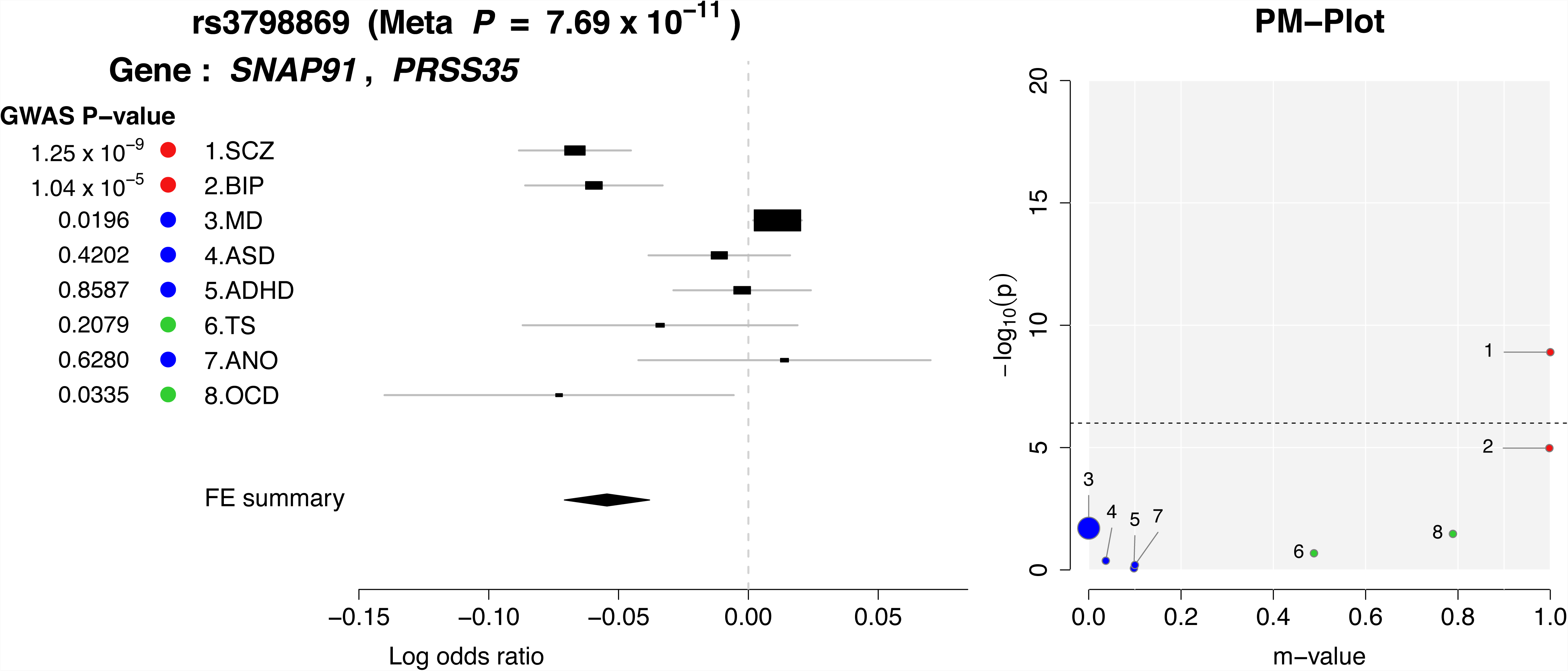

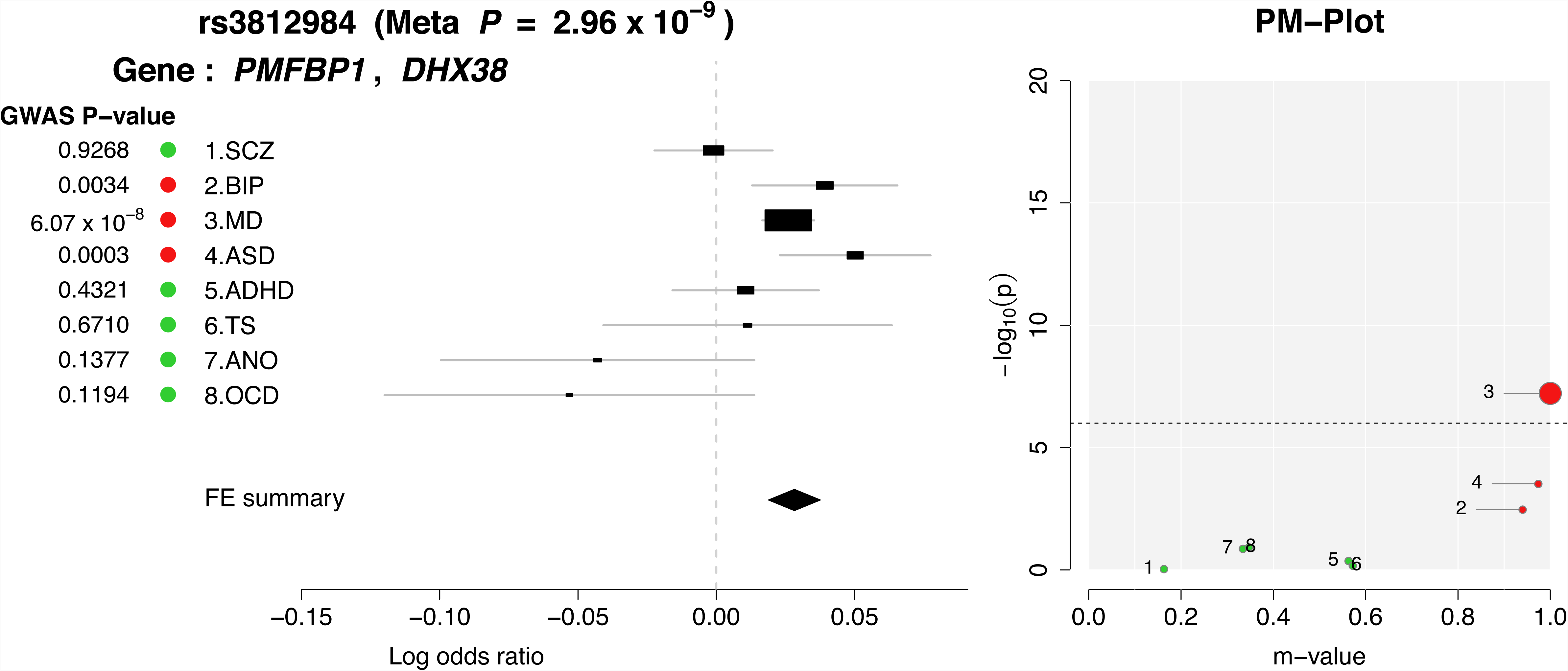

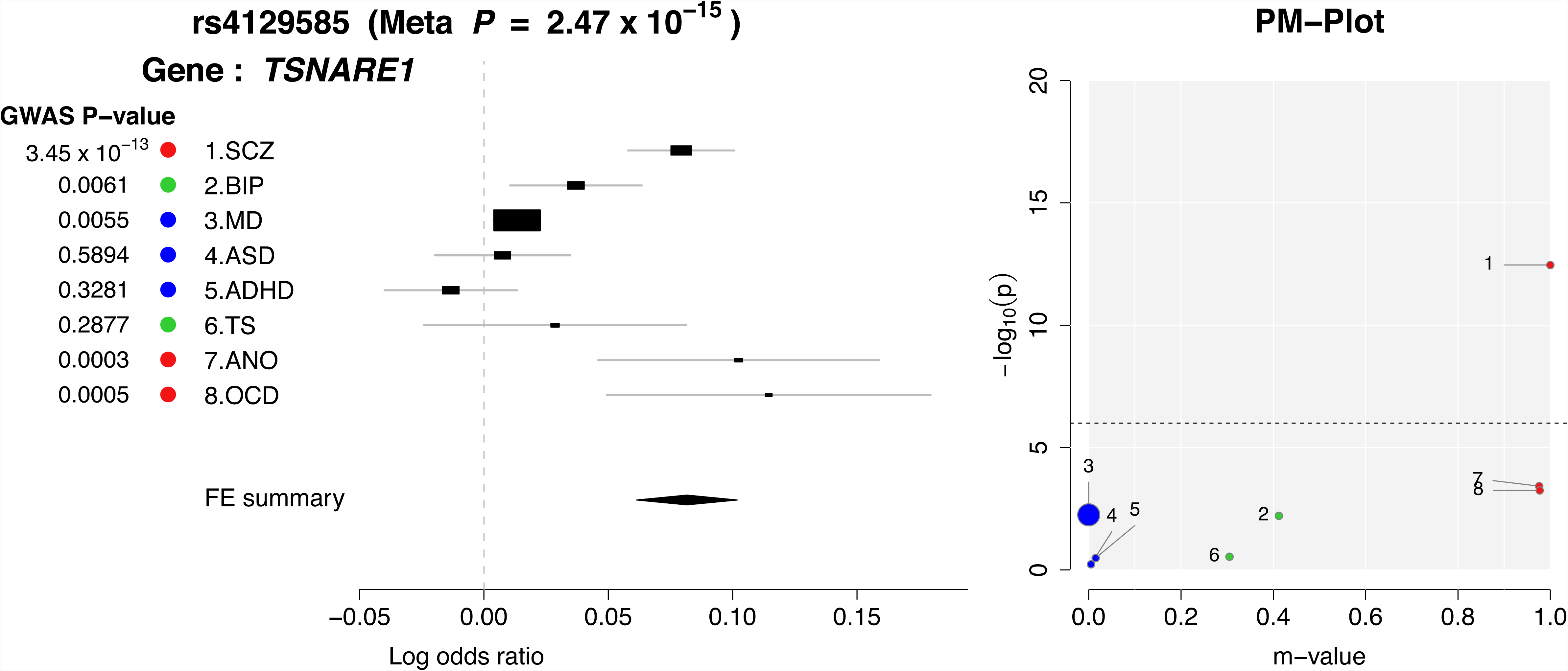

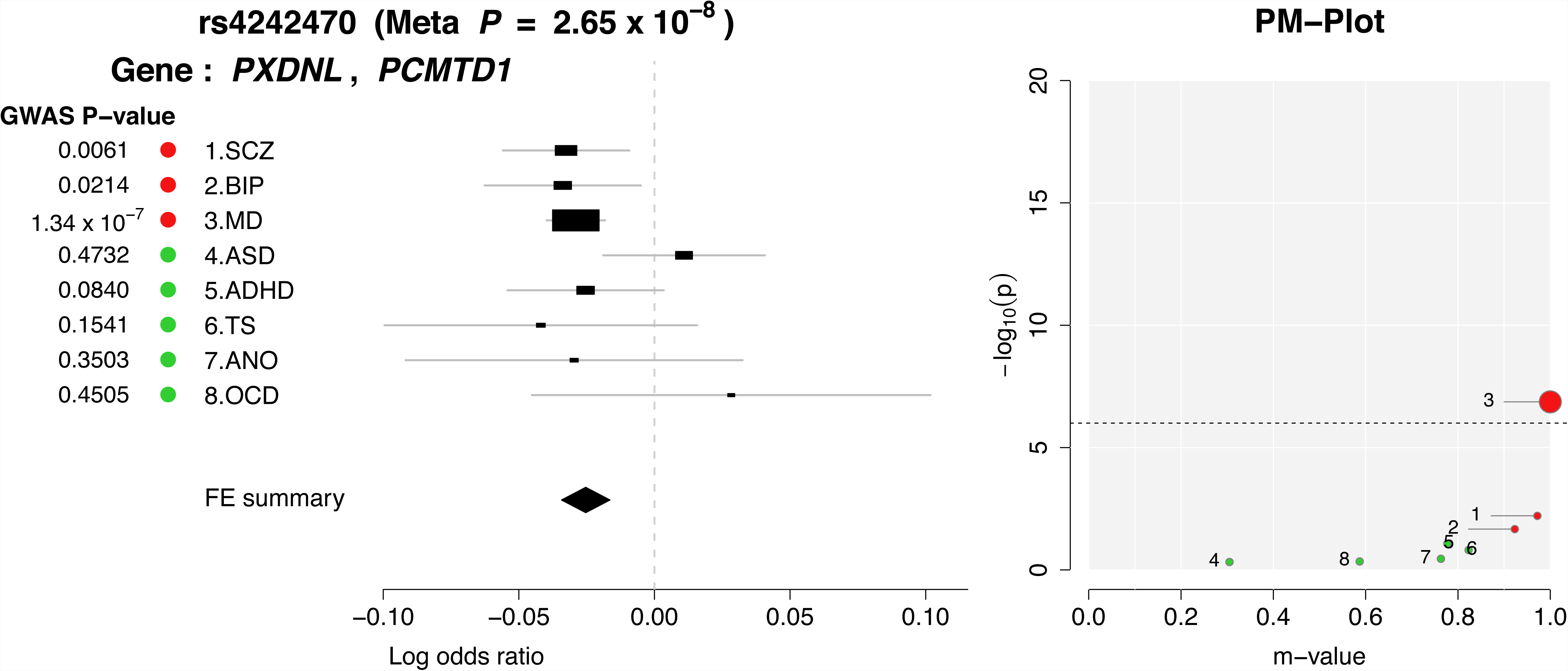

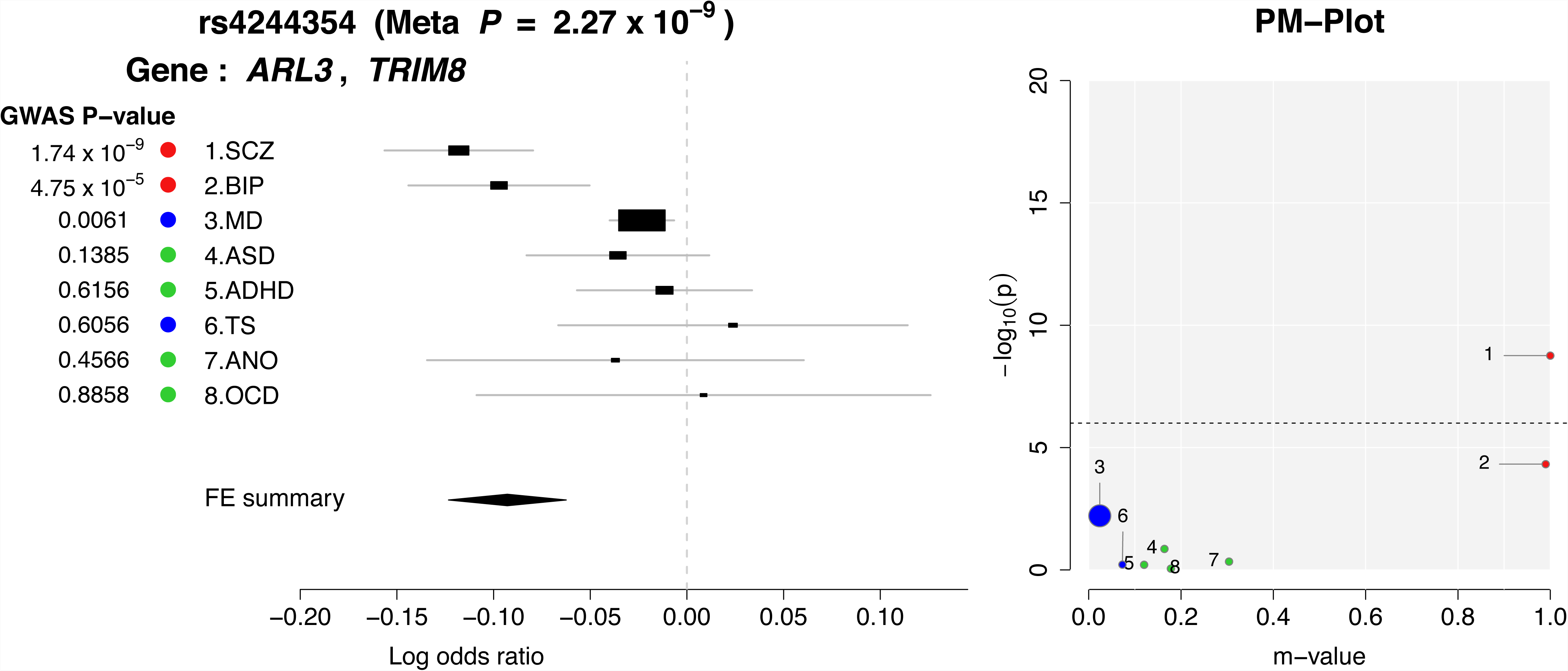

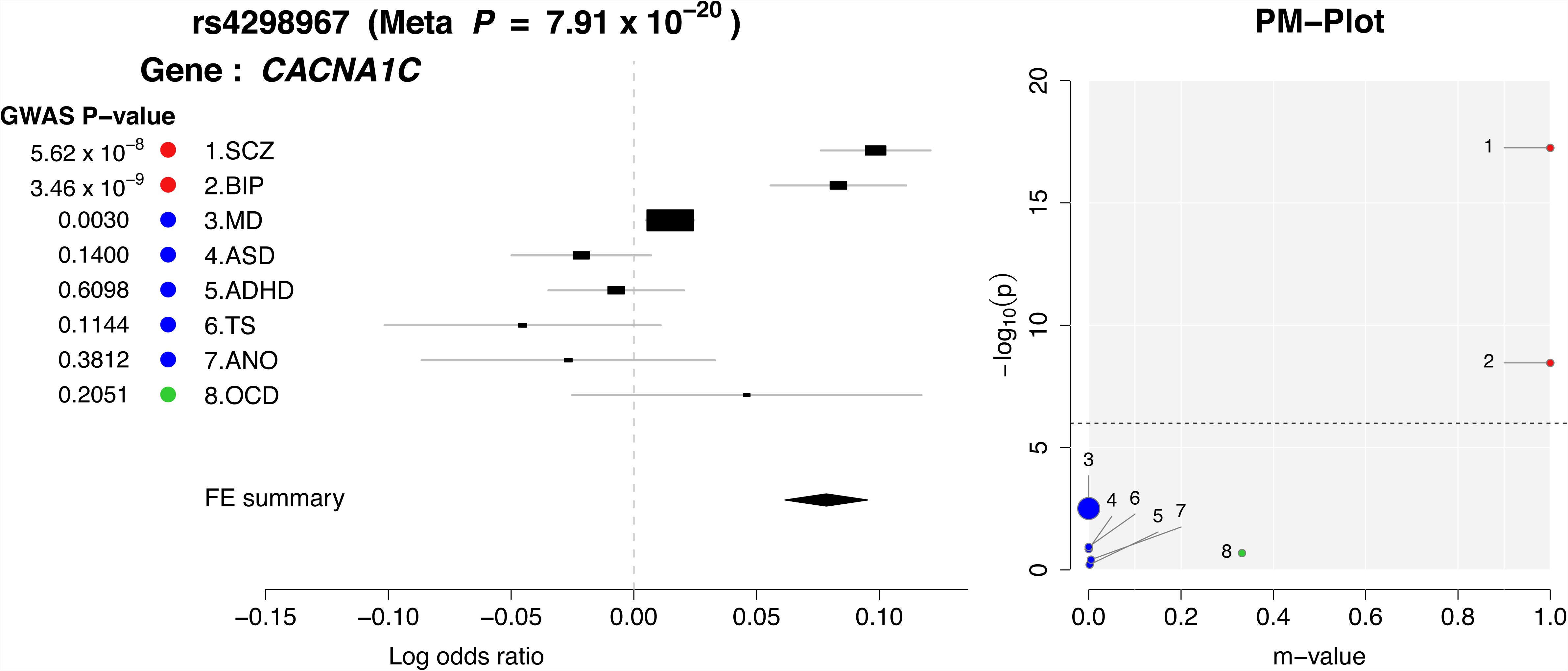

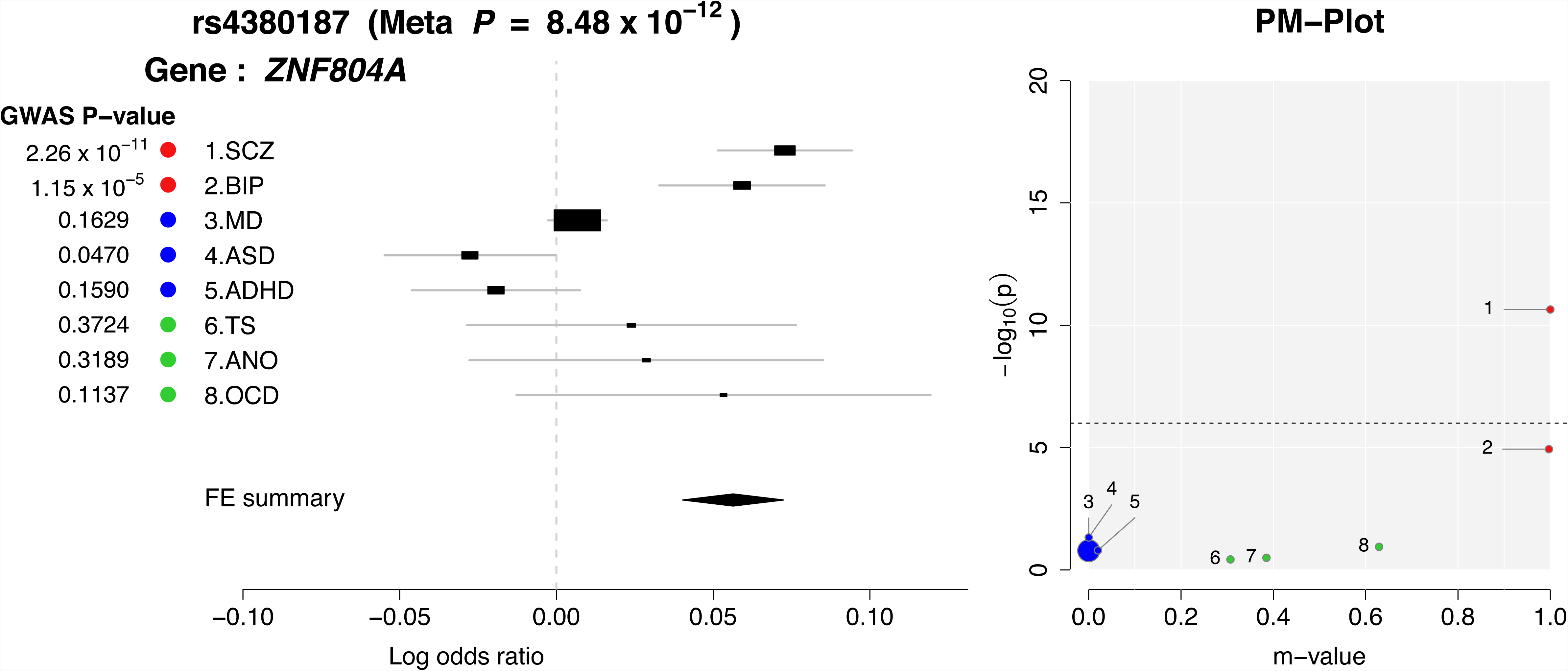

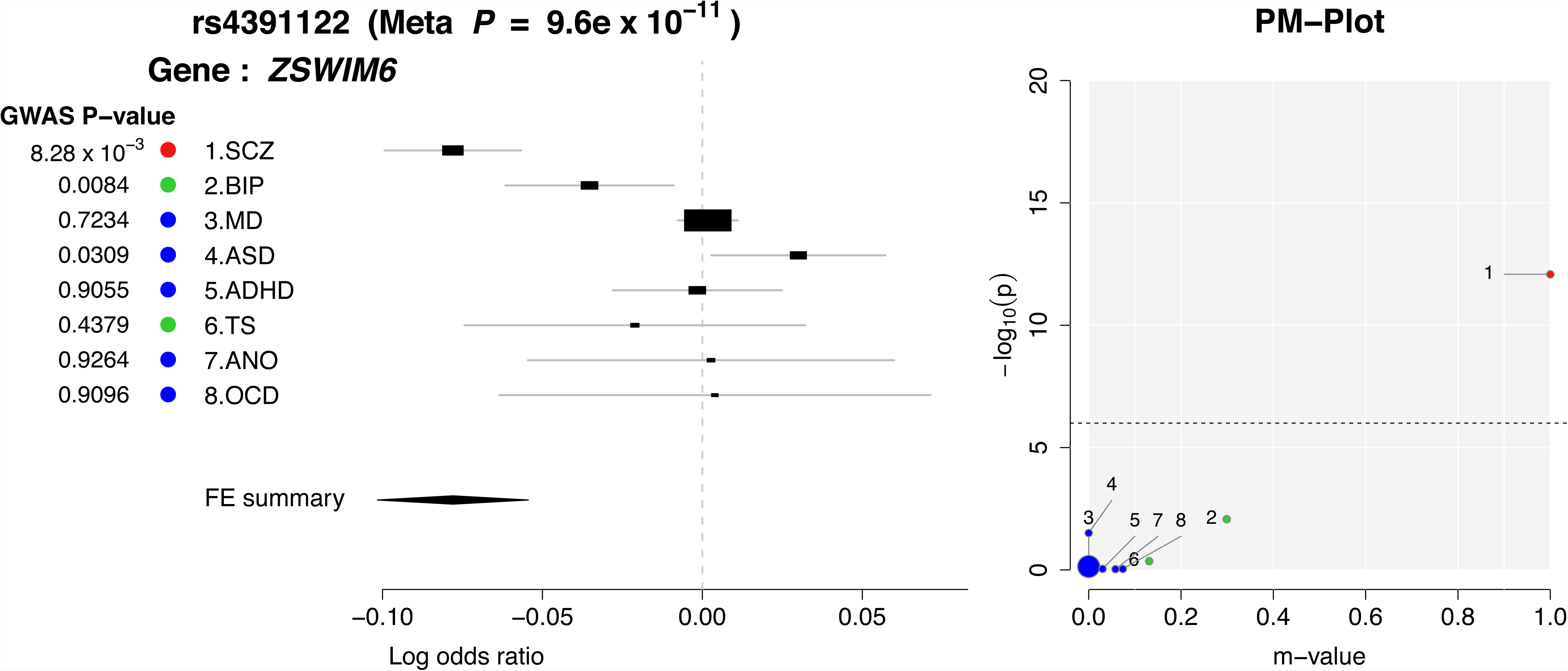

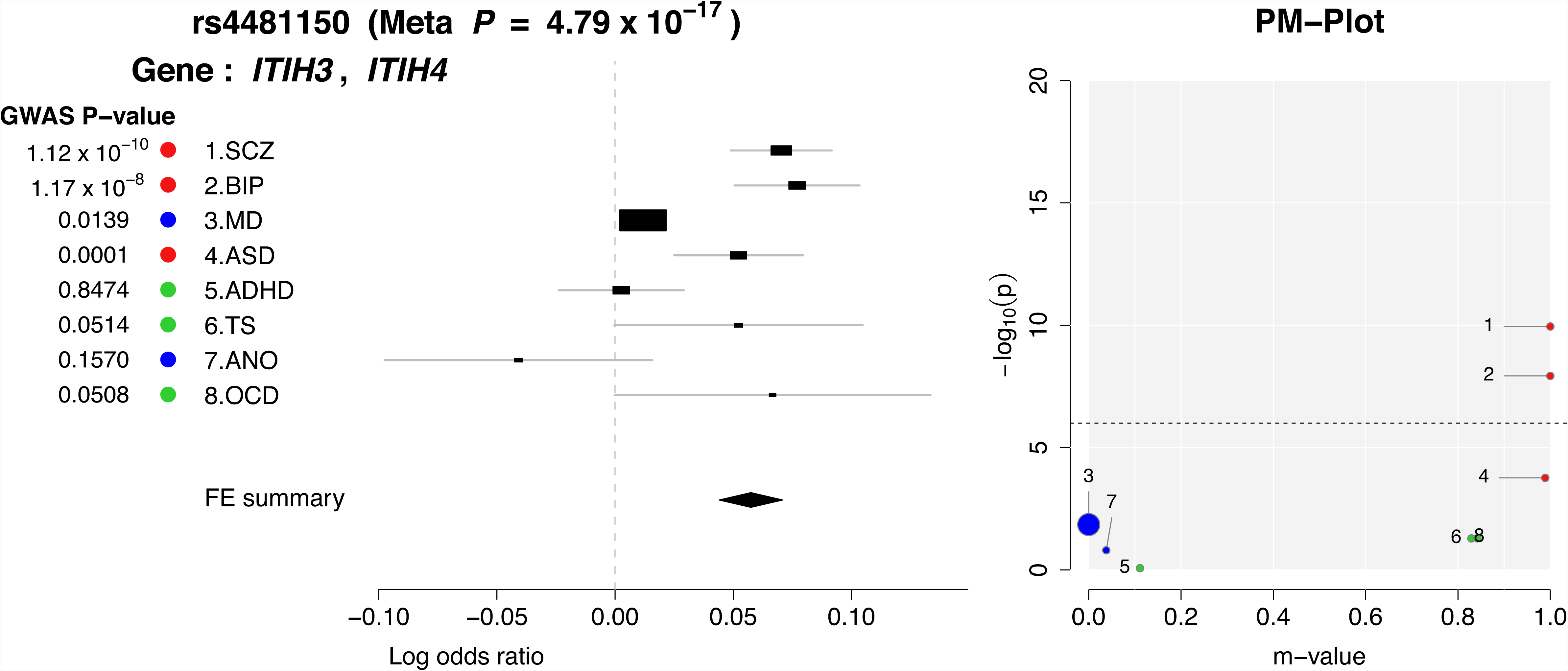

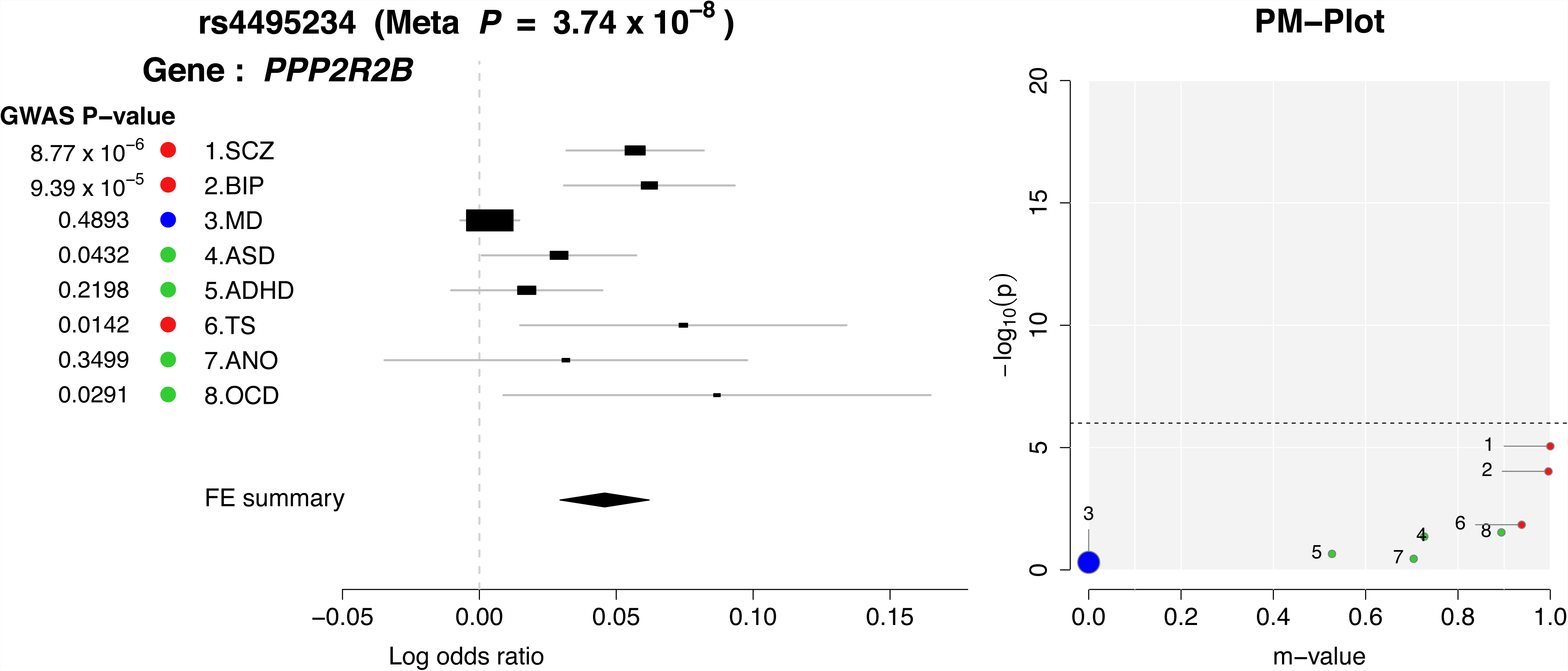

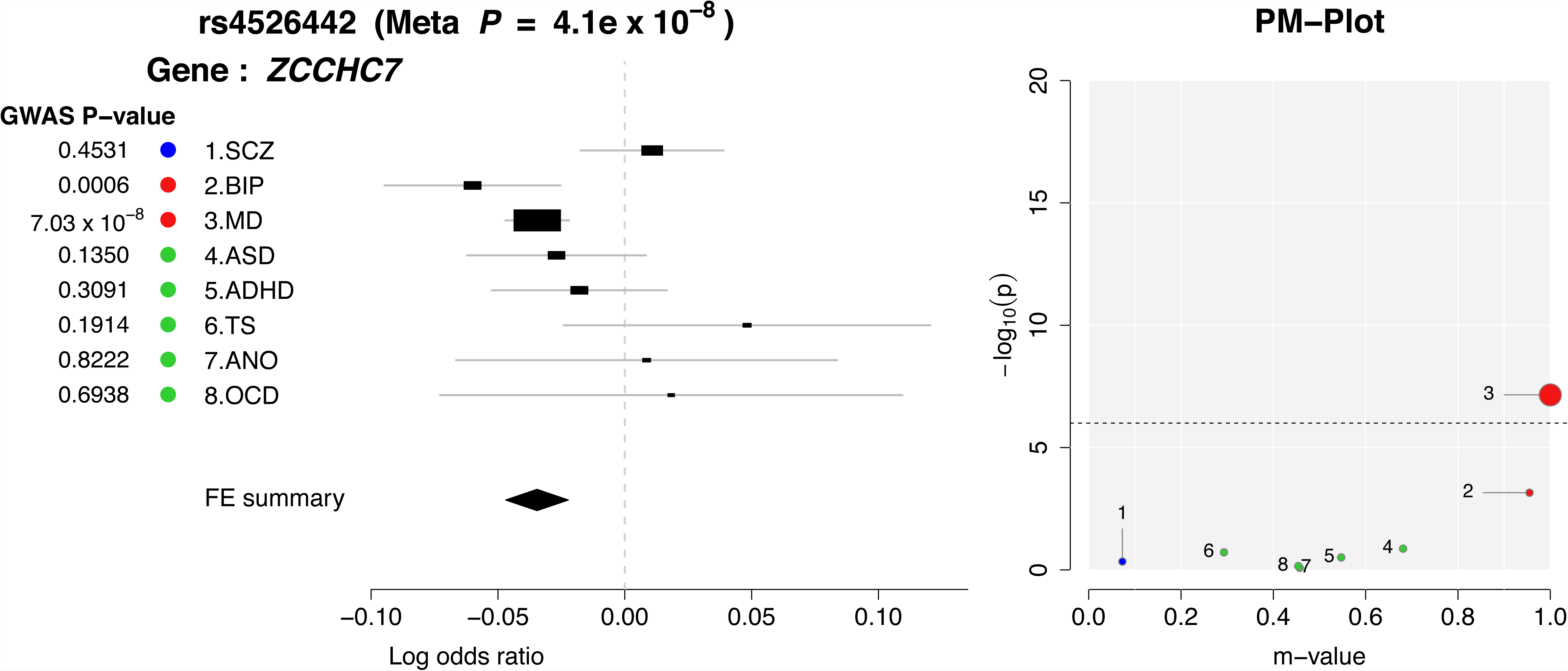

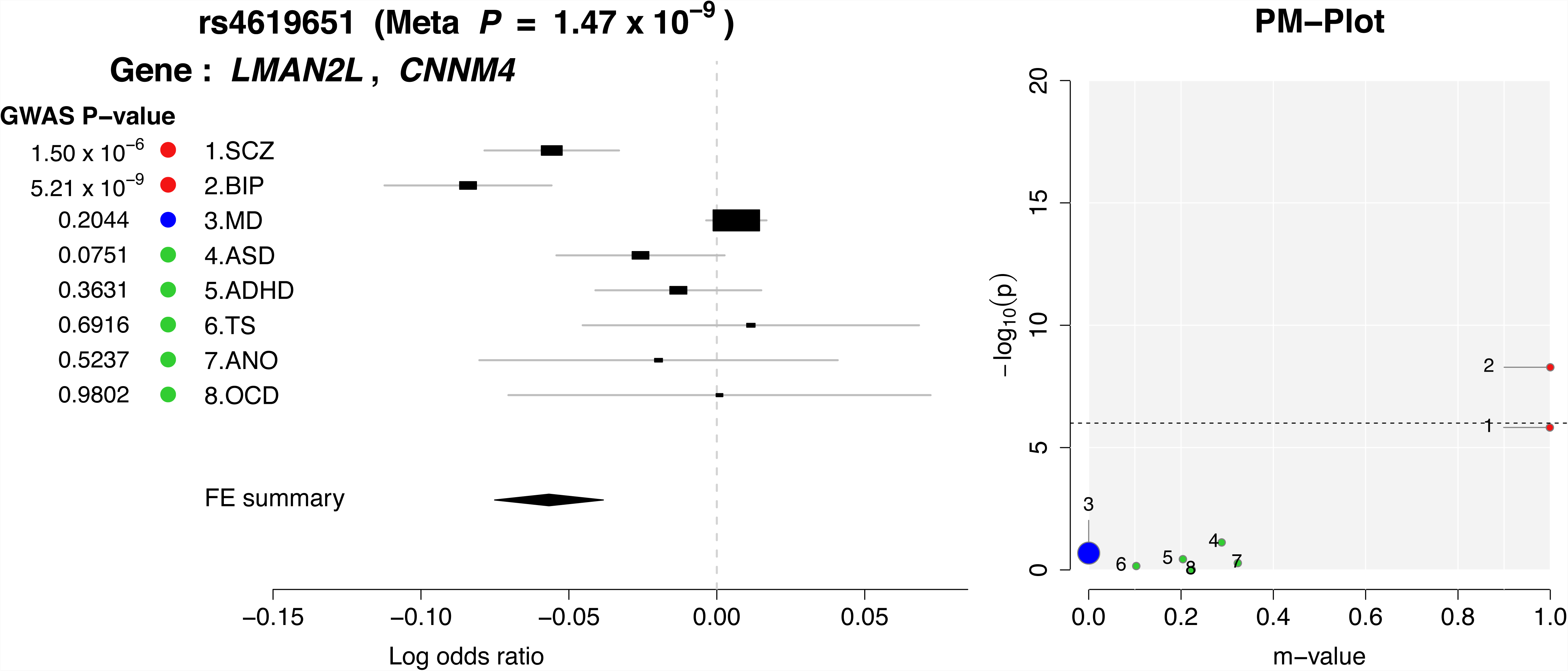

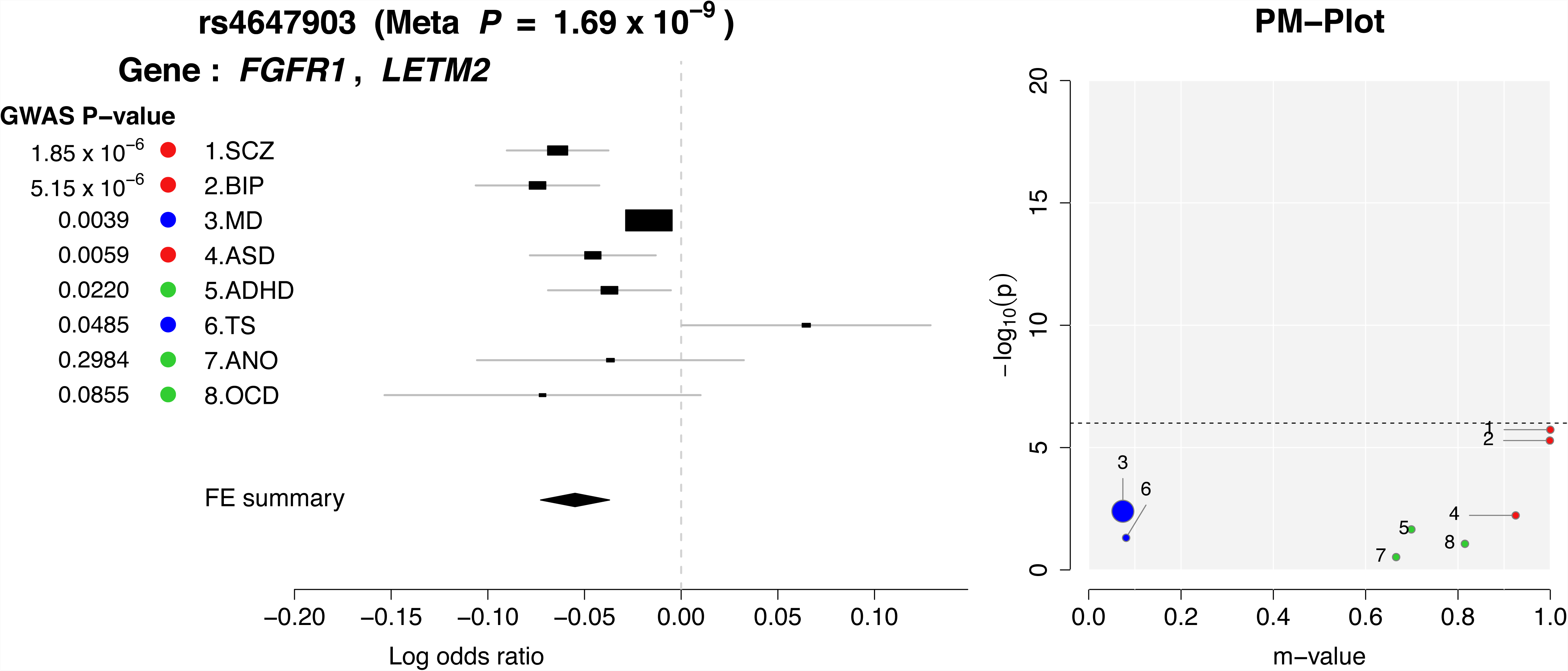

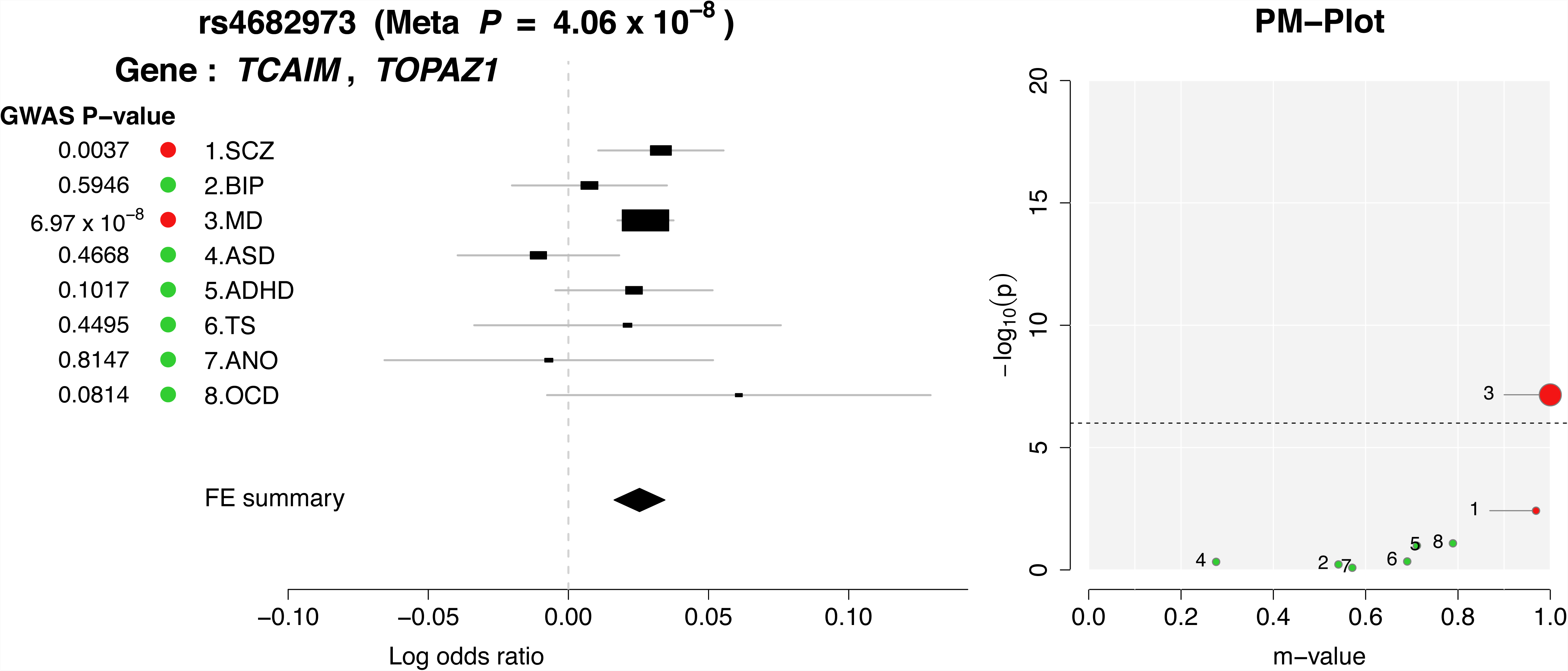

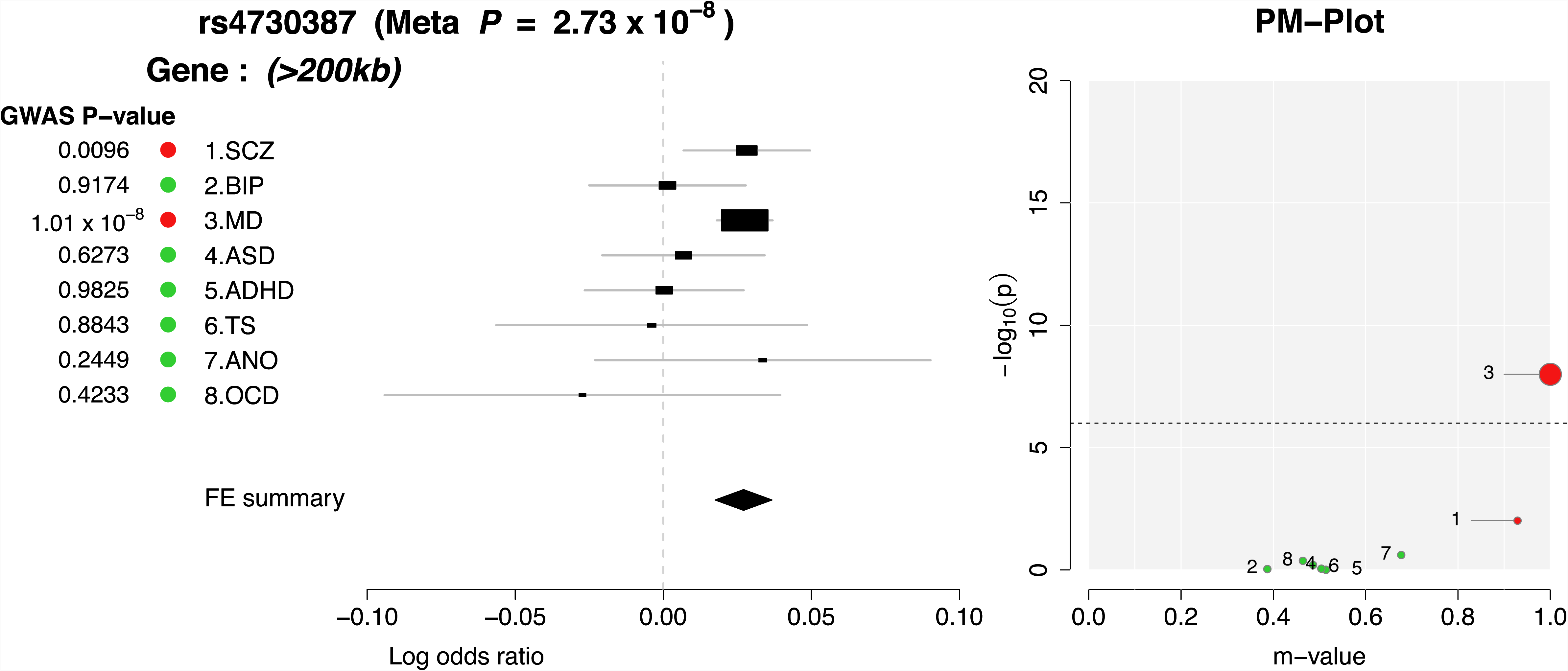

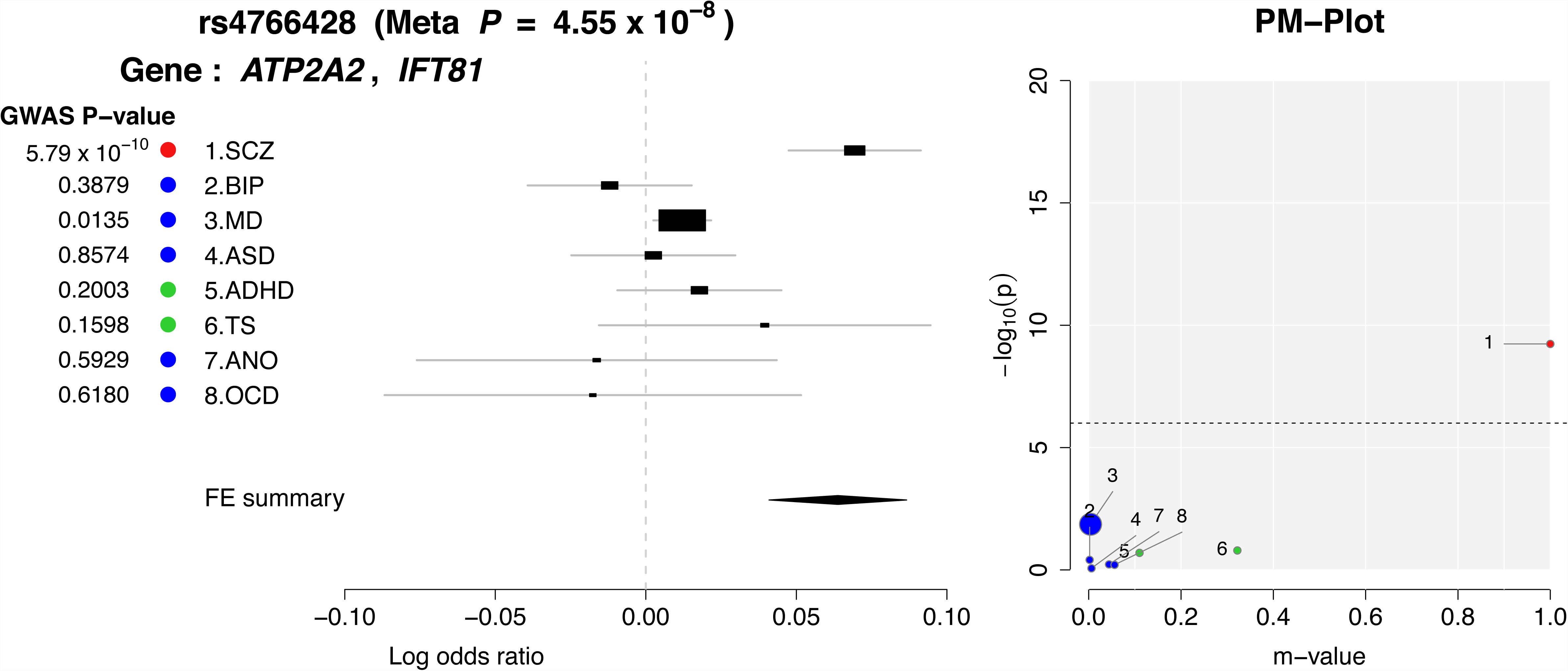

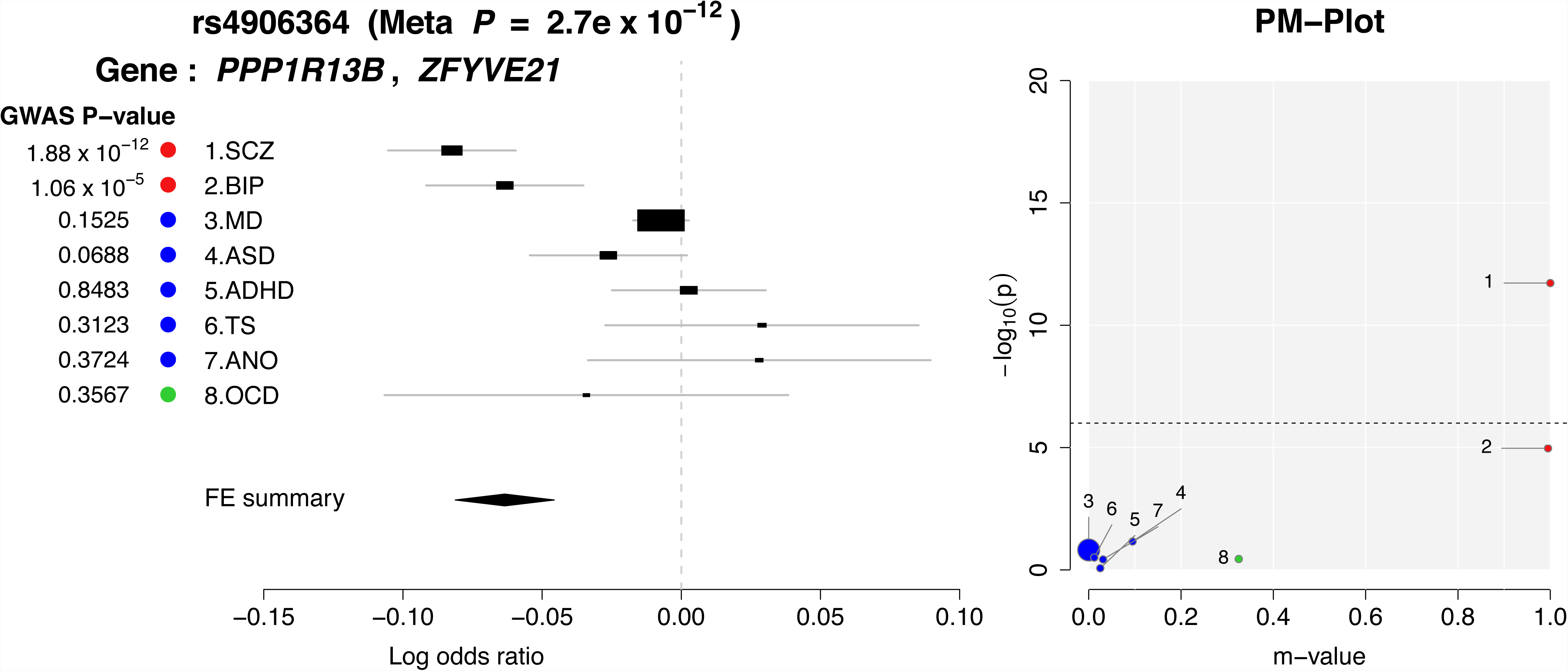

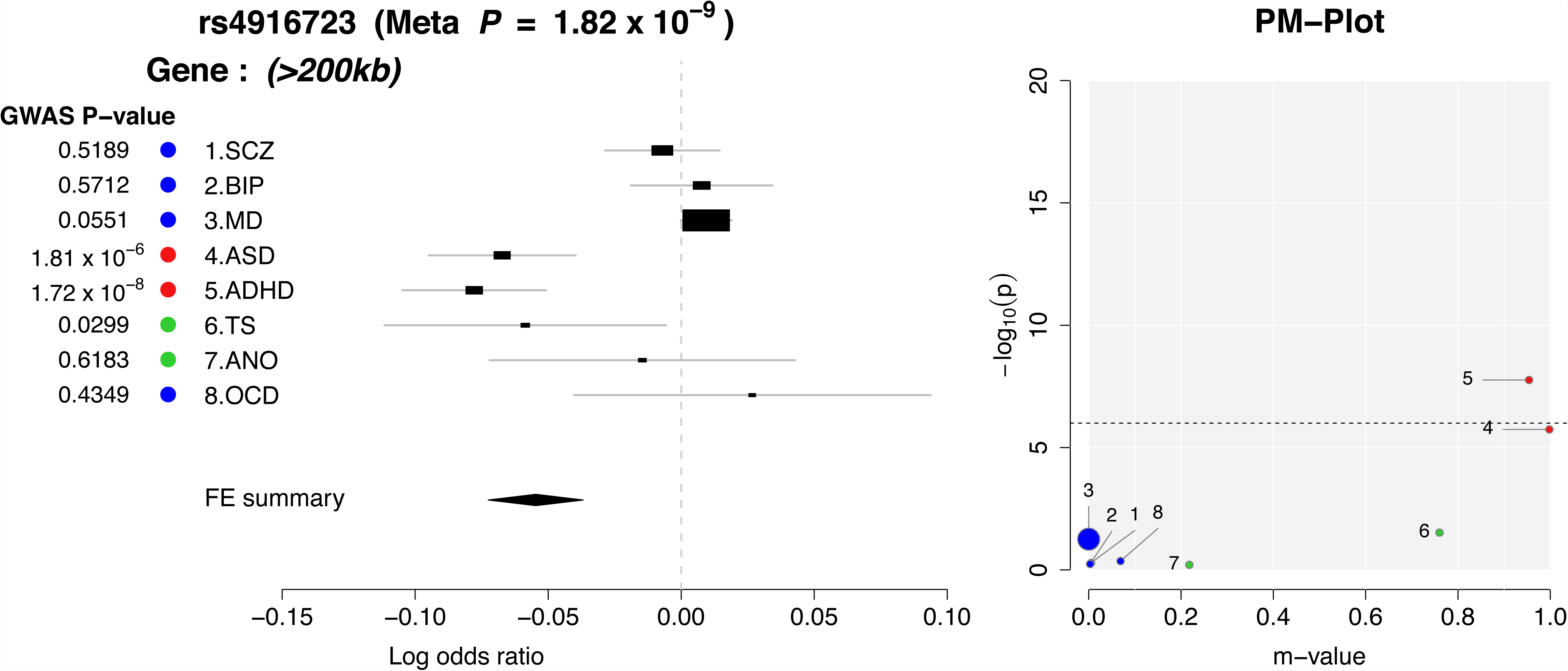

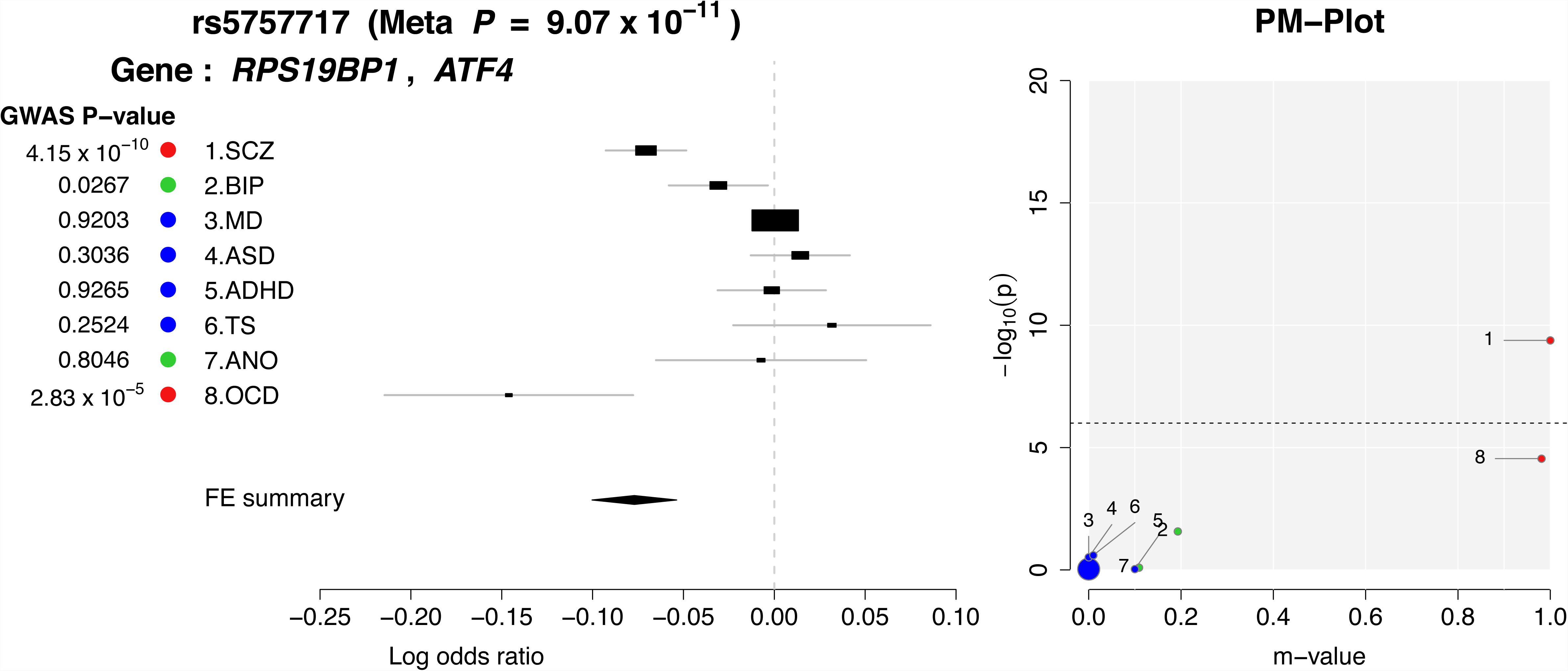

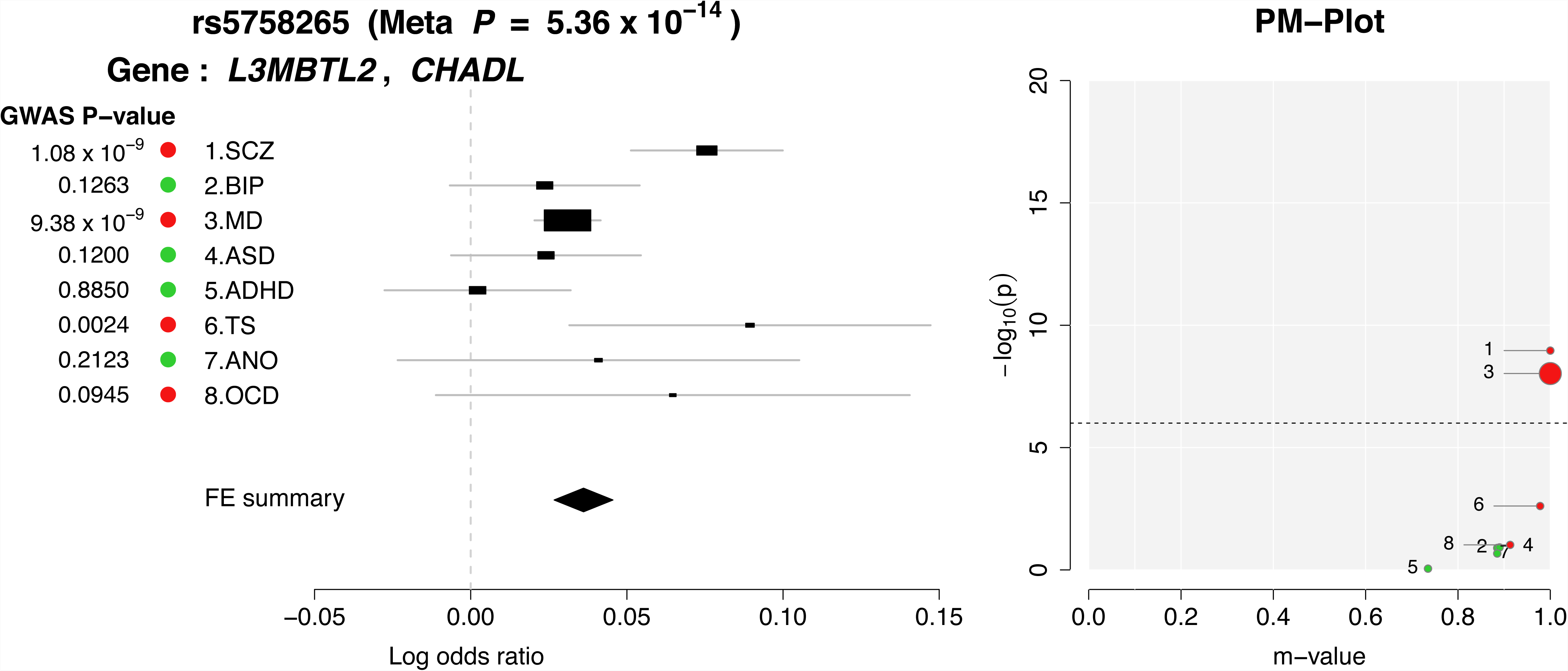

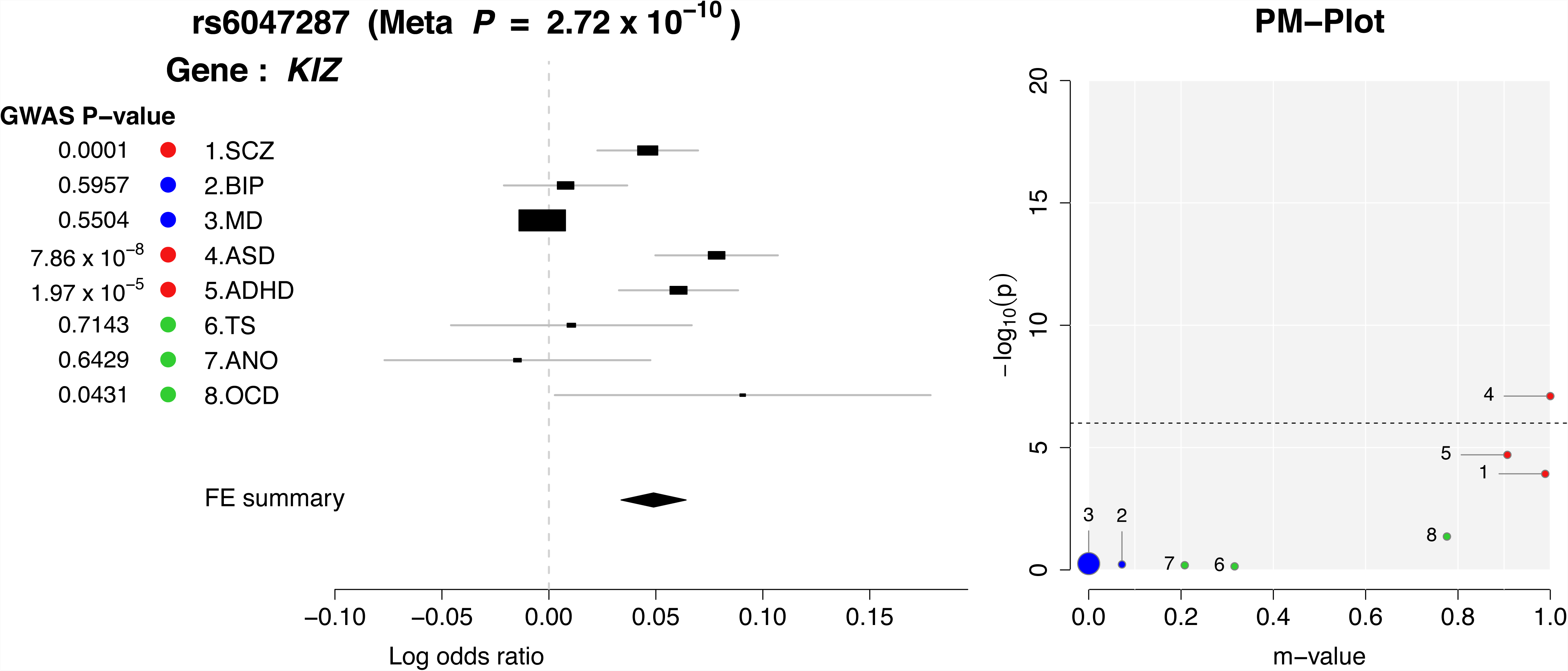

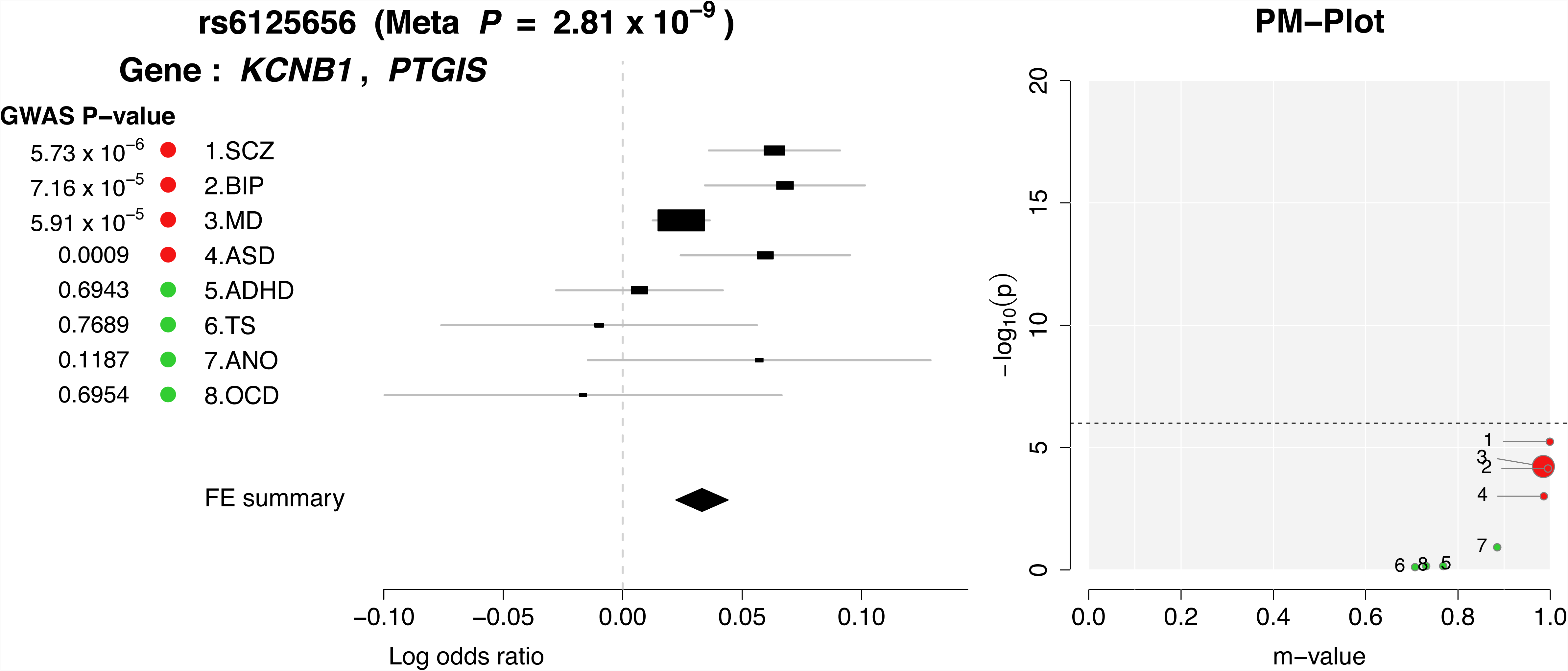

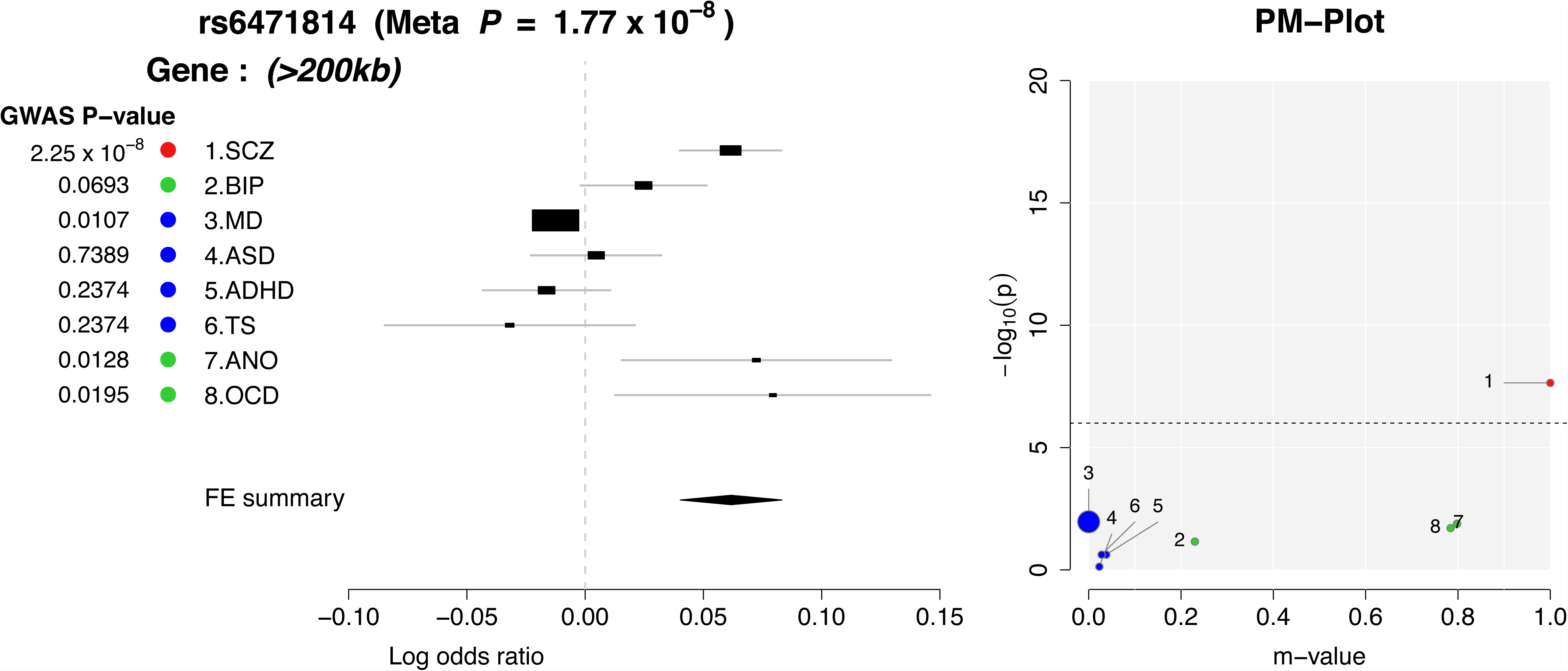

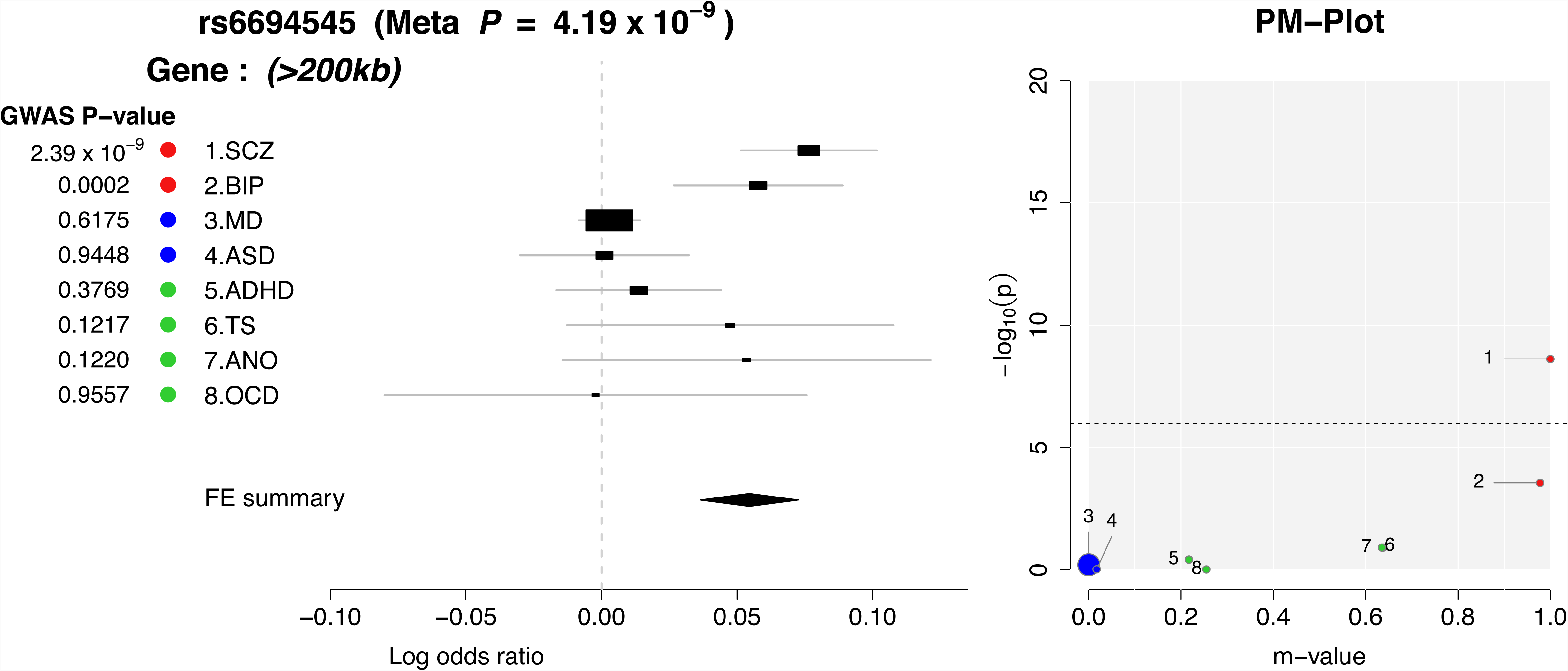

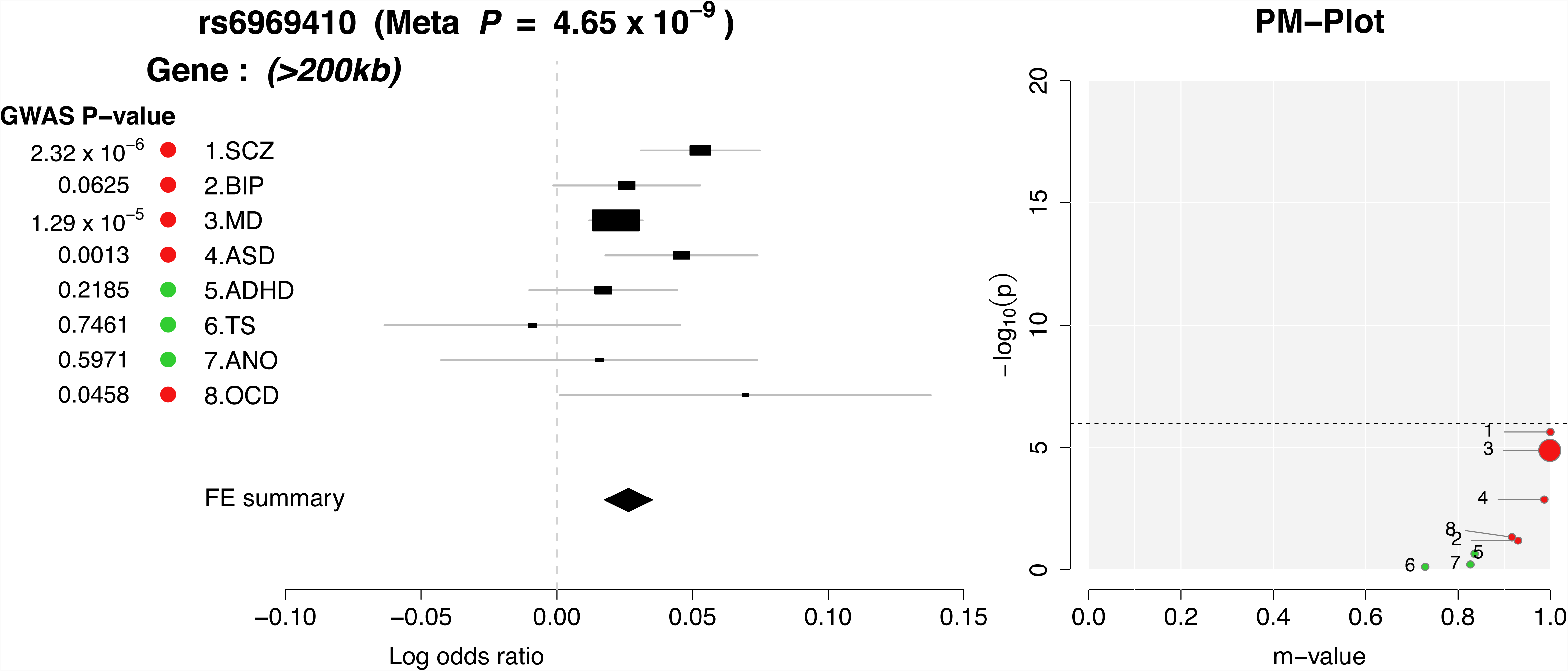

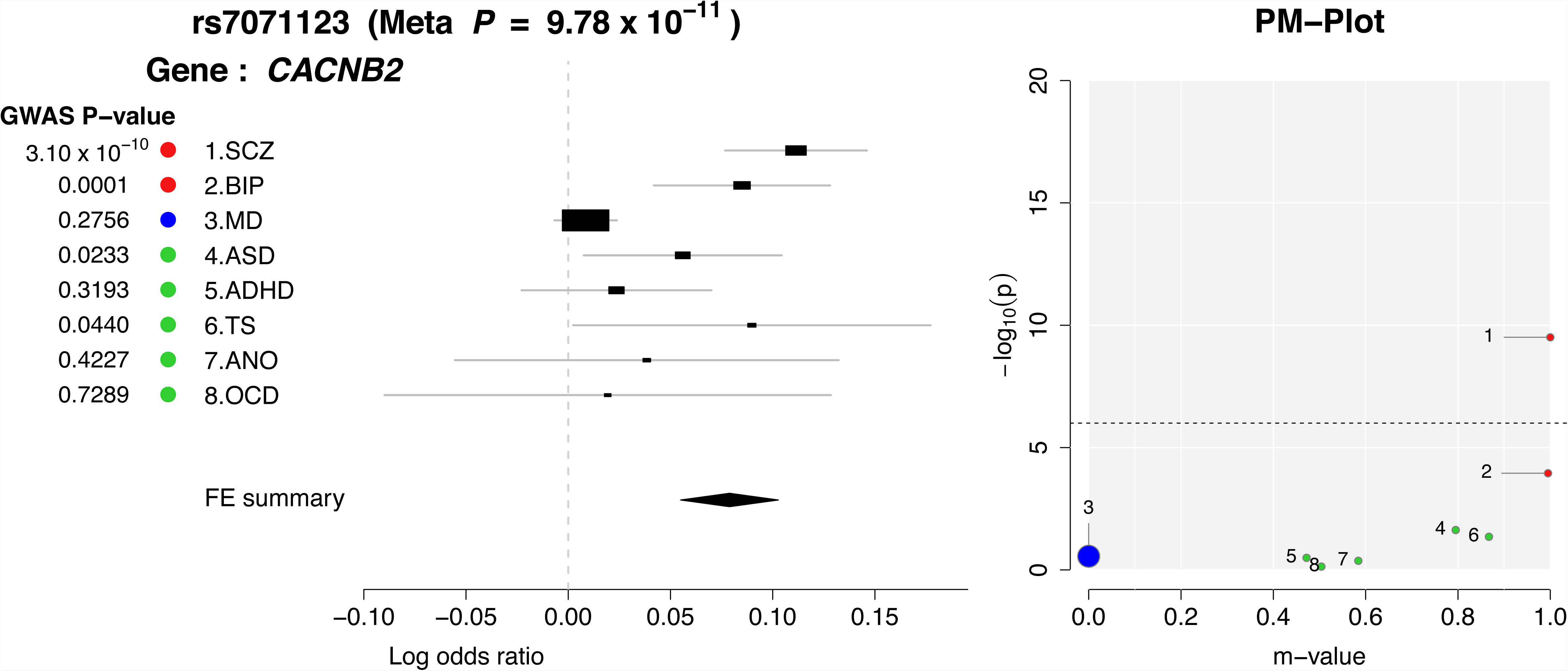

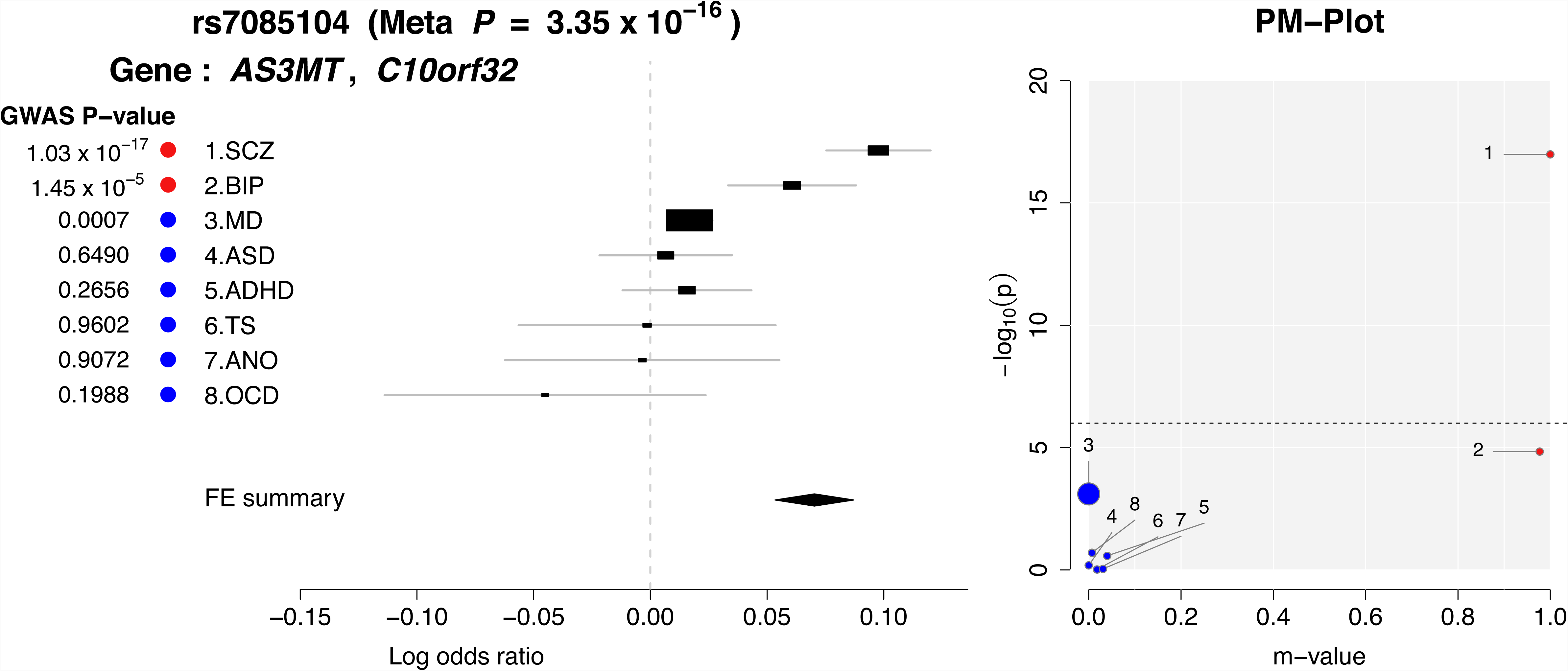

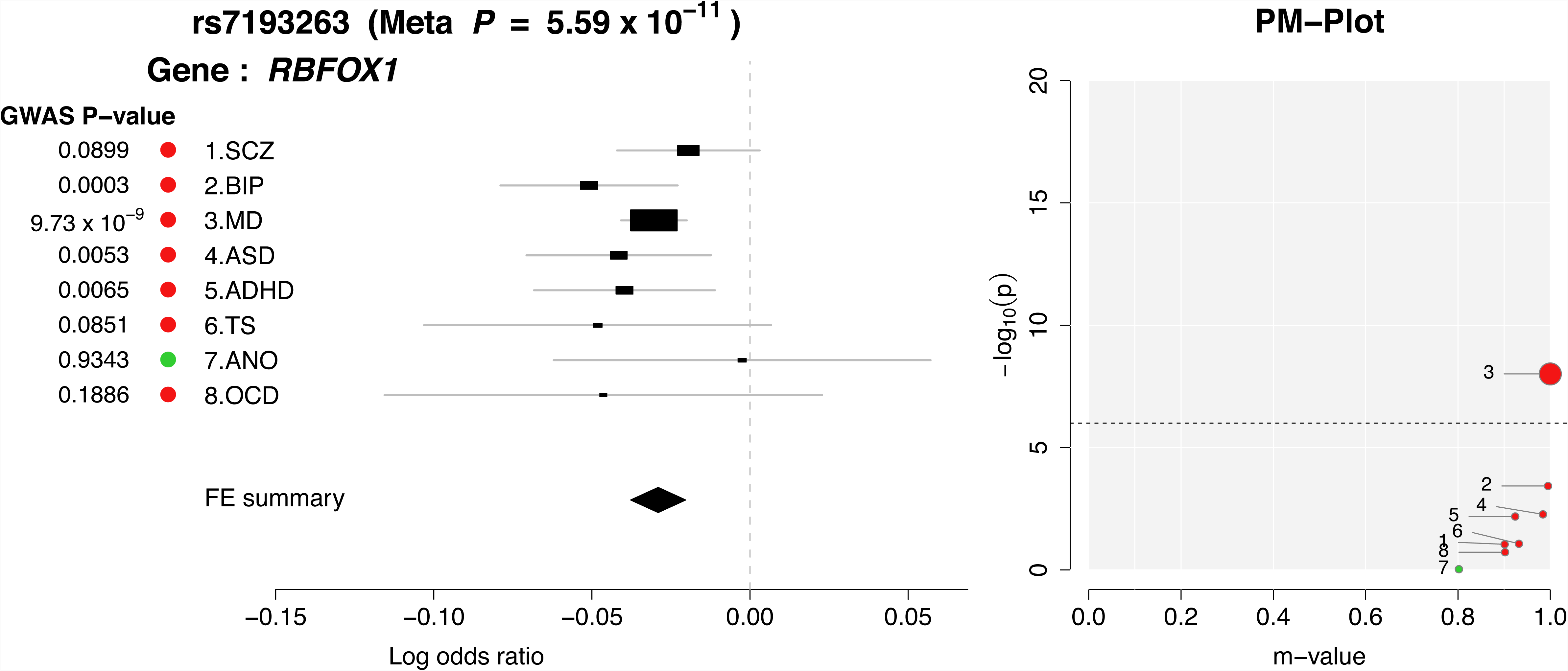

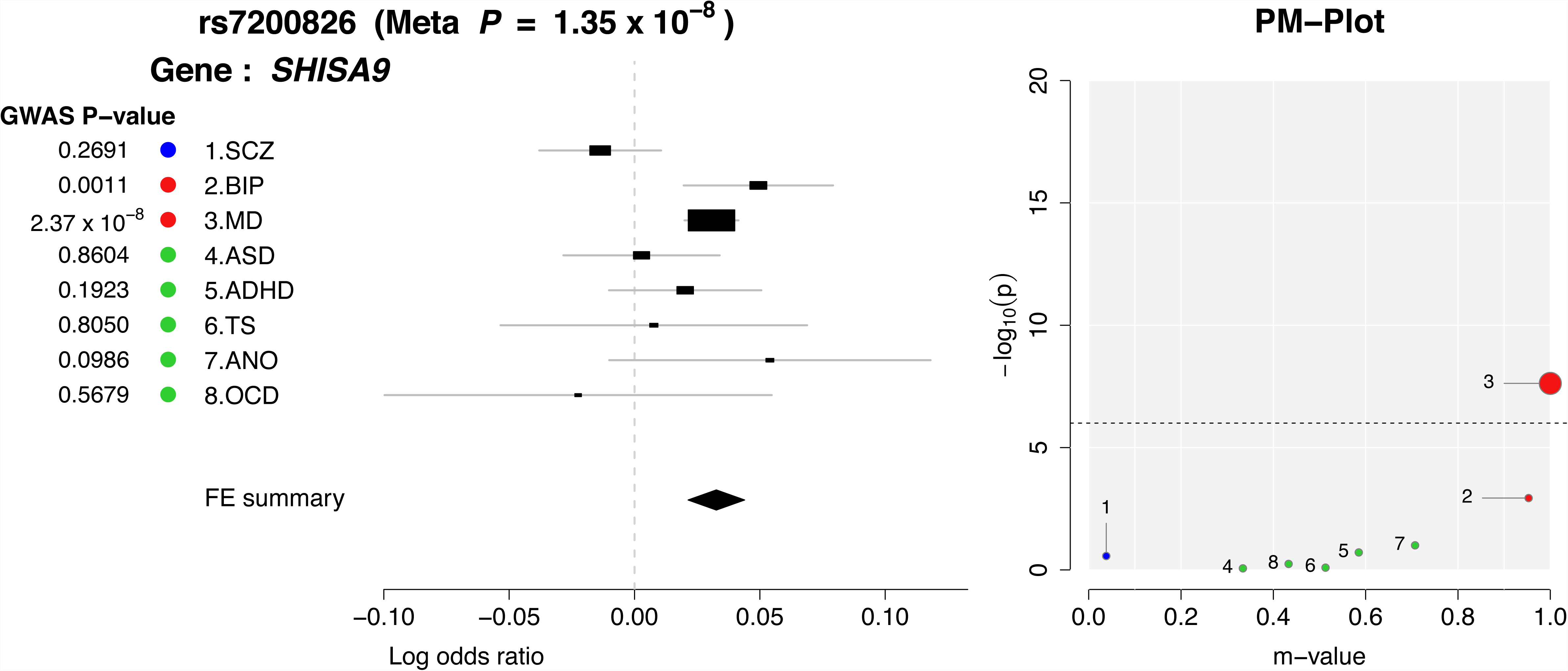

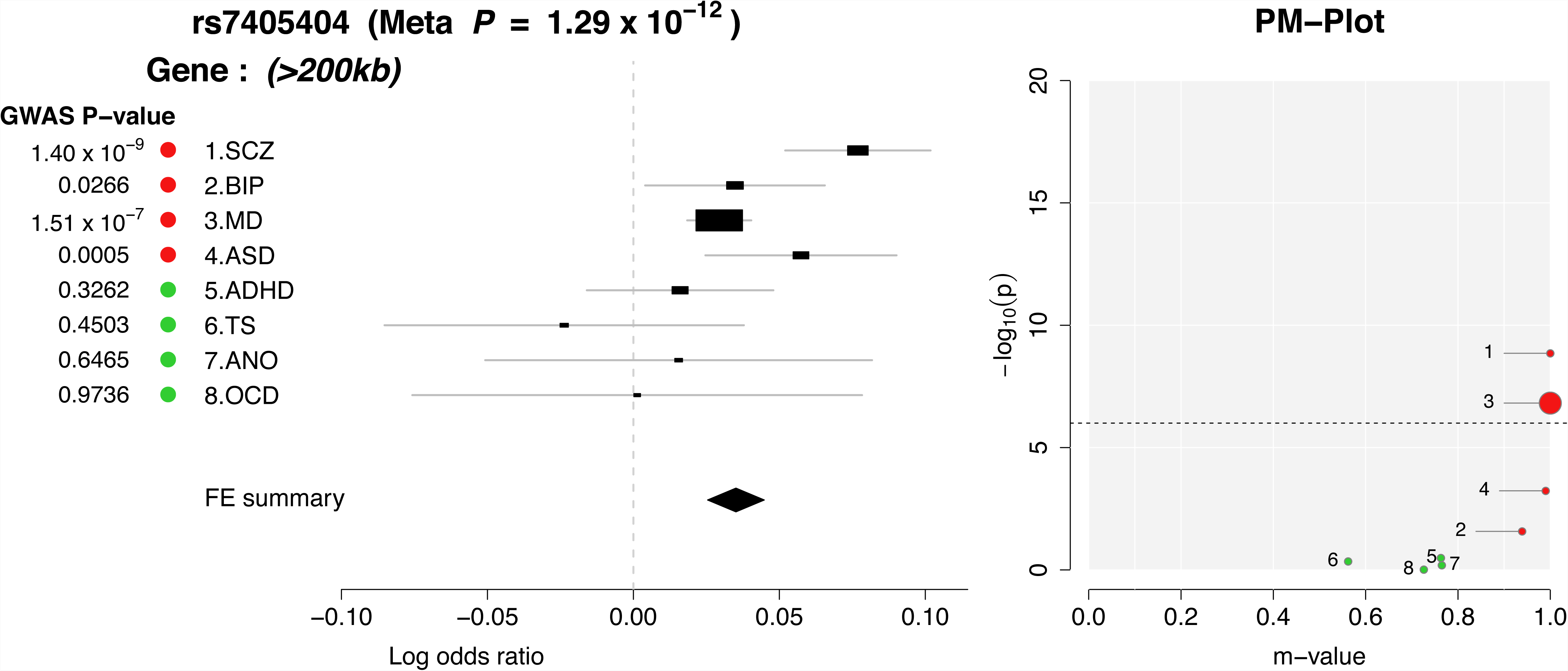

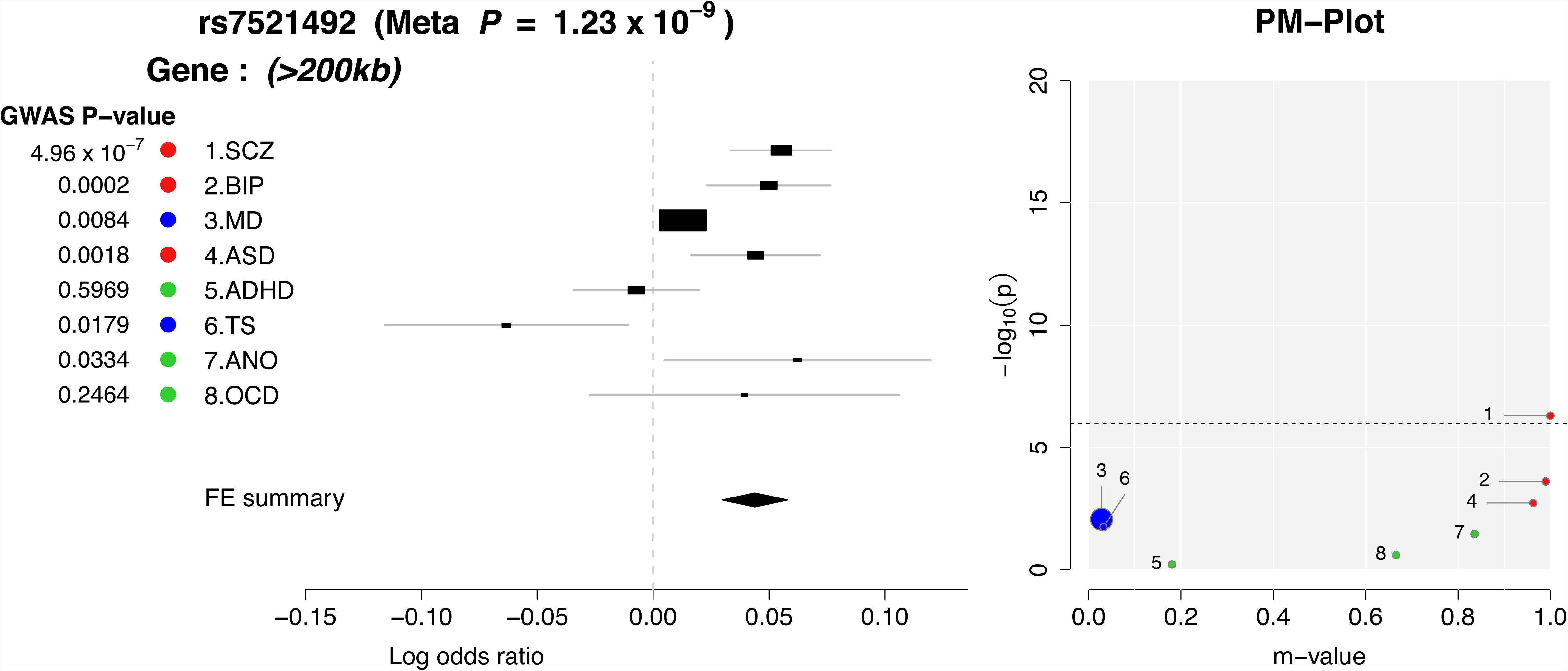

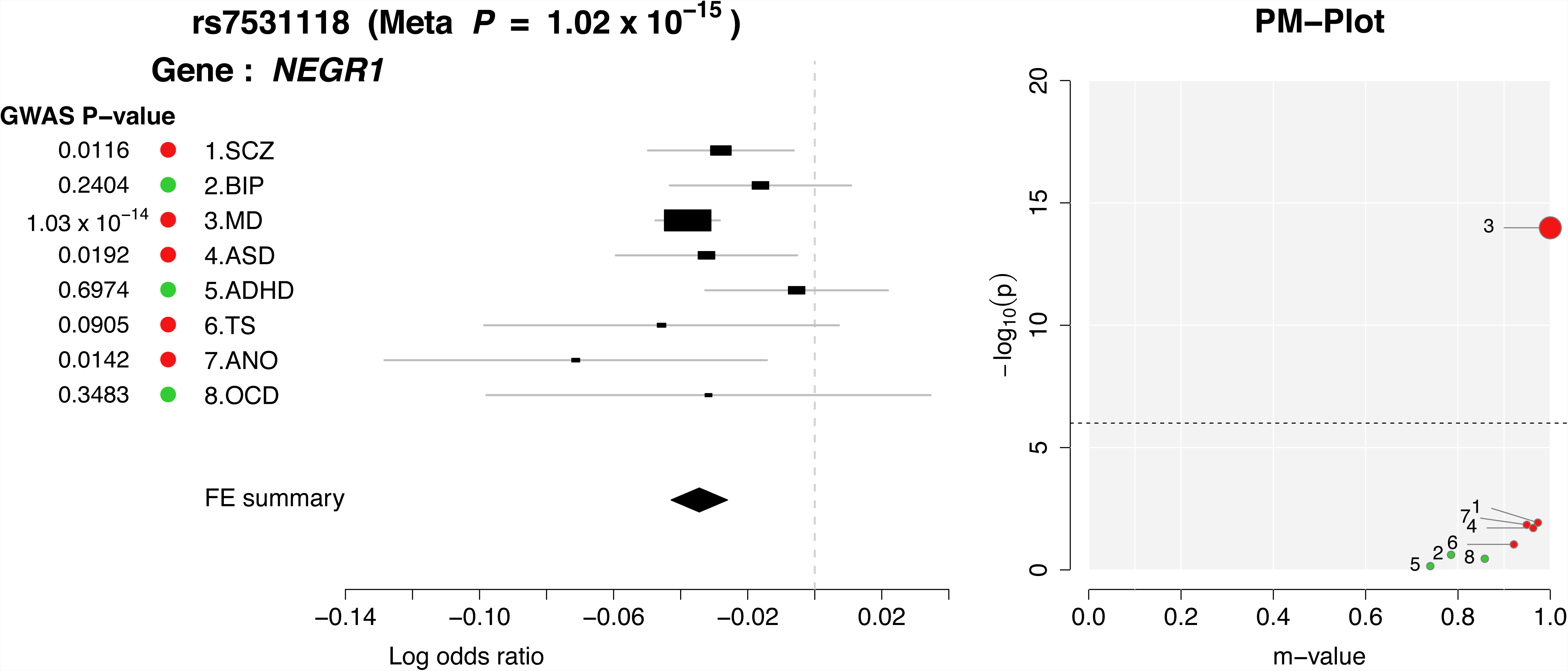

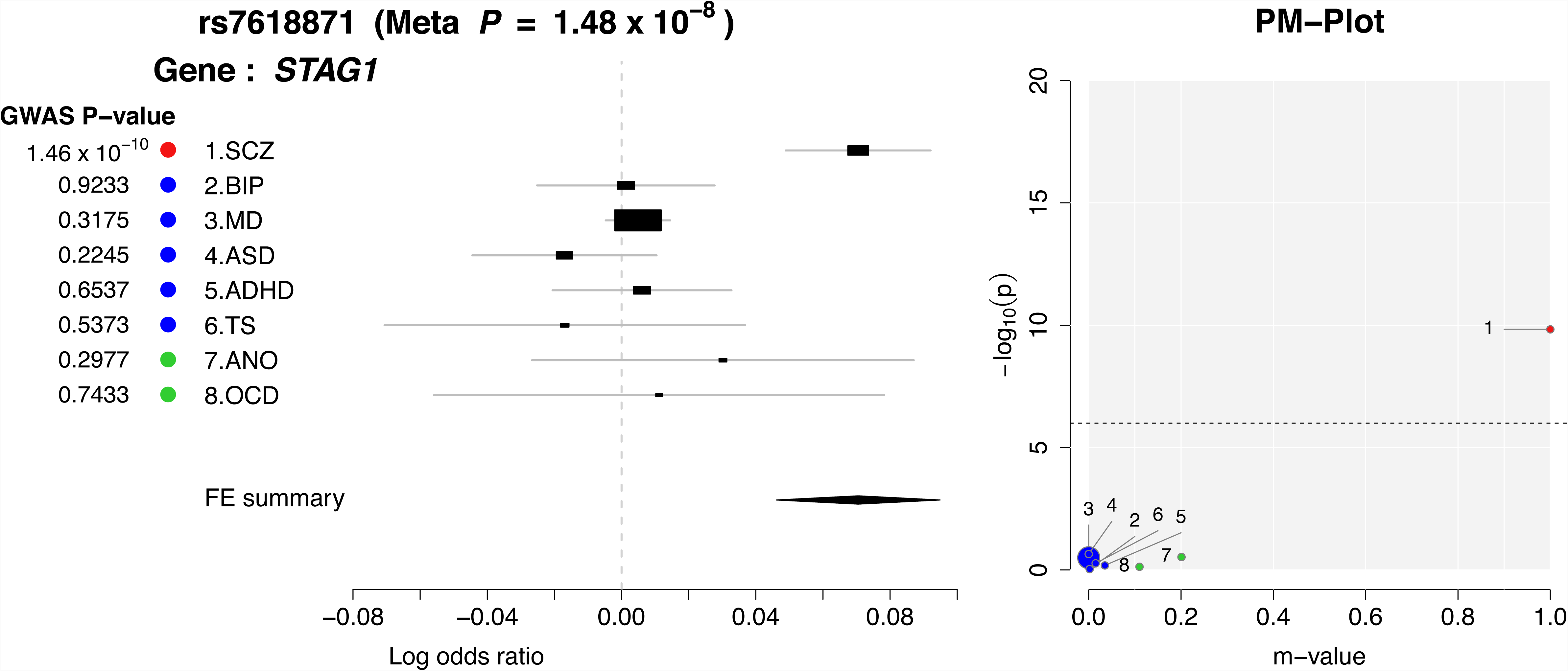

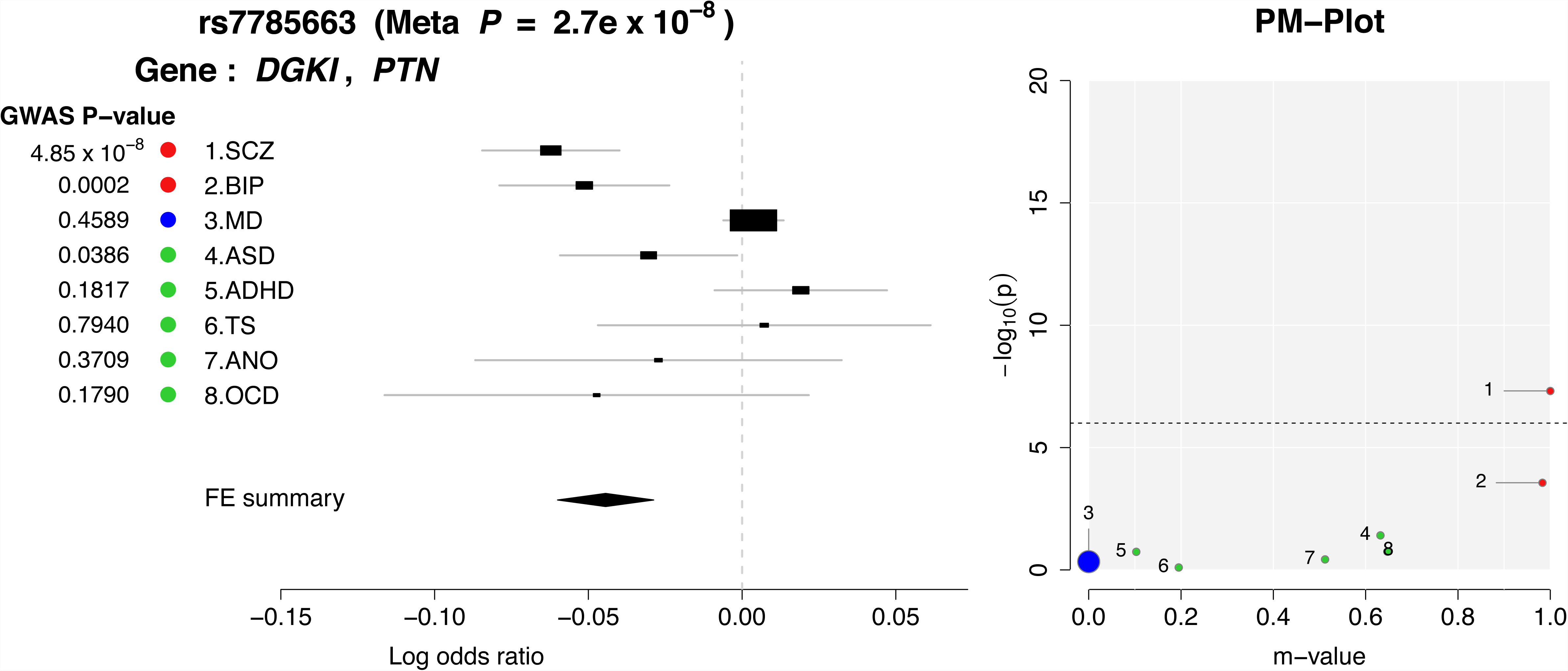

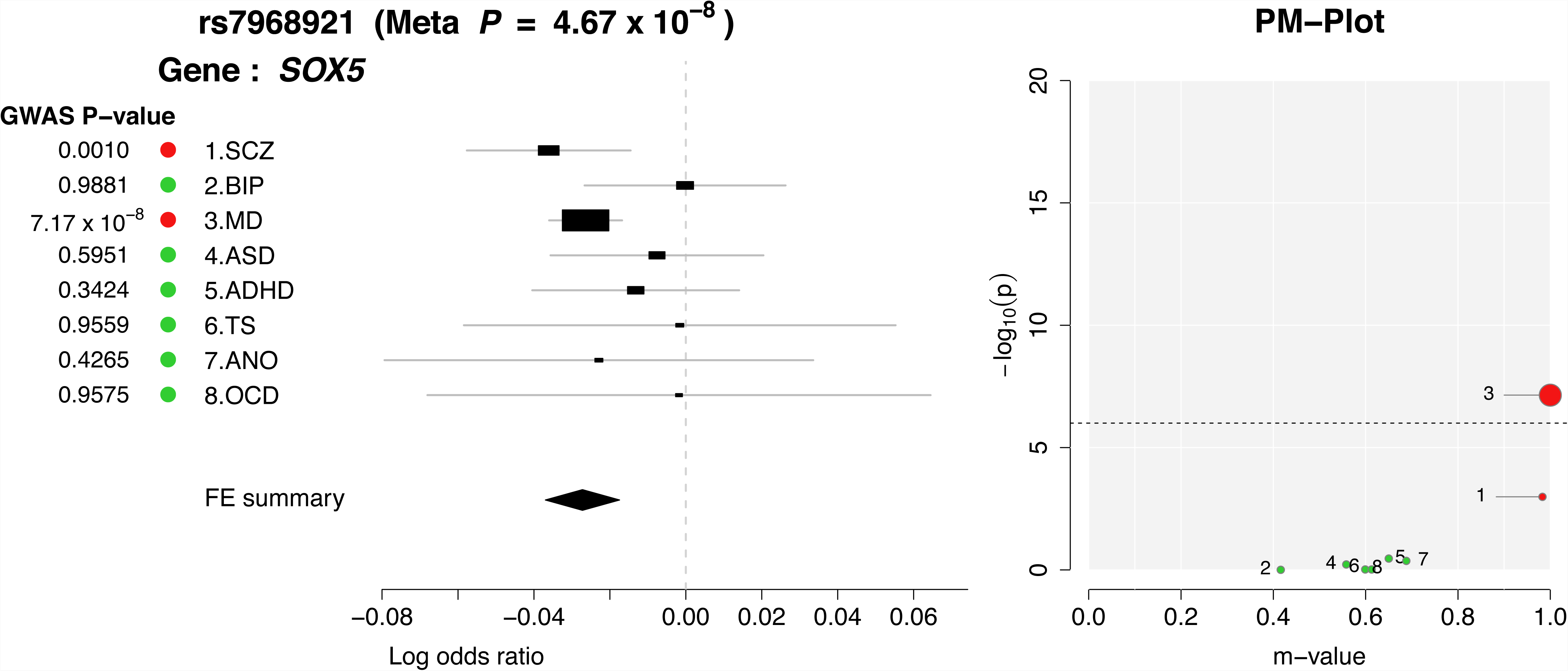

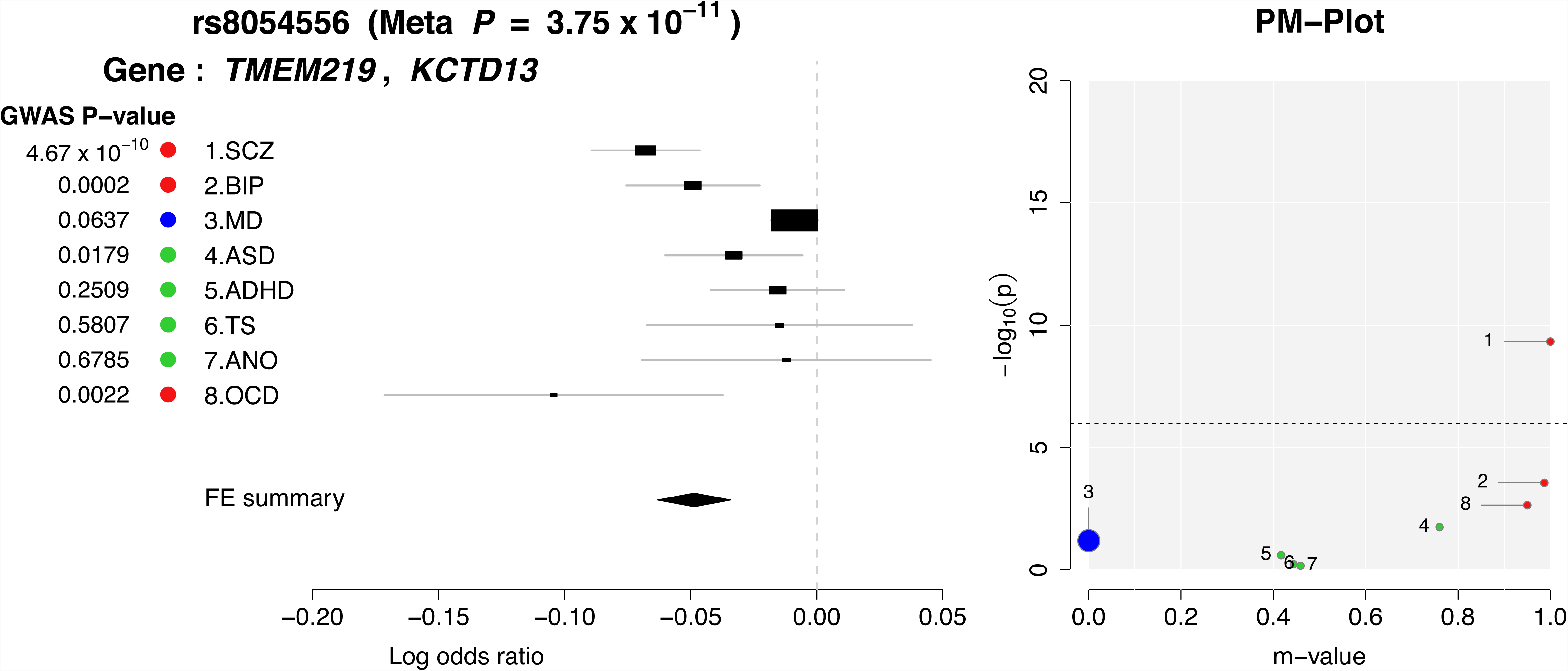

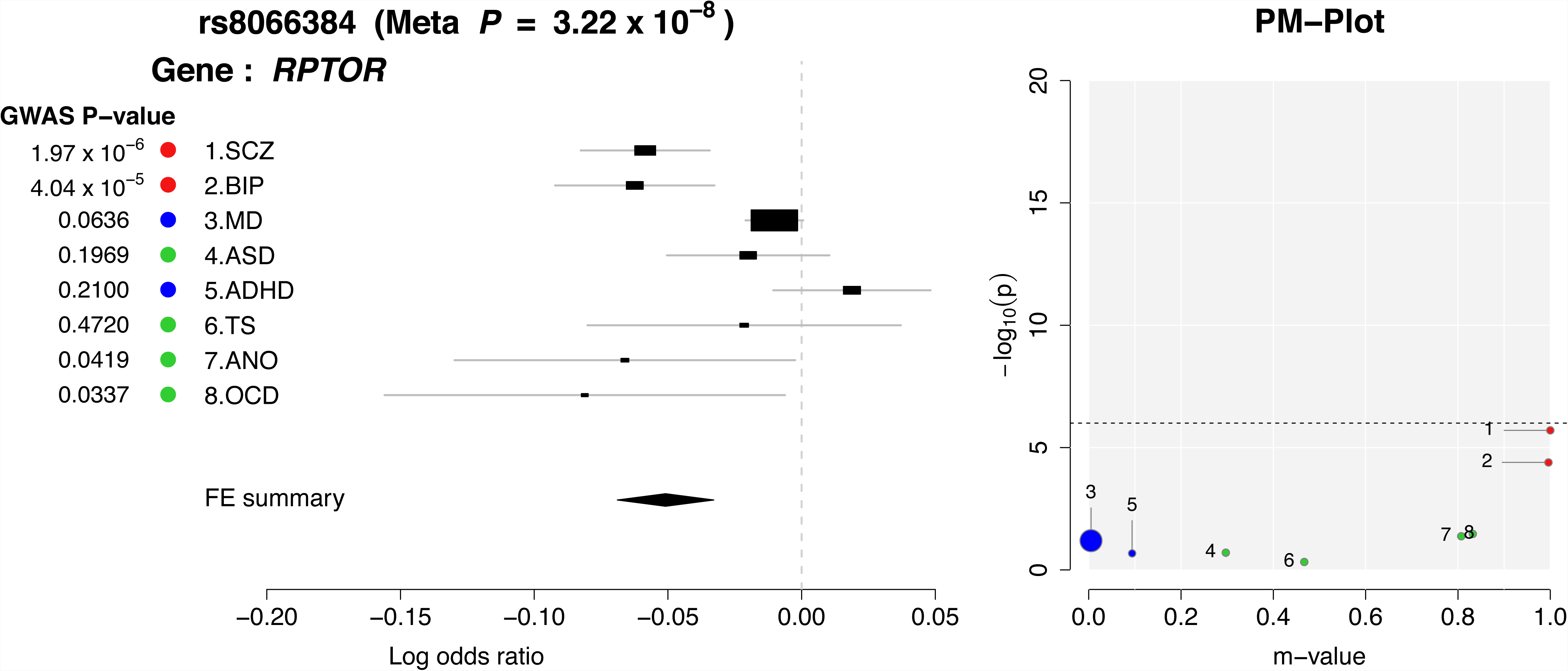

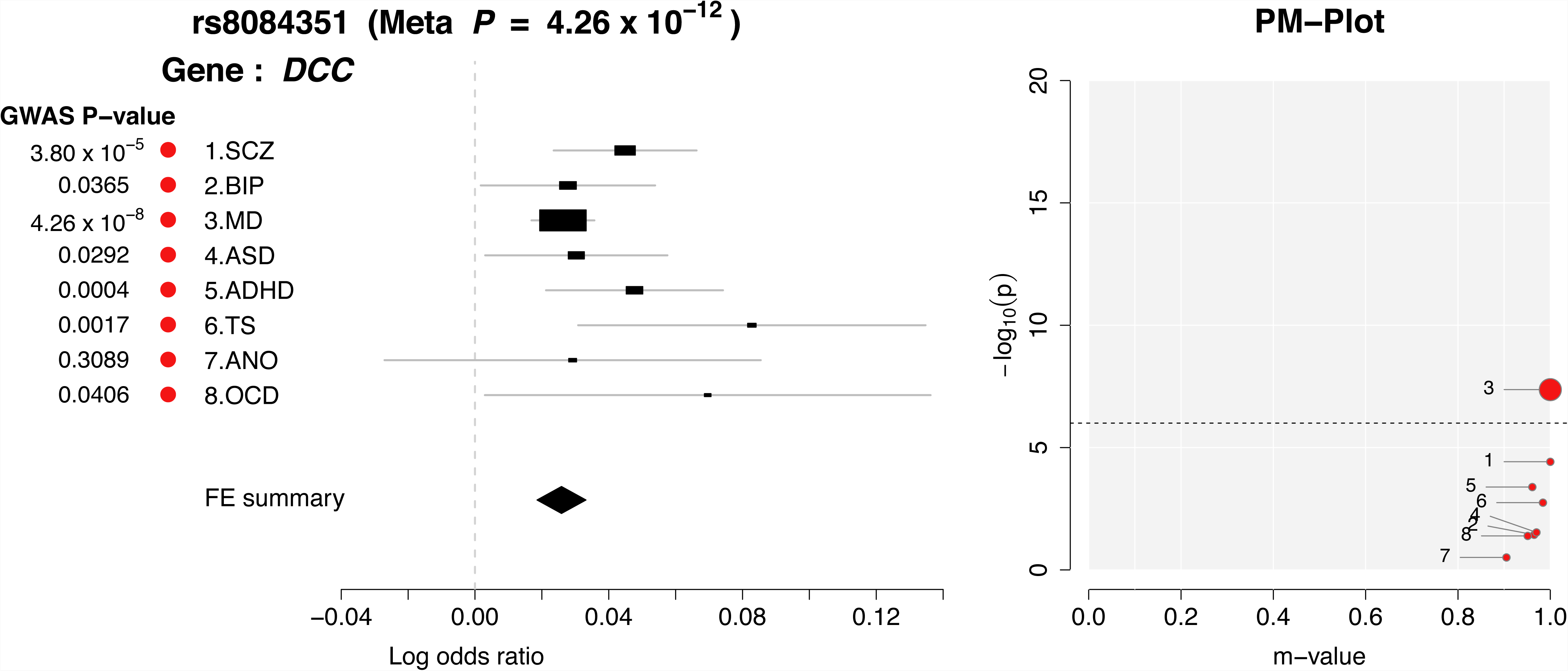

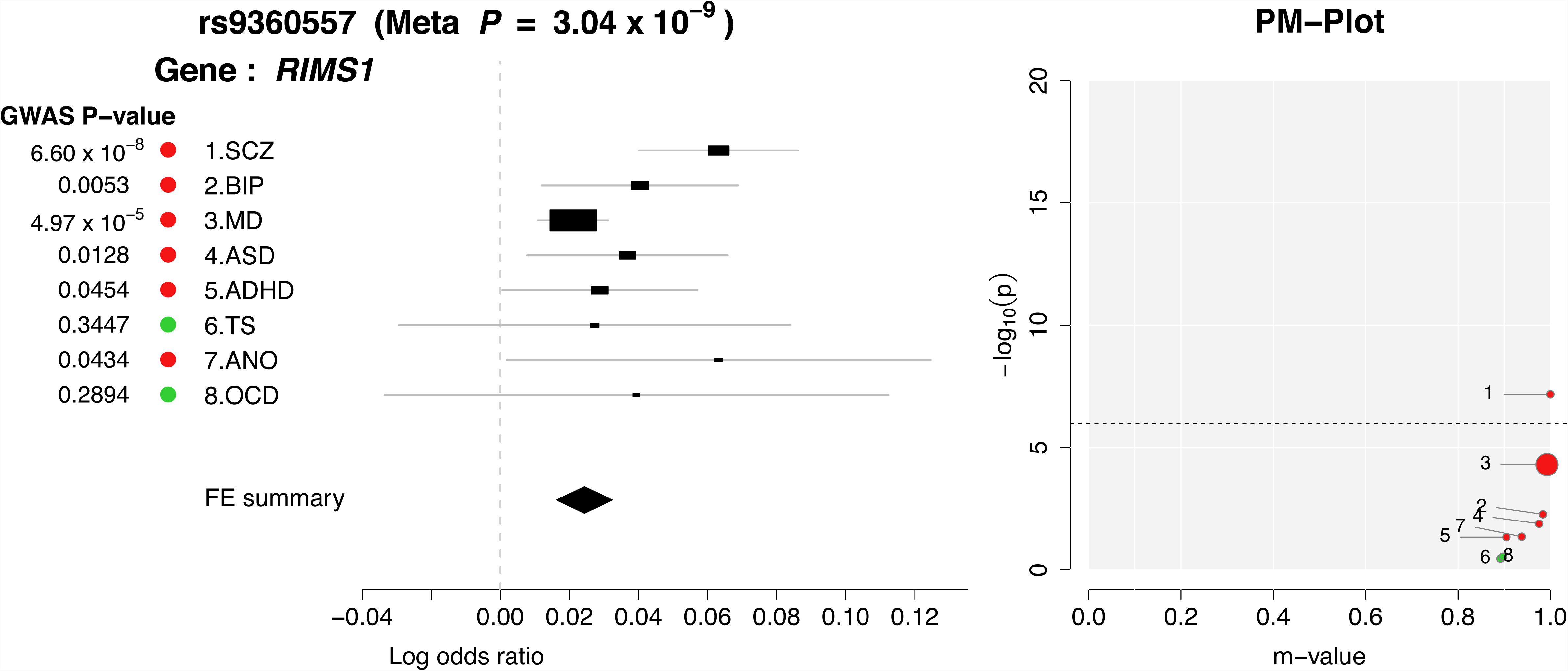

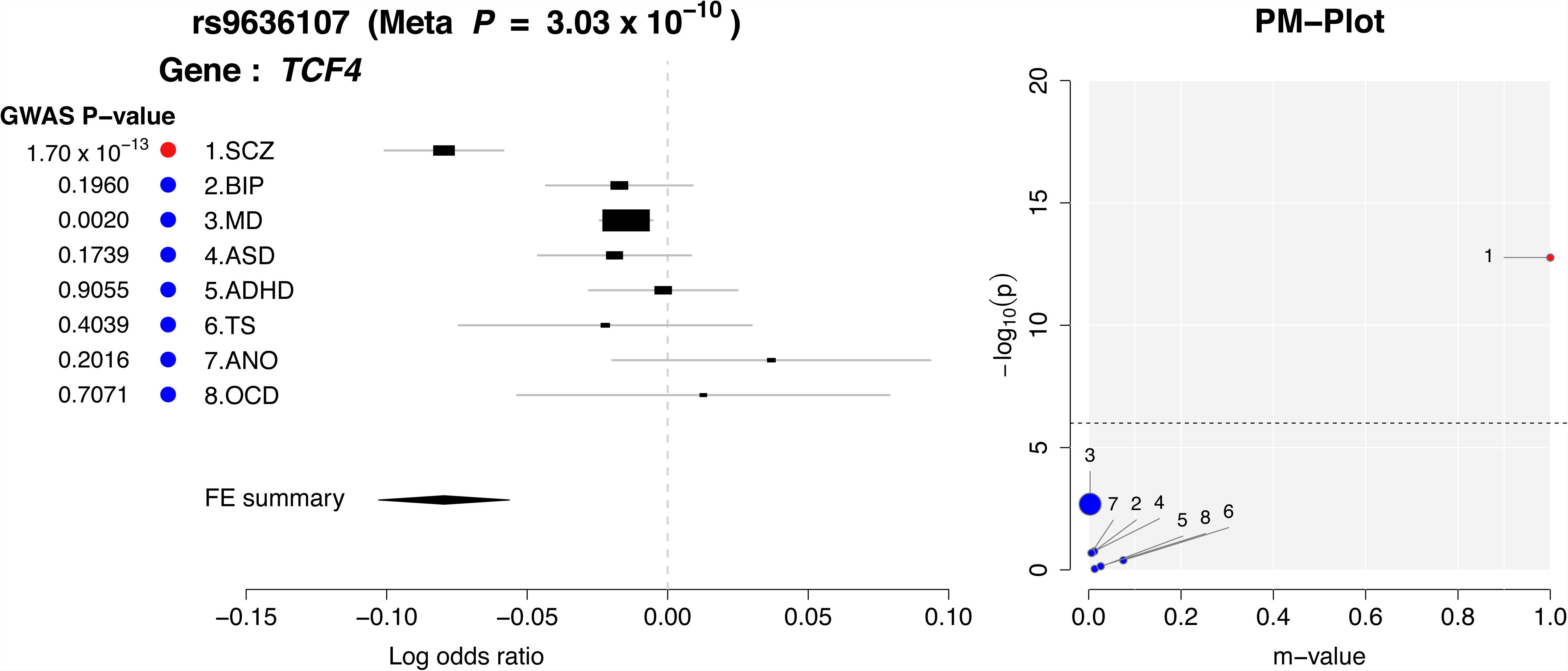

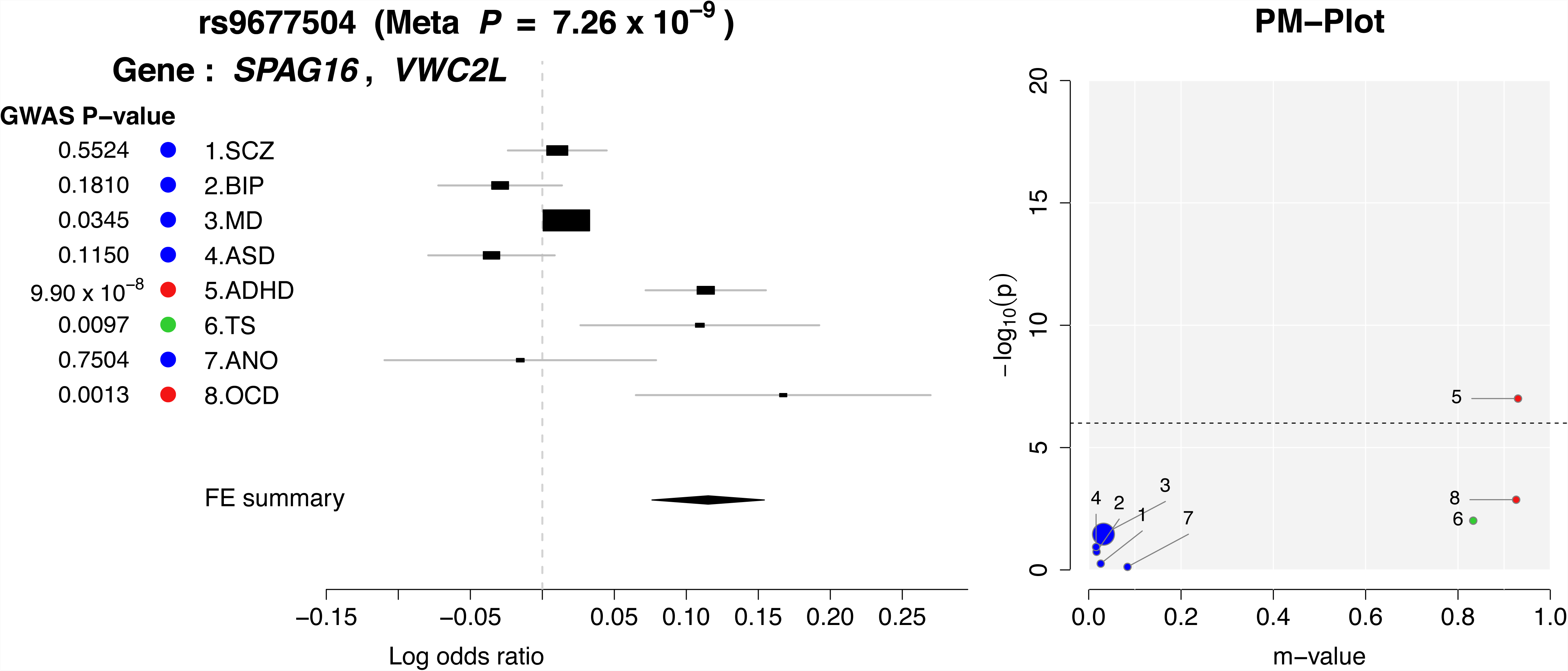

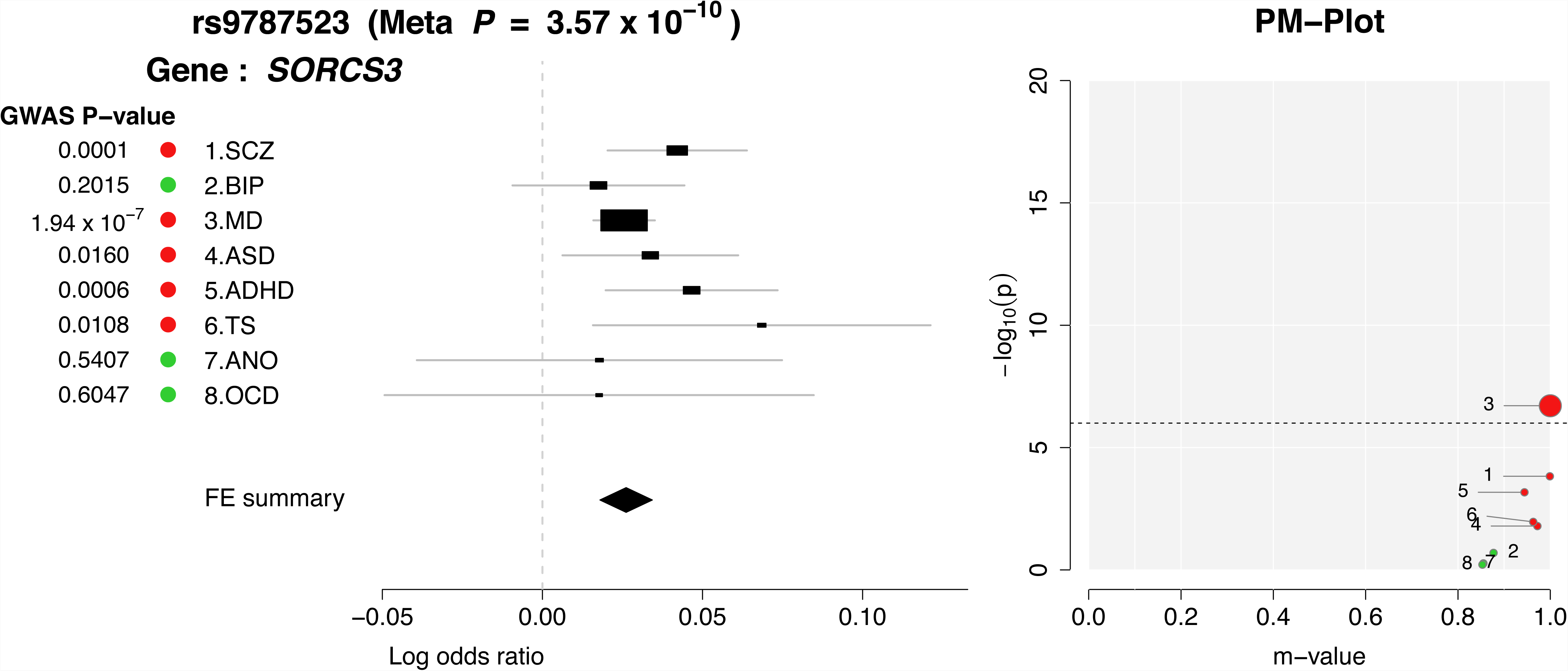

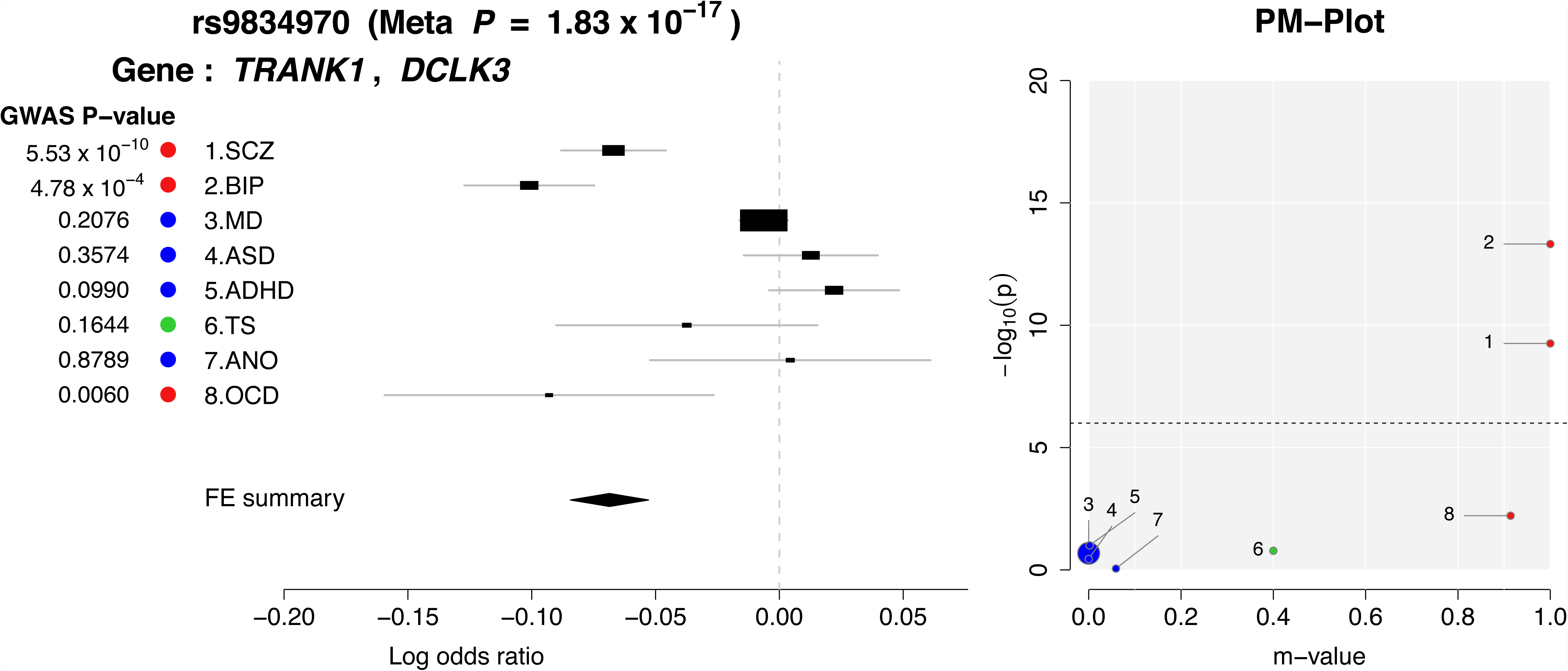

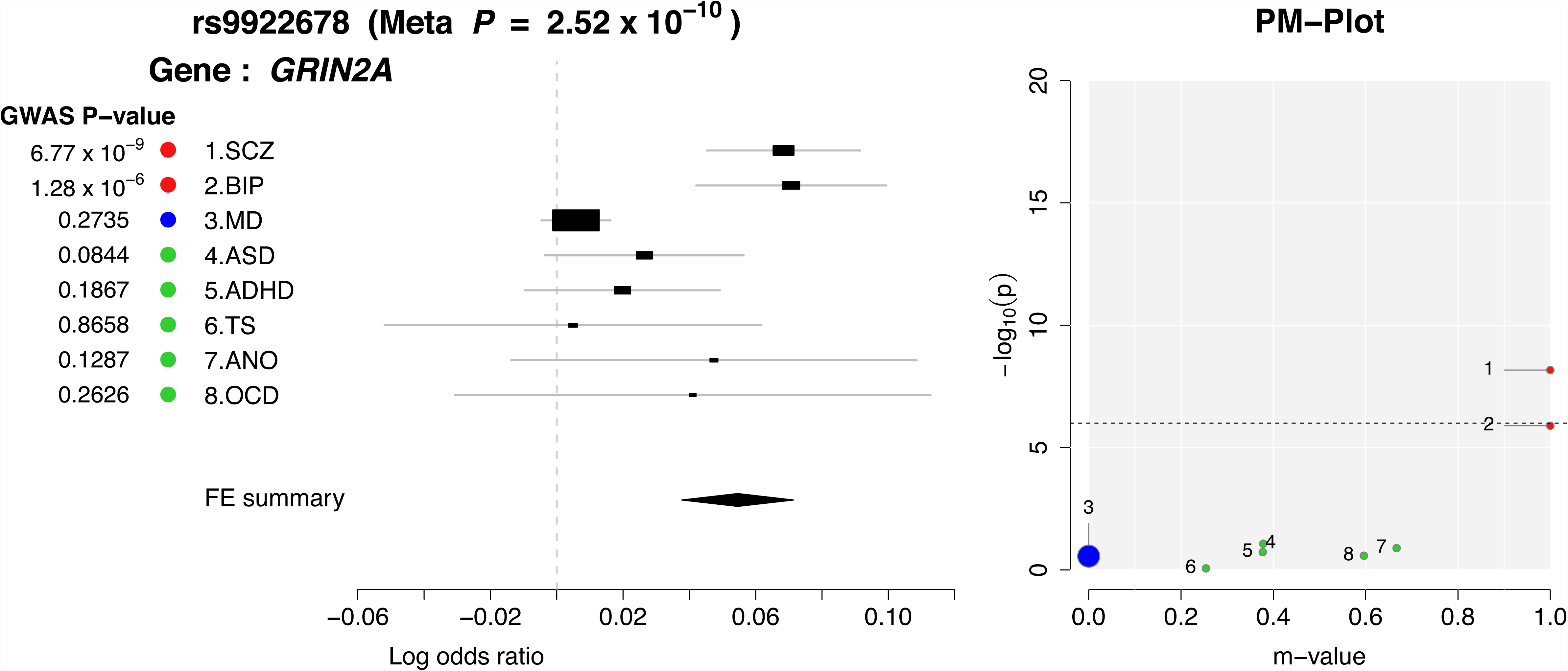

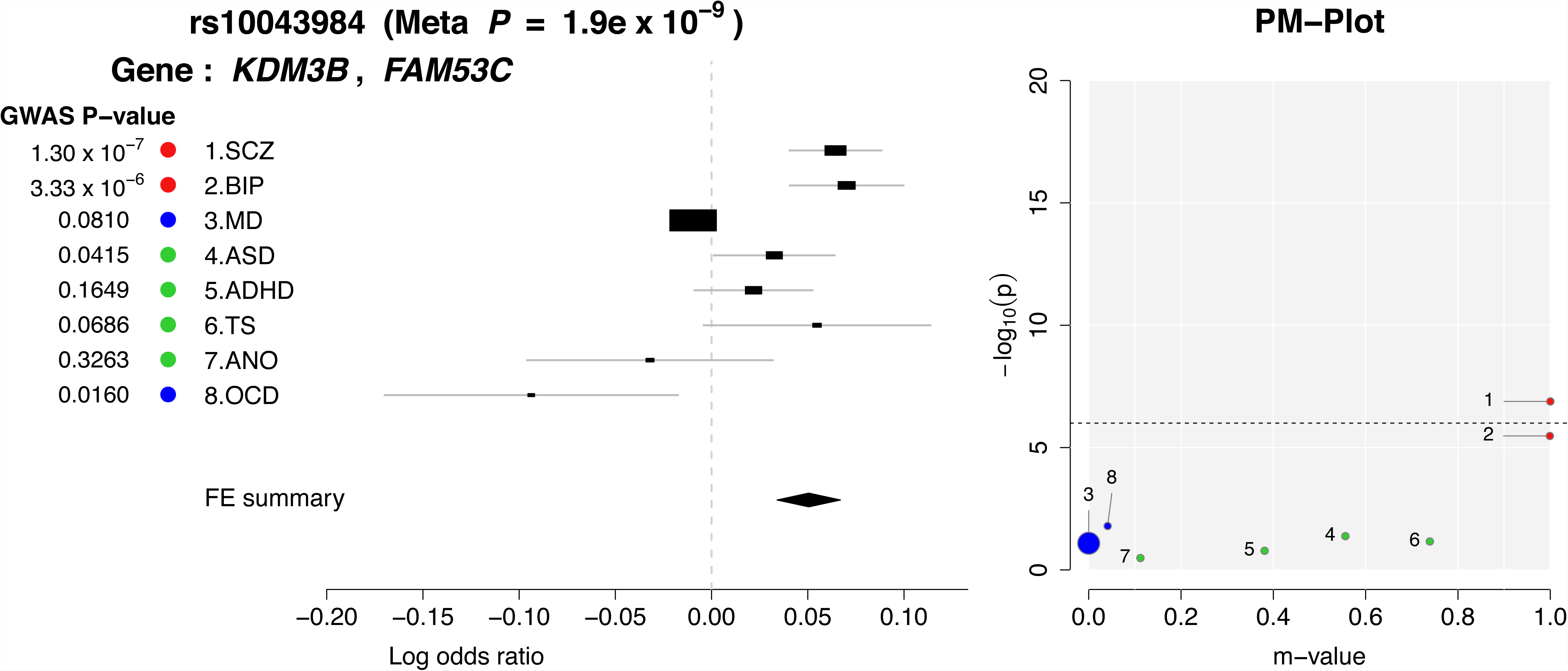

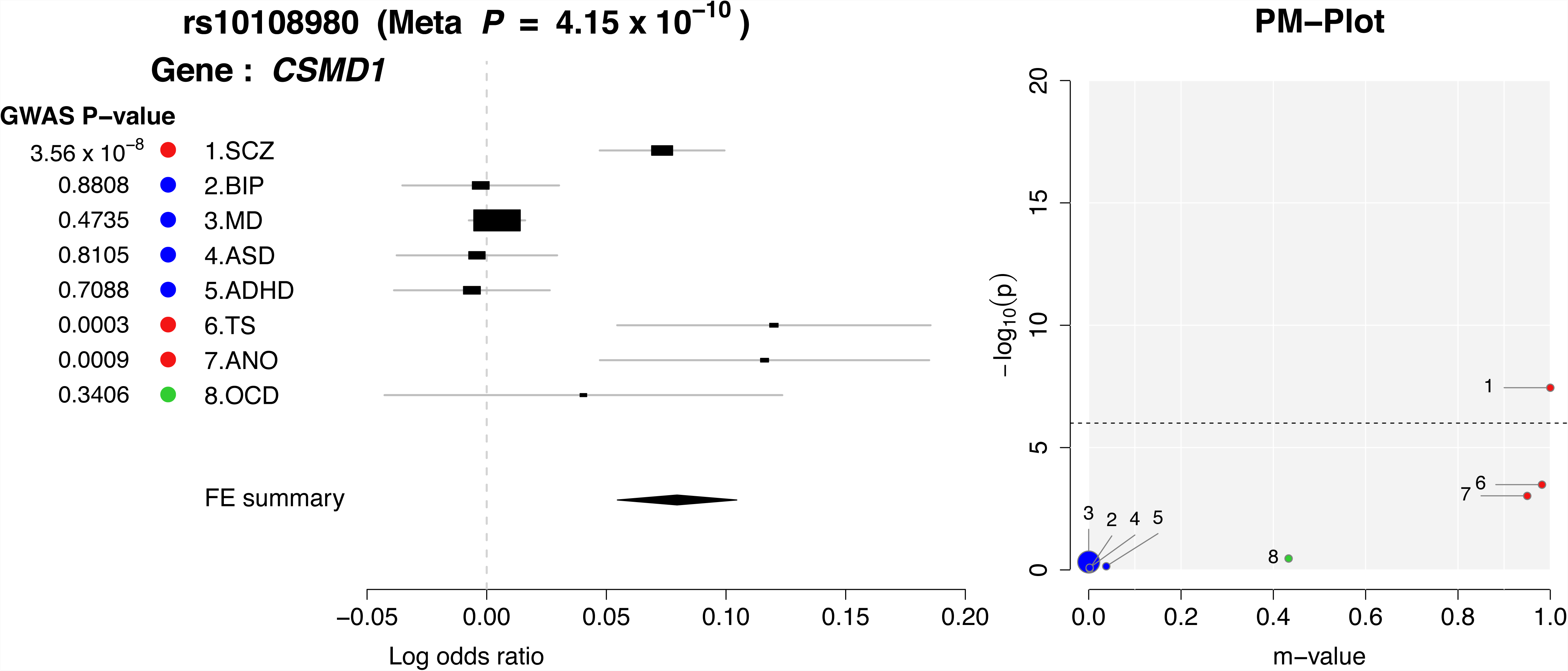

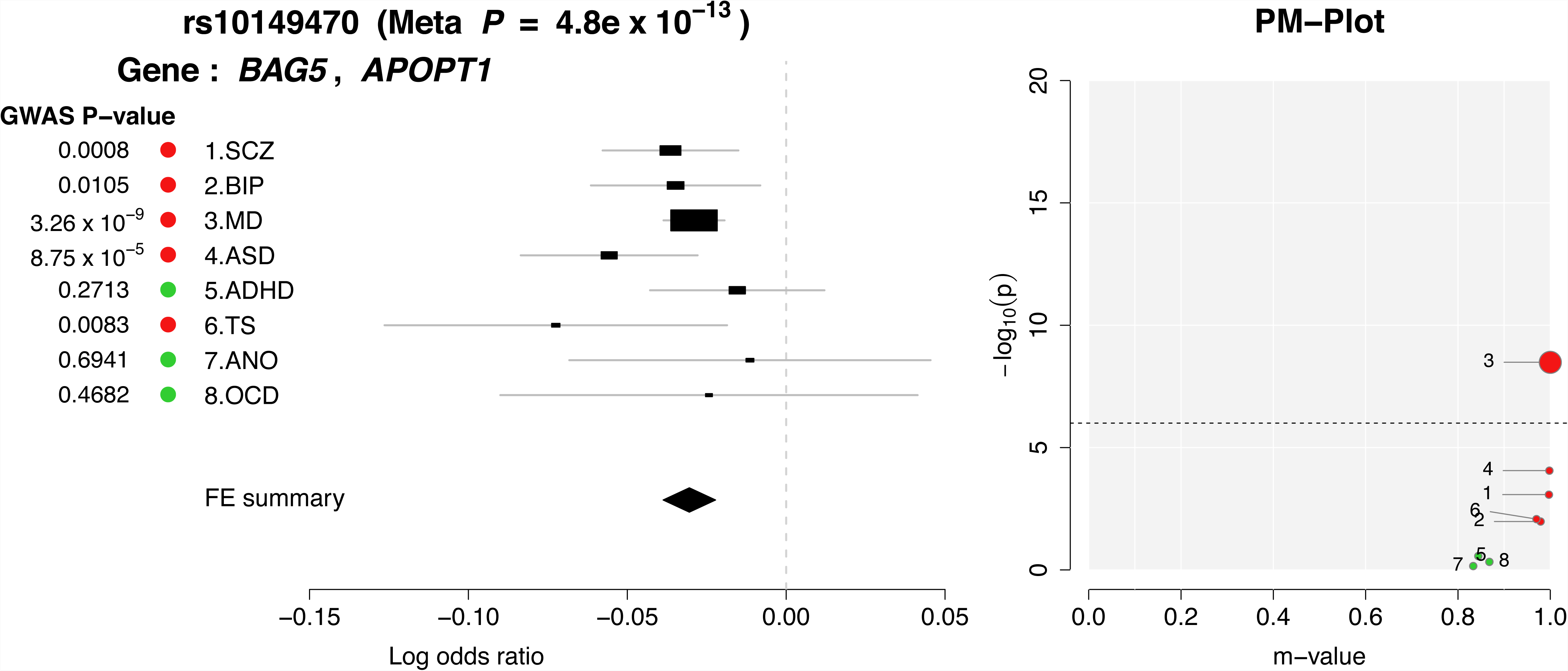

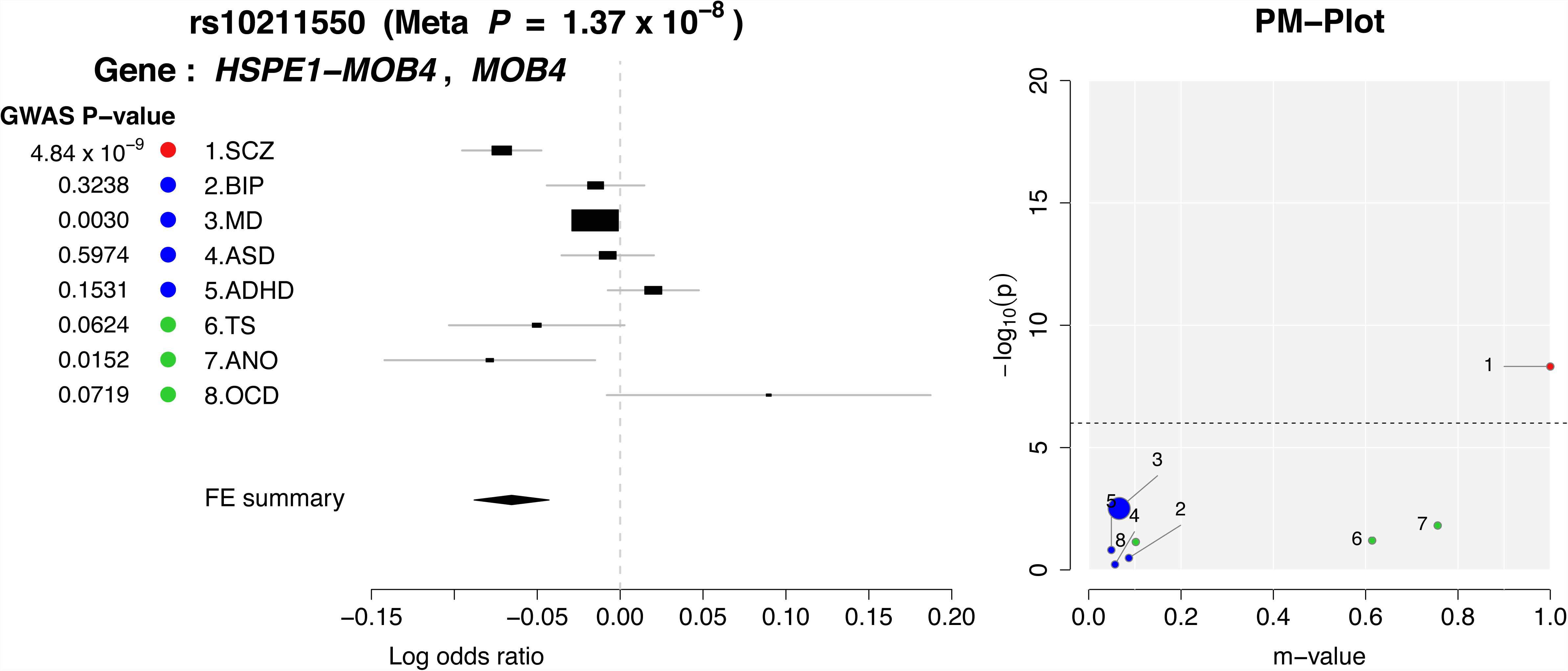

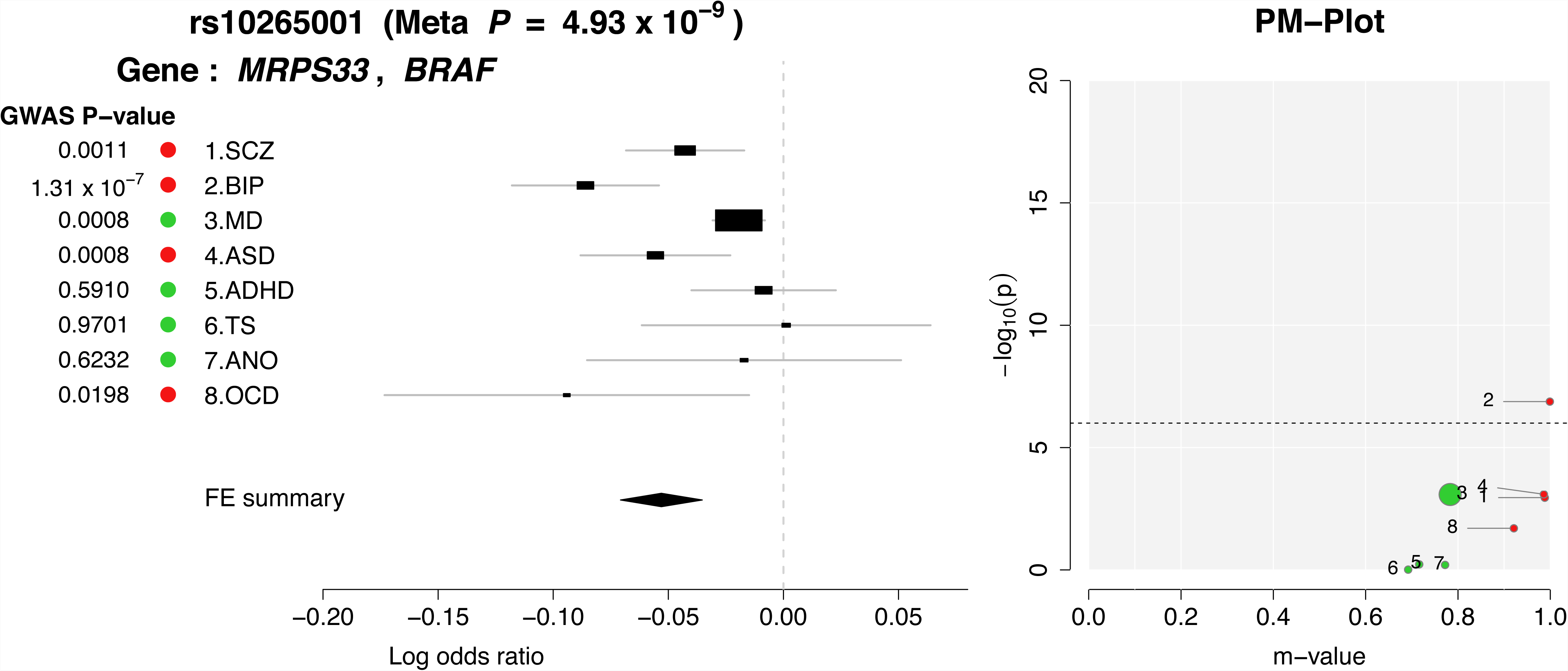

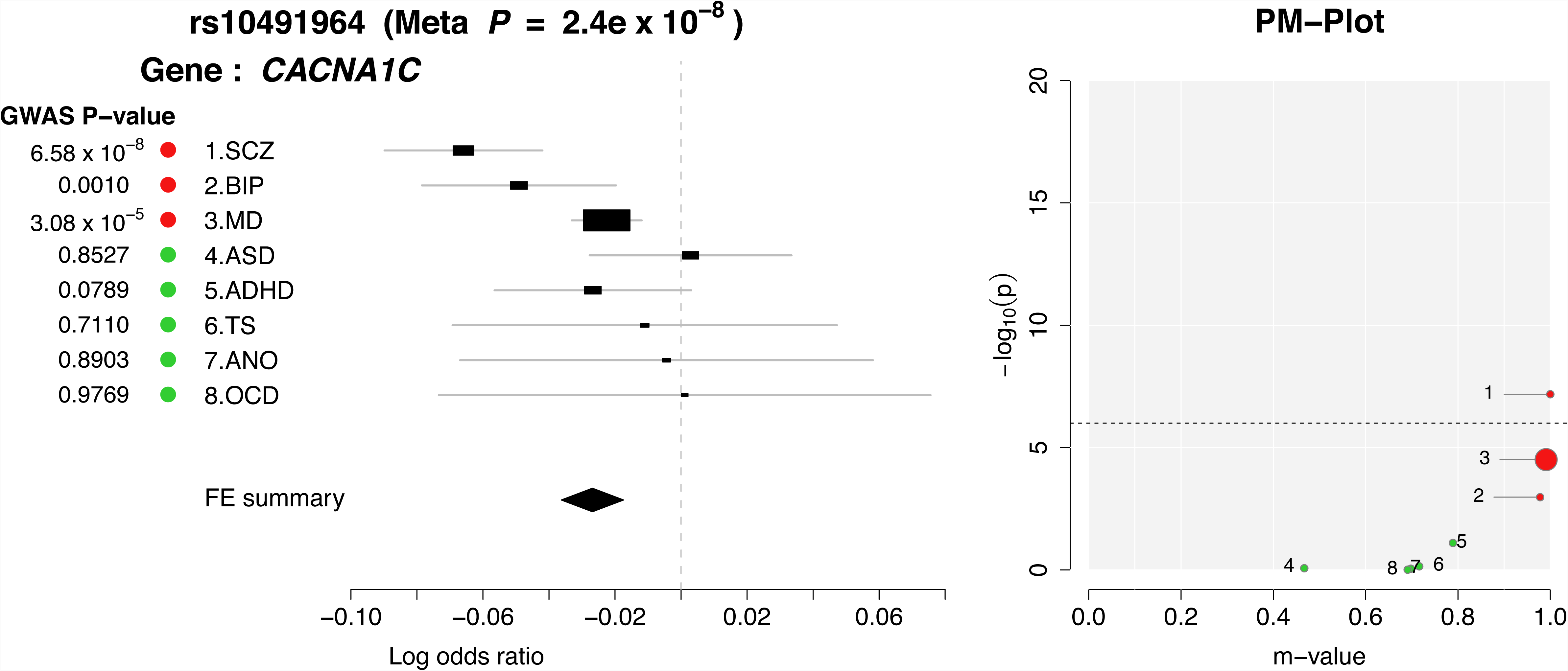

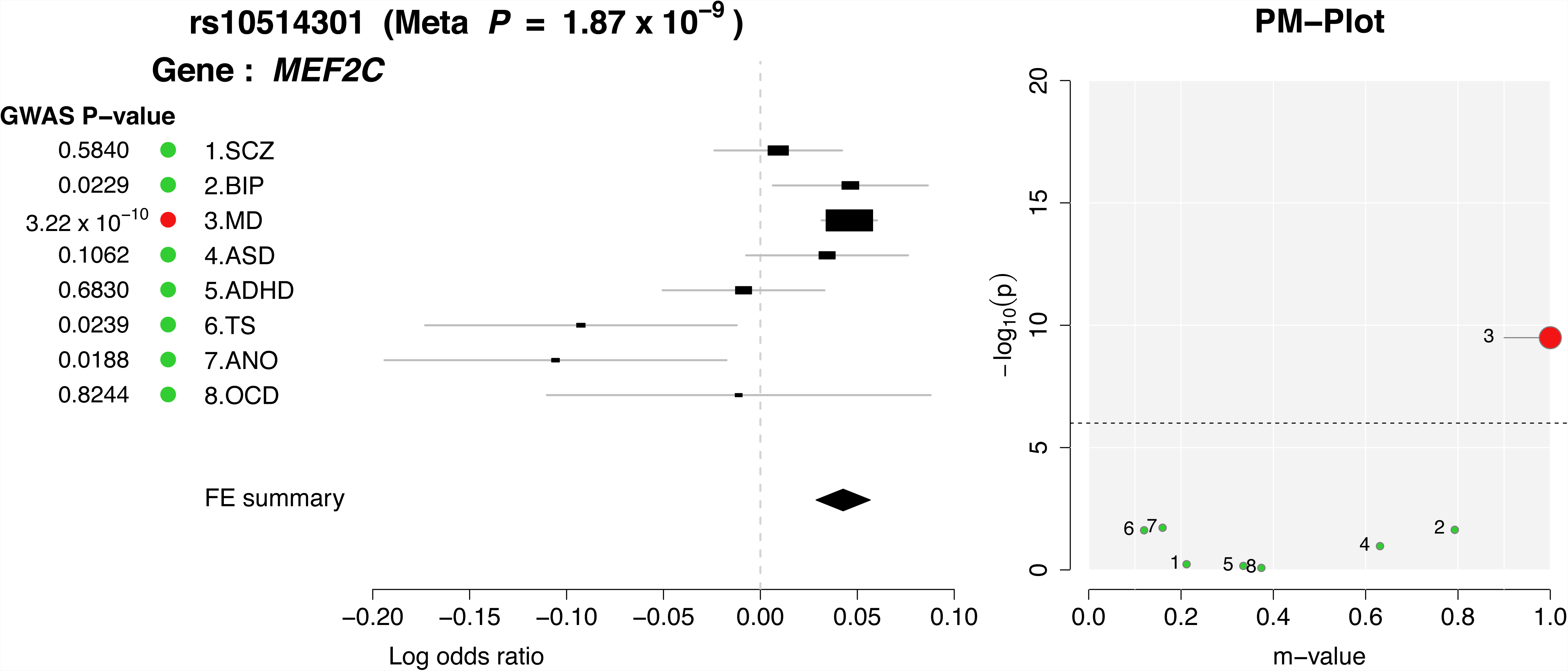

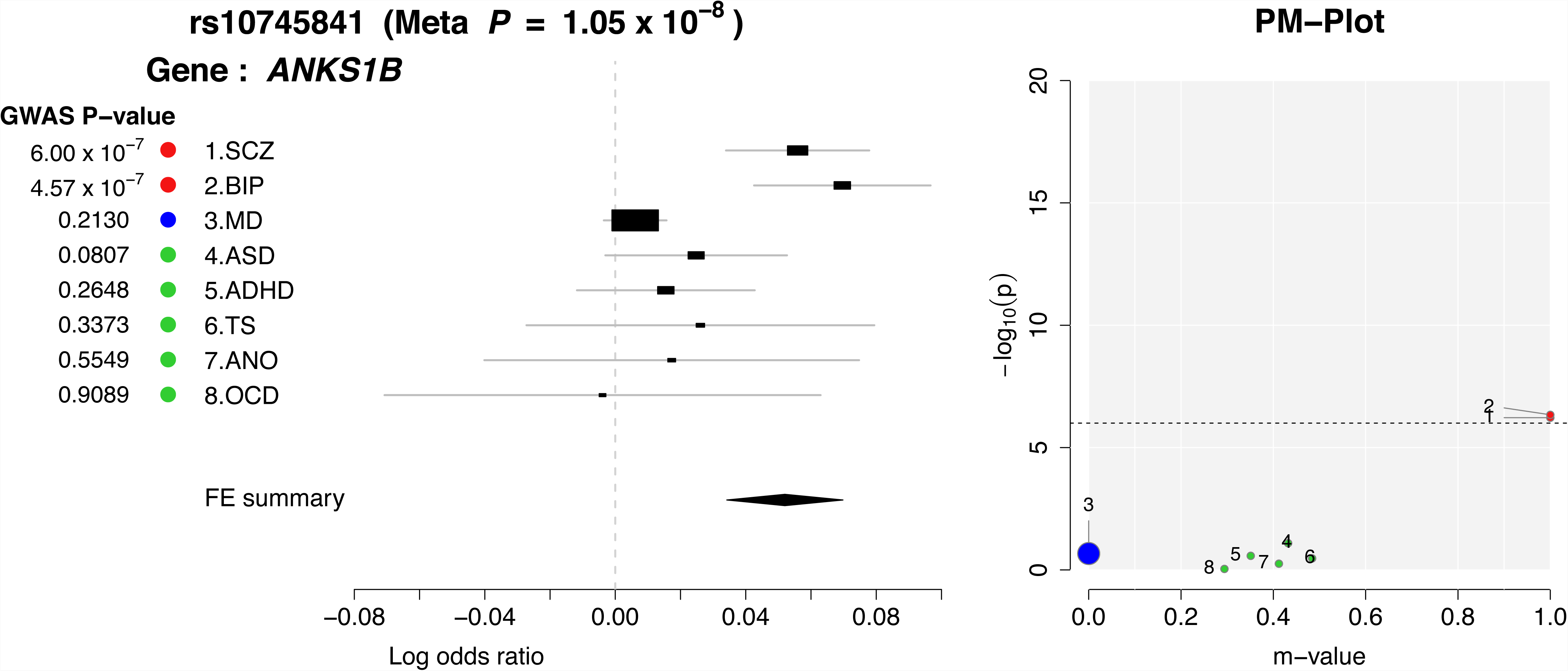

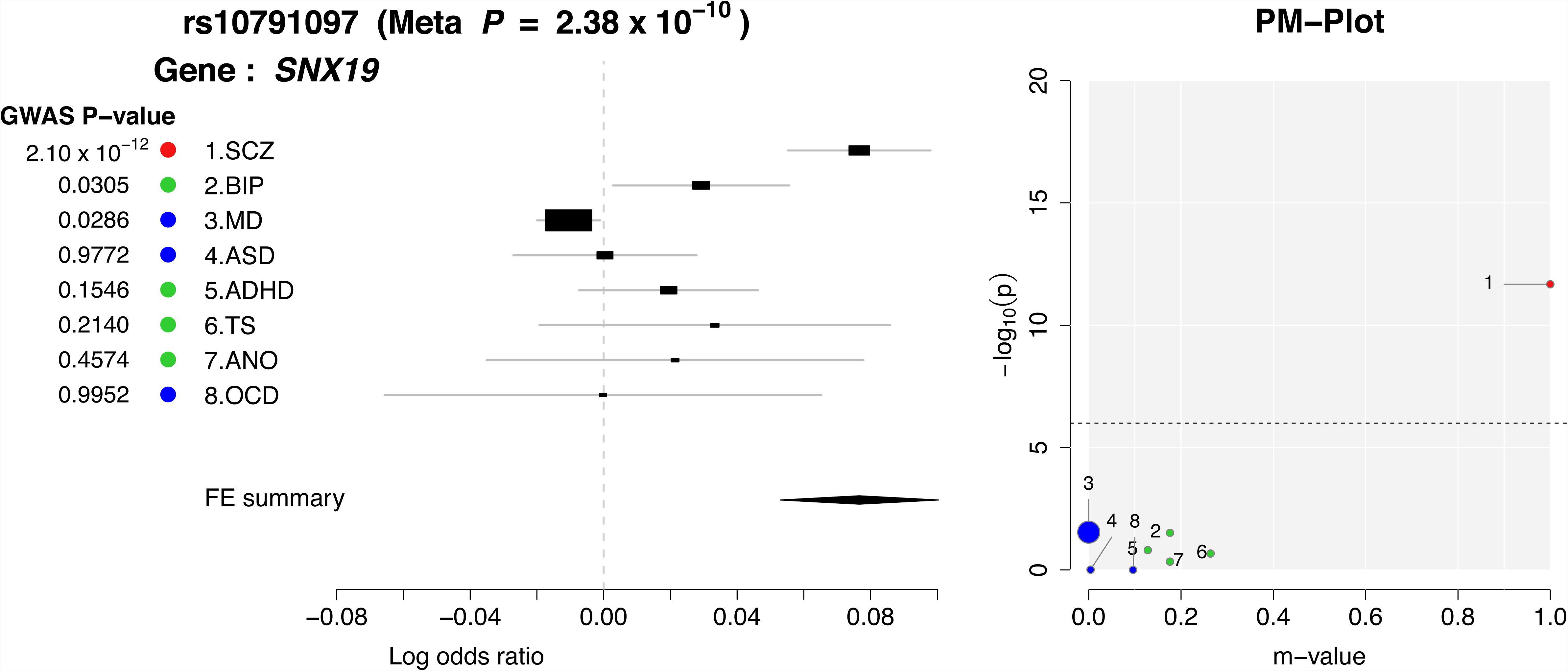

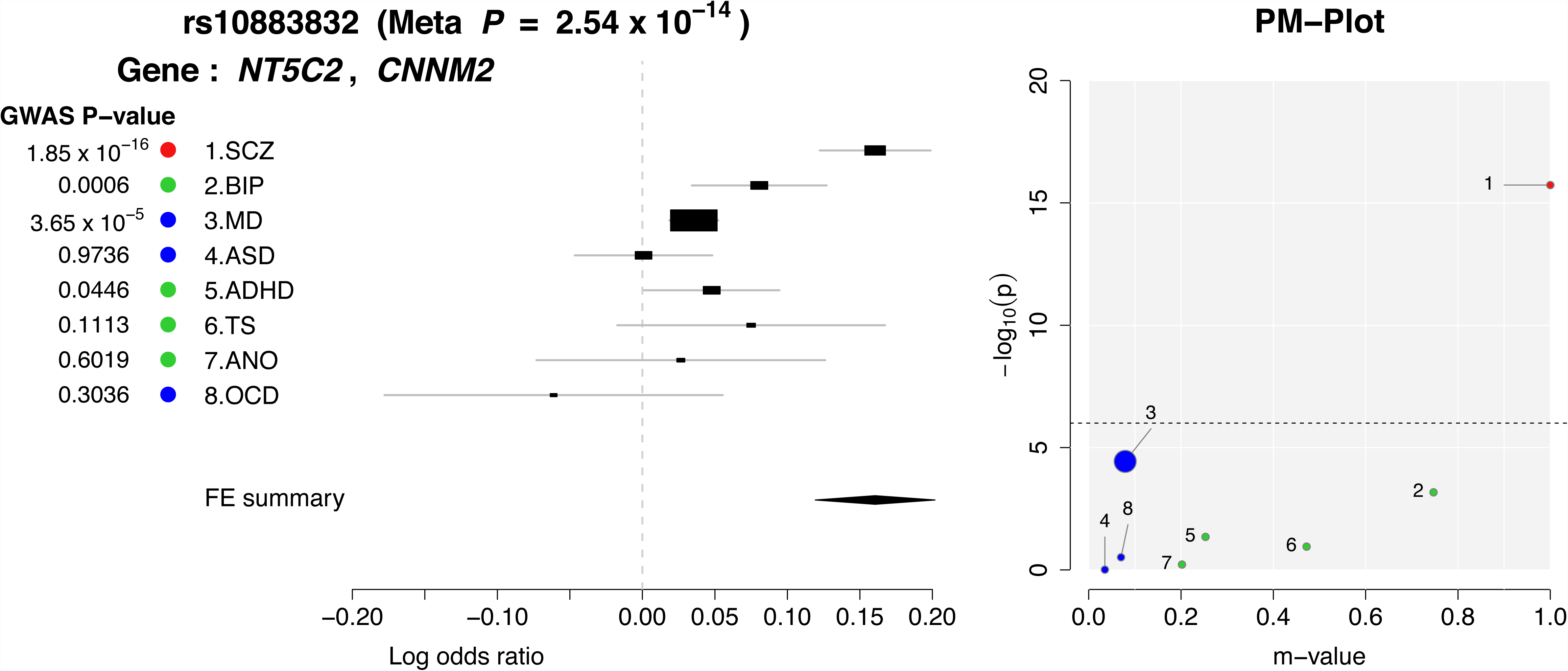

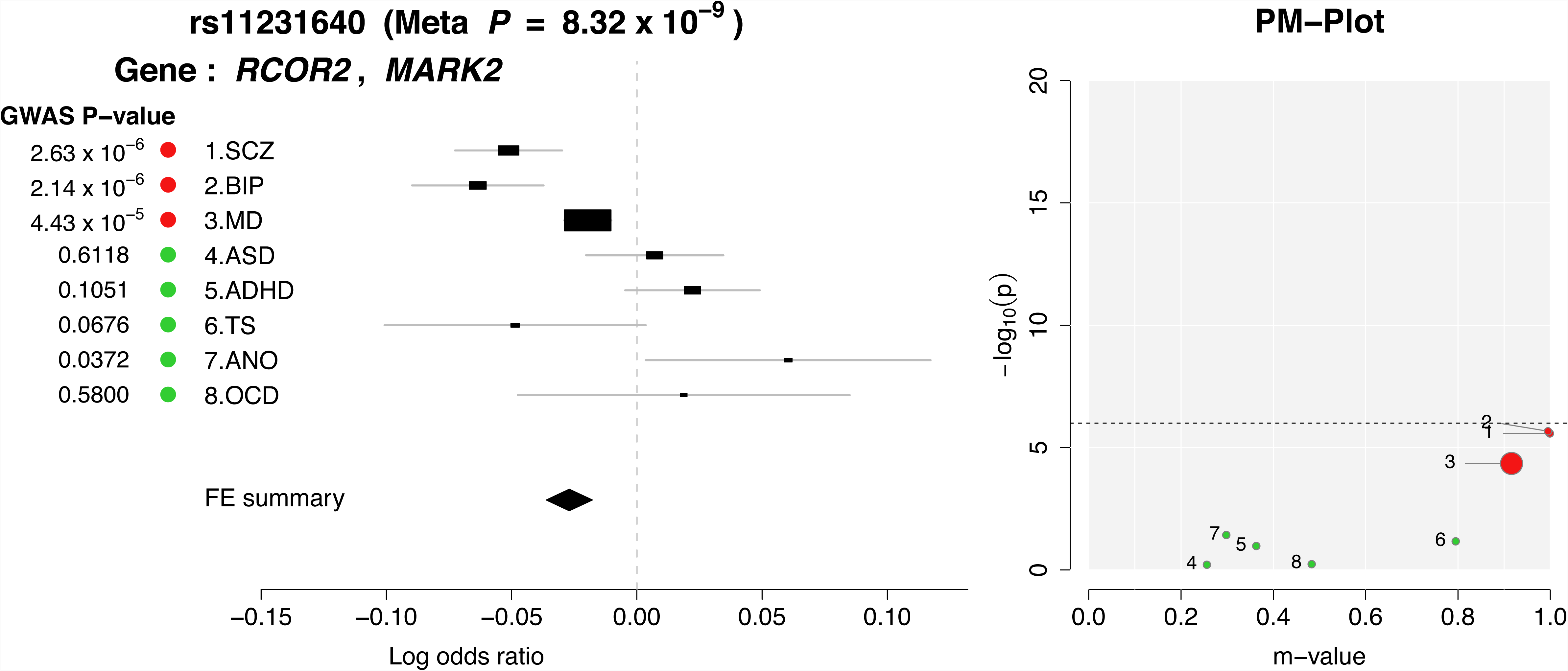

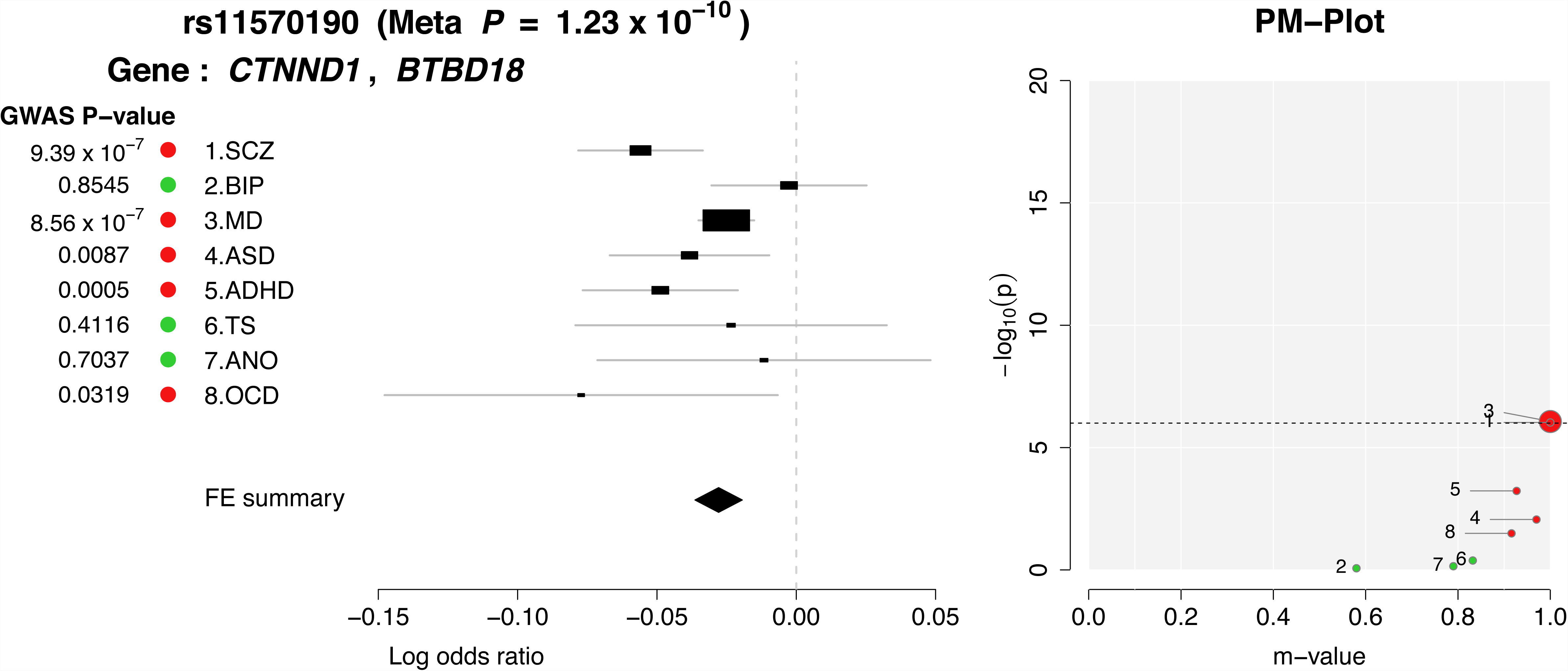

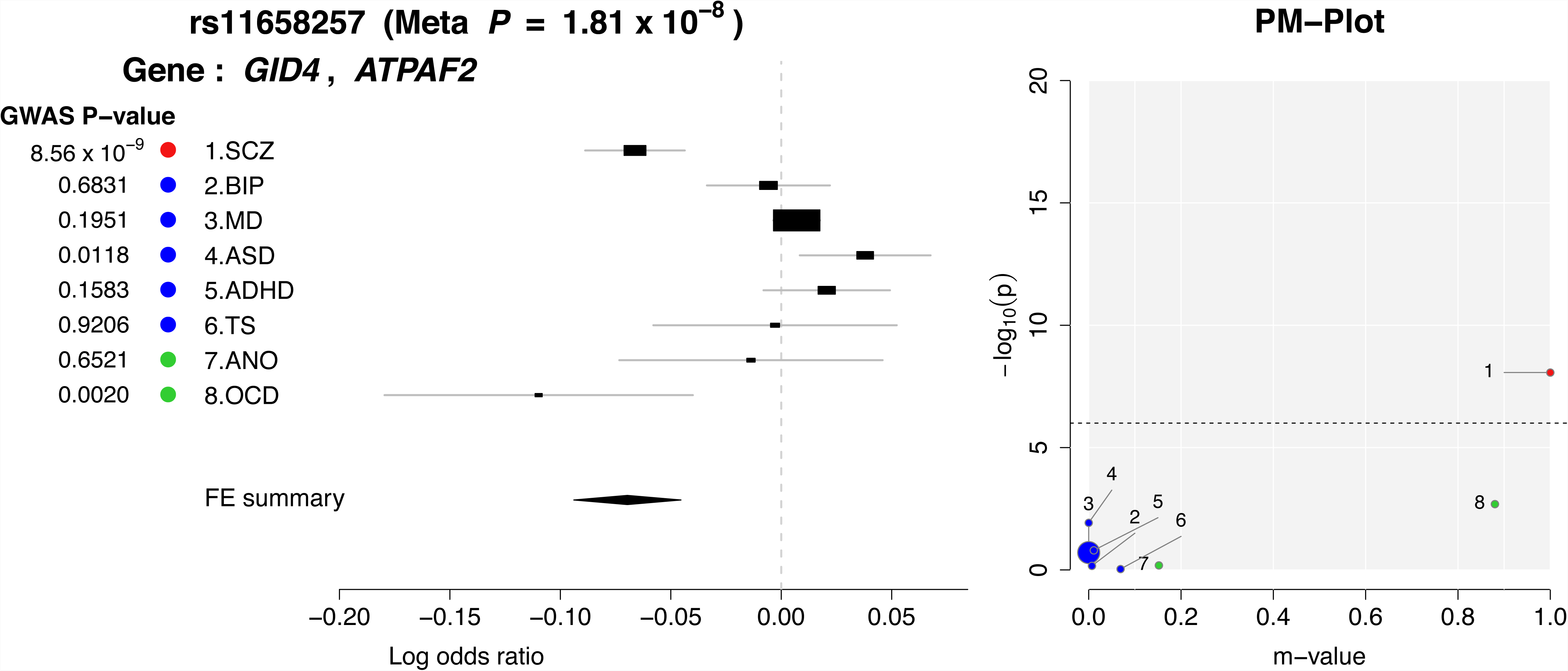

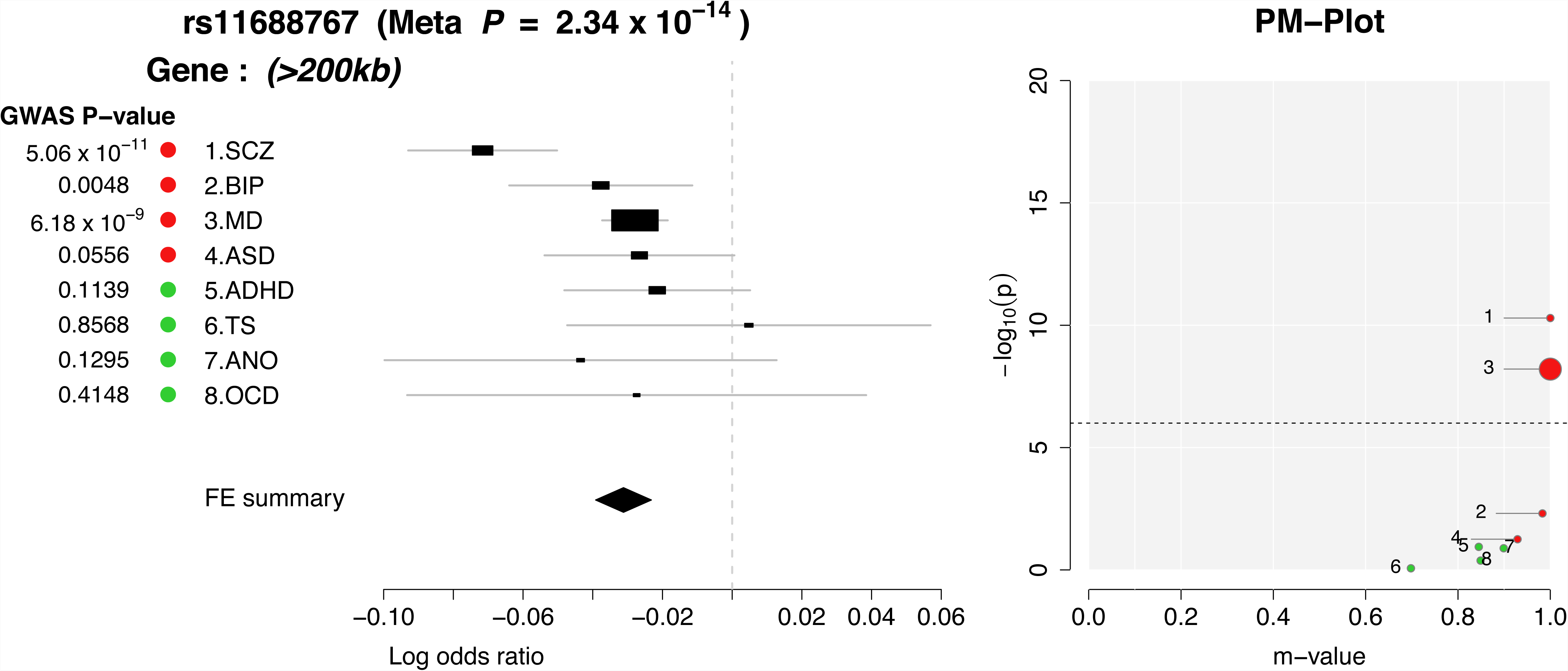

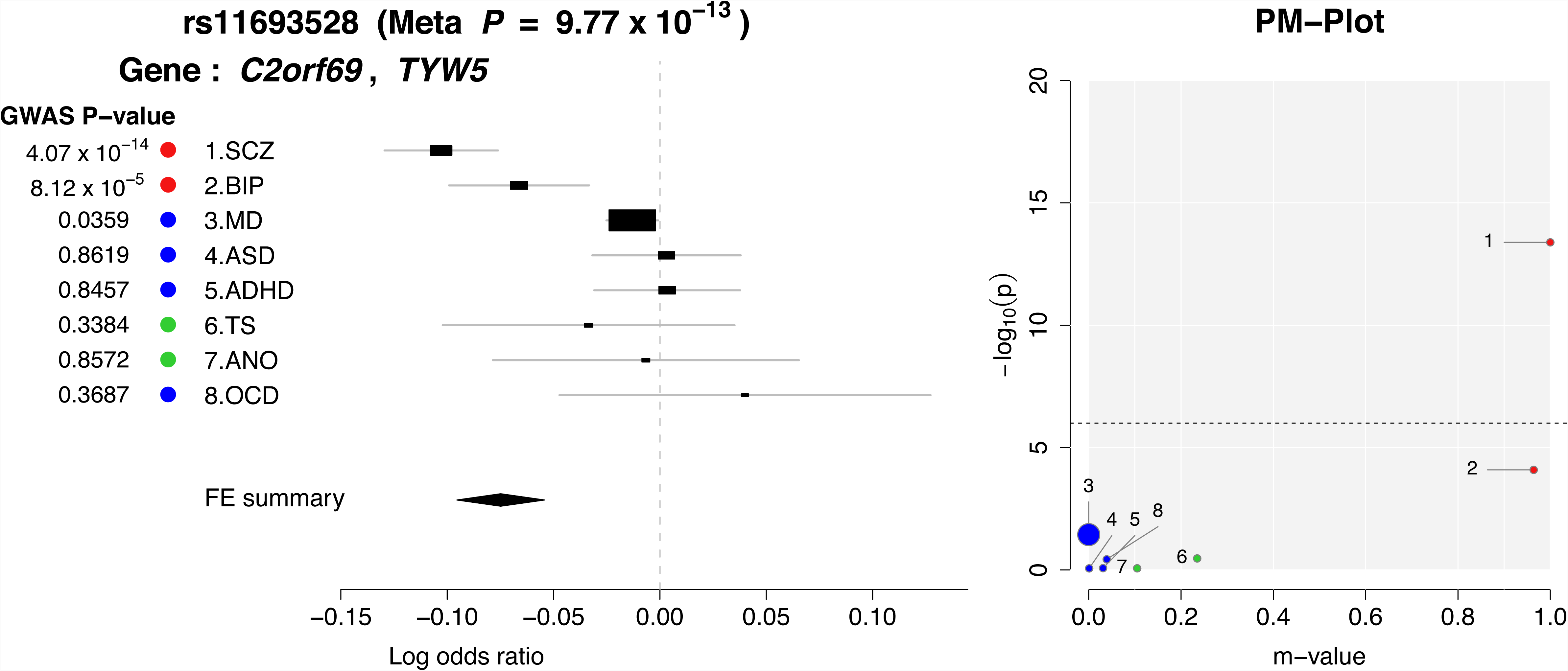

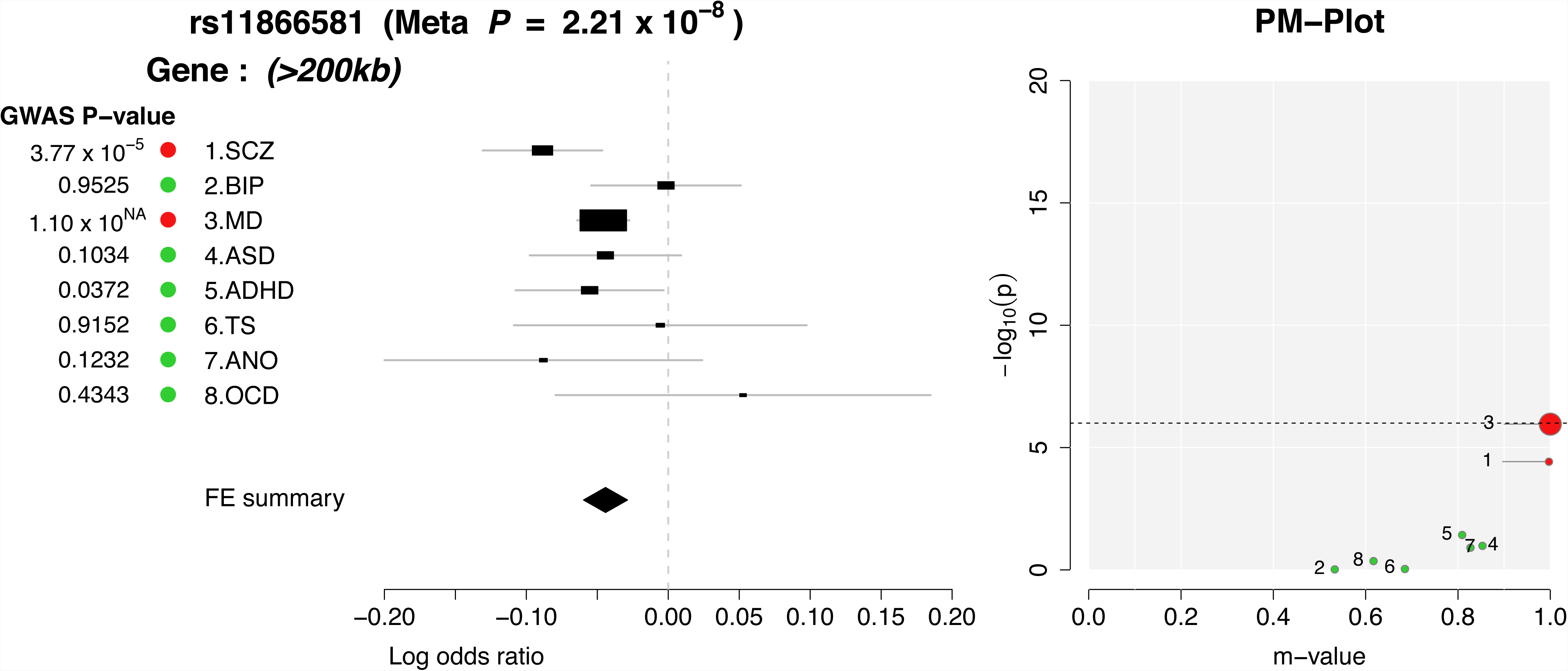

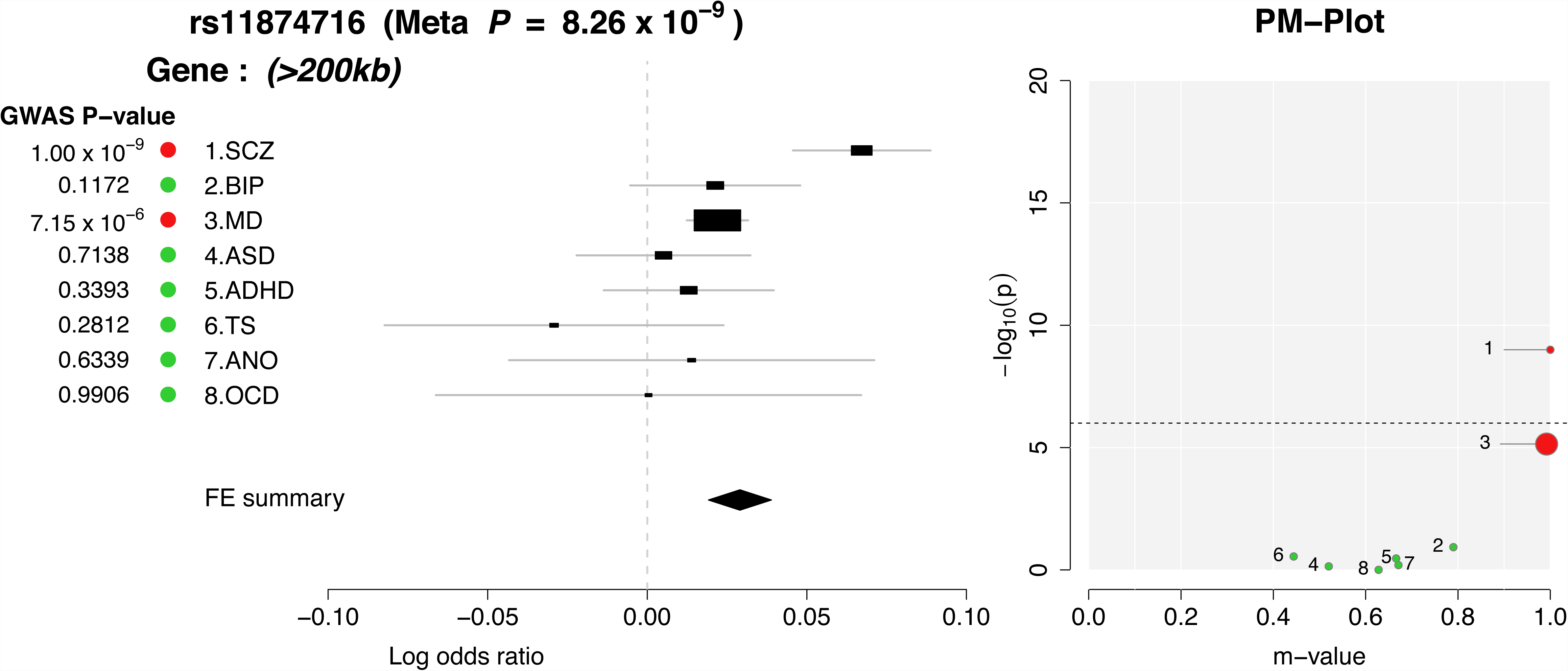

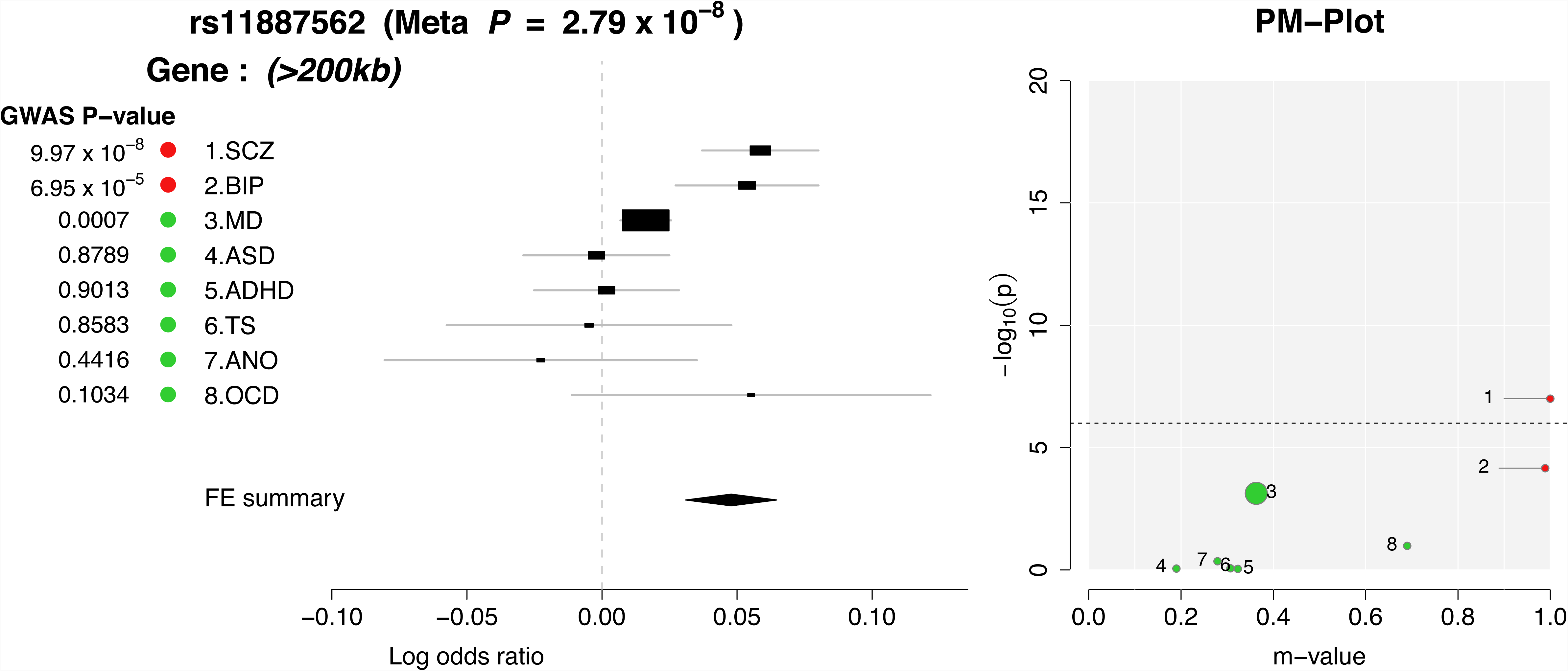

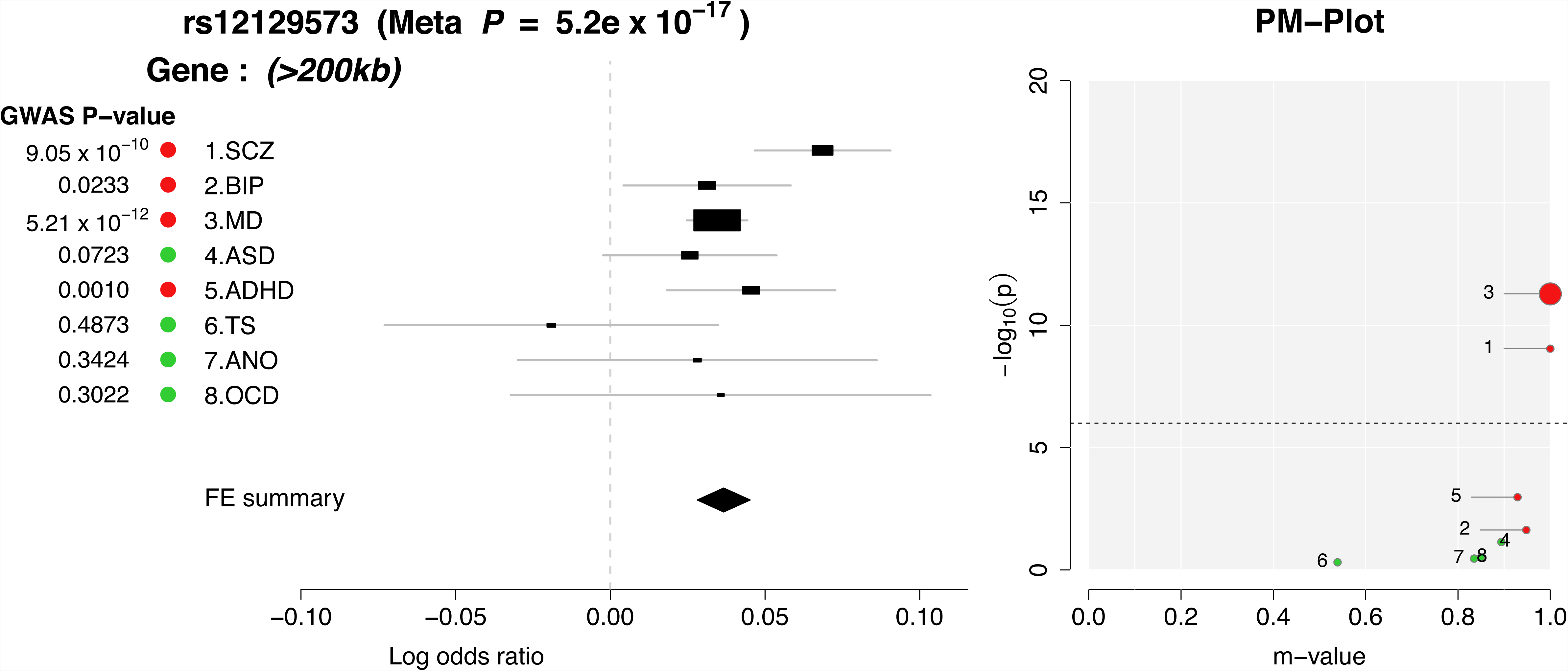

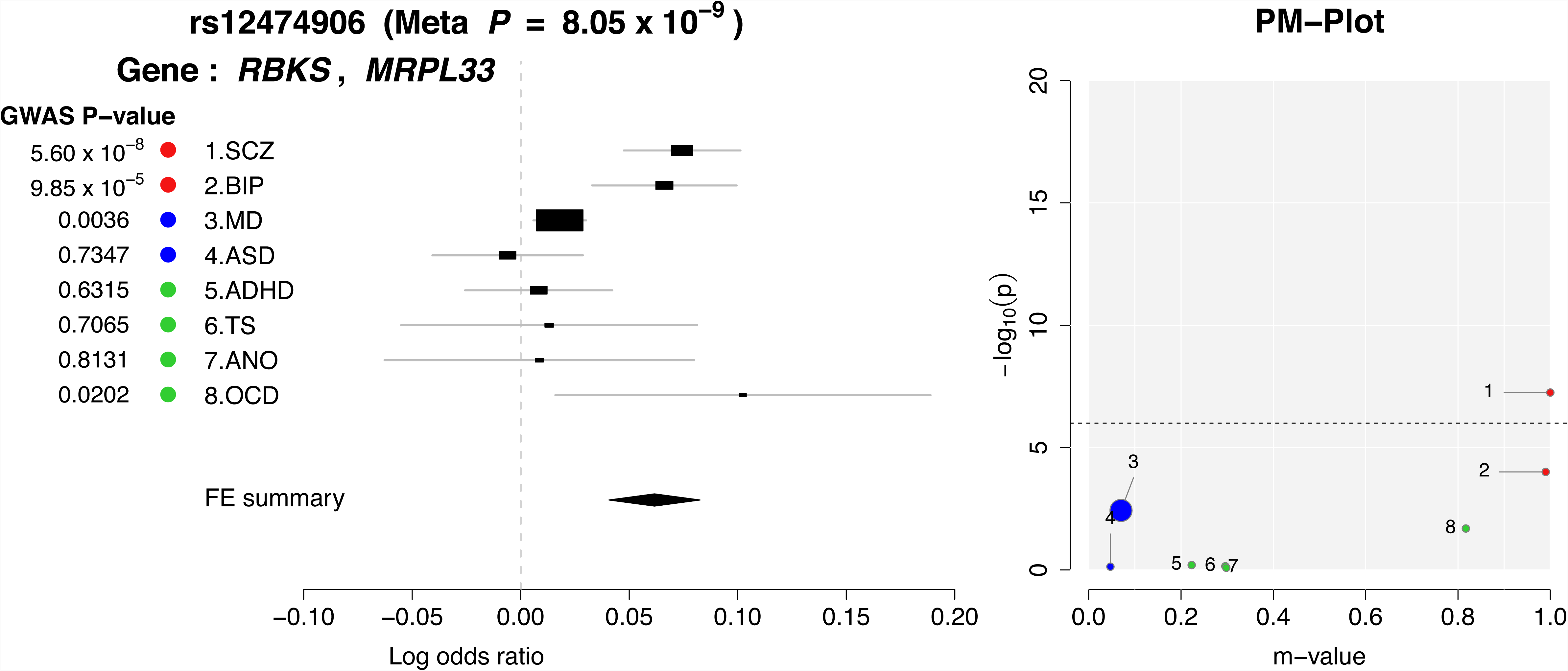

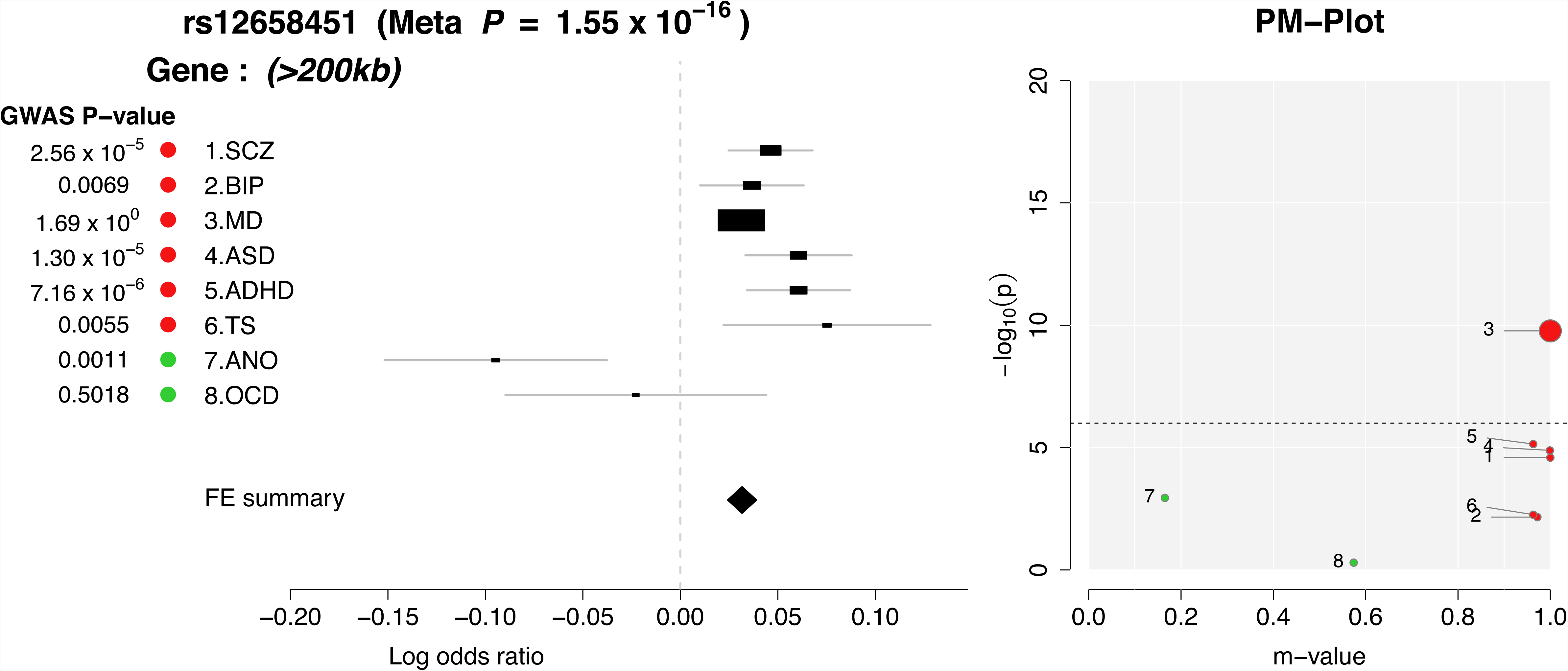

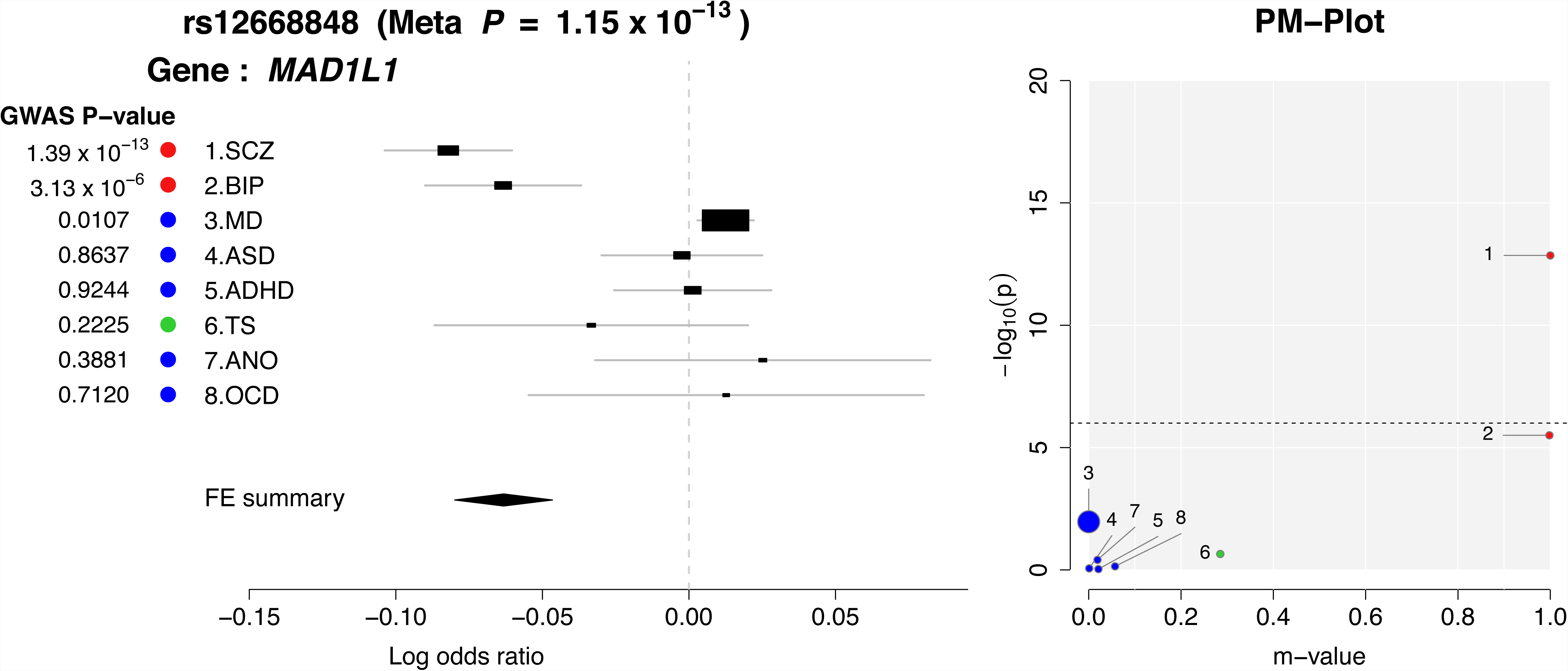

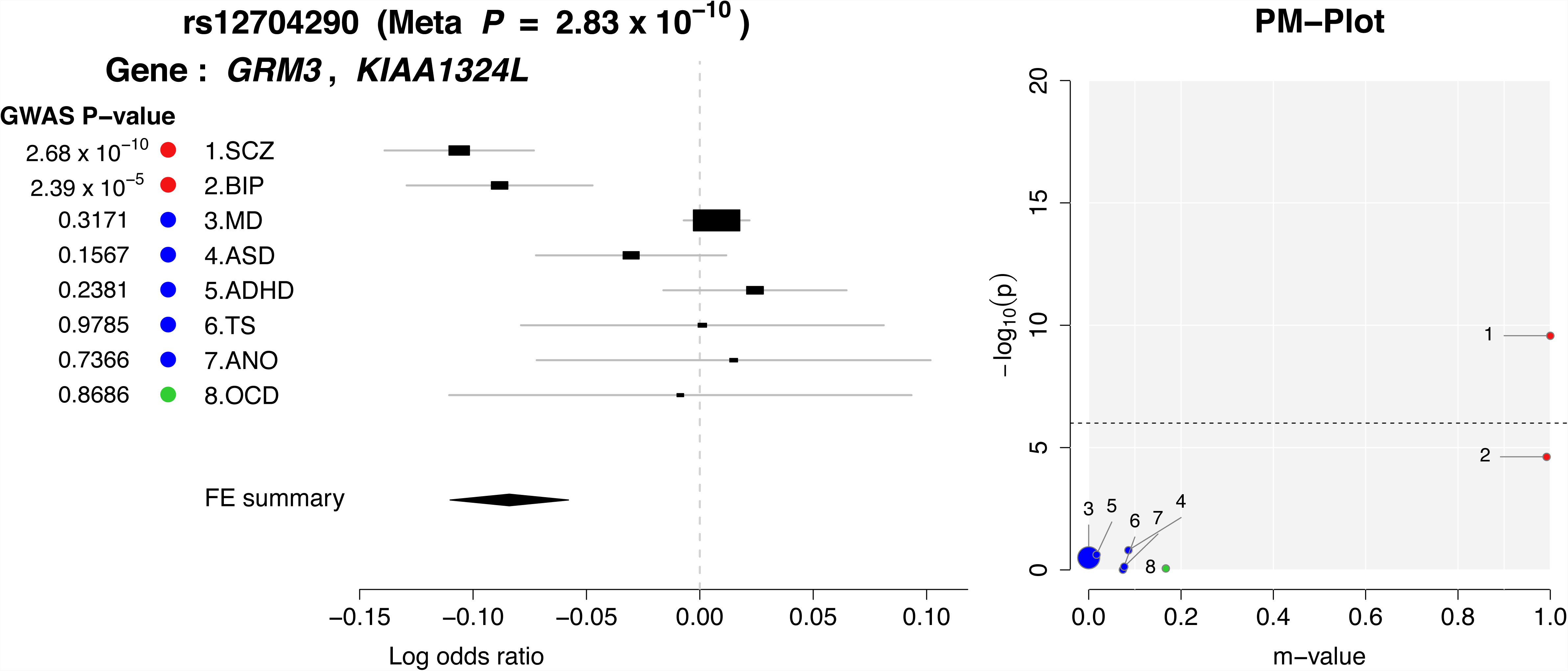

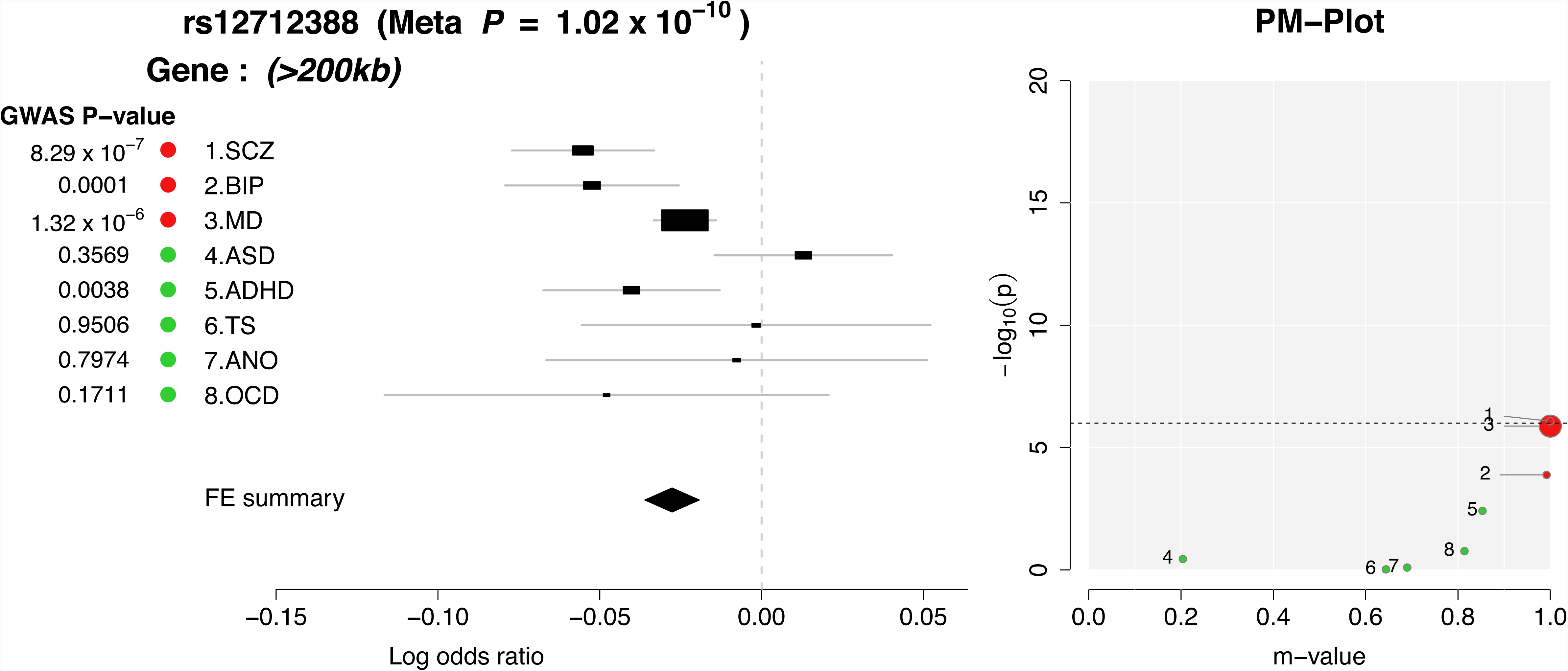

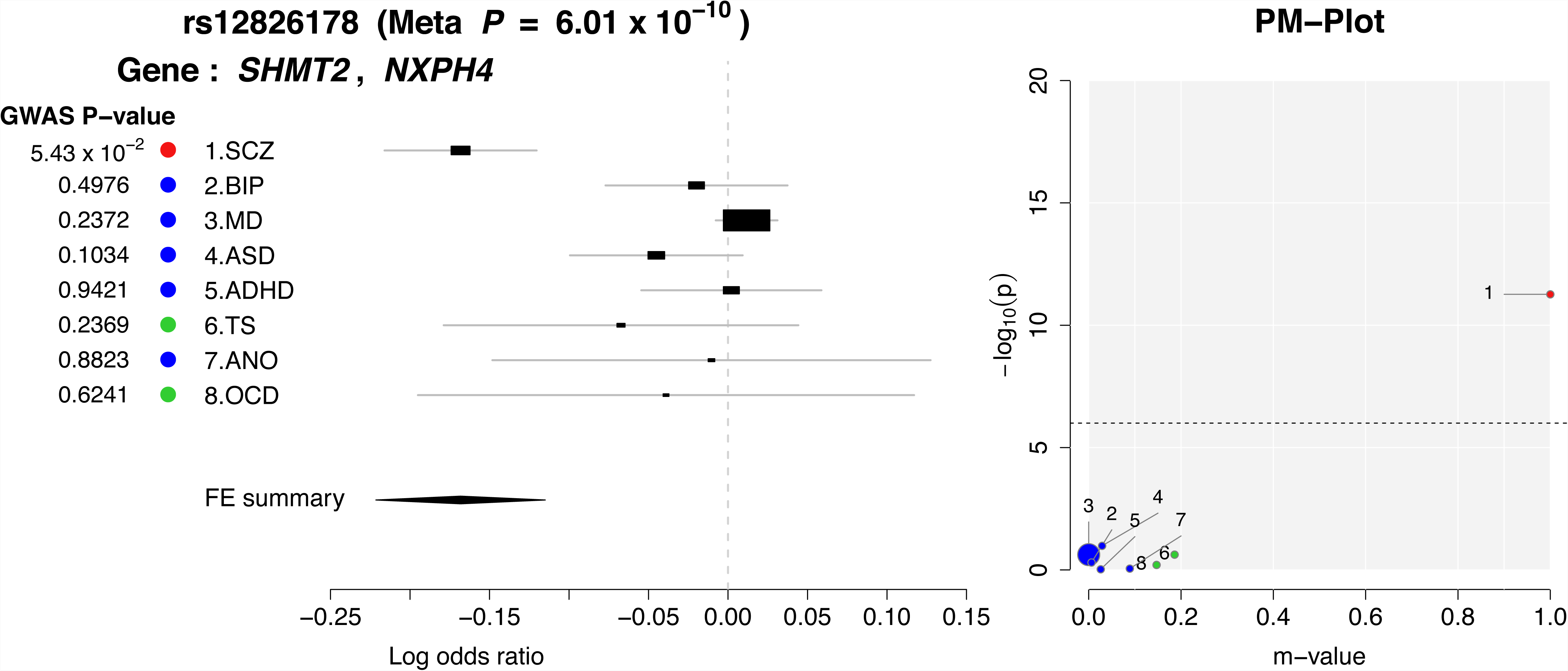

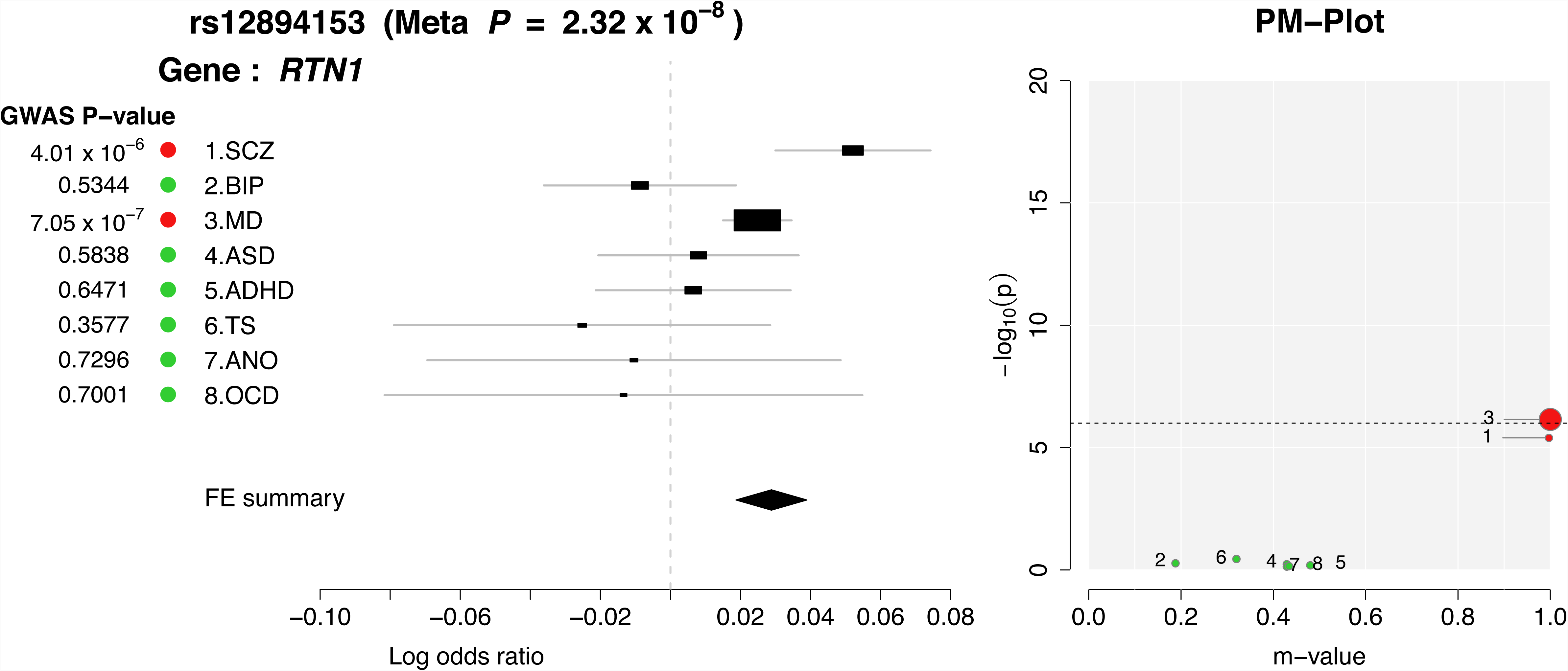

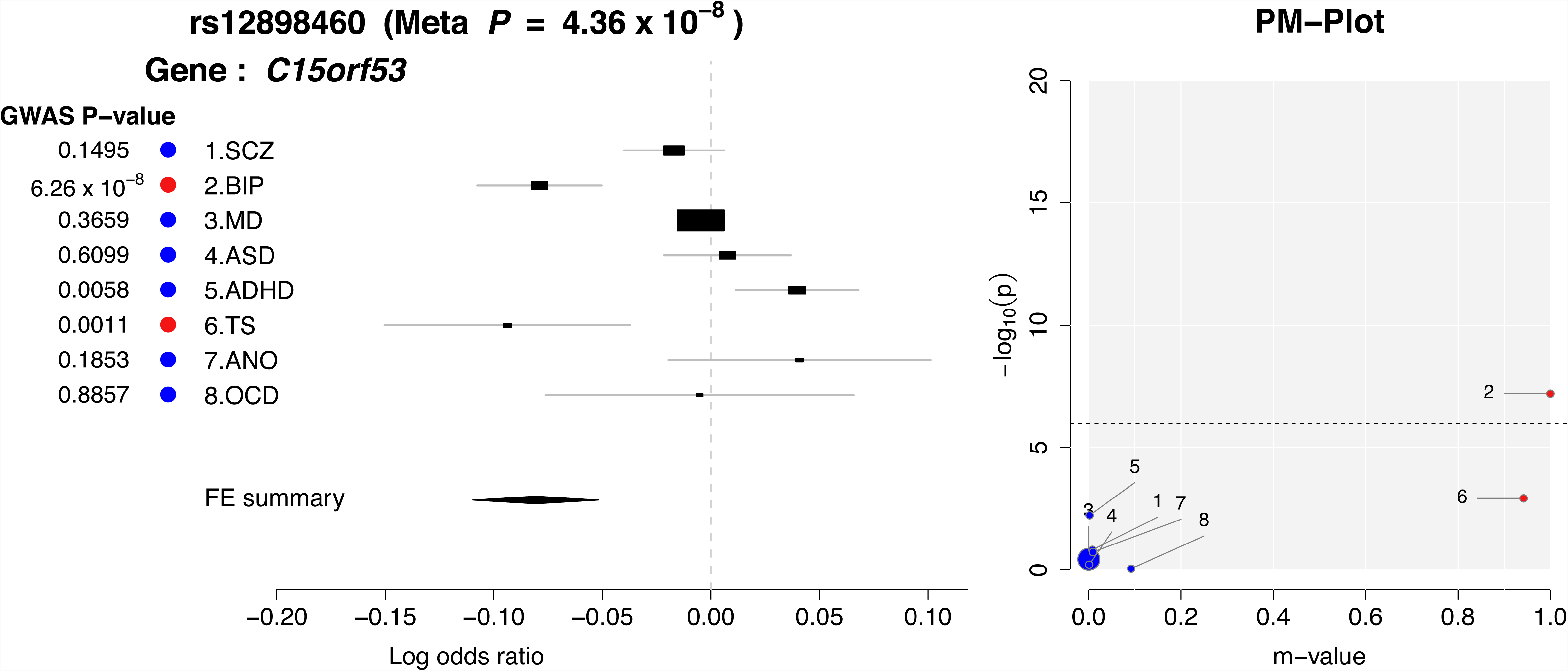

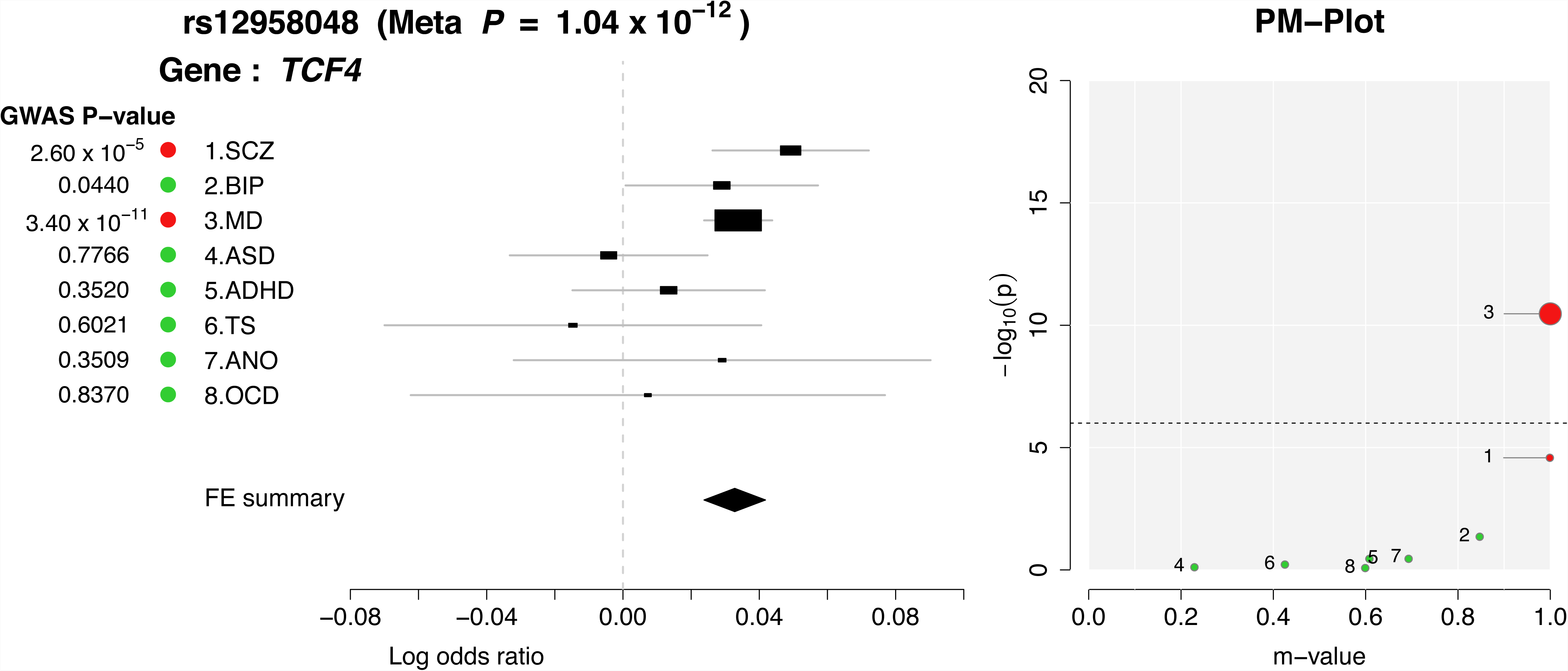

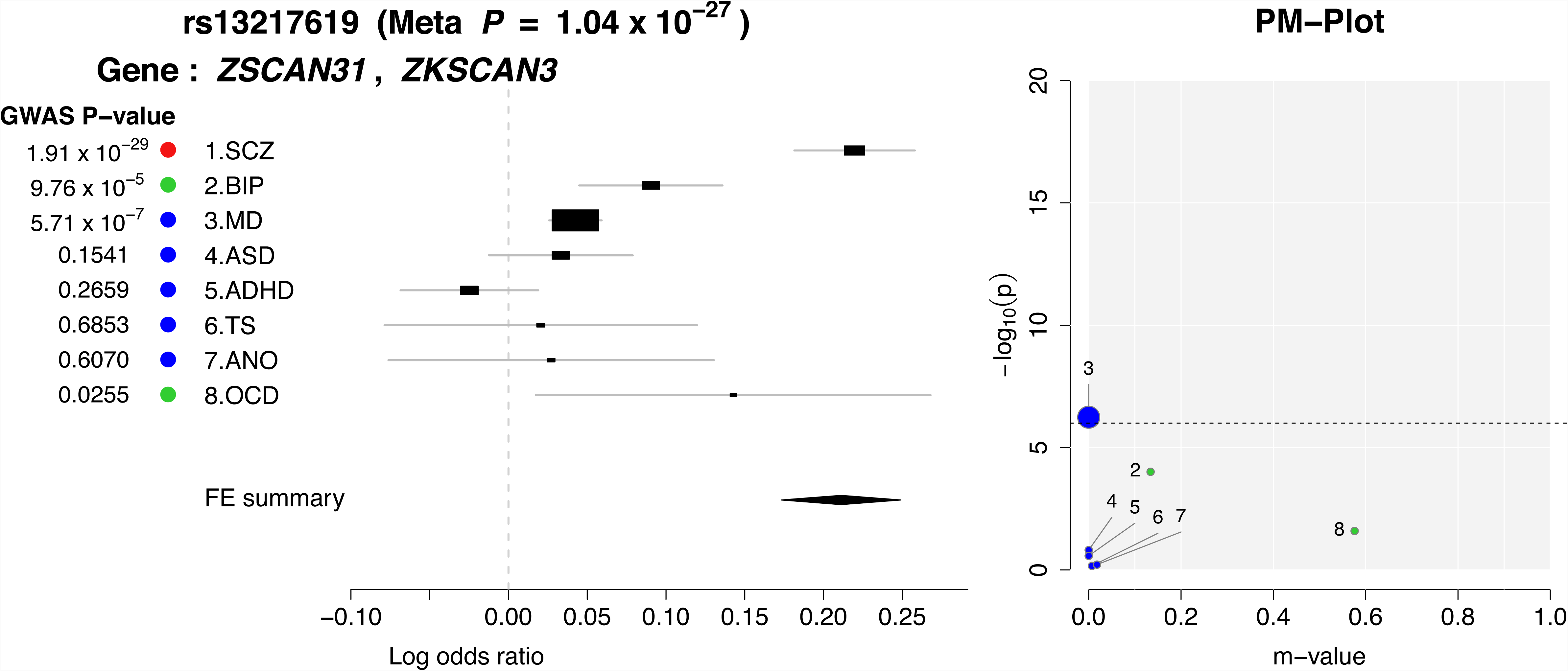

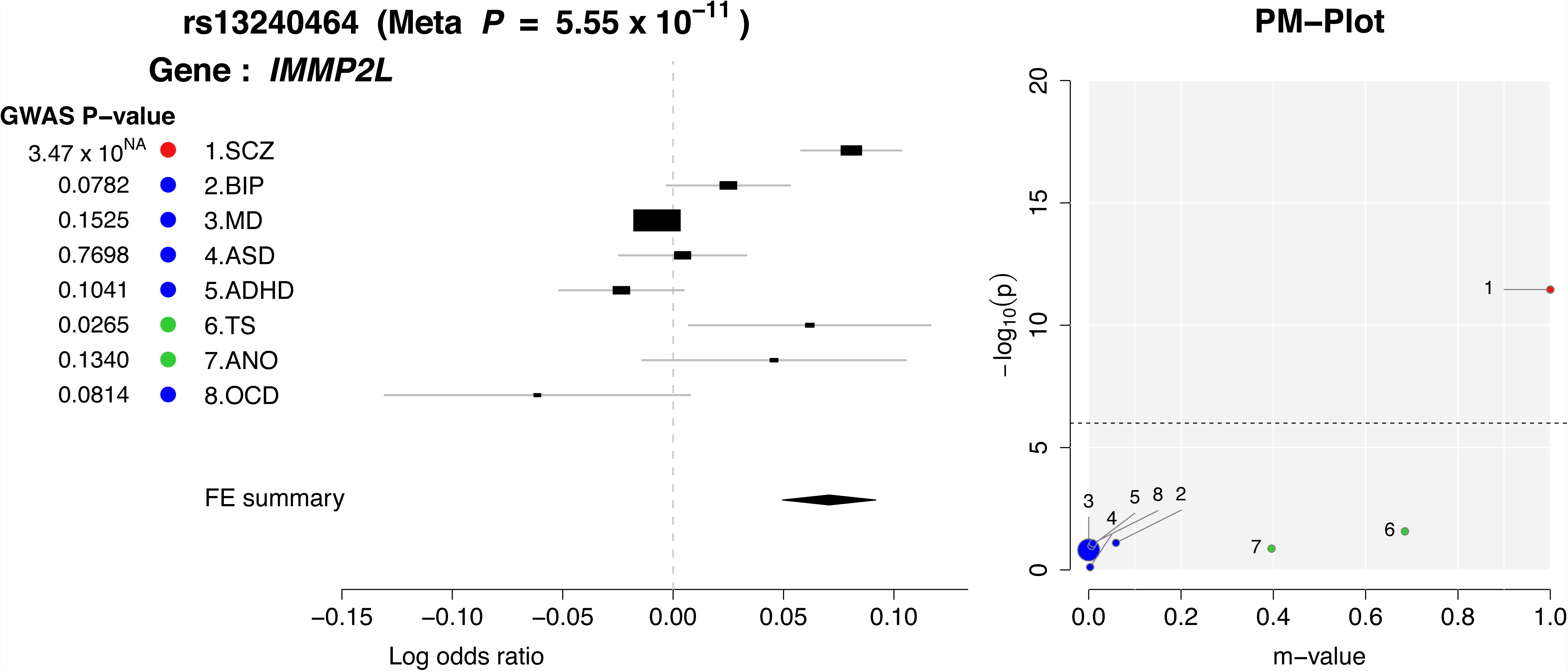

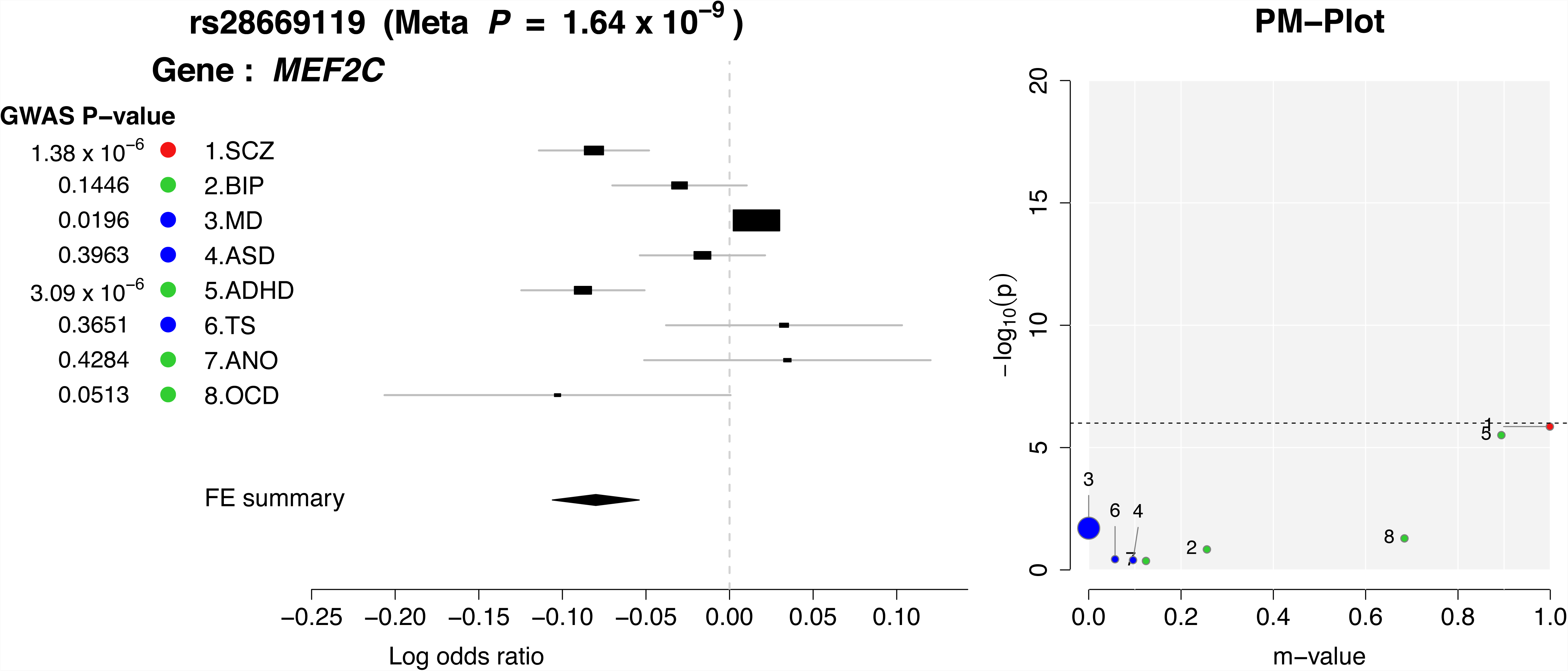

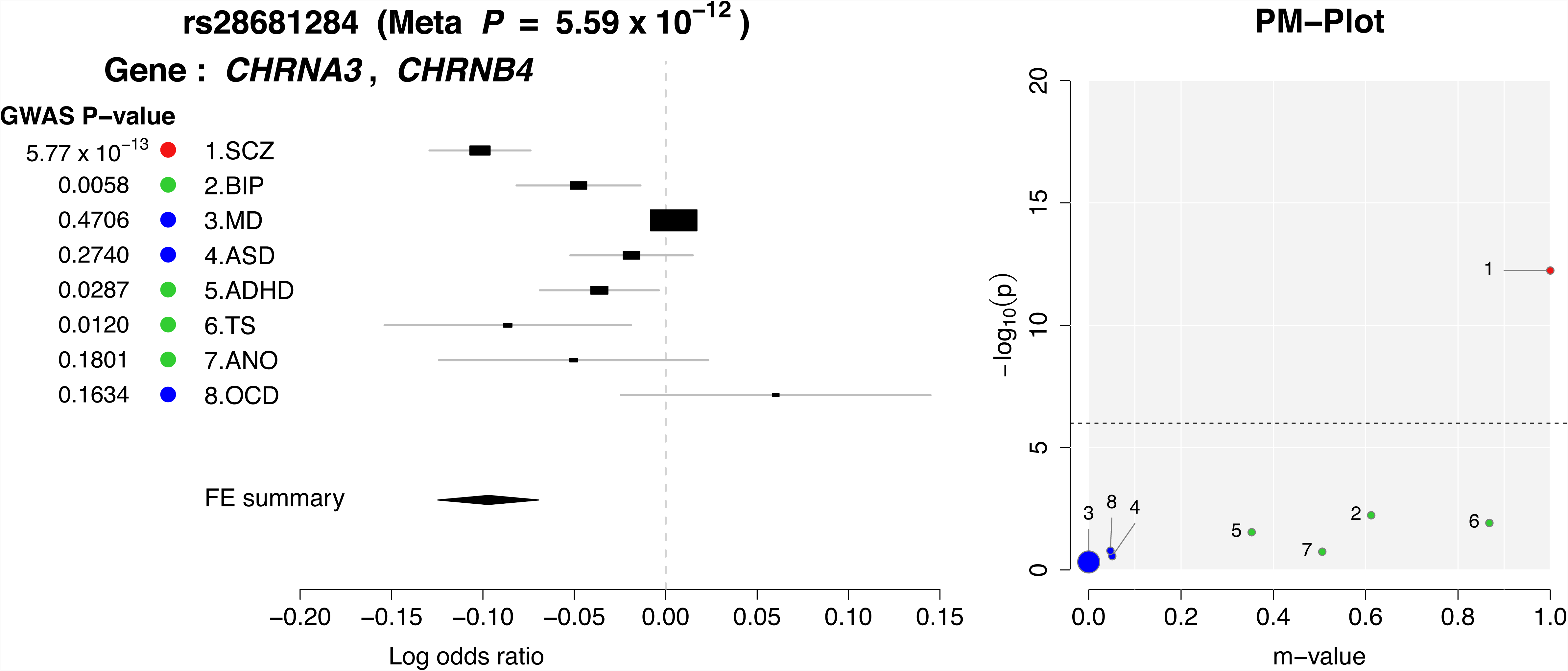

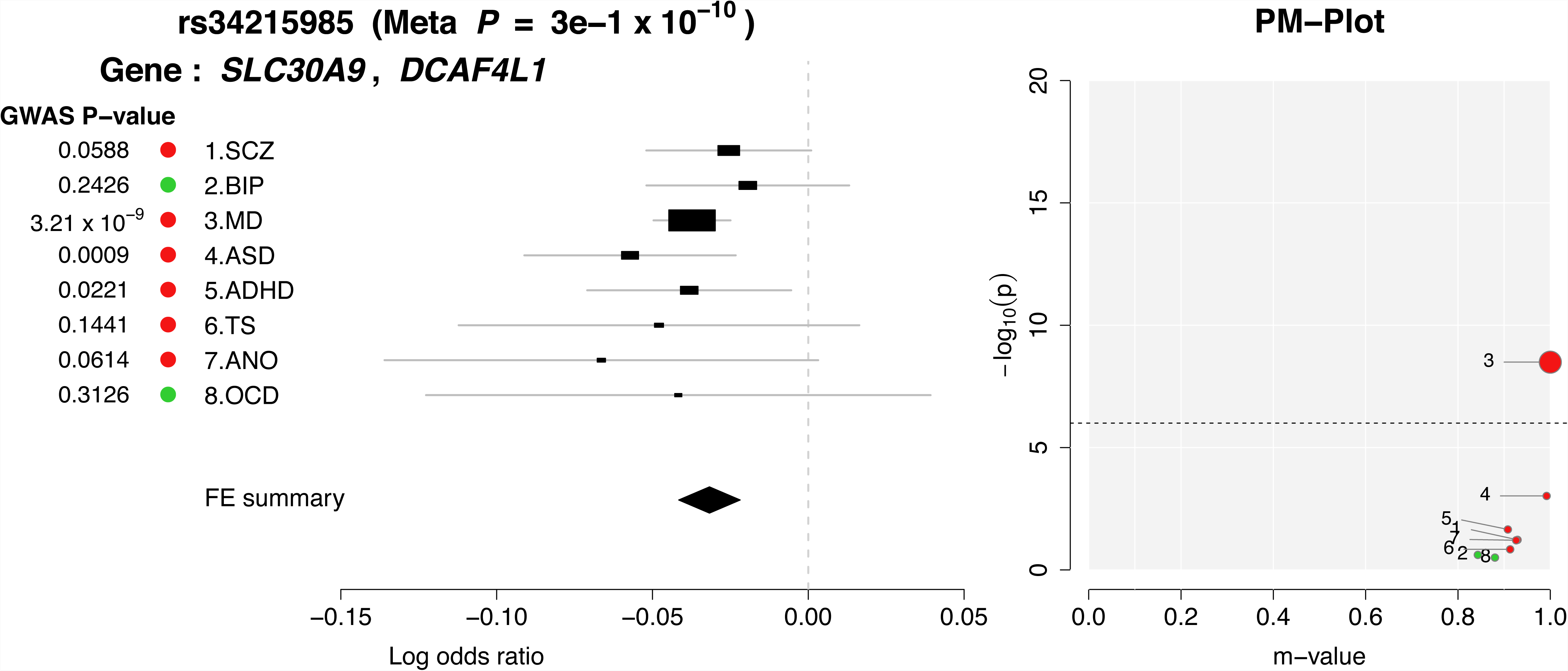

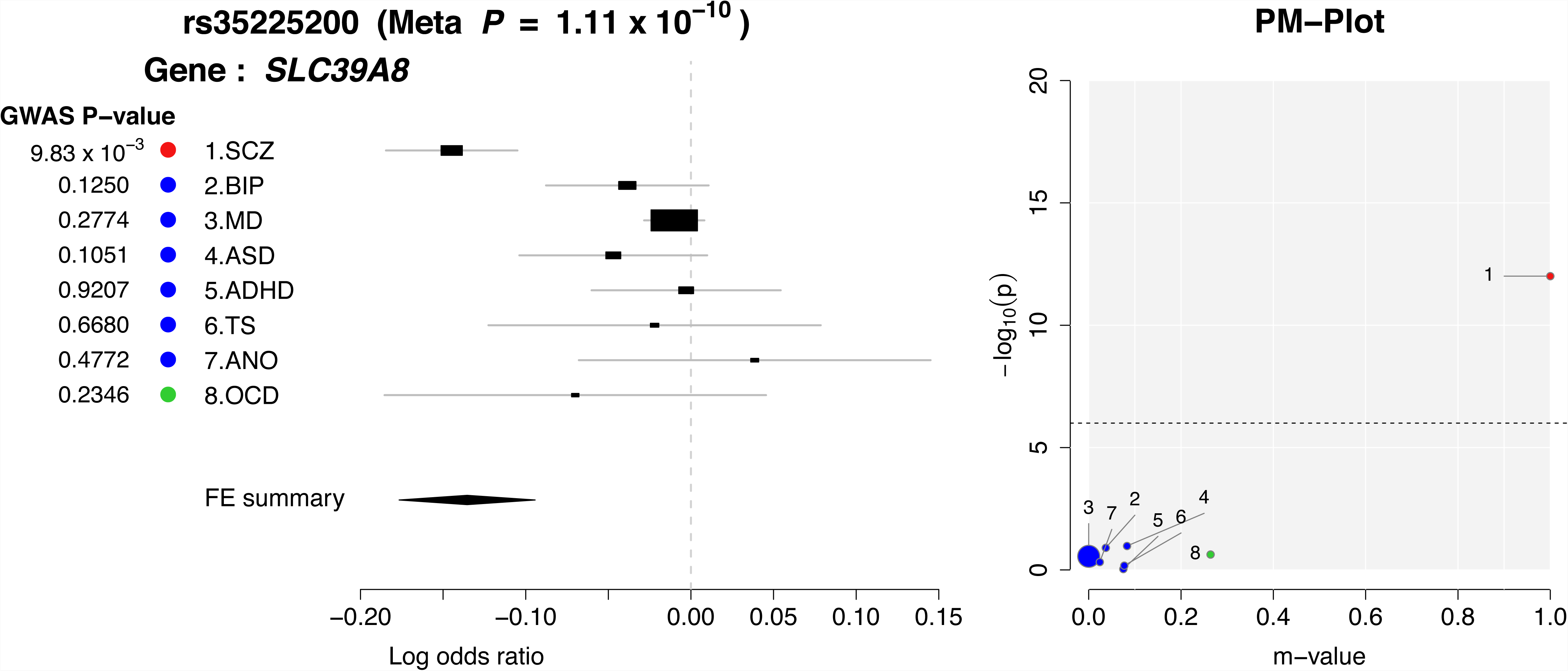

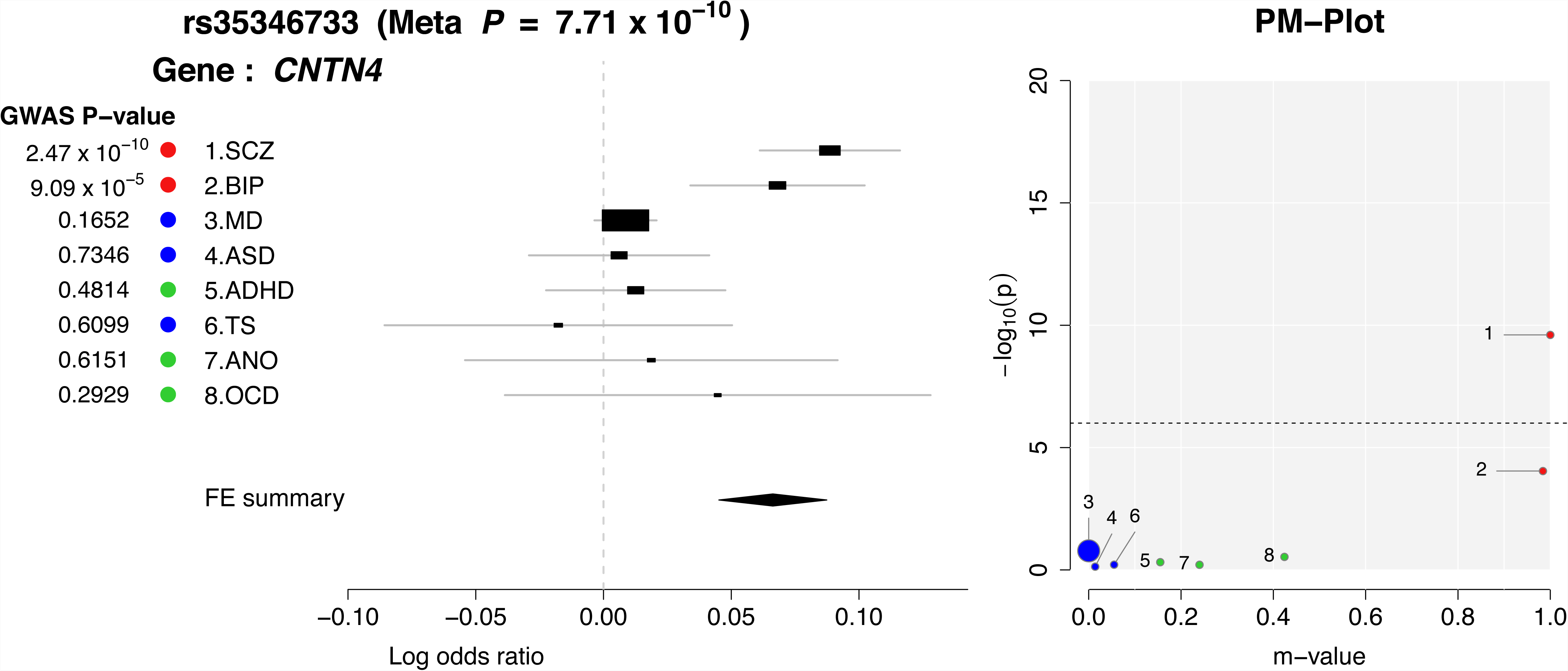

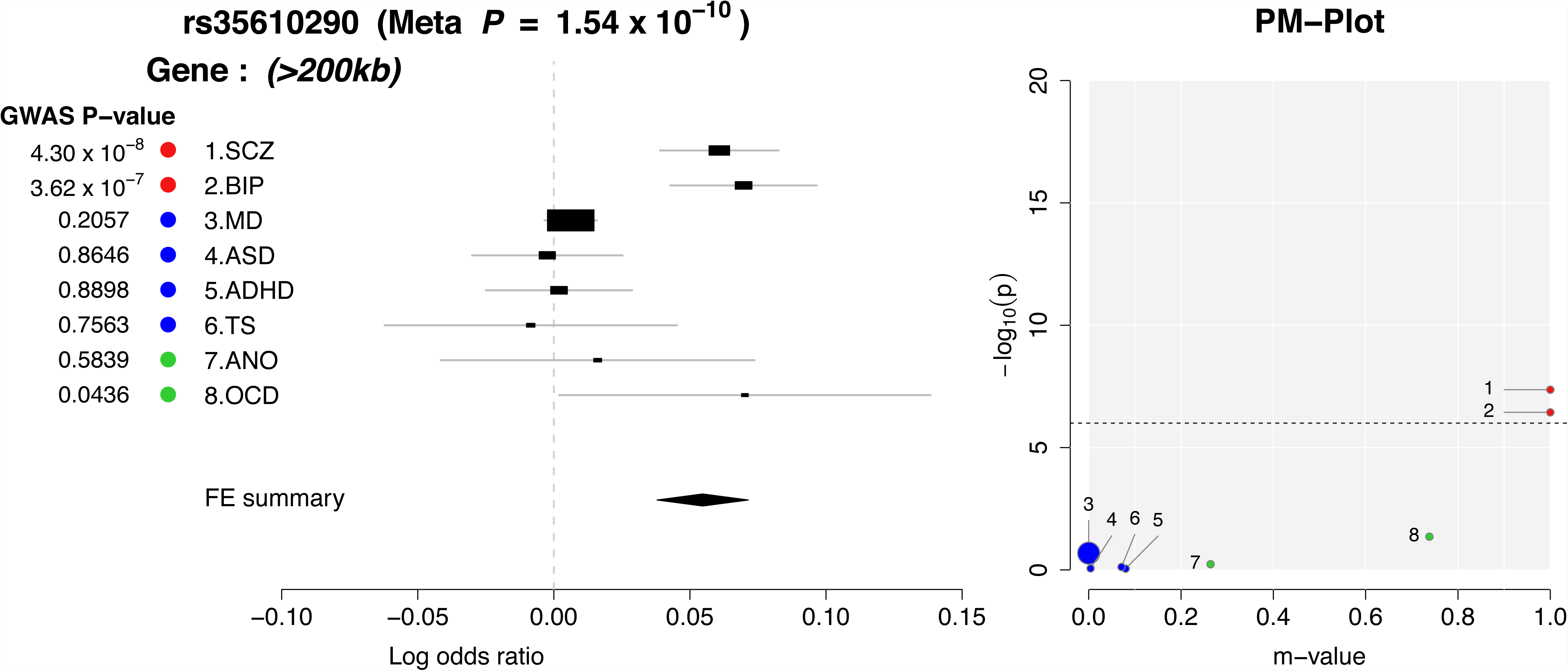

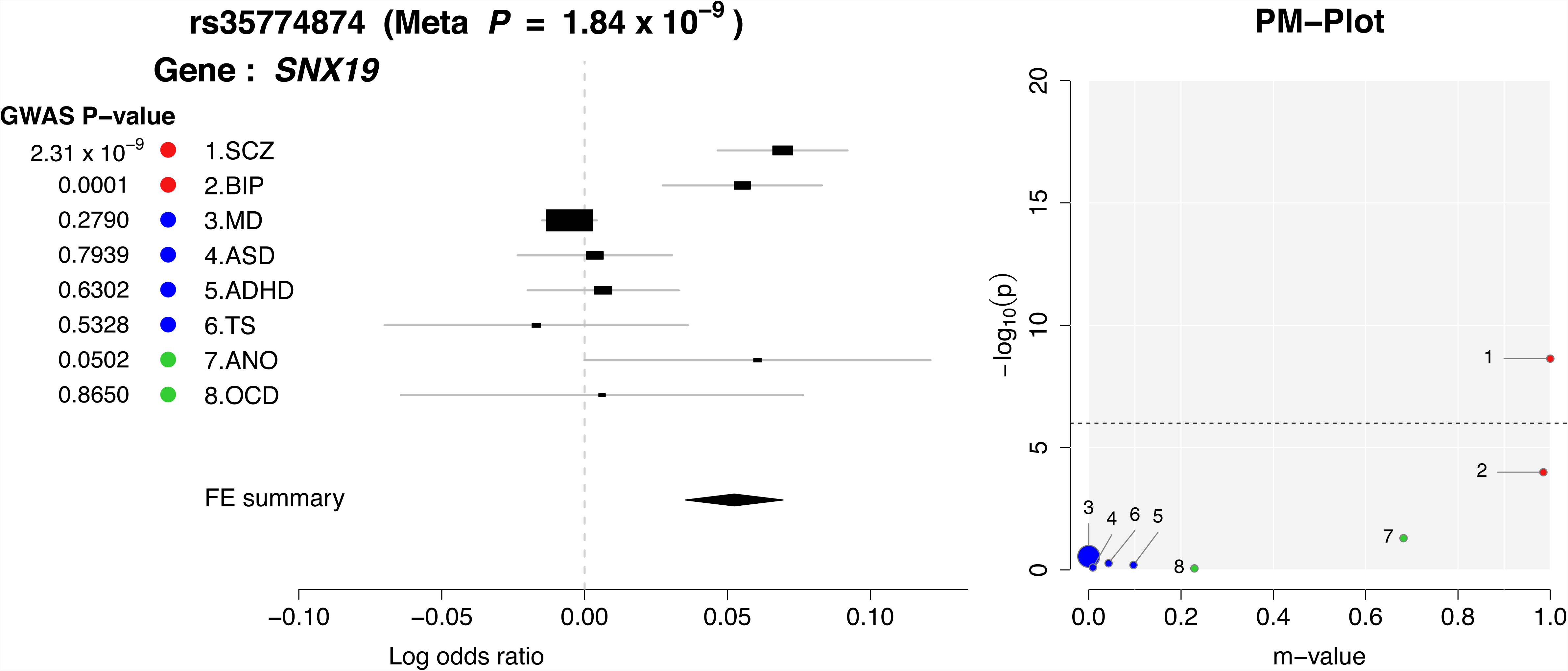

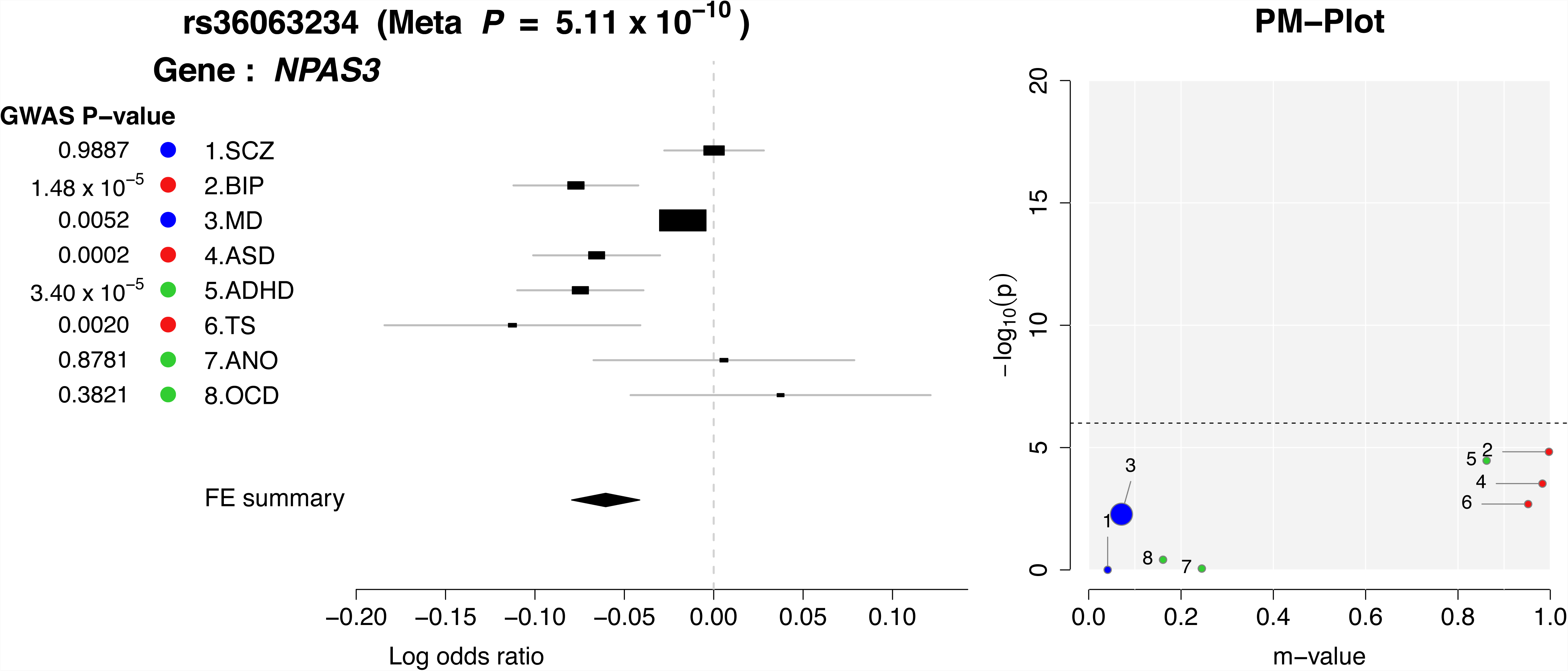

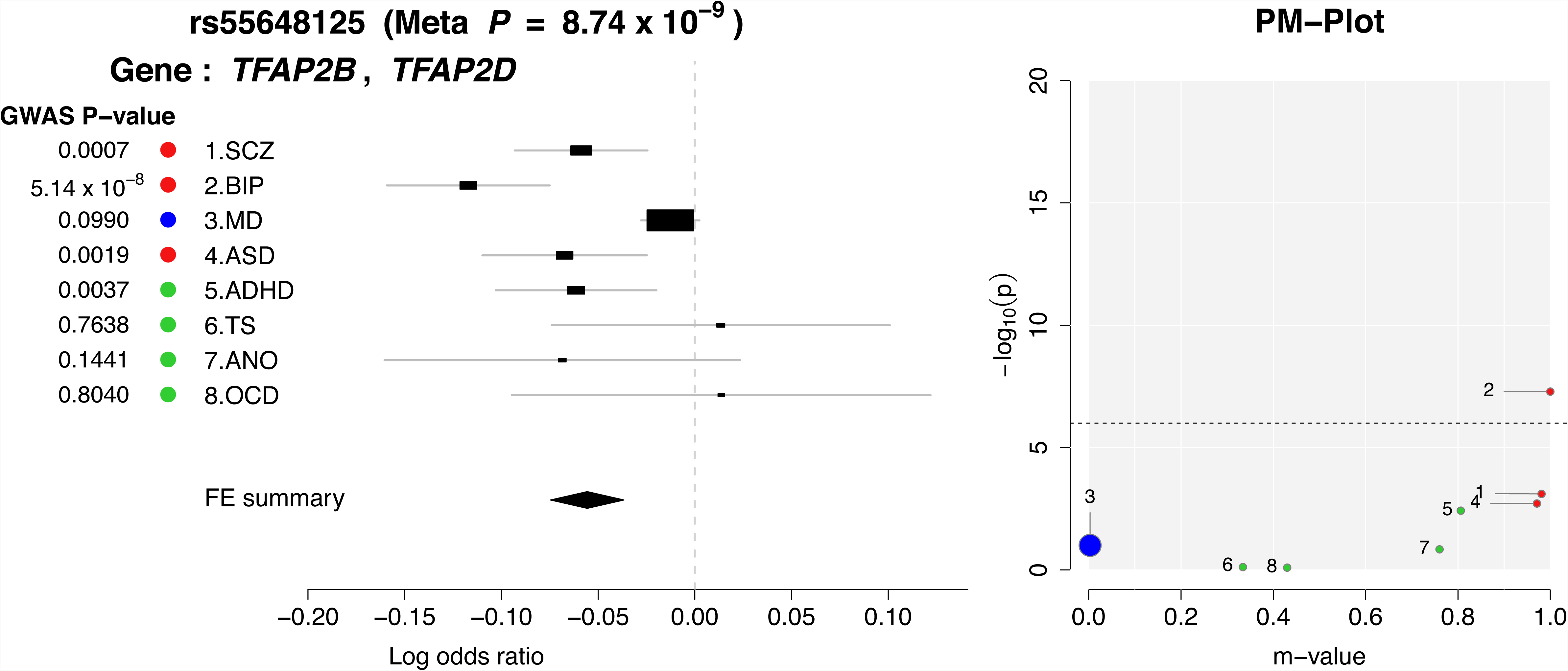

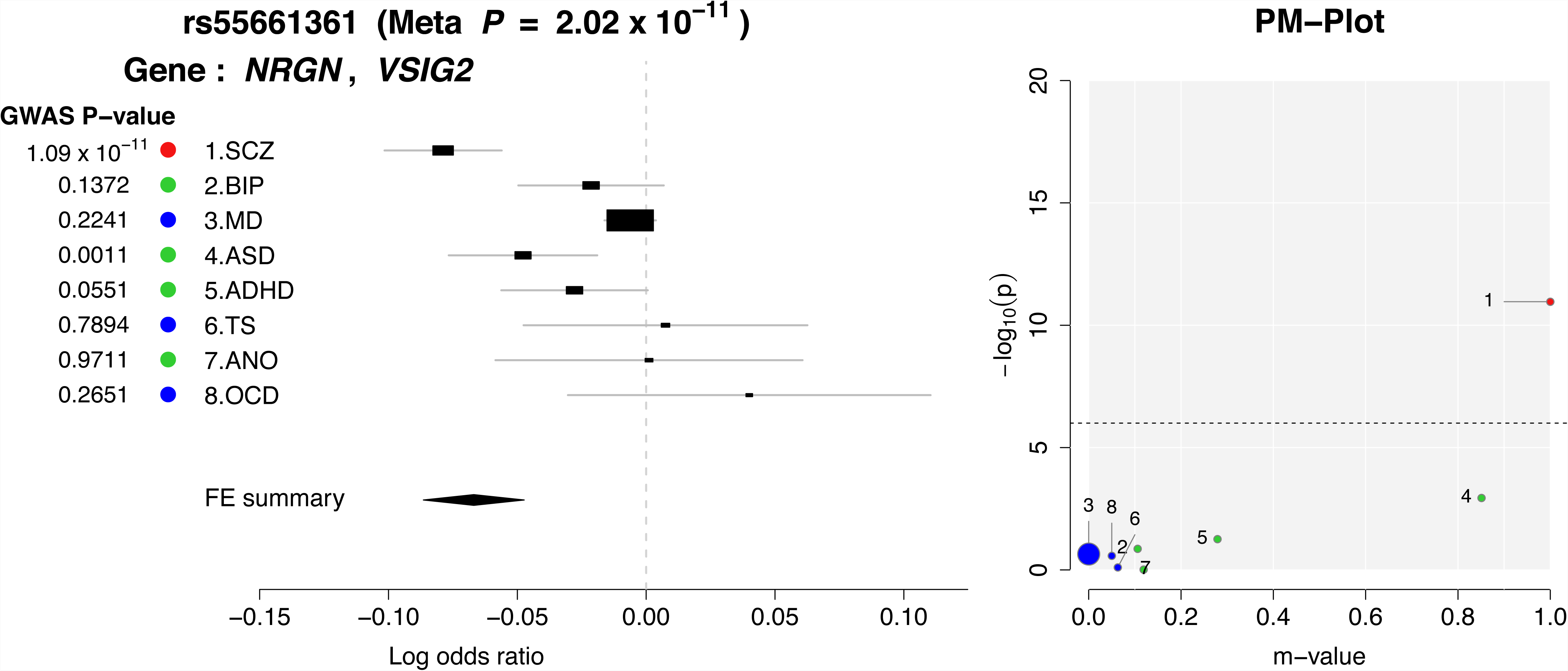

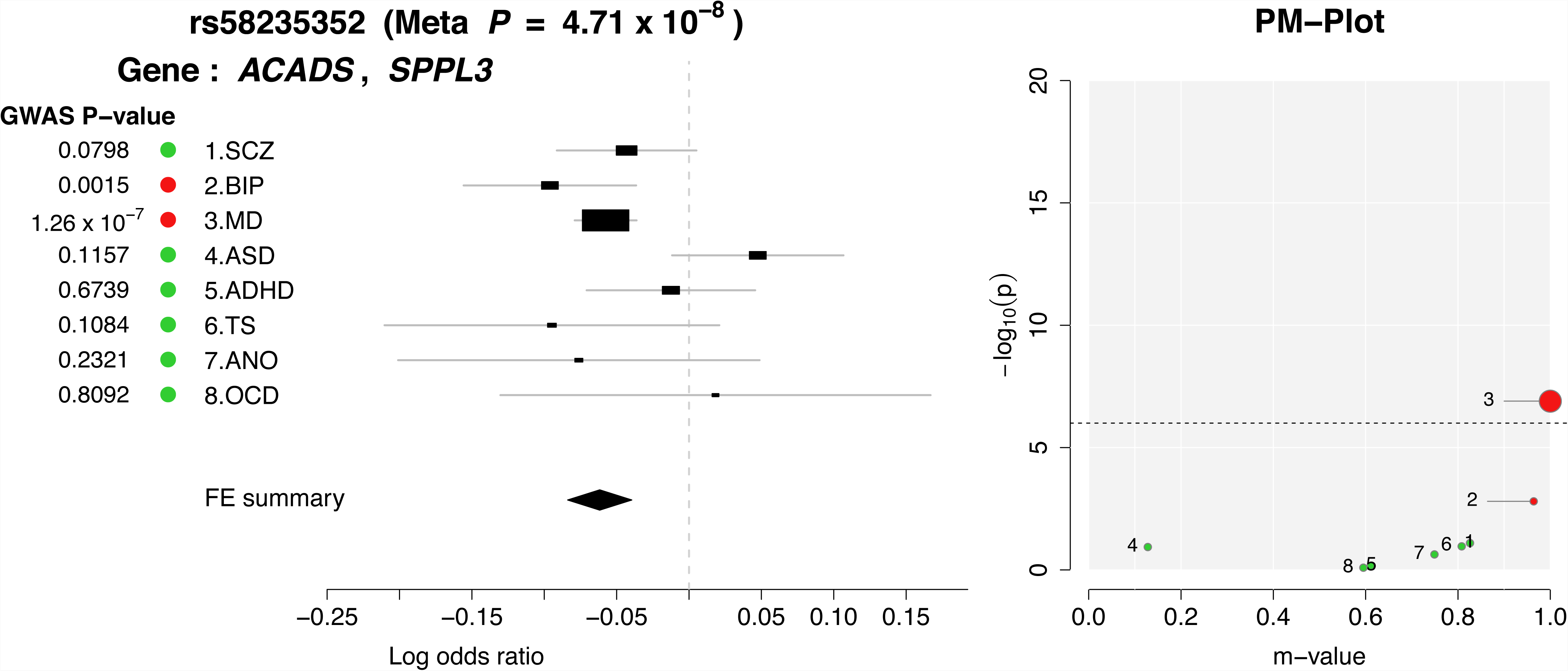

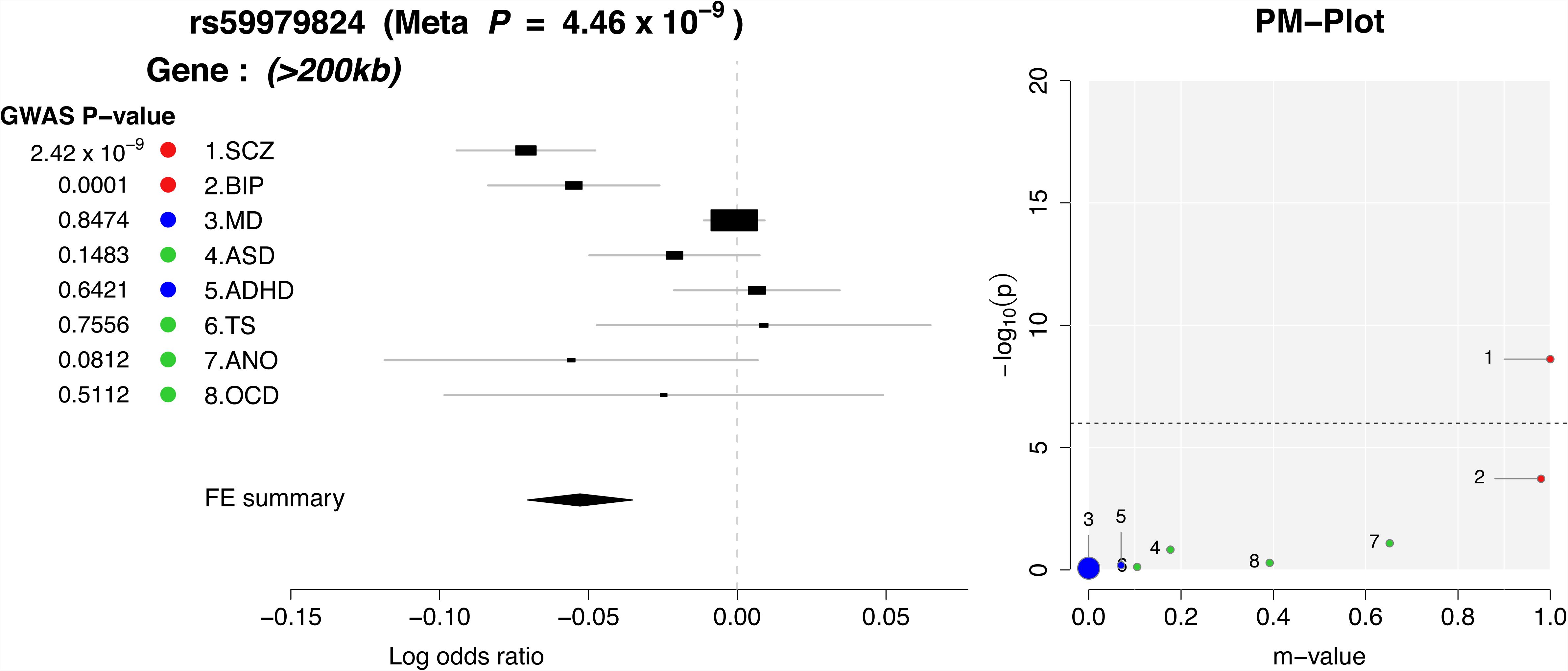

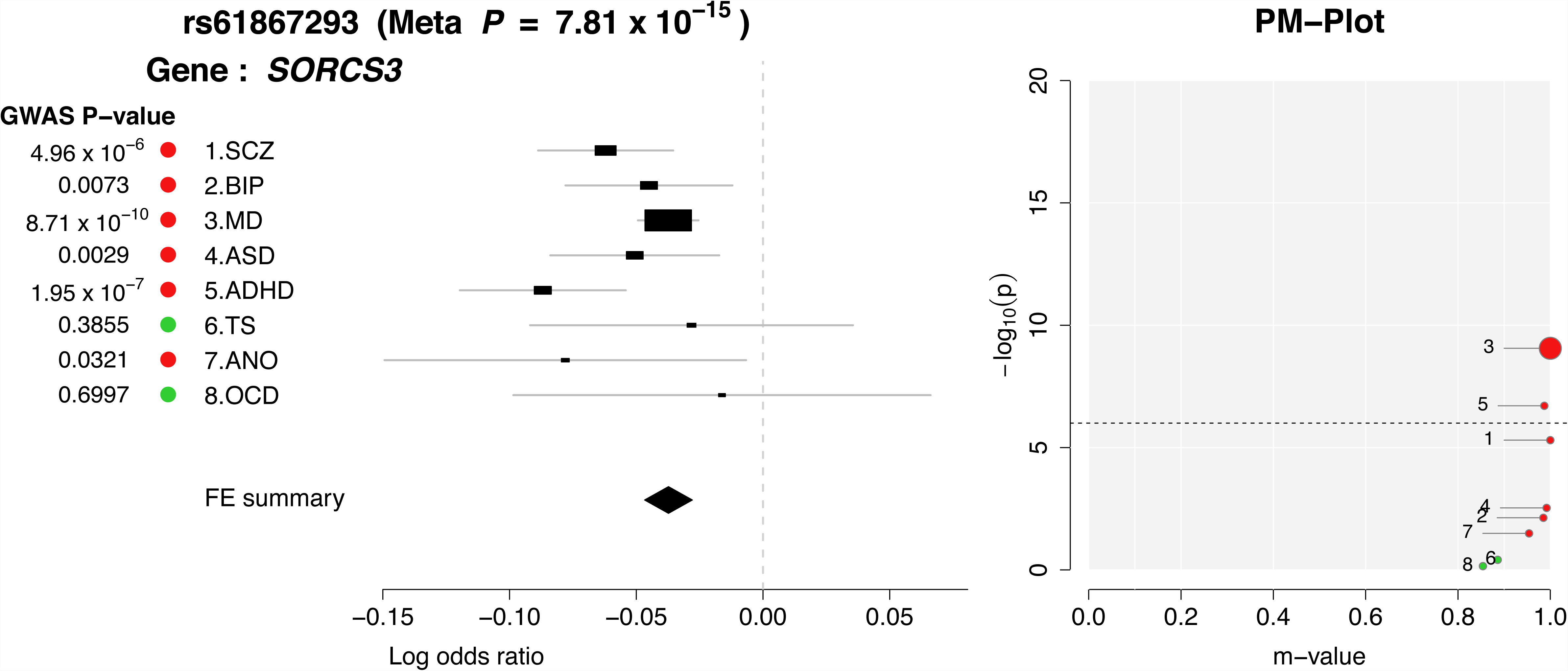

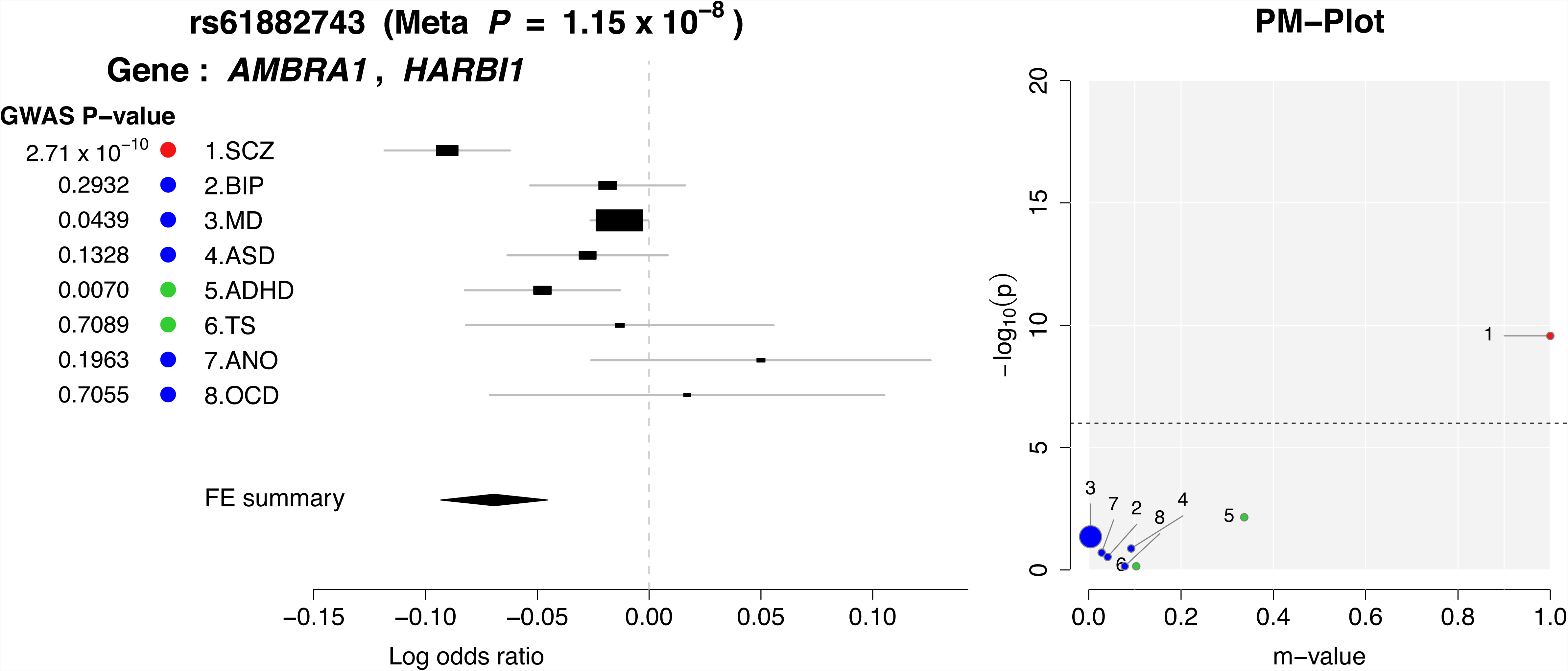

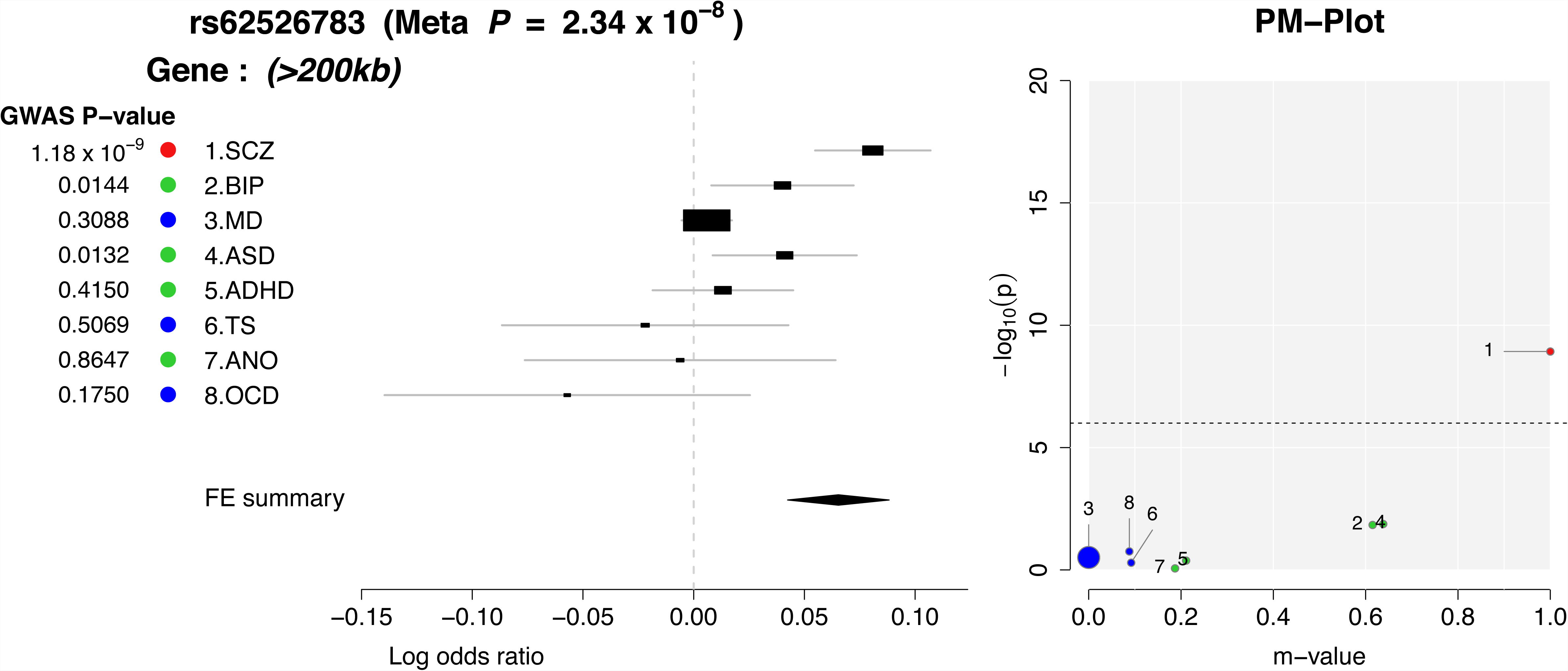

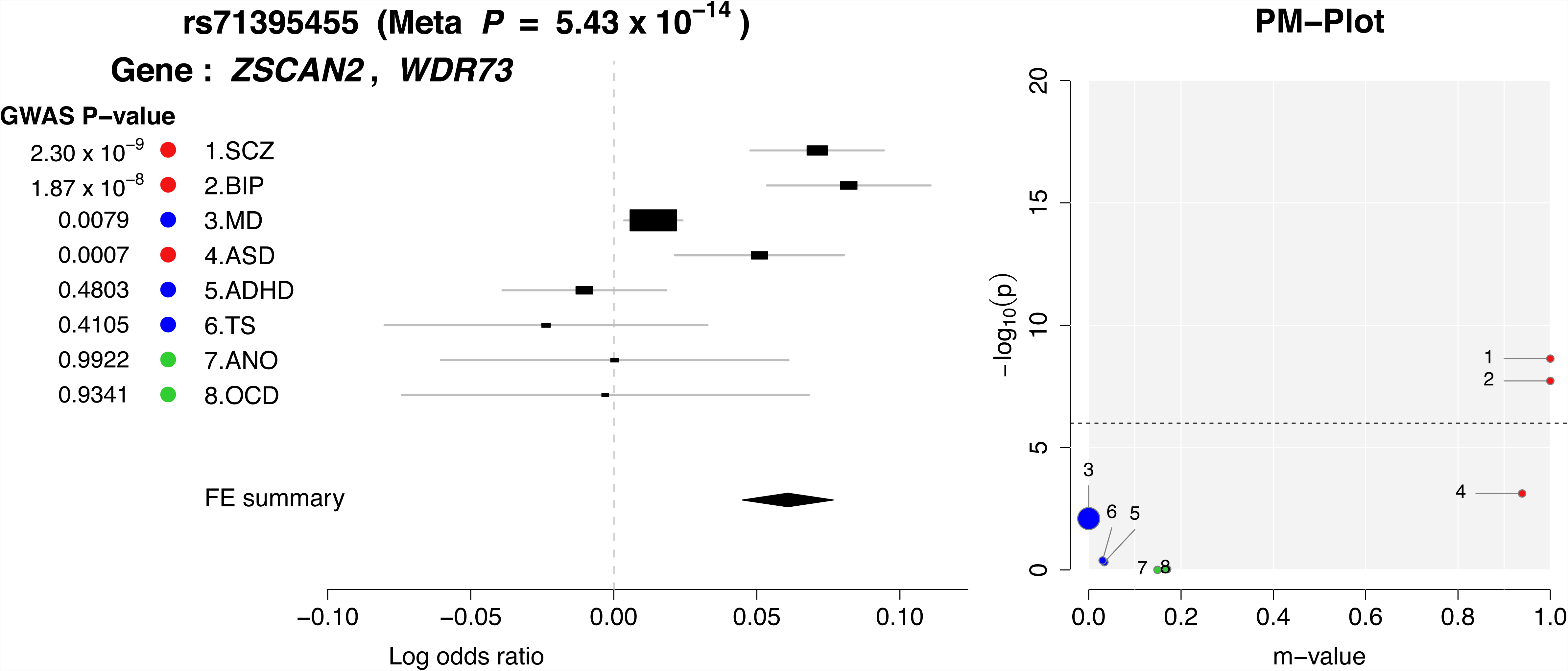

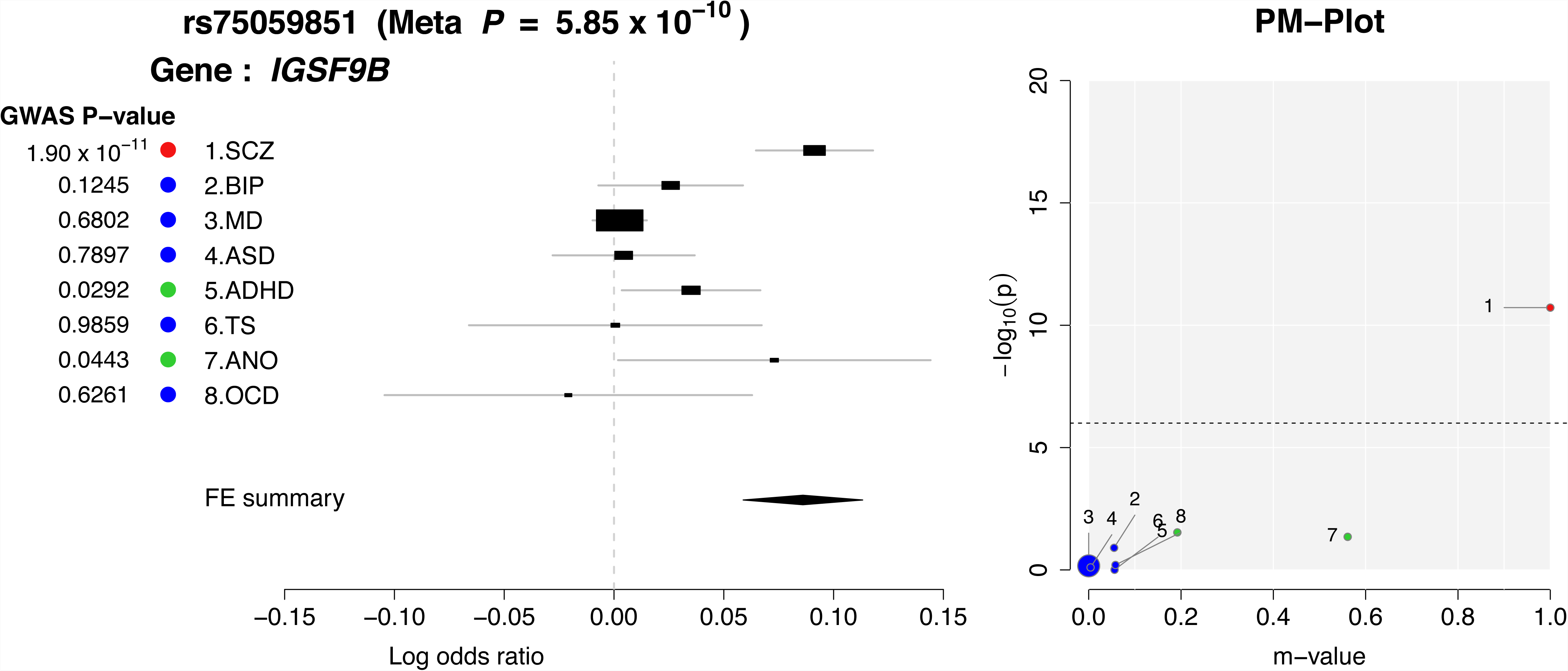

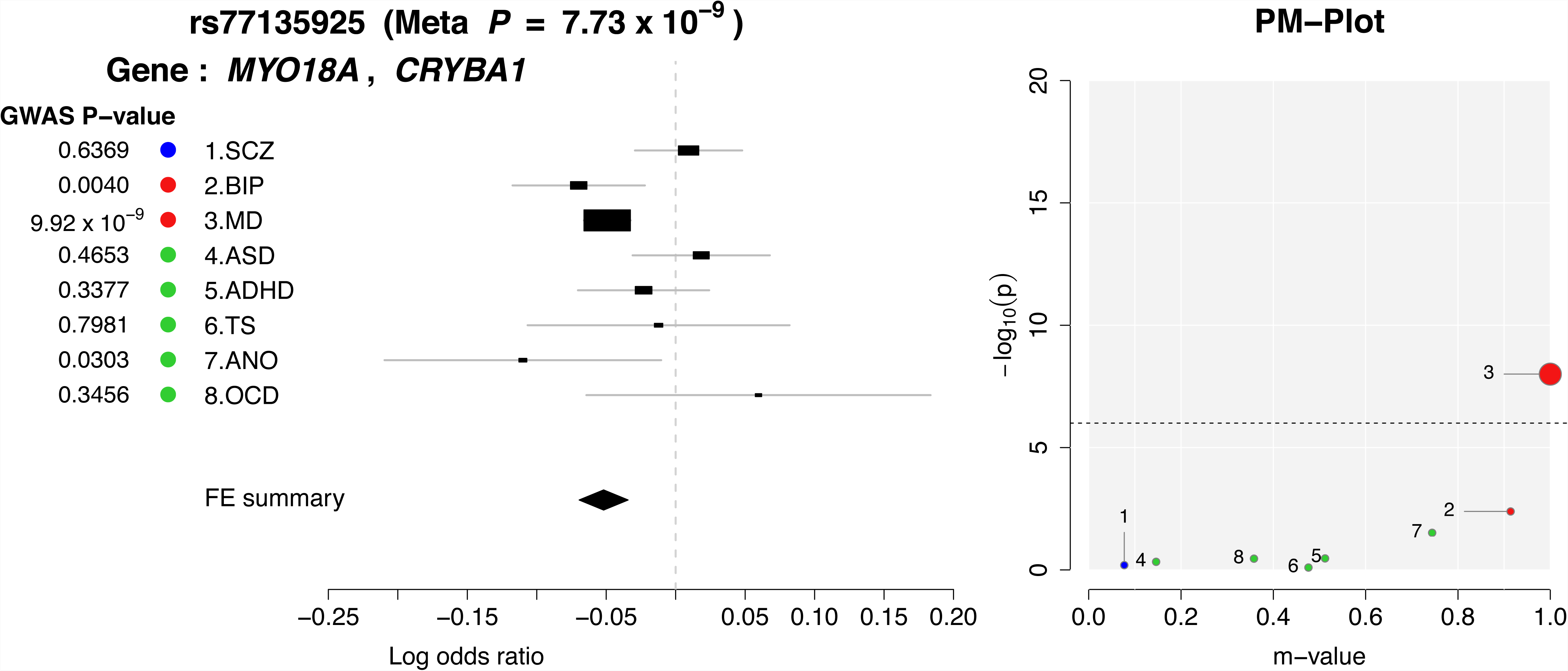

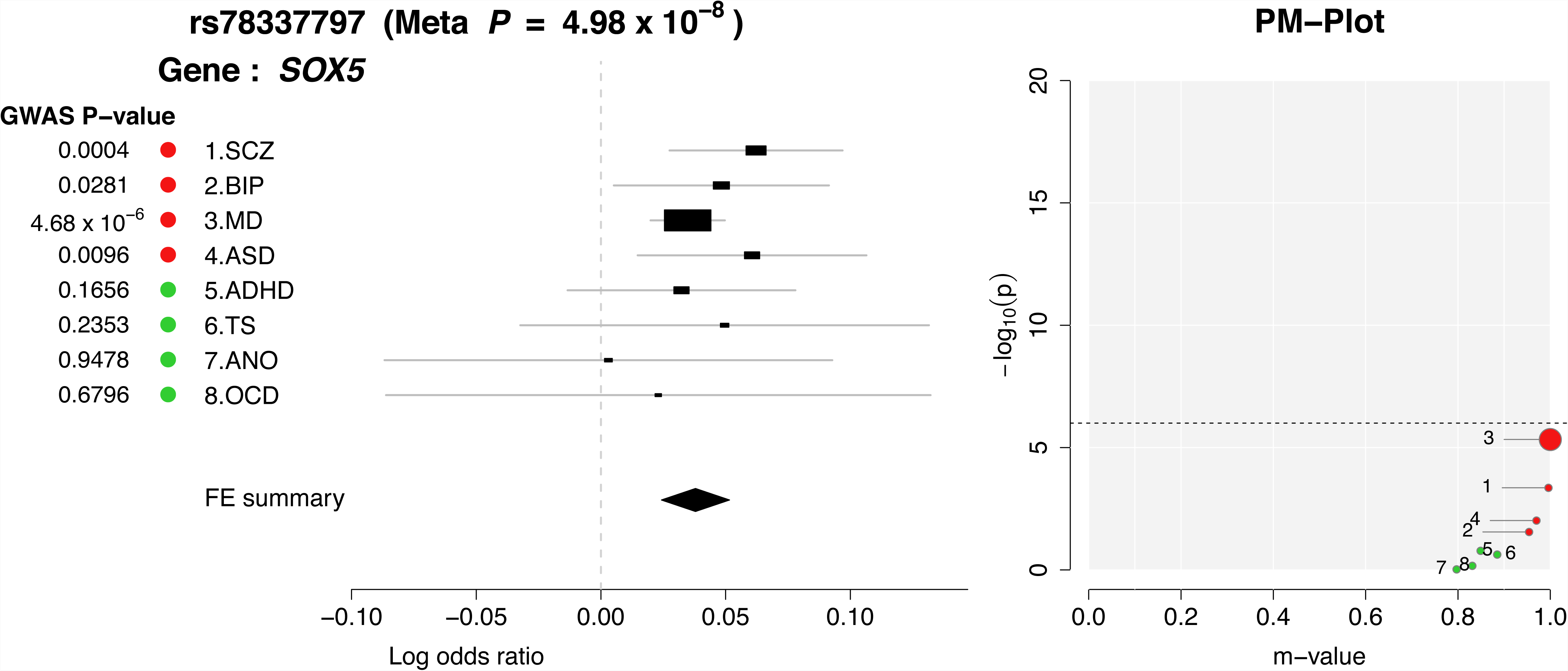

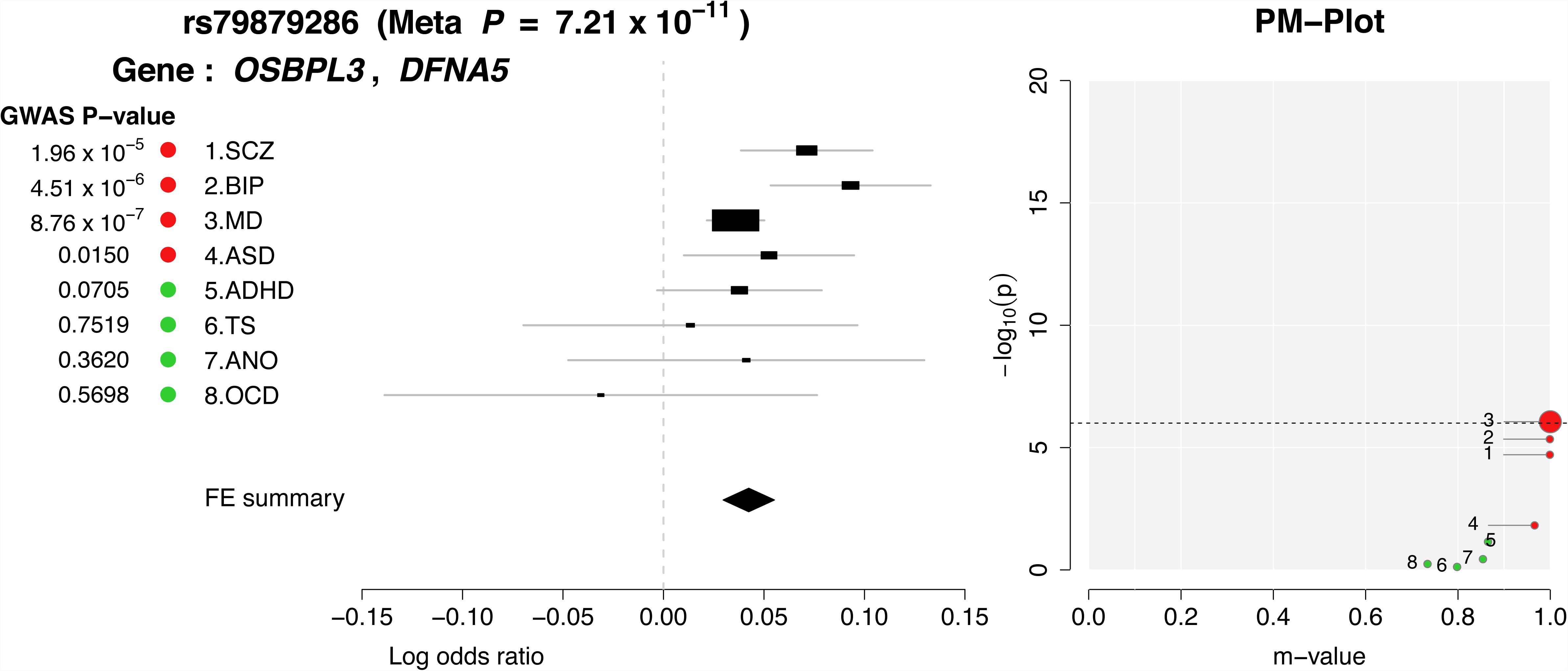

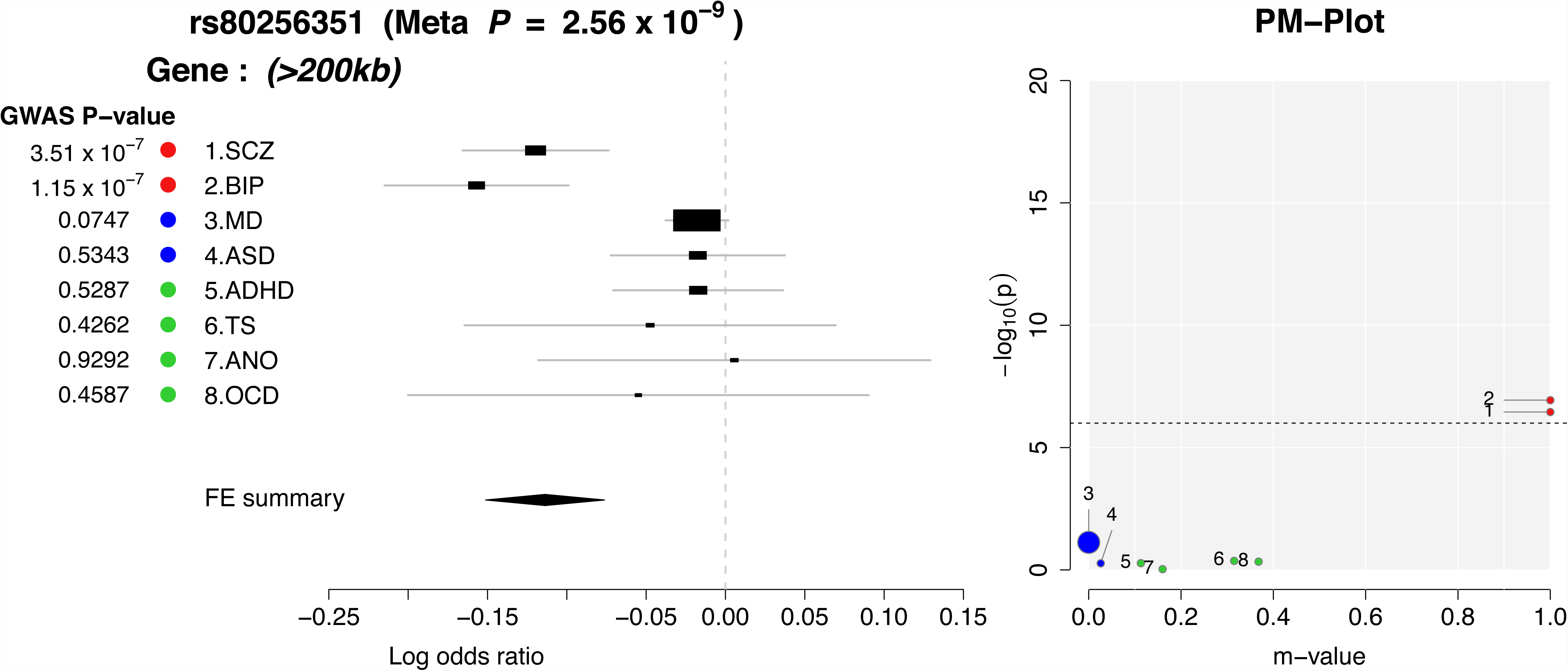

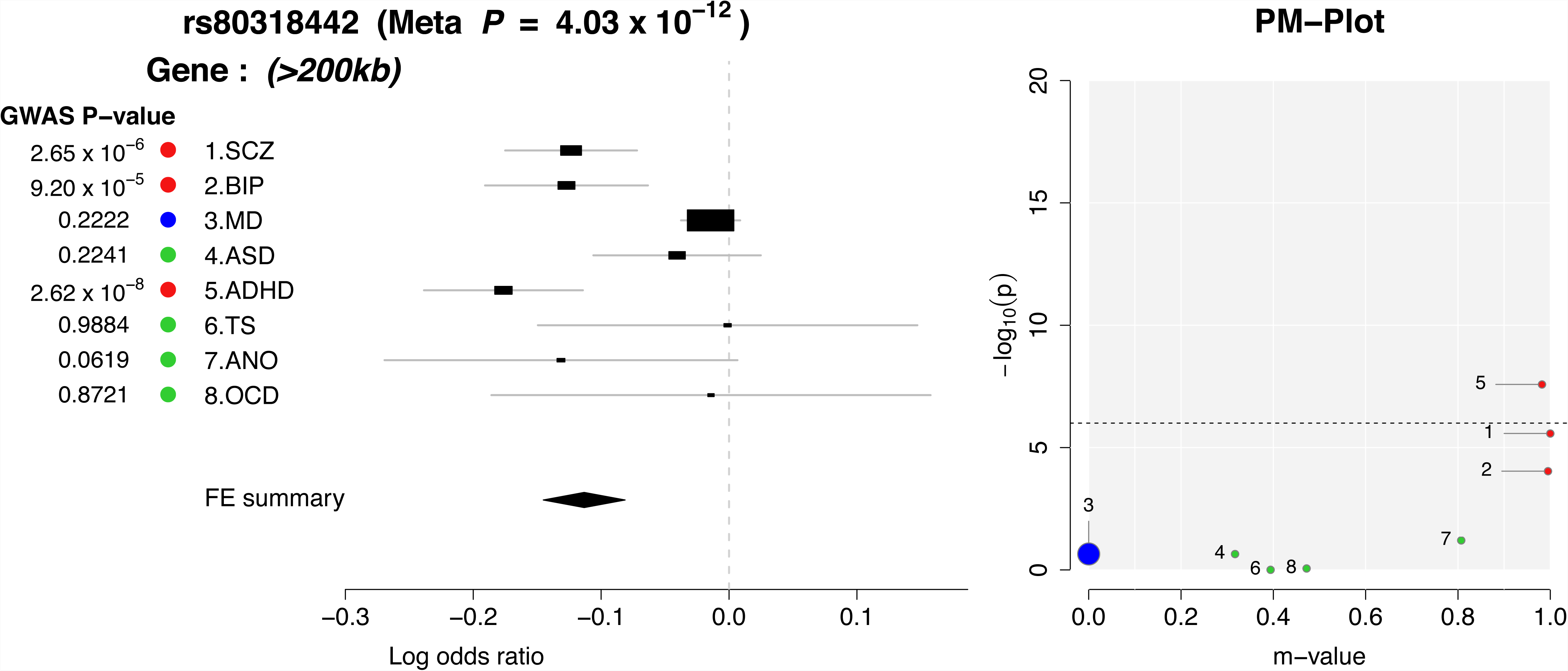

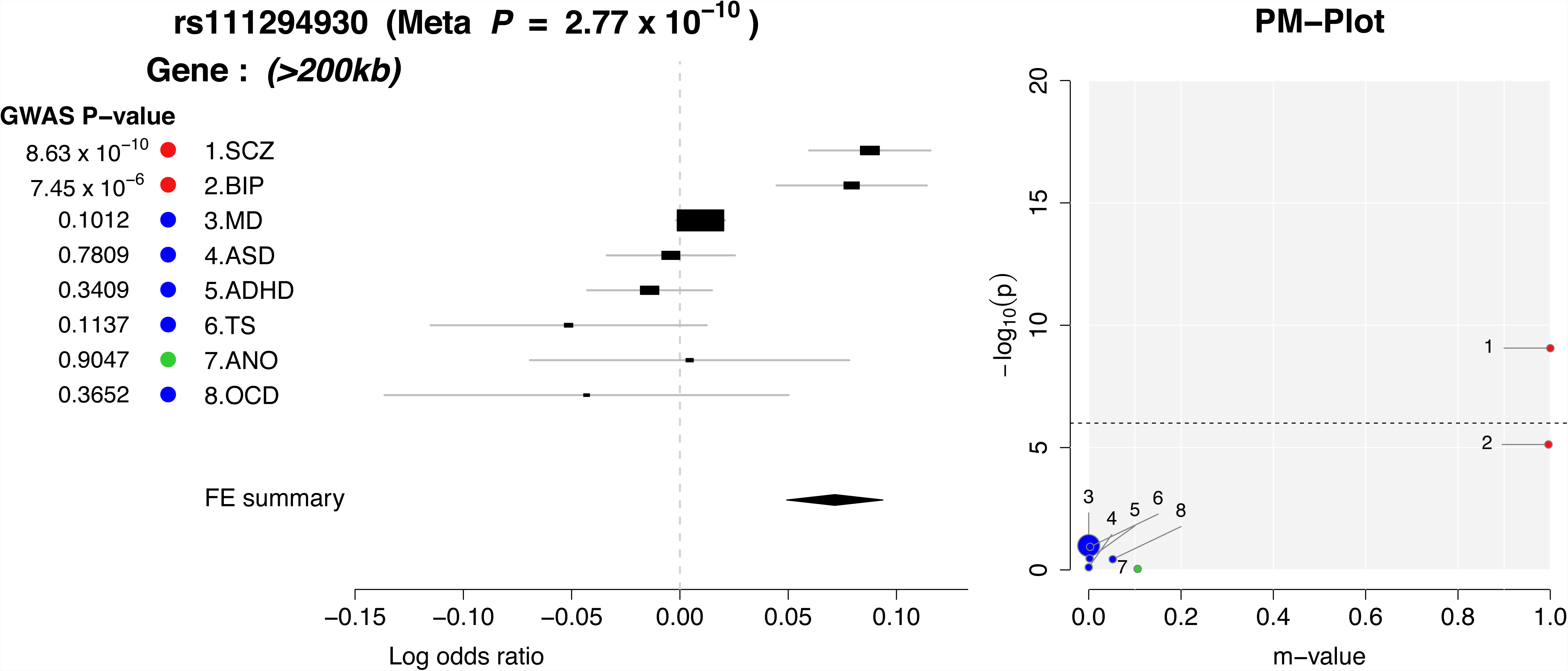

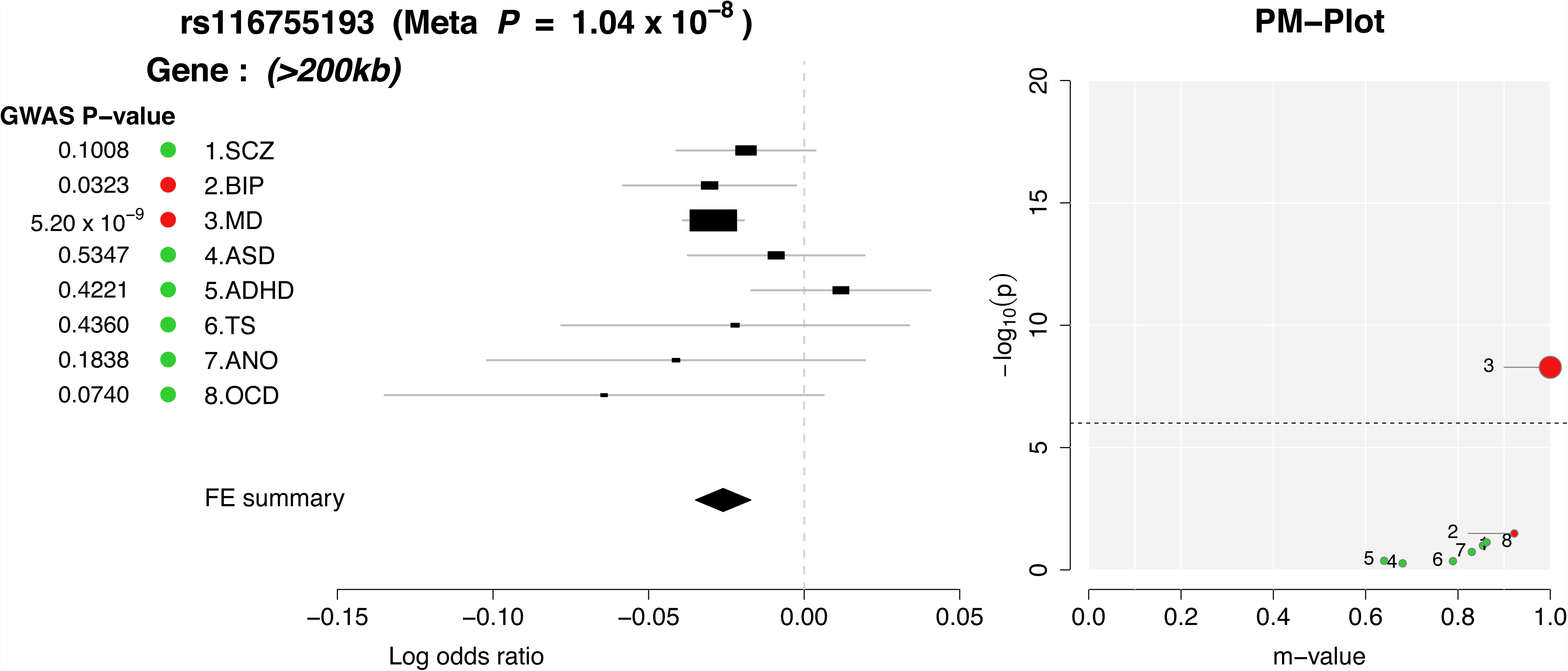

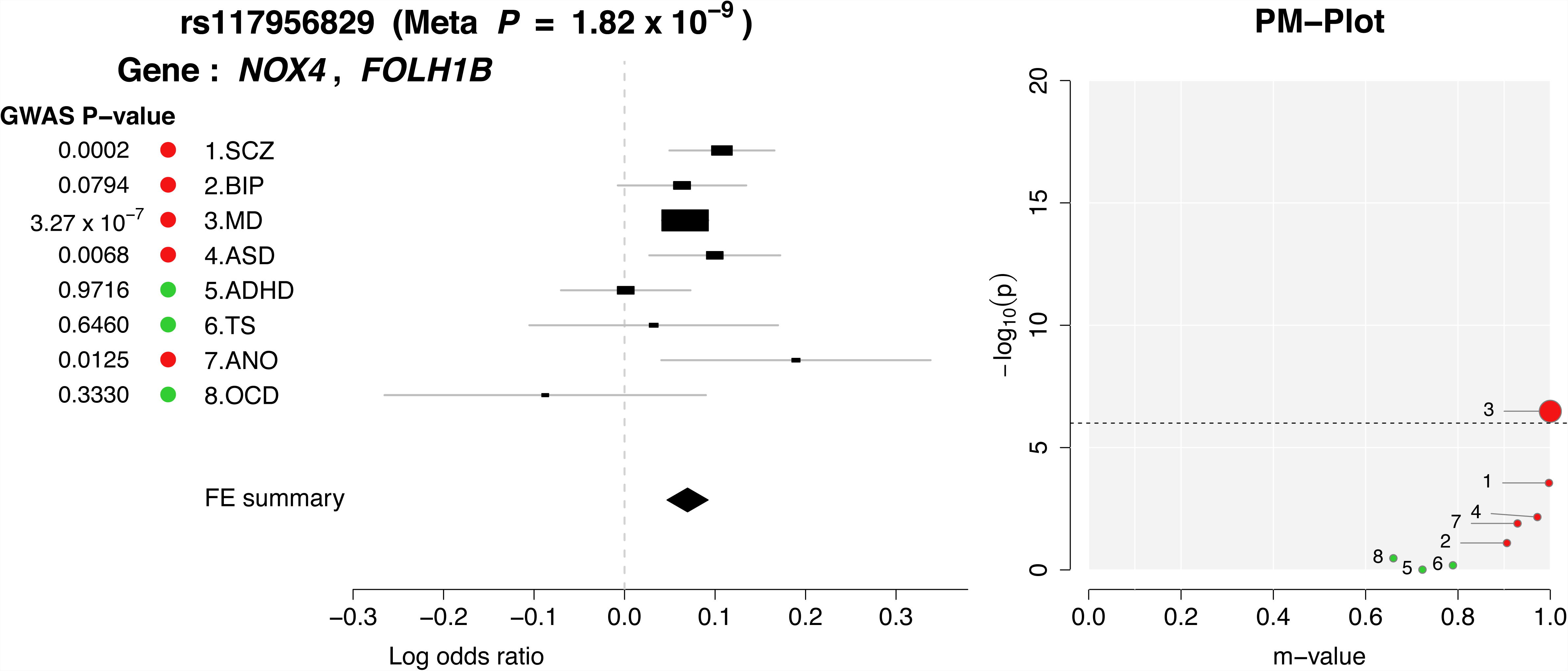

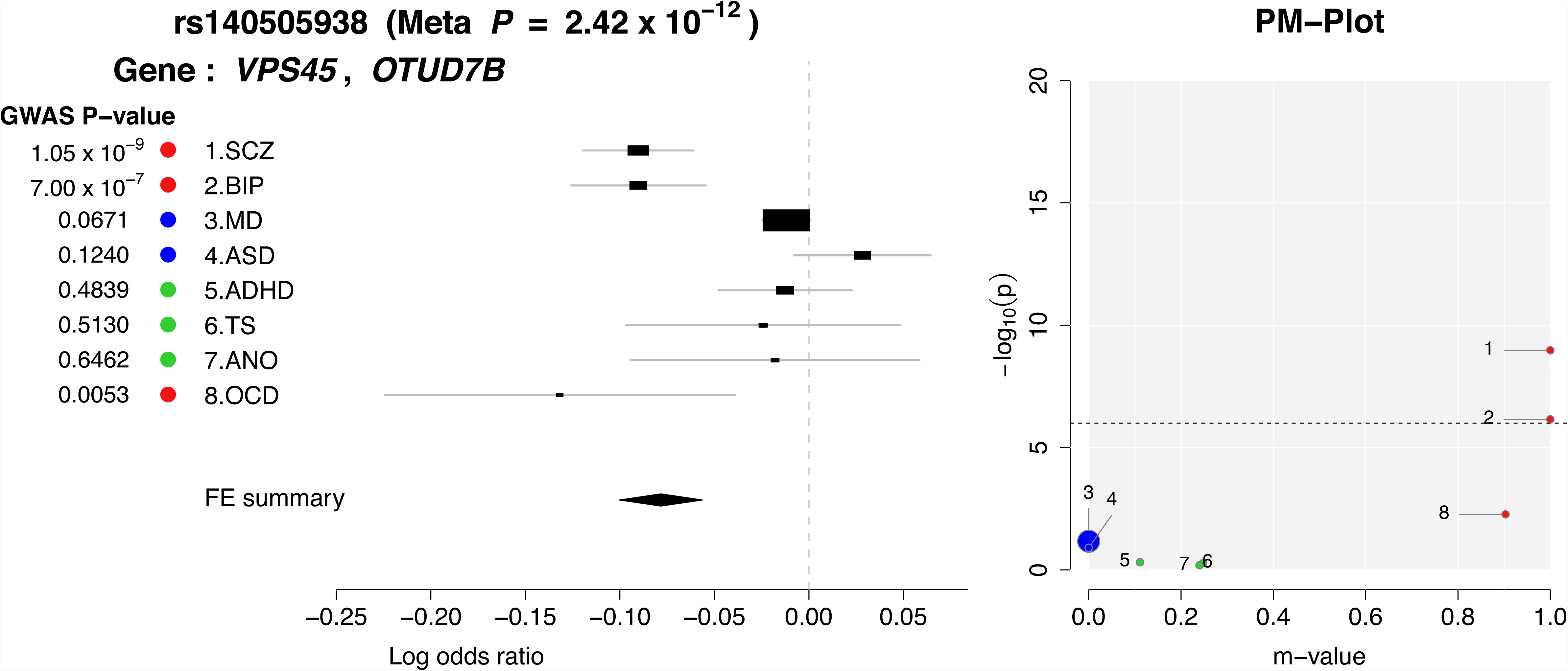

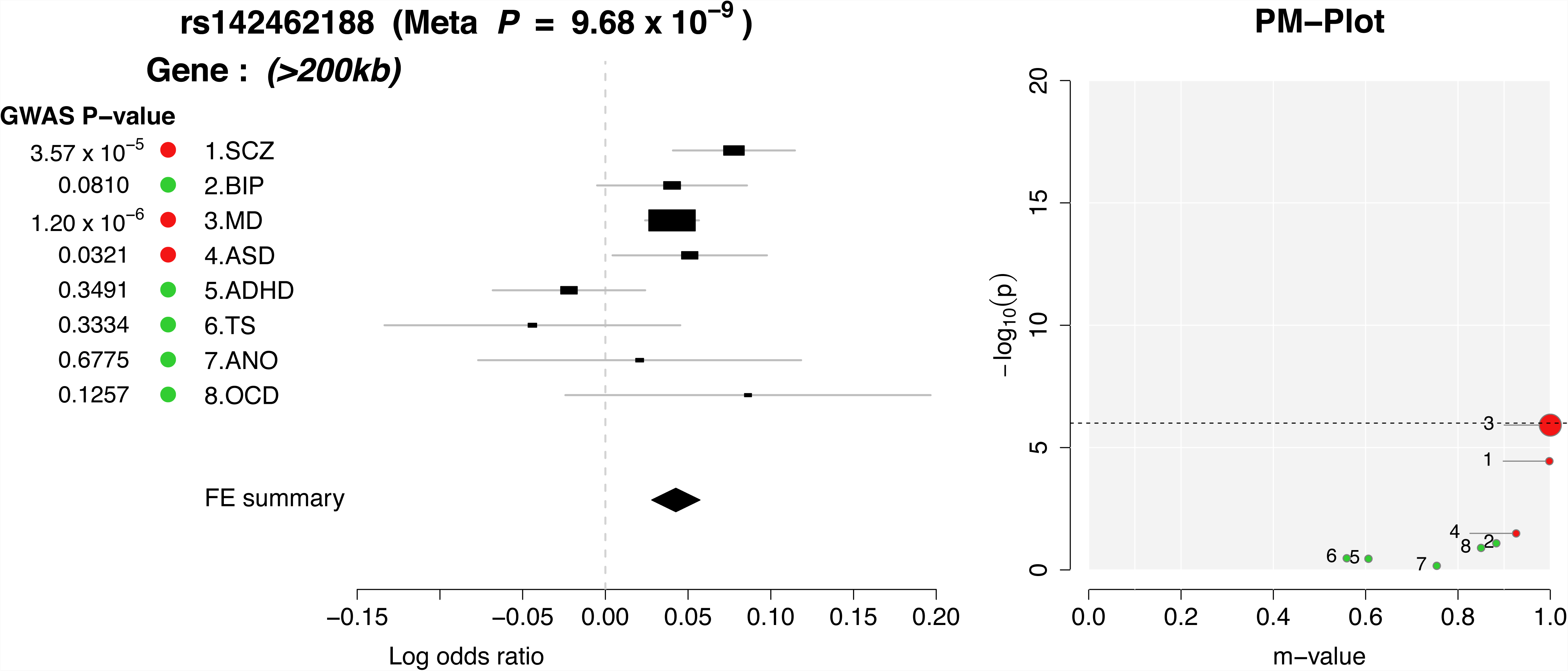

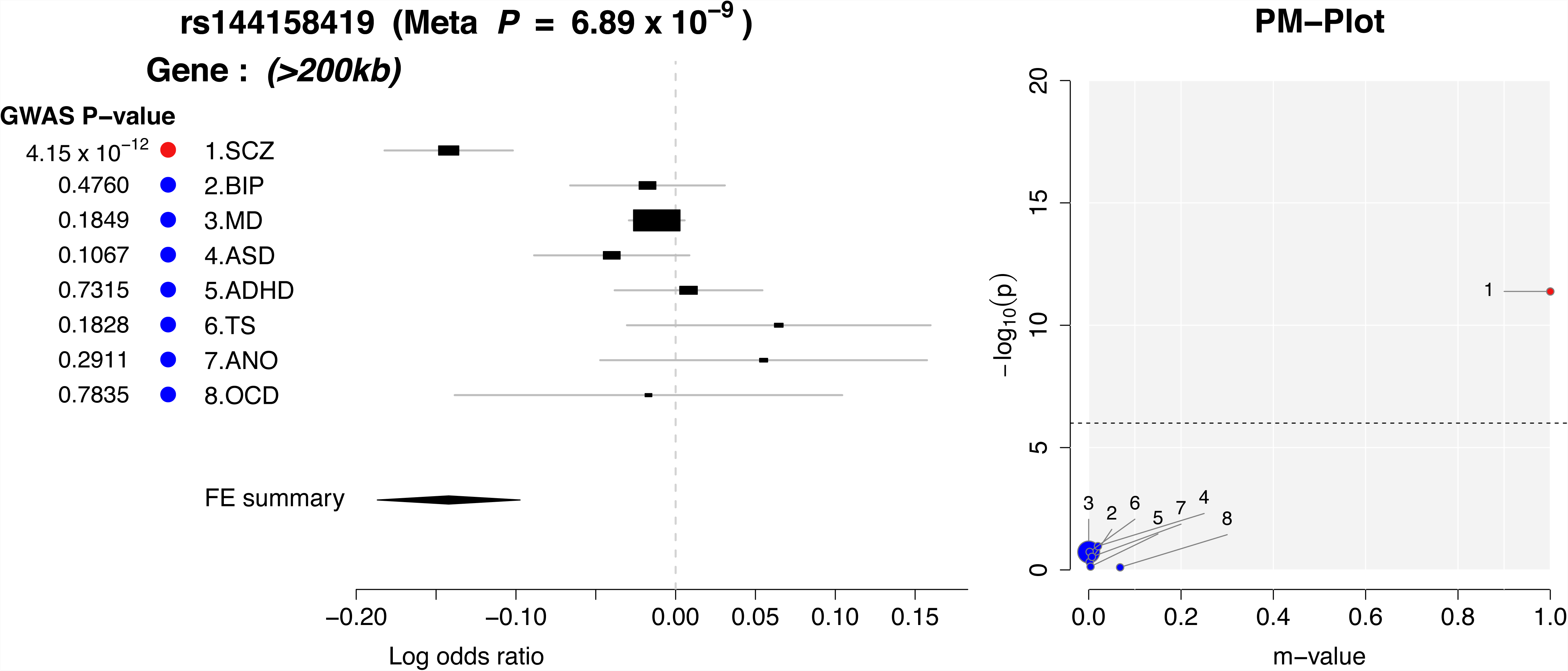

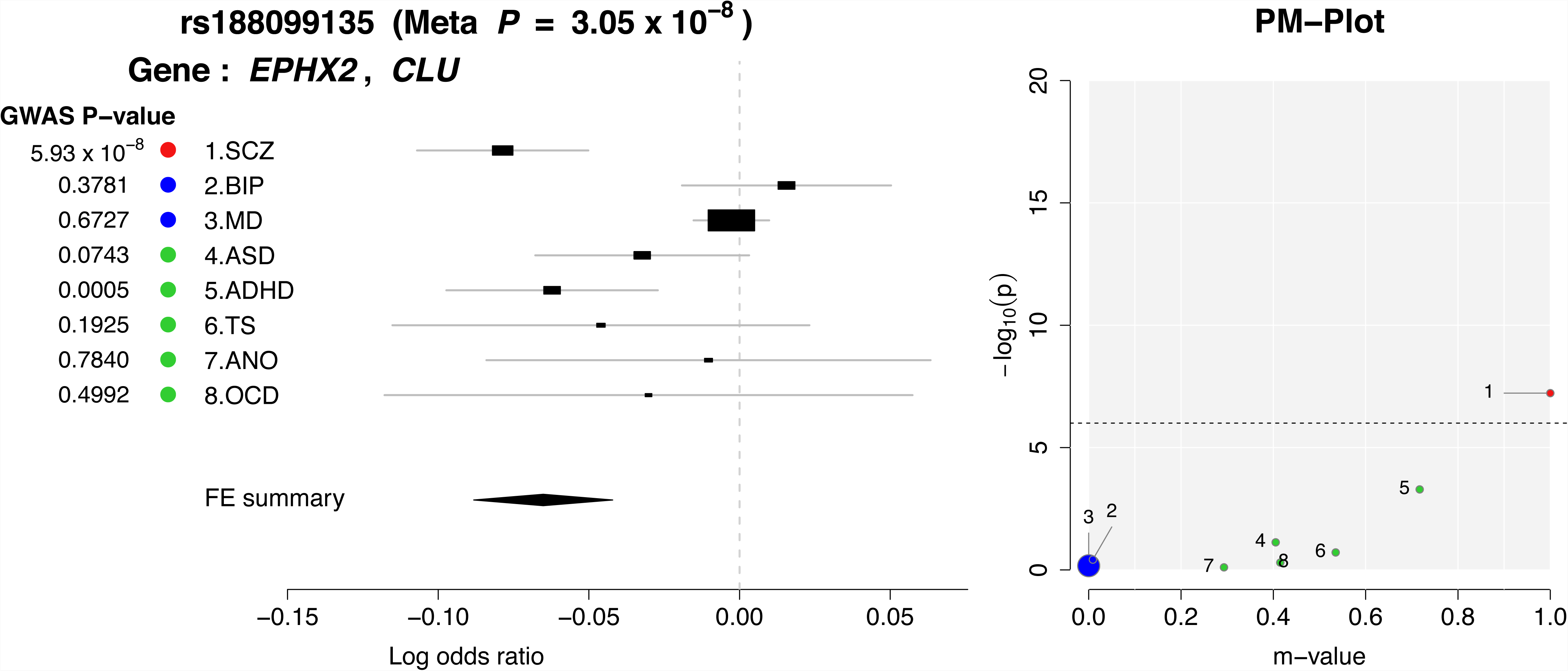

**Figure S3.**
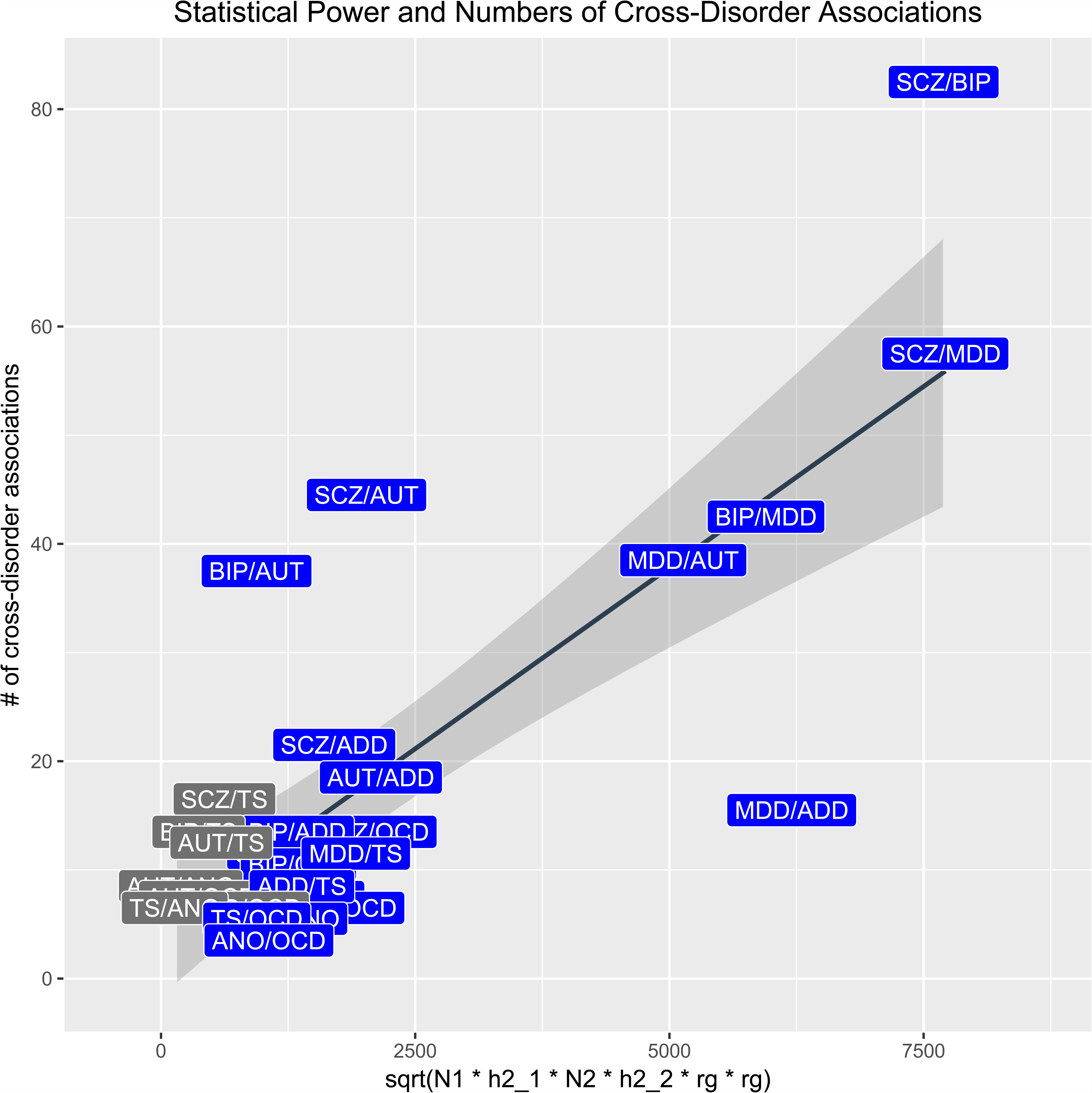
Statistical power and number of cross-disorder associations (Related to Table S4). Power to detect associations across pairs of disorders was plotted with the number of cross-disorder associations identified in the current meta-analysis. For each pair of disorders, power was estimated using the number of cases and heritability for each disorder, as well as the genetic correlation between the disorders. In general, as power increased, so did the number of SNPs identified.

**Figure S4.**
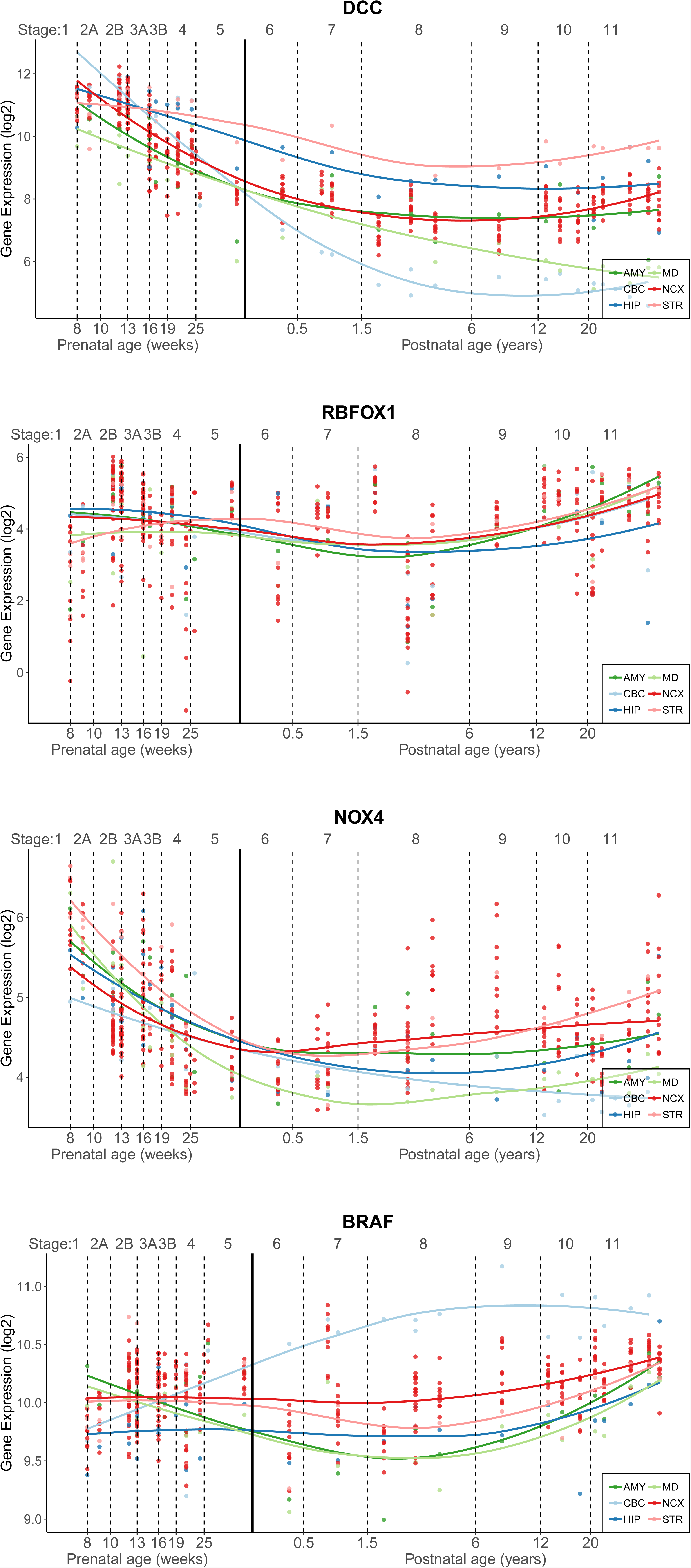
Gene expression of top loci across development. (Related to Figure 3) Gene expression trajectories from a transcriptome atlas of post-mortem brain tissue across development are plotted for four top loci, *DCC, RBFOX1, NOX4* and *BRAF* in six different brain tissue types. AMY = amygdala; MD = mediodorsal nucleus of the thalamus; CBC = cerebellar cortex; NCX = neocortex; HIP = hippocampus; STR = striatum.

**Figure S5.**
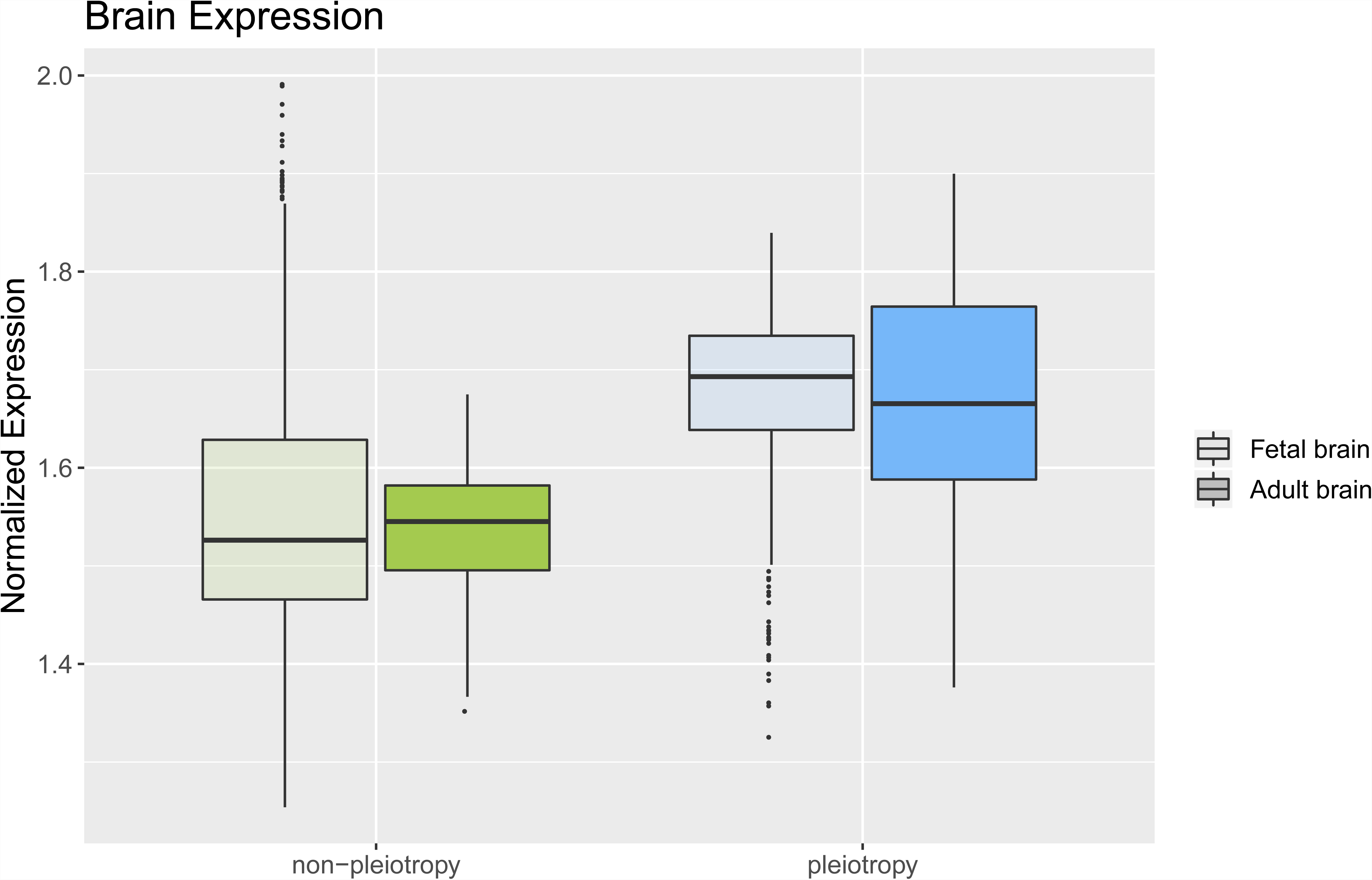
Gene expression in the brain for pleiotropic and non-pleiotropic loci. (Related to Figure 5) Average normalized gene expression in fetal and adult post-mortem brain tissue for pleiotropic (109) and non-pleiotropic (37) loci were plotted. Disorder-specific and pleiotropic risk loci showed a similar level of gene expression in prenatal and postnatal development after multiple testing correction (t-test *p* > 0.025).

## STAR* METHODS

## CONTACT FOR REAGENT AND RESOURCE SHARING

Any inquiries about analytical results or other information should be directed to Phil H. Lee or Jordan W. Smoller.

## EXPERIMENTAL MODEL AND SUBJECT DETAILS

### Genotyped sample description

Genotype data from eight studies of genetic associations with psychiatric disorders conducted by the Psychiatric Genomics Consortium were included in this report. A summary of each study is provided below, however, detailed sample descriptions are available in the primary publication. The lead PI of every cohort included across studies certified that their protocol was approved by their local Ethical Committee. Supplementary Table 1 lists for each disorder the number of cases and controls, the number of loci identified in the single disorder genome-wide association study, and SNP-based heritability.

### Schizophrenia | Ripke et al., 2014

108 loci were identified as associated with schizophrenia in a case-control meta-analysis including 150,064 individuals. For the current study, the 46 case-control cohorts of European ancestry were retained, totaling 33,640 cases and 43,546 controls. Cases were defined as individuals diagnosed with schizophrenia or schizoaffective disorder, which was determined by research-based assessment or clinician diagnosis depending on the sample.

### Bipolar disorder | Stahl et al., 2018

Thirty-two case-control cohorts from Europe, North America, and Australia including 20,352 cases and 31,358 controls of European ancestry were meta-analyzed to identify 30 loci associated with bipolar disorder. Cases met criteria for lifetime diagnosis of bipolar disorder as defined by DSM-IV, ICD-9, or ICD-10, which was established using interview-based structured assessment, clinician-administered checklists, or review of medical records. All subjects in the meta-analysis were included in the current study.

### Major depression | Wray et al., 2018

Seven case-control cohorts were combined to identify 44 loci associated with major depression. The first cohort included 29 case-control samples of European descent where lifetime diagnosis of major depressive disorder was ascertained using structured clinical interviews (DSM-V, ICD-9, ICD-10), clinician-administered checklists, or review of medical records. Six additional cohorts of European ancestry, including the Hyde et al study (23andMe, Inc), determined case status using other methods including national or hospital treatment registers, self-reported symptoms or treatment by a medical professional, or direct interviews. Analyses comparing the original cohort with the additional ones indicated strong correlation of common genetic variants and little evidence of heterogeneity. 130,664 cases and 330,470 controls from these cohorts were included in the current analyses.

### Attention deficit hyperactive disorder | Demontis et al., 2019

Twelve cohorts of European, North American, and Chinese descent were aggregated in a meta-analysis of attention deficit and hyperactive disorder, revealing 12 associated loci. For the first cohort, cases were ascertained using the Danish Psychiatric Central Research Registrar and diagnoses were confirmed by psychiatrists according to ICD-10. The remaining studies included four parent-offspring trio cohorts and seven case-control cohorts. Cases were recruited from clinics, hospitals or through medical registries and diagnosed using research-based assessments administered by clinicians or trained staff. 19,099 cases and 34,194 controls of European ancestry were included in the current study.

### Autism spectrum disorder | Grove et al., 2017

Five family-based cohorts of European descent and a population-based case-control sample from Denmark were combined to discover five loci associated with autism spectrum disorder. In each family study, diagnosis was confirmed for all affected individuals using standard research tools and expert clinical consensus diagnosis. In the population-based cohort, cases were identified using the Danish Psychiatric Central Research Register and were diagnosed with ASD before 2013 by a psychiatrist according to ICD-10. All subjects in this sample were included here (18,381 cases; 27,969 controls).

### Obsessive compulsive disorder | IOCDF-GC and OCGAS, 2018

Individuals of European descent from two cohorts were combined in this meta-analysis including 2,688 cases and 7,037 controls; no loci reached genome-wide significance. Case diagnoses were established using DSM-IV criteria and controls were unscreened. All cases and controls were included in the current analyses.

### Anorexia nervosa | Duncan et al., 2017

3,495 cases from two consortia and 10,982 matched controls from the Psychiatric Genomics Consortium, all of European descent, were meta-analyzed to identify one locus associated with anorexia nervosa. Cases met criteria as defined by DSM-IV for lifetime diagnosis of anorexia nervosa (restricting or binge-purging subtype), bulimia nervosa, or anorexia nervosa – not otherwise specified, anorexia nervosa subtype. All individuals included in the primary study were included in the current analyses.

### Tourette Syndrome | Yu et al., in press, *American Journal of Psychiatry*

Three case-control cohorts and one family-based cohort from Europe and North America including 4,819 cases and 9,488 controls of European ancestry were meta-analyzed to identify one locus associated with Tourette Syndrome. All cases met DSM-IV-TR or DSM-5 criteria for Tourette syndrome, except for 12 cases who met DSM-5 criteria for chronic motor or vocal tic disorder. All cases were recruited by Tourette syndrome specialty clinics or by email/online recruitment combined with validated, web-based phenotypic assessments.

### Genotype quality control, imputation, and association analysis

All primary studies used the standardized PGC ricopili pipeline for quality control, imputation and association testing. Briefly, for each dataset, poor quality SNPs and samples missing >5% SNPs were removed. Next, pre-phasing and imputation were implemented using IMPUTE2 (Howie et al., 2011) and the 1000 Genomes reference panel. High quality SNPs (INFO > 0.8) with low missingness (<1%) were retained. A subset of these markers (MAF > 0.05; pruned for linkage disequilibrium, r^2^ > 0.02) were used to assess relatedness and population stratification. Only one of any pair of related individuals was retained. Each imputed dataset was tested for association with the disease outcome of interest using an additive logistic regression model in PLINK (Purcell et al., 2007) with age, sex, and 10 principal components included as covariates. Finally, a meta-analysis within each disease category was done using an inverse-weighted fixed effects model. After extracting SNPs commonly exist in all eight disorder studies, we removed 3,591 SNPs whose alleles were incompatible. For palindromic SNPs, we compared allele frequencies between eight studies to check strand ambiguity. 50 SNPs with frequency difference greater than 15% from the 1KG reference was excluded. As a result, 6,786,994 autosomal SNPs remained for further analysis.

## QUANTIFICATION AND STATISTICAL ANALYSIS

### Genome-wide SNP-heritability estimation

For each of the eight GWAS disorders, LD Score regression was performed on the summary statistics of individual disease using LDSC to estimate SNP-based heritability in the liability scale and genetic correlation between pairs of disorders (Bulik-Sullivan et al., 2015b). LD scores and weights for European populations were downloaded from the LDSC website (http://www.broadinstitute.org/∼bulik/eur_ldscores/). SNPs were removed if the minor allele frequency is smaller than 5% or an imputation quality score is less than 0.9; MHC region was excluded from the analysis. For single-trait LDSC, the slope of the regression estimates the SNP-based heritability, and the intercept captures the inflation in the summary statistics due to population stratification or other confounding factors. We confirmed that the heritability Z-scores (i.e., a measure of the polygenic signals) are greater than four, and the LDSC intercepts are approximately one and less than *λ*_*GC*_. This suggests that the increase in mean χ2 statistics is mostly due to polygenicity and not due to stratification.

### Factor analysis and genomic SEM

Genomic SEM’s Multivariable LD score regression method (Grotzinger et al., 2018) was first used to estimate the genetic covariance matrix (S) and sampling covariance matrix (V) for the eight psychiatric traits. Quality control for this step included removing SNPs with an MAF < 1%, information scores < .9, SNPs from the MHC region, and filtering SNPs to HapMap3. All SNP effects were standardized using the sumstats function in Genomic SEM. To examine genome-wide factor structure, models using only the genetic covariance and sampling covariance matrix were fit. Genomic SEM provides indices of model fit—standardized root mean square residual (SRMR), model *𝒳*^2^, Akaike Information Criteria (AIC), and Comparative Fit Index (CFI)—that can be used to determine how well the proposed model captures the observed data. Model fit for the common factor model in which the loadings were freely estimated was only fair, (*𝒳*^2^ (20) = 313.94, AIC = 345.9, CFI = .786, SRMR = .149), suggesting that there were nuances in the genetic architecture not fully captured by a single cross-trait index of genetic risk. An exploratory factor analysis (EFA) of the S matrix with three-factors using the promax rotation in the R package factanal was then used to guide construction of a follow-up model (**Supplementary Table 3**). A follow-up confirmatory model with three correlated factors was specified in Genomic SEM based on the EFA parameter estimates (positive standardized loadings > .2 were retained; Figure 2b). This model provided good fit to the data (*𝒳*^2^ (15) = 85.35, AIC = 127.36, CFI = .945, SRMR = .079). Results indicated there was a moderate genetic correlation between the compulsive and mood/psychotic disorders factors (*r*_*g*_ = .43, *SE* = .07), a smaller genetic correlation between the mood/psychotic and early onset factors (*r*_*g*_ = .24, *SE* = .05), and next to no correlation between the compulsive and early onset factors (*r*_*g*_ = < .01, *SE* = .07). A model that included additional negative cross-loadings provided similar fit to the data and highly similar correlations across the genetic factors. Given this consistency in results, the correlated factors model with SNP effects only included positive loadings.

### Summary-data-based meta-analysis

To identify genomic loci shared across multiple neuropsychiatric disorders, we performed primary meta-analysis using the subset-based fixed-effects method ASSET (Bhattacharjee et al., 2012). Standard meta-analysis pools the effect of a given SNP across *K* studies, weighting the effects by the size of the study. In this subset-based meta-analysis, this basic procedure – as well as a null probability distribution for a SNP having no effect in any study – was calculated for all subsets of studies to determine a given SNP’s subset-specific effect. The maximum SNP effect across subsets can be denoted

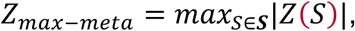

where the absolute value of the subset-specific effect [Z(S)] over class S of all possible subsets of K studies is highest. The numbers of shared subjects across eight disorder studies were identified using the PGC checksum algorithm, and *Z*_*meta*_ was standardized so that covariance between the statistics can be accounted for as previously described (Bhattacharjee et al., 2012; Lin and Sullivan, 2009). Tail probabilities for the distribution of the maximum were then estimated with the discrete local maxima method, which uses the correlation structure of test statistics across subsets. Based on this distribution, a p-value was derived using a one-sided test of the test statistic (Z); each directional test can then be combined using a chi-square method (Fisher’s combined p-value method). Even when correcting for all subset tests (2_*K*_-1), simulations suggest there is a substantial gain in power using this test relative to traditional meta-analysis.

Once SNPs with genome-wide significant association were identified, we identified LD-independent genomic regions using PLINK clumping (–clump-r2=0.4, –clump-kb=500, –clump-p1=5e-08, –clump-p2=5e-02). Genomic regions were merged if they physically overlap using bedtools. Due to extensive LD, the MHC region was considered as one region (chr6:25-35Mb). To detect secondary signals independent of index SNP in each of the candidate cross-disorder loci, conditional analysis was performed with GCTA-COJO (Yang et al., 2012) using meta-analysis summary statistics from ASSET. 1KG EUR population was used as the reference panel for estimating LD. For each genomic region harboring a cross-disorder signal, we tested the presence of any additional associated SNPs using a stepwise procedure (–cojo-slct), conditioning on the primary significant SNP for model initiation. A conditional p-value for each variant was reported, adjusted for genomic control and collinearity. In each region, additional SNPs were selected as a distinct association signal if having a conditional p-value < 1e-06.

### Disease-association modeling

We estimated posterior probabilities for each of the top loci identified from the meta-analysis to quantify disorder-specific effects (Han and Eskin, 2012). This estimation, known as the *m-*value, relies on two assumptions, 1) effects are either present or absent in studies, and 2) if they are present, they are similarly sized across studies. Assume *X*_*i*_ is the observed effect size of study *i*, and *T*_*i*_ is a random variable with value 1 if study *i* has an effect and 0 if not, then the m-value can be estimated using Bayes’ theorem:

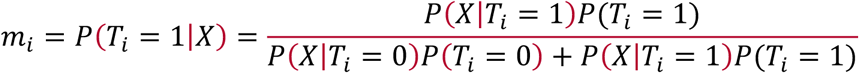

which can then be used to predict whether an effect exists in a given study (>.9) or not (<.1) under the binary effects assumption. For further details, refer to the publication by Han & Eskin (2012).

### Functional annotation and gene-mapping of genome-wide significant variants

For the 146 genome-wide significant variants, gene mapping and functional annotation was conducted using various resources, including SNPNexus (Dayem et al., 2018) and FUMA (Watanabe et al., 2017). Nearest genes and functional consequence of each SNP on gene functions were annotated based on ANNOVAR (Wang et al., 2010). Combined Annotation Dependent Depletion (CADD) score (Kircher, 2014) indexes the deleteriousness of variants computed based on 67 annotation resources. SNPs with the CADD score higher than 12 were considered to confer deleterious effects. The RegulomeDB (Boyle, 2012) provides a categorical score that describes how likely a SNP is likely to play a regulatory role based on the integration of high-throughput datasets. The RDB score of 1a suggests the strongest evidence while the score 7 represents the least support for a regulatory potential. The minChrState and the commonChrState represent the minimum and the most common15-core chromatin state across 127 tissue/cell type predicted by ChrHMM. The chromatin state of less than 8 suggests an open chromatin state. We also performed eQTL mapping, which provides significant *cis*-SNP-gene pairs (up to 1Mb apart) in brain tissue types from GTEx and BRAINEAC.

For chromatin interaction mapping, we first refined the localization of potential causal variants for top 146 lead SNPs. We then used FINEMAP (Benner et al., 2016) to identify credible SNPs. For each region, we considered only SNPs located in the LD region with the lead SNP (r^2^ > 0.6). We then applied the method to calculate the posterior probability of being causal for each of the remaining SNPs. A 95% credible set of SNPs for each region was constructed by ordering the posterior probability from largest to smallest and selecting in the corresponding SNPs up to a cumulative probability of 95%. Credible SNPs were then grouped into those that are located within the promoter or exons and those that are non-coding/intronic. Promoter/exonal SNPs were directly assigned to their target genes using positional mapping, while non-coding/intronic SNPs were assigned to their target genes based on long range interactions (Hi-C) or expression quantitative trait loci (eQTLs). Two Hi-C datasets originated from the human brain (fetal brain Hi-C (Won et al., 2016) and adult brain Hi-C (Wang et al., 2018)) were used to map credible SNPs as previously described (Wang et al., 2018). A colocalization analysis with the recent eQTL dataset from adult prefrontal cortices (PFC) was also used to map 146 GWS loci into their target genes (Wang et al., 2018). In the end, we obtained two sets of candidate genes, one from fetal brain (positional mapping, fetal brain Hi-C), the other from adult brain (positional mapping, adult brain Hi-C, adult brain eQTLs).

### GTEx gene expression enrichment analysis

MAGMA gene-property analysis (de Leeuw et al., 2015) was performed using gene expression data from 83 tissues based on GTEx RNA-seq data (v7). Expression values (RPKM) were log2 transformed with pseudo-count one after winsorization at 50, and average expression values were taken per tissue. Analysis was performed separately for 30 general tissue types and 53 specific tissue types.

### Pathway analysis using Gene Ontology

We used FUMA (Watanabe et al., 2017) to map SNPs to genes and then test for enrichment of specific Gene Ontology functions and pathways among genome-wide significant pleiotropic and disorder-specific SNPs separately. Hypergeometric tests identify any statistical over-representation of genes from the input list (mapped from SNPs) in predefined Gene Ontology gene sets which describe biological processes, molecular functions, and cellular components. Multiple test correction was applied by category.

### Enrichment analysis using brain developmental, regional, and cell-type-specific data

Developmental expression trajectories for candidate genes were plotted using a published transcriptome atlas constructed from post-mortem brain data (Kang et al. 2011). Expression values were log-transformed and centered using the mean expression values for all brain expressed genes. Mean expression values for candidate genes were plotted across prenatal (6-37 weeks post-conception) and postnatal (4 months to 62 years) developmental stages. We used candidate genes identified in fetal brain and adult brain to plot prenatal and postnatal gene expression profiles, respectively.

To obtain genes that show cortical regional enrichment (e.g. frontal cortical enrichment), we computed t-statistics for each gene for a specific cortical region (e.g. frontal cortex) versus all other cortical regions (e.g. parietal cortex, temporal cortex, and occipital cortex, Kang et al. 2011). Top 5% of genes that show specific expression patterns for each cortical region were selected as region-specific genes. These genes were then overlapped with candidate genes by Fisher’s exact test to measure cortex regional enrichment.

Single cell expression profiles from the adult brain (Darmanis et al., 2015) were used to identify cell-type specificity of candidate genes. Single cell expression values were log-transformed and centered using the mean expression values. Average centered expression values for candidate genes were calculated in each cell. Cells were then grouped into cell clusters (neurons, astrocytes, microglia, oligodendrocytes, OPC, and endothelial cells), and a relative expression level for a given cell cluster was calculated by a *scale* function in R.

Developing neural cell-type enrichments were estimated using expression profiles of single-cells taken from fetal cortical laminae. Cell-type specific genes were selected according to a significant Pearson correlation (FDR < 0.05) between the gene and an idealized cluster marker for each cell-type, following the approach described in original publication. Candidate gene enrichment for each set of specifically expressed genes was estimated by logistic regression and adjusting for gene length.

### Comparison with other brain-related traits and diseases

To look for other traits that show an association with our top loci, we examined publicly available GWAS summary statistics of 11 brain-related and 10 non-brain-related traits and disorders. The selected brain-related traits include Alzheimer’s disease (Lambert et al., 2013), anxiety (Otowa et al., 2016), all epilepsies (International League Against Epilepsy Consortium on Complex Epilepsies. Electronic address, 2014), genetic generalized epilepsy (International League Against Epilepsy Consortium on Complex Epilepsies. Electronic address, 2014), non-acquired focal epilepsy (International League Against Epilepsy Consortium on Complex Epilepsies. Electronic address, 2014), intracerebral hemorrhage (Woo et al., 2014), post-traumatic stress disorder (Duncan et al., 2018), ischemic stroke (all) (Malik et al., 2016), cardioembolic stroke (Malik et al., 2016), large-vessel disease(Malik et al., 2016), and small vessel disease(Malik et al., 2016), non-brain-related traits include height (Wood et al., 2014), obesity (Berndt et al., 2013), Crohn’s disease (Goyette et al., 2015; Liu et al., 2015), inflammatory bowel disease (Goyette et al., 2015; Liu et al., 2015), ulcerative colitis (Goyette et al., 2015; Liu et al., 2015), coronary artery disease (Nikpay et al., 2015), sleep duration(Jones et al., 2016), number of children ever born (Barban et al., 2016), subject well-being (Okbay et al., 2016), and neuroticism (Okbay et al., 2016). Data source and sample size information for each individual GWAS can be found in Table S16. All the summary results files were harmonized before use, with reference and effect alleles matched up across studies for each index SNP. We then extracted the association statistics of the lead SNPs—effect sizes with standard errors (or Z-scores) and p-values—with each of the traits listed herein. Where the index variant was not available, the nearest proxy SNP (r^2^ > 0.8) was used for reporting the association.

## DATA AND SOFTWARE AVAILABILITY

The Psychiatric Genetics Consortium (PGC)’s policy is to make genome-wide summary results publicly available. Summary statistics for a combined meta-analysis of eight psychiatric disorders without 23andMe data are available on the PGC web site (https://www.med.unc.edu/pgc/results-and-downloads). Results for 10,000 SNPs for eight disorders including 23andMe are also available on the PGC web site. The summary-level GWAS association statistics for PGC individual disorders are available at the website (https://www.med.unc.edu/pgc/results-and-downloads).

GWAS summary statistics for the 23andMe cohort (Hyde, 2016) must be obtained separately. These can be obtained by individual researchers under an agreement with 23andMe that protects the privacy of the 23andMe participants. Contact Aaron Petrakovitz to apply for access to the data.

## Supplementary Tables

**Table S1. Summary of eight neuropsychiatric disorder datasets (Related to Table 1).** Disease-specific cases and controls included in the meta-analysis, number of individual GWAS loci, and liability-based SNP heritability estimates are provided. Heritability was estimated from available European summary statistics using LD score regression.

**Table S2. Genetic correlations estimated by LD score regression. (Related to Figure 1)** Genetic correlations and standard errors between pairs of disorders were estimated from European GWAS summary statistics using LD score regression.

**Table S3. Results from EFA of genetic covariance matrix (Related to Figure 1 and Figure S1)**We estimated the genetic covariance matrix across eight psychiatric traits using genomic SEM and then performed an exploratory factor analysis, which revealed three factors. Individual disease loadings on each factor are listed as well as correlations between factors.

**Table S4. List of 146 lead SNPs (Related to Figure 3 and Figure S2)** The cross-disorder fixed-effects meta-analysis identified 136 loci and within these loci, multi-SNP-based conditional analyses identified 10 additional SNPs. The details of these 146 lead SNPs, as well as the posterior probability of disease-specific associations estimated using a Bayesian statistical framework are listed.

**Table S5. Disorder-specific association (Related to Figure 3 and Figure S2)** Disease-specific associations are listed for all 146 lead SNPs.

**Table S6. Loci with opposite directional effects (Related to Figure 4)** SNPs with a cross-disorder meta-analysis *p* ≤ 1×10^−6^ that showed disorder-specific effects in opposite directions are listed along with the effects for each disease.

**Table S7. Functional annotation of 146 lead SNPs (Related to Table S4)** Functional annotation for the genome-wide significant SNPs was done using SNPNexus, FUMA, and ANNOVAR. SNPs were mapped to the nearest gene within 100kb when possible and various functional consequences were annotated including combined annotation dependent depletion (CADD) scores, which reflect the deleteriousness of variants based on 67 annotation resources, and likelihood SNP plays a regulatory role based on the RegulomeDB (1 = strongest; 7 = weakest).

**Table S8. Brain eQTL and Hi-C data annotation based on FUMA database (Related to Table S4)** Genome-wide significant SNPs were tested for any eQTL effects in brain tissues. Significant *cis*-SNP-gene pairs (up to 1Mb apart) identified in GTEx (v7) or BRAINEAC are listed along with associated p-values. Additionally, fine-mapping for the 146 lead SNPs was done using an efficient summary statistics-based Bayesian method to identify a 99% credible set of SNPs for each region. These SNPs were used to identify long-range interactions with the regions harboring a candidate SNP using brain Hi-C datasets. The Hi-C tissue, regions, and p-values were reported.

**Table S9. GWAS catalog data for lead SNPs (Related to Table S4)** The NHGRI-EBI GWAS Catalog was queried for each SNP to identify all traits previously associated with a lead SNP (p-value < 1×10^−5^), along with the corresponding effect sizes and p-values.

**Table S10. GTEx gene enrichment analysis using MAGMA (Related to Figure 5)** To investigate enrichment of lead SNPs within certain tissue types, we performed a MAGMA gene-property analysis using gene expression data from 83 tissues based on GTEx RNA-seq data (v7). Enrichment statistics for each tissue are listed.

**Table S11. Tissue enrichment analysis for pleiotropic risk loci using MAGMA (GTEx v7) (Related to Figure 5)** To evaluate pleiotropic-specific effects, we assessed enrichment for the 109 pleiotropic lead SNPs using MAGMA and 83 tissues based on GTEx v7. The statistics for each tissue are listed.

**Table S12. Tissue enrichment analysis for disorder-specific risk loci using MAGMA (GTEx v7) (Related to Figure 5)** To evaluate disorder-specific effects, we assessed enrichment for non-pleiotropic lead SNPs using MAGMA and 83 tissues based on GTEx v7. The statistics for each tissue are listed.

**Table S13. Gene ontology analysis for pleiotropic loci (Related to Result - *Functional characterization of pleiotropic risk loci*)** We used Gene Ontology data to identify any enrichment among the 109 pleiotropic SNPs for specific biological processes (GO_bp), molecular functions (GO_mf), or cellular components (GO_cc). Gene sets for which significant enrichment was identified are listed along with raw and adjusted p-values.

**Table S14. GO enrichment analysis for disease-specific risk loci (Related to Result - *Functional characterization of pleiotropic risk loci*)** We used Gene Ontology data to identify any enrichment among the 37 disease-specific loci for specific biological processes (GO_bp), molecular functions (GO_mf), or cellular components (GO_cc). Gene sets for which significant enrichment was identified are listed along with raw and adjusted p-values.

**Table S15. GWAS Catalog enrichment analysis for top genome-wide significant loci (Related to Result - *Functional characterization of pleiotropic risk loci*)** For traits in the NHGRI-EBI GWAS Catalog, we mapped genome-wide significant loci to genes and then assessed the overlap between genes for a given trait and genes linked to the top loci for the current meta-analysis. We calculated p-values associated with the degree of overlap relative to what would have been expected by chance.

**Table S16. Comparison of disease-specific vs. pleiotropic risk loci by various functional and genomic features (Related to Result - *Functional characterization of pleiotropic risk loci*)** Functional and genomic features were compared between pleiotropic and non-pleiotropic loci. Statistics from t-tests, as well as ANOVAs, comparing these two groups were reported.

**Table S17. Genetic correlation analysis of top cross-disorder loci with 28 brain-related traits (Related to Result - *Relationship between cross-disorder genetic risk and other brain-related traits and diseases***) We used LD score regression to estimate genetic correlations between cross-disorder risk and 28 other publicly available brain-related traits. These effects, along with data source and sample size information for each individual GWAS were listed.

**Table S18. GWAS catalog enrichment analysis for pleiotropic risk loci (Related to Result - *Relationship between cross-disorder genetic risk and other brain-related traits and diseases***) We used Gene Ontology data to identify any enrichment among the 109 pleiotropic SNPs for brain-related traits. Gene sets for which significant enrichment was identified are listed along with raw and adjusted p-values.

